# PHI: a Galaxy-based workflow for reproducible prophage-host interaction analysis and standardized viral-genomics reporting

**DOI:** 10.64898/2025.12.02.691814

**Authors:** Joao Pedro Saraiva, Felipe Borim Corrêa, Matthias Bernt, Nawras Ghanem, Esteban Nieto, Rodolfo Brizola Toscan, Lukas Y. Wick, Antonis Chatzinotas

**Affiliations:** Helmholtz Centre for Environmental Research - UFZ, Department of Applied Microbial Ecology, 04318 Leipzig, Germany; Helmholtz Centre for Environmental Research - UFZ, Department of Computational Biology and Chemistry, 04318 Leipzig, Germany; Department of Biochemistry and Biotechnology, University of Thessaly, Viopolis, 41500, Larissa, Greece; Małopolska Centre of Biotechnology, Jagiellonian University, Kraków, Poland; Institute of Biology, Leipzig University, 04103 Leipzig, Germany; German Centre for Integrative Biodiversity Research (iDiv) Halle-Jena-Leipzig, 04103 Leipzig, Germany

**Author notes:** Authors contributed equally to this study (Joao Pedro Saraiva, Felipe Borim Corrêa). Corresponding authors: Joao Saraiva,; Antonis Chatzinotas.

**Keywords:** Bioinformatics, galaxy-server, virology, host-phage interactions

## Abstract

**Background:** Viruses that infect bacteria, known as bacteriophages or phages, are widespread in nature and play important roles in shaping microbial communities and ecosystem functions. Some phages can integrate into bacterial genomes as “prophages”, where they may influence the biology of their host by carrying genes that affect metabolism, virulence, or environmental adaptation. Despite their importance, studying prophages and their interactions with bacterial hosts remains challenging because it typically requires combining many complex computational tools and can be resource-intensive.

**Results:** In this study, we introduce the Prophage-Host Interaction Toolkit (PHI), a user-friendly and automated workflow available through the Galaxy platform. PHI brings together multiple established tools into a single, reproducible pipeline that identifies candidate prophages, evaluates their quality, predicts host relationships, and characterizes key functional genes. Importantly, all results are summarized in an interactive report that simplifies interpretation. When applied to a mock community composed of 22 bacteria as a workflow demonstration, PHI detected 41 prophages across 14 hosts, classifying them into high- and medium-quality phage genomes. Host assemblies exhibited > 99 % completeness and < 1 % contamination for most genomes, while DefenseFinder revealed between 3 and 24 antiviral systems per genome.

**Conclusions:** By removing installation barriers and consolidating the outputs of multiple established tools, PHI lowers the barrier to advanced phage analysis, enabling both specialists and non-experts to explore phage-host interactions and their implications in areas such as microbiome research, biotechnology, and environmental science.

## Background

Phages are viruses that infect bacterial cells. After infection, lytic viruses can replicate while lysogenic phages can integrate their genome into the chromosome of their bacterial hosts [1]. Prophages often carry auxiliary metabolic genes (AMGs) that affect host metabolic and broader ecosystem functions [2, 3]. The automated discovery and comprehensive profiling (e.g., taxonomic assignment, functional annotation, and life-cycle prediction) of phages within host genomes remains challenging due to the complexity of building bioinformatic pipelines and useful summaries and visualizations of their results. Existing workflows typically focus on either identification and gene annotation [4, 5] or specific interaction aspects [6], lacking a comprehensive approach that addresses a wide spectrum of phage-host relationships within genomic data.

Several methods have advanced the field of phage identification and host prediction, such as PHASTEST [5], VirSorter2 [7], and CHERRY [8] but they often provide limited information on both host and phage perspectives, require extensive computational resources, or lack user-friendly interfaces, thereby limiting their use by the scientific community.

We present the Prophage-Host Interaction Toolkit (PHI), a comprehensive and automated framework within the Galaxy [9] platform (e.g., publicly usable via https://usegalaxy.eu), for the identification and profiling of phages within host genomes that unifies established tools into a reproducible workflow. PHI overcomes the limitations of existing tools, such as the complexity and difficulty that arise with installing and running multiple tools. PHI also compiles all results in a single report, facilitating knowledge extraction and enabling researchers from diverse backgrounds to benefit from bioinformatics applications without requiring extensive technical training.

Key features of PHI include its modular architecture, allowing for easy updates and customization. PHI incorporates tools for reproducible detection and contextual profiling of candidate prophages, identification of phage-host pairs, and analysis of CRISPR spacer relationships. Furthermore, it provides comprehensive visualization tools for depicting phage genomic features and host interaction networks, facilitating the interpretation of complex data.

The following sections describe the PHI toolkit, including the integrated tools and its application to a mock-community case study, demonstrating its utility for reproducible workflow execution and integrated reporting.

### Implementation

#### Workflow

The workflow is implemented in Galaxy [9] and it is publicly available through the Galaxy Europe server (https://usegalaxy.eu/u/maze/w/phi-toolkit). To achieve the full functionality described in this manuscript, we pre-tailored the pipeline with established tools and analyses that generate an HTML report through an R Markdown script. Additionally, the flexibility of Galaxy allows for full customization of the pipeline. Any modification or expansion of the pipeline is possible as long as the desired tools are available on the server and the code used to generate the report is updated.

#### Pipeline description

A simplified schematic of the workflow is shown in Figure 1. A more detailed workflow schema produced by Galaxy’s workflow editor is available in the Additional File 1: Figure S1). The PHI pipeline accepts bacterial genomes in FASTA format as input files. These genomes are submitted in parallel to the following host-analysis tools: CheckM2 1.0.2+galaxy1 using the CheckM diamond DB [10] to assess the quality, GTDB-Tk 2.4.1+galaxy0 using the full DB release 220 [11, 12] to assign taxonomy, Defense-Finder 2.0.1+galaxy1 using models 2.0.2 [13] to detect known anti-phage systems, and finally geNomad 1.11.1+galaxy0 using reference data version 1.9 [14] to predict and annotate prophages. Once a genome has been processed and prophages have been identified by geNomad, several virus-analysis tools also run in parallel, which are: (i) CheckV 1.0.3+galaxy0 using reference data version 1.5 [15] to evaluate the quality of prophage genomes, (ii) dRep compare 3.6.2+galaxy1 using an ANI threshold of 0.95 [16] to cluster and compare prophage genomes recovered from the same host, thereby distinguishing identical from distinct prophage sequences, (iii) ABRicate 1.0.1 [17] using the Virulence Factor Database (VFDB) [18] to identify virulence genes in the predicted prophage genomes with the, (iv) iPHoP 1.3.3+galaxy0 [6] to predict other potential hosts and (v) VIBRANT 1.2.1+galaxy2 using reference data version 1.2.1 [19] to identify auxiliary metabolic genes (AMGs) in the prophages. Unless stated otherwise, default parameters were used. The dRep ANI threshold to form secondary clusters was set to 0.95 instead of the default 0.99 to decrease clustering stringency. The database used by ABRicate was set to VFDB instead of the default Resfinder since we are interested in virulence factors. Technically, CheckV and VIBRANT can also predict prophages, but we decided to restrict this task to geNomad due to its regular and active maintenance and to avoid redundancy in predictions. The workflow concludes with the automated report generation via R Markdown, producing a structured HTML summary (Additional File 2: Figure S2). The generated HTML report is organized into multiple thematic sections that integrate host, prophage, and interaction-level analyses. In addition to summary tables of genome quality, taxonomy, and prophage content, the report includes interactive visualizations such as host-virus association networks, CRISPR spacer-protospacer matching outputs, and genome-level feature maps. These components allow users to explore relationships between prophages and their predicted hosts, assess host range, and evaluate genomic context in a unified framework. Tool versions, parameters, and database releases are documented inside the workflow run to ensure reproducibility.

**Figure 1.**
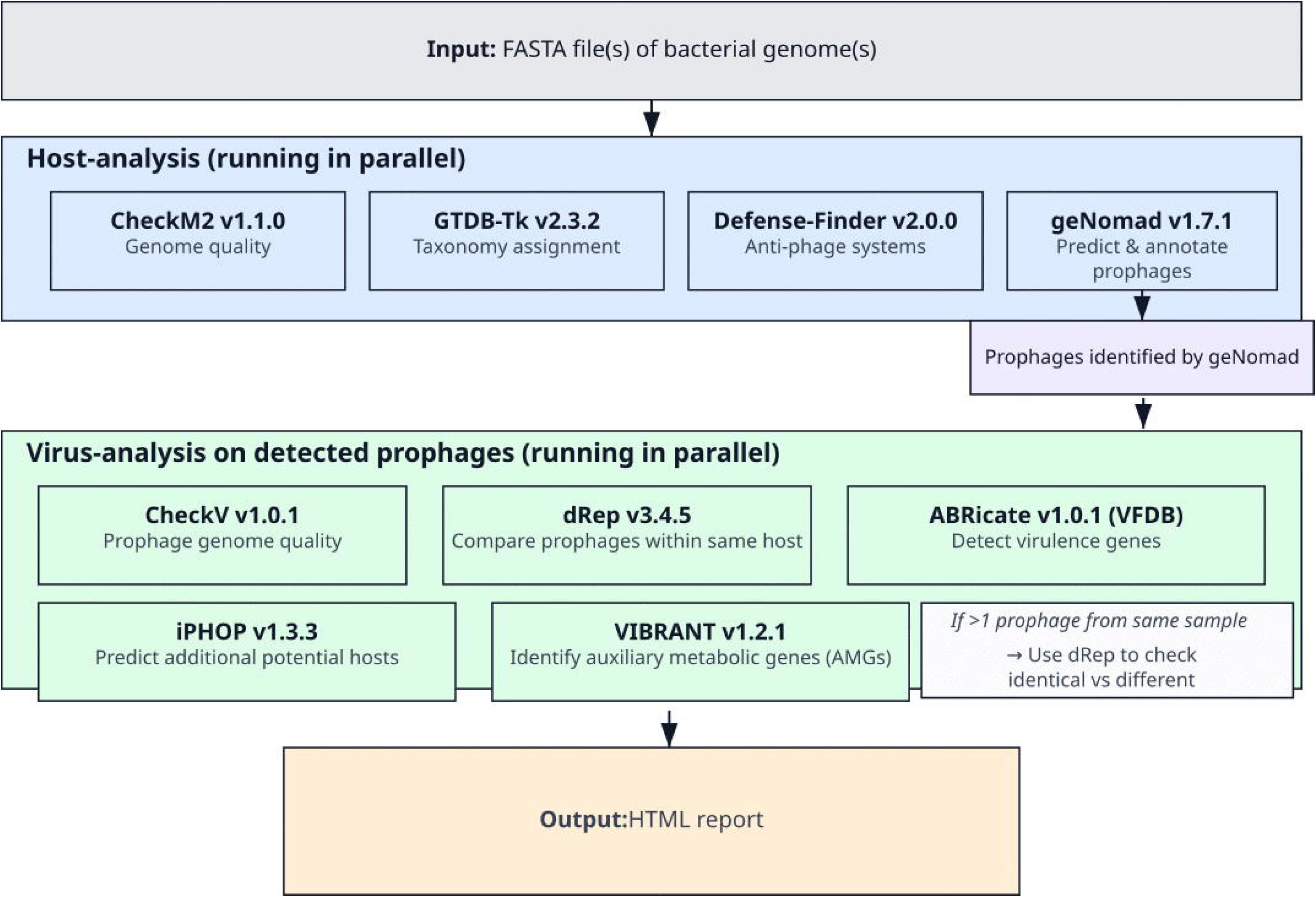
Workflow of PHI toolkit. The user submits one or more FASTA files for processing. The pipeline begins with the parallel execution of the host analysis tools for each submitted genome. Prophages identified by geNomad [14] are then submitted to several virus analysis tools (which are also executed in parallel). If no candidate prophage or viral sequence is identified for a genome, downstream viral analyses are skipped for that genome and the final report records the absence of viral outputs. To generate the HTML report, a script written in R Markdown renders an interactive and user-friendly HTML document. This script first loads several key outputs for each tool, plus the input host genomes. Host genome analysis includes: the geNomad summary table (*_virus_summary.tsv), which reports detected prophages within bacterial genomes; the CheckM2 quality report (quality_report.tsv), which contains completeness and contamination estimates; the taxonomic classification from GTDB-Tk (gtdbtk.bac120.summary.tsv); and the predicted defense systems identified by DefenseFinder (*_defense_finder_systems.tsv). Prophage genome analysis includes: the VIBRANT annotations of auxiliary metabolic genes (VIBRANT_AMG_individuals_*_virus.tsv); the CheckV quality summary file (quality_summary.tsv); the host prediction results from iPHoP (Host_prediction_to_genome_m90.csv); the prophage clustering and comparison output from dRep (Cdb.csv) and the clustering dendrogram (Primary_clustering_dendrogram.pdf); and the virulence factor annotations generated by Abricate (*_virus_vfdb.tsv).

#### Workflow decision logic and edge-case handling

PHI first runs host-level analyses on all submitted genome FASTA files and then triggers downstream viral analyses only when geNomad reports candidate viral or prophage sequences. If no candidate viral sequences are recovered for a genome, the downstream viral modules (CheckV, dRep, ABRicate/VFDB, iPHoP, and VIBRANT) are skipped for that genome and the final report indicates that no viral output was available. If only one or too few viral sequences are available, dereplication or clustering outputs are not interpreted as comparative evidence. Missing module outputs are reported as unavailable rather than silently omitted. These rules make the workflow transparent and reproducible, but they do not modify the underlying predictions generated by each integrated tool.

To calculate the quality classes for host genomes that are displayed on the Overview Table, we used the quality definitions described by the Minimum Information about a Single Amplified Genome (MISAG)/ Minimum Information about a Metagenome-Assembled Genome (MIMAG) standards [20]. ‘High-quality draft’ indicates that a Single Amplified Genome (SAG) or Metagenome-Assembled Genome (MAG) is >90% complete with less than 5% contamination. ‘Medium-quality draft’ SAGs and MAGs are those genomes with completeness estimates of ≥50% and less than 10% contamination. All other SAGs and MAGs (<50% complete with <10% contamination) should be reported as ‘low-quality drafts’ [21]. The quality score is defined as completeness – 5* contamination [22].

### Demonstration dataset

To demonstrate workflow execution, output integration, and report interpretability, we applied PHI to 22 bacterial genomes from a defined mock community dataset [23]. This community is routinely used as a baseline for testing bioinformatics analyses and data interpretation. As we were focusing on phages, we downloaded the FASTA sequences of the complete genomes of 22 bacterial strains from the National Centre for Biotechnology Information (NCBI), except for the reference genome NC_010067.1, which was marked as "Record suppressed". The mock community is hereafter named as MBARC22. The accession numbers for each genome are shown in the HTML report (Additional File 2: Figure S2).

## Results

### Workflow-level runtime and resource usage

The total runtime for the MBARC22 dataset was approximately 3.6 hours. GTDB-Tk required the highest amount of CPU cores (32). In terms of memory, iPHoP was the most demanding, requiring over 130 GB of RAM. In contrast to CheckM and GTDB-Tk which processes all genomes in a single run, all other tools process each genome separately. Of the latter tools, geNomad required the longest time to process a genome on average (21 minutes) while Abricate was the quickest (nine seconds). The average memory peaks and runtimes are shown in Table 1.

**Table 1.**
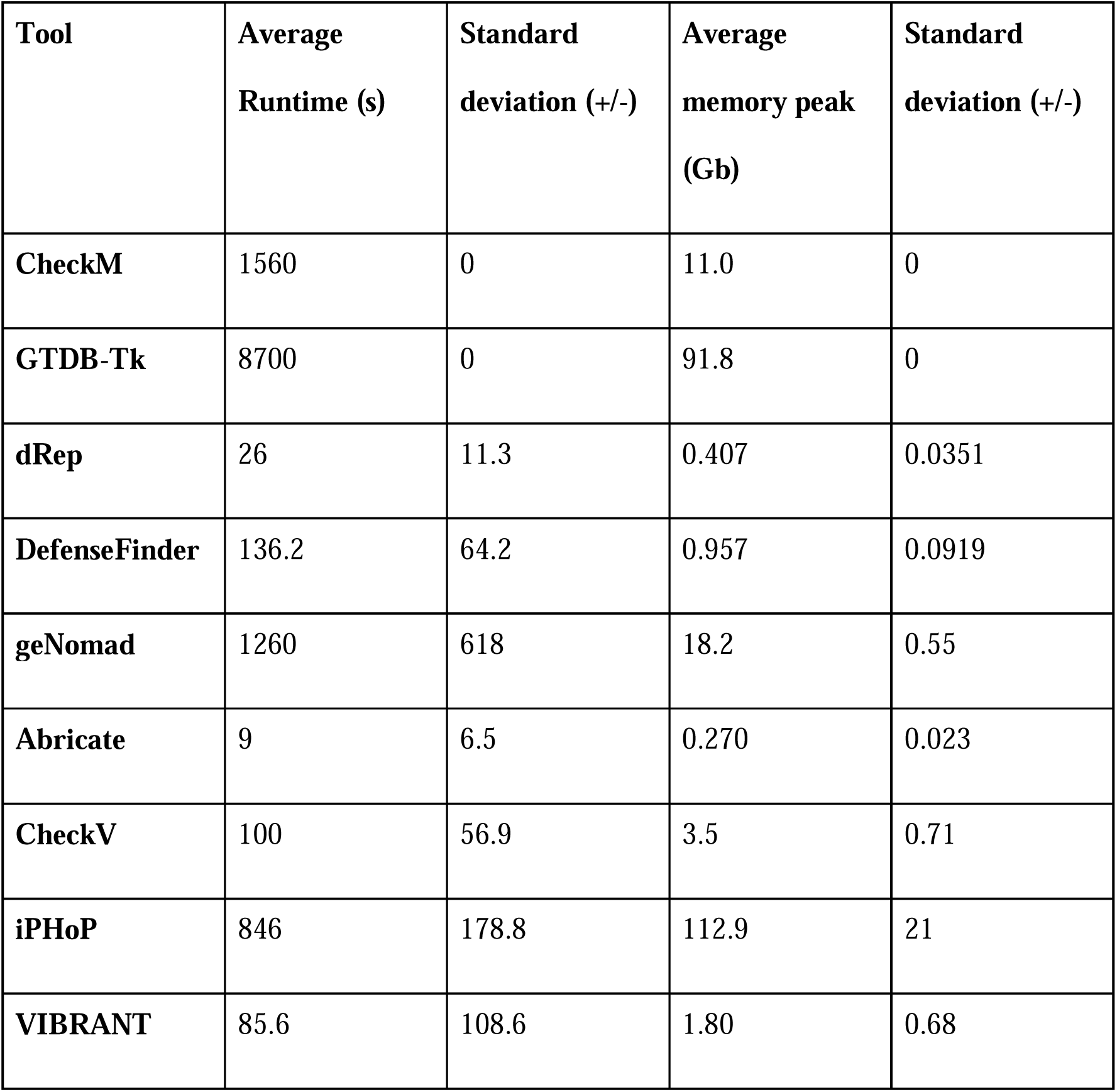
Average computational resources used by each tool for processing MBARC22.

The complete values of memory peaks and runtime per run are available in Additional File 3: Table S1. Note that the resource requirements exceed the resources typically available to researchers. Additionally, this resource-use analysis should be interpreted as computational profiling of the workflow rather than as a predictive-performance benchmark of the underlying tools.

To further evaluate scalability, we performed an additional analysis in which individual tool runtimes were grouped by input file-size intervals, and mean runtime values were calculated for each interval (Additional File 4: Table S2). This analysis showed that runtime did not increase linearly with file size for most tools. Lightweight tools such as ABRicate remained below one minute on average across the tested intervals, whereas tools involving larger database searches or more complex sequence characterization, including geNomad, CheckV, and DefenseFinder, showed greater runtime variability. This suggests that runtime is influenced not only by input file size, but also by sequence composition, contig structure, number of detected viral regions, and fixed initialization costs. iPHoP showed relatively stable runtime across the tested small-file intervals but remained the most memory-intensive step, indicating that memory availability rather than runtime is the main practical constraint for full PHI analyses. Batch-level tools such as GTDB-Tk and CheckM2 should be interpreted separately because they process the complete submitted genome set simultaneously rather than individual genome files.

Implementation in Galaxy improves accessibility because usegalaxy.eu provides public access to the required computational infrastructure.

### Host genome analyses

#### Completeness and Contamination

Except for NC_014008 (14.1% completeness), most bacterial genomes were classified as high-quality drafts (>90% completeness, <5% contamination) (Figure 2).

**Figure 2.**
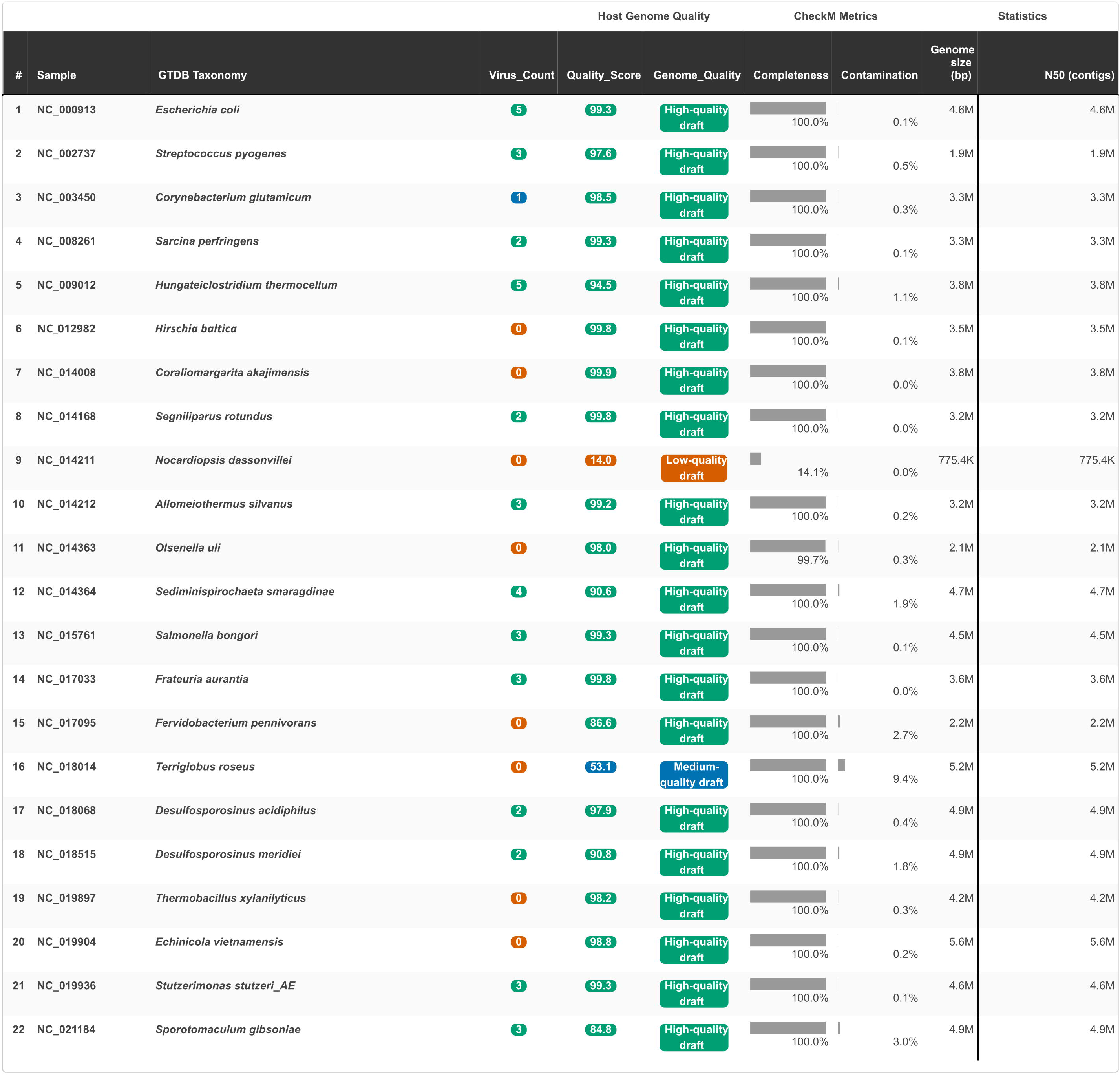
Visualization of the HTML report showing a summary of host genome metrics. The table shows the sample IDs, the taxonomy assignment, the number of viruses identified in the sample, quality metrics such as completeness and contamination, and genome statistics.

#### Detection and Distribution of Antiviral Defense Systems Across Samples

PHI integrated DefenseFinder outputs to summarize the presence of prokaryotic antiviral defense systems across multiple genome assemblies (Table 2). The results revealed a diverse array of defense mechanisms distributed unevenly among the samples. The number of detected defense systems per genome ranged from 3 to 24, with NC_021184.1 exhibiting the highest count (24 systems) and NC_014168.1 presenting the fewest (3 systems). In terms of diversity, NC_021184.1 displayed the greatest richness in both system types (13 unique types) and subtypes (n=18), followed by NC_019904.1, which contained 12 system types and 13 subtypes. Conversely, samples such as NC_014168.1 showed limited diversity, with only 2 unique types. Gene content associated with defense systems also varied substantially. The sample NC_021184.1 encoded the largest number of defense-related genes (n=75), while NC_014168.1 had the fewest (n=4). These results illustrate how PHI integrates and reports DefenseFinder outputs to enable comparison of antiviral defense repertoires across genomes. The observed variation highlights the level of detail that can be captured using the DefenseFinder module within the Phage Toolkit.

**Table 2.**
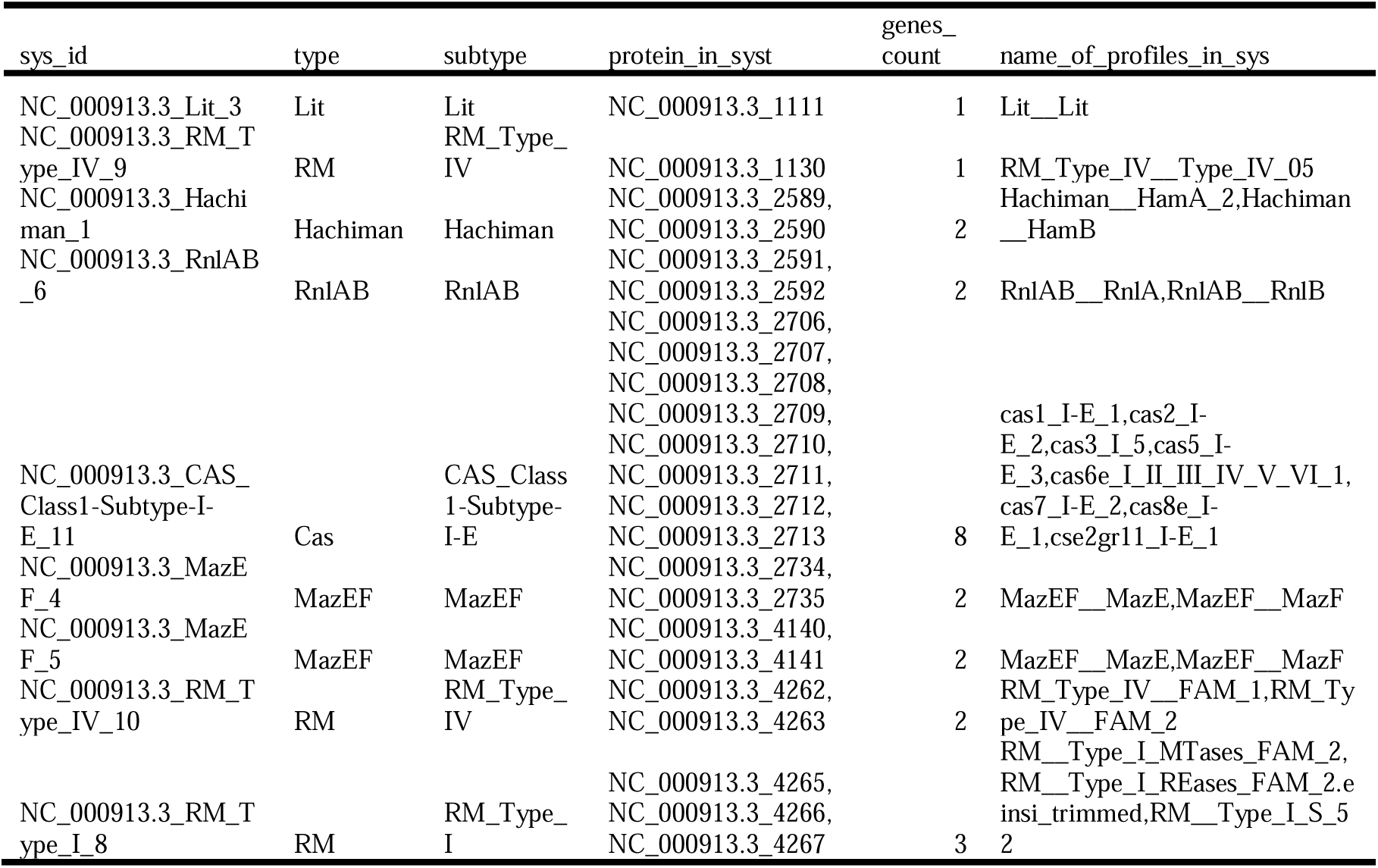
Filtered DefenseFinder (13) results presented in the HTML report. The filtered contents include the identification of defense systems, the type, subtype, and their locations in the contig list.

#### Identification and quality of prophages

In the demonstration dataset, the report showed that 14 out of the 22 analyzed bacterial genomes were hosting candidate prophages (Figure 2). A total of 41 prophages were found in the sample pool, with seven and nine being classified as high- and medium-quality, respectively. The sequences were also categorized using MIUViG standards. Prophages carried varying numbers of genes, typically ranging between 19 and 58 genes, with sequence lengths ranging from approximately 9.8kb to 28.6kb. A visual representation of the identified prophages, their length, and location within the host genome is provided in Figure 3.

**Figure 3.**
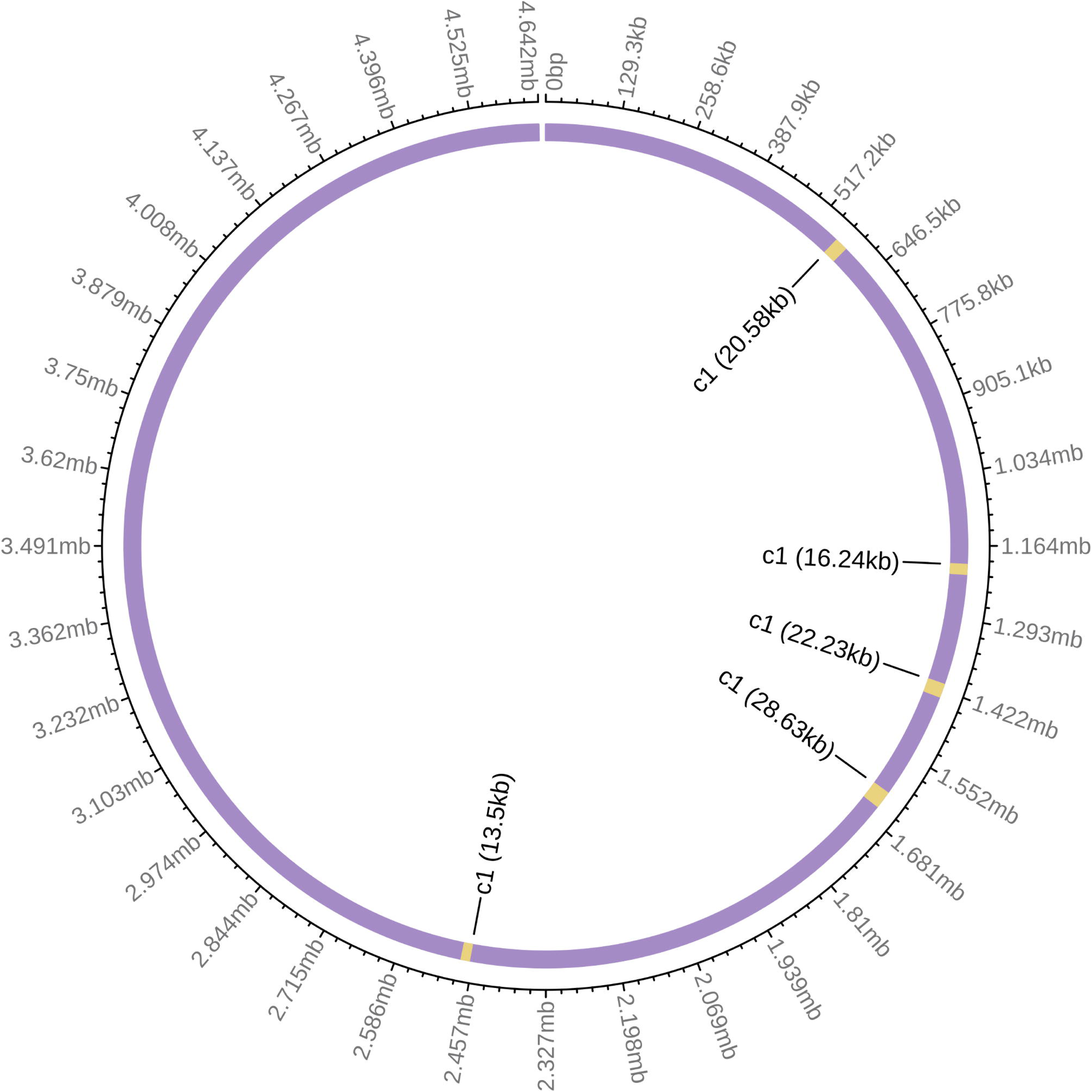
Example visualization of the Host Genome Ideogram. The graph shows the location and length of the predicted prophages.

#### Prophage Genome annotation

A visual representation of the prophage genome and features such as quality, number of genes (and viral genes) is shown for each predicted prophage (Figure 4). The integrated annotation outputs reported 4350 predicted genes (783 marked as viral□hallmark), with the most common annotations being capsid and structural connector proteins. No plasmid signatures, AMR, or conjugation genes were predicted.

**Figure 4.**
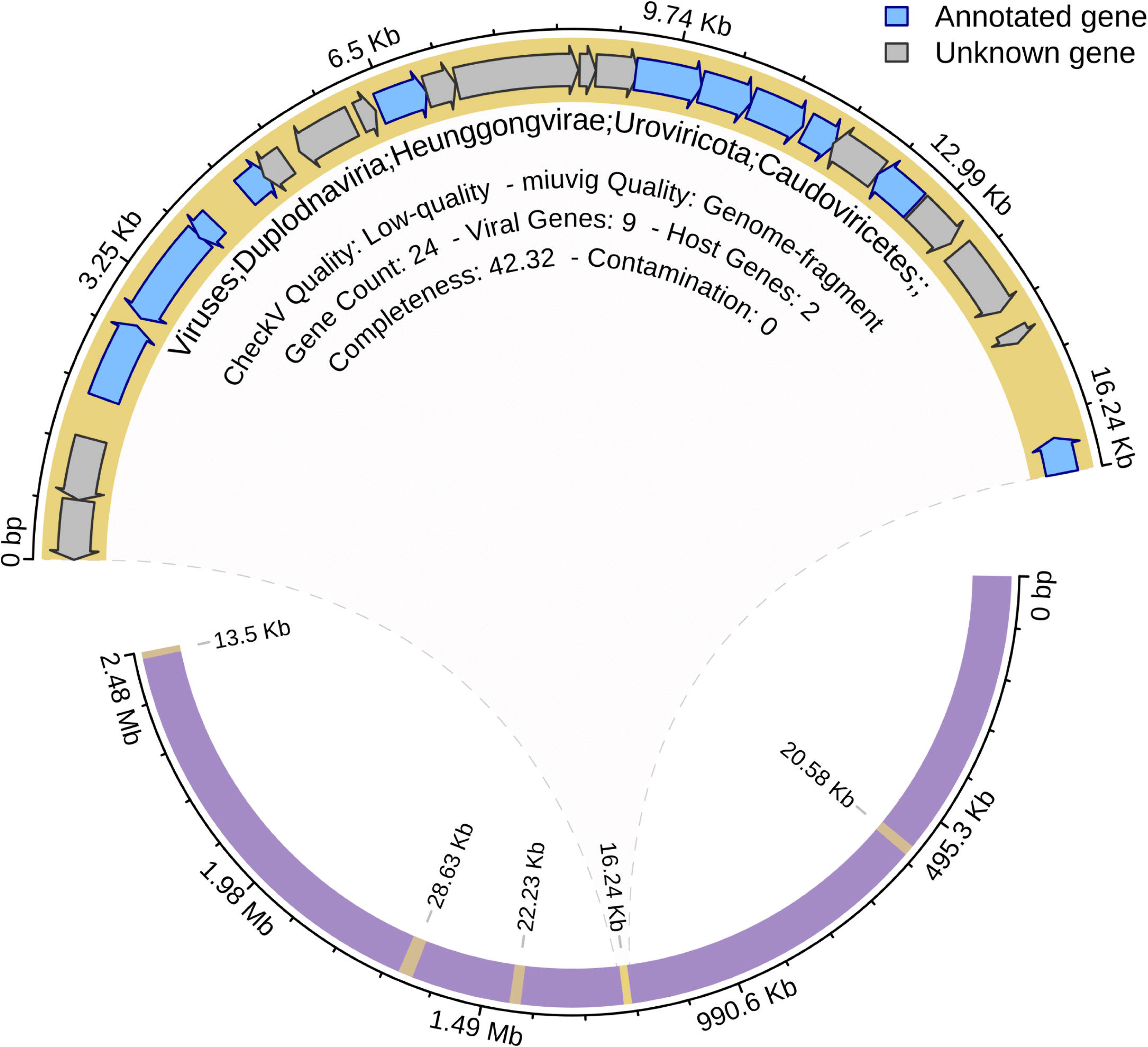
Example visualization of a prophage genome and its associated features. The PHI report generates a separate visualization tab for each predicted phage, showing sequence location, quality, taxonomy, and annotated genes.

## Conclusions

The PHI Toolkit, available through the Galaxy platform, addresses a critical bottleneck in phage research by enabling worldwide access to powerful, yet accessible, computational analyses for both bioinformatics experts and non-experts. Rather than introducing novel predictive methods and algorithms, PHI consolidates tools into a streamlined web interface with a modular, cloud-hosted architecture, eliminating the need for local hardware and advanced bioinformatics expertise. Researchers can therefore avoid manual data wrangling and devote their efforts to biological discovery rather than software installation, maintenance, or resource management. Because each module is version-tracked and can be updated, users can rely on established and actively maintained tools without manual upgrades, while still retaining the flexibility to swap in new tools as the field evolves. Furthermore, the integration of these results within a single interactive report facilitates biological interpretation by linking host identity, prophage content, genome context, and functional annotations. For instance, host genome ideograms and prophage feature maps allow users to examine the genomic location, size, and characteristics of predicted prophages within their bacterial hosts, while CRISPR spacer matching tables provide additional evidence for potential host-phage relationships. When combined with prophage annotations, virulence screening, AMG detection, and host prediction outputs, this integrated structure enables rapid identification of biologically relevant patterns that would be difficult to infer from individual result files alone. Future development of PHI will focus on expanding the workflow with additional prophage-detection, host-prediction, annotation, and visualization modules as new methods become available. Beyond general phage–host interaction studies, PHI may support applied microbiome research by enabling systematic screening of prophage content, virulence-associated genes, auxiliary metabolic genes, and host defense systems in biotechnologically relevant bacteria. In the future, this could include microbial strains considered for bioinoculants or low-risk pesticide applications [24, 25], provided that dedicated validation datasets are used and the outputs are interpreted as part of a broader strain-characterization and risk-assessment framework.

## Availability and requirements

**Project name:** PHI: Prophage-Host Interaction Toolkit

**Project home page:** https://github.com/Helmholtz-UFZ/galaxy-tools/tree/main/tools/phi-toolkit

**Galaxy workflow:** https://usegalaxy.eu/published/workflow?id=c62d65832377e376

**Operating system(s):** Platform independent through Galaxy

**Programming language:** Galaxy workflow format; R/R Markdown for report generation; integrated tools use their respective languages

**Other requirements:** None

**License:** Galaxy Web Portal Service Agreement (https://usegalaxy.org/static/terms.html)

**Any restrictions to use by non-academics:** None

## List of abbeviations

AMG: Auxiliary metabolic gene
ANI: Average nucleotide identity
GTDB-Tk: Genome Taxonomy Database Toolkit
HTML: HyperText Markup Language
MAG: Metagenome-assembled genome
MIMAG: Minimum Information about a Metagenome-Assembled Genome
MISAG: Minimum Information about a Single Amplified Genome
MIUViG: Minimum Information about an Uncultivated Virus Genome
NCBI: National Center for Biotechnology Information
PHI: Prophage-Host Interaction
SAG: Single amplified genome
VFDB: Virulence Factor Database

## Declarations

### Ethics approval and consent to participate

Not applicable.

### Consent for publication

Not applicable.

### Availability of data and materials

The PHI Toolkit consists of the Galaxy workflow itself and a utility Galaxy tool (the final step of the workflow) that integrates the results of all used tools. The utility tool is developed at https://github.com/Helmholtz-UFZ/galaxy-tools/tree/main/tools/phi-toolkit and has been deployed to the Galaxy tool shed (which allows installation on any Galaxy instance). The full set of results can be exported from the Galaxy server via https://usegalaxy.eu/workflows/invocations/e3ee09d9ef3af4a4). Tool, workflow and workflow invocation has been deposited at Zenodo (https://zenodo.org/records/20050288). The MBARC22 genome accessions used for the demonstration are listed in Additional File S2 and were obtained from NCBI, except for NC_010067.1, which was suppressed and therefore excluded.

### Authors’ contributions

JS and FBC conceived the workflow. JS, FBC, MB, NG, EN, and RBT contributed to workflow implementation, tool integration, testing, and report generation. JS drafted the original manuscript. LW and AC supervised the study and contributed to manuscript development. All authors reviewed and approved the final manuscript.

## Acknowledgments

Not applicable.

---

**Additional File 1: Figure S1. Detailed workflow schema of the PHI toolkit**, showing the Galaxy-native integration of prophage detection, viral quality assessment, host prediction, dereplication, genome quality control, taxonomy, defense-system screening, virulence-factor screening, antimicrobial-resistance screening, functional annotation, and final report generation.

**Additional File 2: Figure S2. Example PHI HTML report for the MBARC22 dataset.** Structured HTML report generated by the PHI workflow for the MBARC22 mock-community dataset. The report summarizes genome accession numbers and integrates host, prophage, and interaction-level outputs, including genome quality, taxonomy, prophage content, host–virus association networks, CRISPR spacer–protospacer matching results, genome-level feature maps, host-range assessment, and genomic-context visualizations.

**Additional File 3: Table S1. Complete computational-resource table for PHI runs.** Complete per-run resource-usage table for the MBARC22 analysis. It contains the individual runtime and memory-peak values underlying the average computational-resource summary reported in Table 1, including tool-specific runtime and memory requirements for CheckM2, GTDB-Tk, dRep, DefenseFinder, geNomad, ABRicate, CheckV, iPHoP, and VIBRANT.

**Additional File 4: Table S2. File-size grouped runtime and scalability analysis.** Scalability analysis of PHI tool runtimes grouped by input file-size intervals. For each tool, runtimes were summarized as mean values within file-size bins to evaluate whether runtime increased with input size.

