## Supplementary figures and images for "PHI: a Galaxy-based workflow for reproducible prophage-host interaction analysis and standardized viral-genomics reporting"

### Figure S1

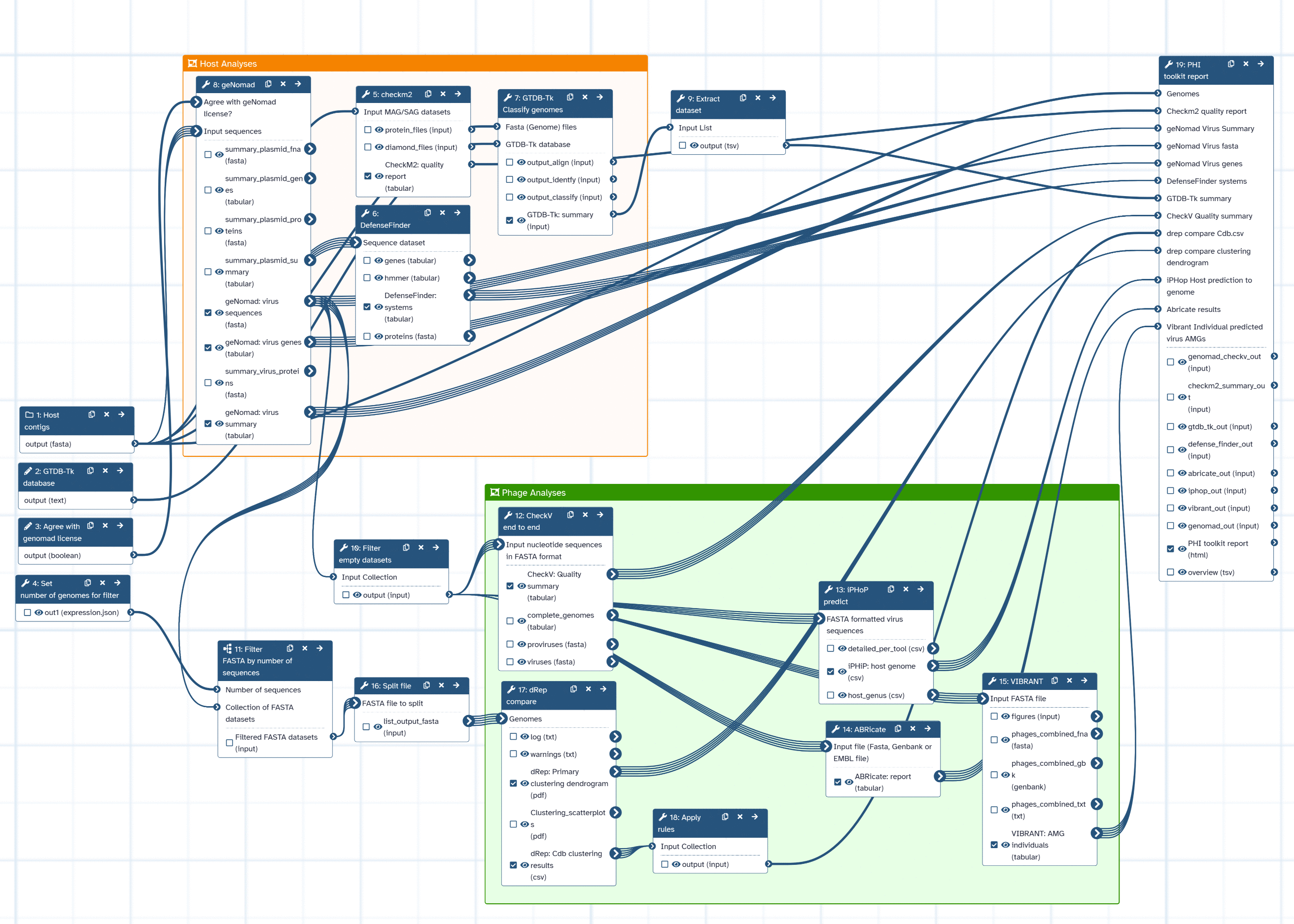
