## Supplementary material for "PHI: a Galaxy-based workflow for reproducible prophage-host interaction analysis and standardized viral-genomics reporting": Figure S2: Suppl. File S2.html

PHI Prophage-Host Interaction Toolkit report


### PHI Prophage-Host Interaction Toolkit report

##### Toolkit for the Detection, Comparison, and Annotation of Prophages in Bacterial Genomes.

###### May 05, 2026

- 1 Summary
- 2 Results
  - 2.1 NC\_000913
    - 2.1.1 Host Genome
    - 2.1.2 Prophages
    - 2.1.3 vOTUs
    - 2.1.4 Virulence Genes

---

### 1 Summary

This table provides sample-by-sample information on detected viruses
and key host genome statistics. It includes taxonomy, virus count,
genome quality classification, CheckM2 metrics (completeness and
contamination), and genome assembly statistics such as size and N50.

|  | | | | Host Genome Quality | | CheckM Metrics | | Statistics | |
| --- | --- | --- | --- | --- | --- | --- | --- | --- | --- |
| # | Sample | GTDB Taxonomy | Virus\_Count | Quality\_Score | Genome\_Quality | Completeness | Contamination | Genome size (bp) | N50 (contigs) |
| 1 | NC\_000913 | *Escherichia coli* | 5 | 99.3 | High-quality draft | 100.0% | 0.1% | 4.6M | 4.6M |
| 2 | NC\_002737 | *Streptococcus pyogenes* | 3 | 97.6 | High-quality draft | 100.0% | 0.5% | 1.9M | 1.9M |
| 3 | NC\_003450 | *Corynebacterium glutamicum* | 1 | 98.5 | High-quality draft | 100.0% | 0.3% | 3.3M | 3.3M |
| 4 | NC\_008261 | *Sarcina perfringens* | 2 | 99.3 | High-quality draft | 100.0% | 0.1% | 3.3M | 3.3M |
| 5 | NC\_009012 | *Hungateiclostridium thermocellum* | 5 | 94.5 | High-quality draft | 100.0% | 1.1% | 3.8M | 3.8M |
| 6 | NC\_012982 | *Hirschia baltica* | 0 | 99.8 | High-quality draft | 100.0% | 0.1% | 3.5M | 3.5M |
| 7 | NC\_014008 | *Coraliomargarita akajimensis* | 0 | 99.9 | High-quality draft | 100.0% | 0.0% | 3.8M | 3.8M |
| 8 | NC\_014168 | *Segniliparus rotundus* | 2 | 99.8 | High-quality draft | 100.0% | 0.0% | 3.2M | 3.2M |
| 9 | NC\_014211 | *Nocardiopsis dassonvillei* | 0 | 14.0 | Low-quality draft | 14.1% | 0.0% | 775.4K | 775.4K |
| 10 | NC\_014212 | *Allomeiothermus silvanus* | 3 | 99.2 | High-quality draft | 100.0% | 0.2% | 3.2M | 3.2M |
| 11 | NC\_014363 | *Olsenella uli* | 0 | 98.0 | High-quality draft | 99.7% | 0.3% | 2.1M | 2.1M |
| 12 | NC\_014364 | *Sediminispirochaeta smaragdinae* | 4 | 90.6 | High-quality draft | 100.0% | 1.9% | 4.7M | 4.7M |
| 13 | NC\_015761 | *Salmonella bongori* | 3 | 99.3 | High-quality draft | 100.0% | 0.1% | 4.5M | 4.5M |
| 14 | NC\_017033 | *Frateuria aurantia* | 3 | 99.8 | High-quality draft | 100.0% | 0.0% | 3.6M | 3.6M |
| 15 | NC\_017095 | *Fervidobacterium pennivorans* | 0 | 86.6 | High-quality draft | 100.0% | 2.7% | 2.2M | 2.2M |
| 16 | NC\_018014 | *Terriglobus roseus* | 0 | 53.1 | Medium-quality draft | 100.0% | 9.4% | 5.2M | 5.2M |
| 17 | NC\_018068 | *Desulfosporosinus acidiphilus* | 2 | 97.9 | High-quality draft | 100.0% | 0.4% | 4.9M | 4.9M |
| 18 | NC\_018515 | *Desulfosporosinus meridiei* | 2 | 90.8 | High-quality draft | 100.0% | 1.8% | 4.9M | 4.9M |
| 19 | NC\_019897 | *Thermobacillus xylanilyticus* | 0 | 98.2 | High-quality draft | 100.0% | 0.3% | 4.2M | 4.2M |
| 20 | NC\_019904 | *Echinicola vietnamensis* | 0 | 98.8 | High-quality draft | 100.0% | 0.2% | 5.6M | 5.6M |
| 21 | NC\_019936 | *Stutzerimonas stutzeri\_AE* | 3 | 99.3 | High-quality draft | 100.0% | 0.1% | 4.6M | 4.6M |
| 22 | NC\_021184 | *Sporotomaculum gibsoniae* | 3 | 84.8 | High-quality draft | 100.0% | 3.0% | 4.9M | 4.9M |

### 2 Results

**Select sample to show:**

## 2.1 NC\_000913

##### 2.1.1 Host Genome

**GTDB-Tk taxonomy**:

classification

d\_\_Bacteria;p\_\_Pseudomonadota;c\_\_Gammaproteobacteria;o\_\_Enterobacterales;f\_\_Enterobacteriaceae;g\_\_Escherichia;s\_\_Escherichia
coli

**CheckM2 Summary**:

| total\_contigs | contig\_n50 | completeness | contamination |
| --- | --- | --- | --- |
| 1 | 4641652 | 100 | 0.13 |

**Defense-Finder Systems**:

| sys\_id | type | subtype | sys\_beg | sys\_end | protein\_in\_syst | genes\_count | name\_of\_profiles\_in\_sys |
| --- | --- | --- | --- | --- | --- | --- | --- |
| NC\_000913.3\_Lit\_3 | Lit | Lit | NC\_000913.3\_1111 | NC\_000913.3\_1111 | NC\_000913.3\_1111 | 1 | Lit\_\_Lit |
| NC\_000913.3\_RM\_Type\_IV\_9 | RM | RM\_Type\_IV | NC\_000913.3\_1130 | NC\_000913.3\_1130 | NC\_000913.3\_1130 | 1 | RM\_Type\_IV\_\_Type\_IV\_05 |
| NC\_000913.3\_Hachiman\_1 | Hachiman | Hachiman | NC\_000913.3\_2589 | NC\_000913.3\_2590 | NC\_000913.3\_2589,NC\_000913.3\_2590 | 2 | Hachiman\_\_HamA\_2,Hachiman\_\_HamB |
| NC\_000913.3\_RnlAB\_6 | RnlAB | RnlAB | NC\_000913.3\_2591 | NC\_000913.3\_2592 | NC\_000913.3\_2591,NC\_000913.3\_2592 | 2 | RnlAB\_\_RnlA,RnlAB\_\_RnlB |
| NC\_000913.3\_CAS\_Class1-Subtype-I-E\_11 | Cas | CAS\_Class1-Subtype-I-E | NC\_000913.3\_2706 | NC\_000913.3\_2713 | NC\_000913.3\_2706,NC\_000913.3\_2707,NC\_000913.3\_2708,NC\_000913.3\_2709,NC\_000913.3\_2710,NC\_000913.3\_2711,NC\_000913.3\_2712,NC\_000913.3\_2713 | 8 | cas1\_I-E\_1,cas2\_I-E\_2,cas3\_I\_5,cas5\_I-E\_3,cas6e\_I\_II\_III\_IV\_V\_VI\_1,cas7\_I-E\_2,cas8e\_I-E\_1,cse2gr11\_I-E\_1 |
| NC\_000913.3\_MazEF\_4 | MazEF | MazEF | NC\_000913.3\_2734 | NC\_000913.3\_2735 | NC\_000913.3\_2734,NC\_000913.3\_2735 | 2 | MazEF\_\_MazE,MazEF\_\_MazF |
| NC\_000913.3\_MazEF\_5 | MazEF | MazEF | NC\_000913.3\_4140 | NC\_000913.3\_4141 | NC\_000913.3\_4140,NC\_000913.3\_4141 | 2 | MazEF\_\_MazE,MazEF\_\_MazF |
| NC\_000913.3\_RM\_Type\_IV\_10 | RM | RM\_Type\_IV | NC\_000913.3\_4262 | NC\_000913.3\_4263 | NC\_000913.3\_4262,NC\_000913.3\_4263 | 2 | RM\_Type\_IV\_\_FAM\_1,RM\_Type\_IV\_\_FAM\_2 |
| NC\_000913.3\_RM\_Type\_I\_8 | RM | RM\_Type\_I | NC\_000913.3\_4265 | NC\_000913.3\_4267 | NC\_000913.3\_4265,NC\_000913.3\_4266,NC\_000913.3\_4267 | 3 | RM\_\_Type\_I\_MTases\_FAM\_2,RM\_\_Type\_I\_REases\_FAM\_2.einsi\_trimmed,RM\_\_Type\_I\_S\_52 |

**Genomad and CheckV Summary**:

| seq\_name | length | gene\_count | viral\_genes | checkv\_quality | miuvig\_quality | taxonomy | topology | coordinates |
| --- | --- | --- | --- | --- | --- | --- | --- | --- |
| NC\_000913.3\|provirus\_1196867\_1213107 | 16241 | 24 | 9 | Low-quality | Genome-fragment | Viruses;Duplodnaviria;Heunggongvirae;Uroviricota;Caudoviricetes;; | Provirus | 1196867-1213107 |
| NC\_000913.3\|provirus\_1412000\_1434224 | 22225 | 27 | 13 | Low-quality | Genome-fragment | Viruses;Duplodnaviria;Heunggongvirae;Uroviricota;Caudoviricetes;; | Provirus | 1412000-1434224 |
| NC\_000913.3\|provirus\_2463012\_2476510 | 13499 | 19 | 12 | Low-quality | Genome-fragment | Viruses;Duplodnaviria;Heunggongvirae;Uroviricota;Caudoviricetes;; | Provirus | 2463012-2476510 |
| NC\_000913.3\|provirus\_563848\_584430 | 20583 | 32 | 13 | Low-quality | Genome-fragment | Viruses;Duplodnaviria;Heunggongvirae;Uroviricota;Caudoviricetes;; | Provirus | 563848-584430 |
| NC\_000913.3\|provirus\_1627517\_1656149 | 28633 | 44 | 10 | Low-quality | Genome-fragment | Viruses;Duplodnaviria;Heunggongvirae;Uroviricota;Caudoviricetes;; | Provirus | 1627517-1656149 |

**Host Genome Ideogram with Phages**:

In
this circular plot, **“c”** indicates the contig, and the
number that follows (e.g., **c1**) represents the contig
number. If multiple contigs are present in the genome, each will be
shown with a distinct label (e.g., **c1**,
**c2**, etc.).

##### 2.1.2 Prophages

**Select prophage to show:**

###### 2.1.2.1 Phage ID: NC\_000913.3|provirus\_1196867\_1213107

**Phage-Host Genome Ideogram:**

**Genes Annotation (geNomad):**

| gene | length | marker | annotation\_accessions | annotation\_description |
| --- | --- | --- | --- | --- |
| NC\_000913.3\|provirus\_1196867\_1213107\_1105 | 666 | NA | NA | NA |
| NC\_000913.3\|provirus\_1196867\_1213107\_1106 | 696 | NA | NA | NA |
| NC\_000913.3\|provirus\_1196867\_1213107\_1107 | 894 | GENOMAD.088167.PP | PF10463;COG2856 | Peptidase U49 |
| NC\_000913.3\|provirus\_1196867\_1213107\_1108 | 1128 | GENOMAD.195357.VP | PF09003;K21039 | Bacteriophage lambda integrase, Arm DNA-binding domain |
| NC\_000913.3\|provirus\_1196867\_1213107\_1109 | 246 | GENOMAD.095038.VV | PF07825 | Excisionase-like protein |
| NC\_000913.3\|provirus\_1196867\_1213107\_1110 | 342 | GENOMAD.226053.VP | PF11080;K18840 | Endoribonuclease GhoS |
| NC\_000913.3\|provirus\_1196867\_1213107\_1111 | 309 | GENOMAD.226995.VP | NA | NA |
| NC\_000913.3\|provirus\_1196867\_1213107\_1112 | 675 | NA | NA | NA |
| NC\_000913.3\|provirus\_1196867\_1213107\_1113 | 201 | GENOMAD.218084.VP | PF15943;COG4197 | NA |
| NC\_000913.3\|provirus\_1196867\_1213107\_1114 | 558 | GENOMAD.058510.VV | PF06892 | Phage regulatory protein CII (CP76) |
| NC\_000913.3\|provirus\_1196867\_1213107\_1115 | 339 | GENOMAD.102216.VV | NA | NA |
| NC\_000913.3\|provirus\_1196867\_1213107\_1116 | 1368 | GENOMAD.159591.VP | NA | NA |
| NC\_000913.3\|provirus\_1196867\_1213107\_1117 | 183 | GENOMAD.088953.VV | PF10003 | NA |
| NC\_000913.3\|provirus\_1196867\_1213107\_1118 | 474 | GENOMAD.044416.VV | TIGR01537 | NA |
| NC\_000913.3\|provirus\_1196867\_1213107\_1119 | 750 | GENOMAD.110264.VV | PF04865;COG3948 | Phage-related baseplate assembly protein |
| NC\_000913.3\|provirus\_1196867\_1213107\_1120 | 585 | GENOMAD.113504.VP | PF10076;COG3778 | Uncharacterized protein YmfQ in lambdoid prophage, DUF2313 family |
| NC\_000913.3\|provirus\_1196867\_1213107\_1121 | 630 | GENOMAD.044962.VV | PF21882 | Putative tail fiber protein gp53-like, C-terminal |
| NC\_000913.3\|provirus\_1196867\_1213107\_1122 | 375 | GENOMAD.045203.VV | PF02413 | Caudovirales tail fibre assembly protein, lambda gpK |
| NC\_000913.3\|provirus\_1196867\_1213107\_1123 | 603 | GENOMAD.140378.VP | PF02413 | NA |
| NC\_000913.3\|provirus\_1196867\_1213107\_1124 | 537 | GENOMAD.191048.VP | COG4675 | Microcystin-dependent protein (function unknown) |
| NC\_000913.3\|provirus\_1196867\_1213107\_1125 | 555 | NA | NA | NA |
| NC\_000913.3\|provirus\_1196867\_1213107\_1126 | 834 | GENOMAD.139104.VV | PF13395 | NA |
| NC\_000913.3\|provirus\_1196867\_1213107\_1127 | 165 | NA | NA | NA |
| NC\_000913.3\|provirus\_1196867\_1213107\_1128 | 405 | GENOMAD.227190.VP | PF07166;COG5562 | Prophage-encoded protein YbcV, DUF1398 family |

###### 2.1.2.2 Phage ID: NC\_000913.3|provirus\_1412000\_1434224

**Phage-Host Genome Ideogram:**

**Genes Annotation (geNomad):**

| gene | length | marker | annotation\_accessions | annotation\_description |
| --- | --- | --- | --- | --- |
| NC\_000913.3\|provirus\_1412000\_1434224\_1320 | 1236 | GENOMAD.159447.VP | PF12167;K14059;TIGR01634;COG4385 | phage tail protein, P2 protein I family |
| NC\_000913.3\|provirus\_1412000\_1434224\_1321 | 216 | GENOMAD.114098.VV | PF06806 | Putative excisionase (DUF1233) |
| NC\_000913.3\|provirus\_1412000\_1434224\_1322 | 210 | GENOMAD.223267.VP | PF06688;K19780 | NA |
| NC\_000913.3\|provirus\_1412000\_1434224\_1323 | 810 | GENOMAD.127355.VP | PF03837;TIGR00616;COG3723 | recombinase, phage RecT family |
| NC\_000913.3\|provirus\_1412000\_1434224\_1324 | 2601 | GENOMAD.084315.VV | PF06630;K10906 | Enterobacterial exodeoxyribonuclease VIII |
| NC\_000913.3\|provirus\_1412000\_1434224\_1325 | 276 | NA | NA | NA |
| NC\_000913.3\|provirus\_1412000\_1434224\_1326 | 171 | GENOMAD.110847.VV | PF04181 | NA |
| NC\_000913.3\|provirus\_1412000\_1434224\_1327 | 222 | GENOMAD.074955.VV | NA | NA |
| NC\_000913.3\|provirus\_1412000\_1434224\_1328 | 612 | NA | NA | NA |
| NC\_000913.3\|provirus\_1412000\_1434224\_1329 | 156 | GENOMAD.222124.VP | PF07151 | NA |
| NC\_000913.3\|provirus\_1412000\_1434224\_1330 | 135 | NA | NA | NA |
| NC\_000913.3\|provirus\_1412000\_1434224\_1331 | 477 | GENOMAD.115833.VV | PF00376;COG5606;TIGR00673;K10123 | MerR family regulatory protein |
| NC\_000913.3\|provirus\_1412000\_1434224\_1332 | 297 | GENOMAD.054651.VV | PF15943;COG4197 | DNA-binding transcriptional regulator YdaS, prophage-encoded, Cro superfamily |
| NC\_000913.3\|provirus\_1412000\_1434224\_1333 | 423 | GENOMAD.072359.VV | PF06254 | Putative bacterial toxin ydaT |
| NC\_000913.3\|provirus\_1412000\_1434224\_1334 | 858 | GENOMAD.195765.VP | PF07120;COG3756 | Uncharacterized conserved protein YdaU, DUF1376 family |
| NC\_000913.3\|provirus\_1412000\_1434224\_1335 | 747 | GENOMAD.047125.VV | PF16463;COG5437;TIGR02126 | Phage tail tube protein family |
| NC\_000913.3\|provirus\_1412000\_1434224\_1336 | 561 | GENOMAD.144793.VV | PF07789 | NA |
| NC\_000913.3\|provirus\_1412000\_1434224\_1337 | 300 | GENOMAD.219285.VP | PF03245 | Bacteriophage Rz lysis protein |
| NC\_000913.3\|provirus\_1412000\_1434224\_1338 | 1425 | NA | NA | NA |
| NC\_000913.3\|provirus\_1412000\_1434224\_1339 | 264 | GENOMAD.159709.VV | NA | NA |
| NC\_000913.3\|provirus\_1412000\_1434224\_1340 | 360 | NA | NA | NA |
| NC\_000913.3\|provirus\_1412000\_1434224\_1341 | 1029 | GENOMAD.120741.VP | COG5281 | Phage-related minor tail protein |
| NC\_000913.3\|provirus\_1412000\_1434224\_1342 | 132 | NA | NA | NA |
| NC\_000913.3\|provirus\_1412000\_1434224\_1343 | 981 | NA | NA | NA |
| NC\_000913.3\|provirus\_1412000\_1434224\_1344 | 3363 | GENOMAD.020617.VV | NA | NA |
| NC\_000913.3\|provirus\_1412000\_1434224\_1345 | 576 | GENOMAD.167268.VP | PF02413 | Caudovirales tail fibre assembly protein, lambda gpK |
| NC\_000913.3\|provirus\_1412000\_1434224\_1346 | 591 | NA | NA | NA |
| NC\_000913.3\|provirus\_1412000\_1434224\_1347 | 234 | GENOMAD.221767.PV | PF10965 | Biofilm development protein YmgB/AriR |

###### 2.1.2.3 Phage ID: NC\_000913.3|provirus\_2463012\_2476510

**Phage-Host Genome Ideogram:**

**Genes Annotation (geNomad):**

| gene | length | marker | annotation\_accessions | annotation\_description |
| --- | --- | --- | --- | --- |
| NC\_000913.3\|provirus\_2463012\_2476510\_2314 | 1059 | NA | NA | NA |
| NC\_000913.3\|provirus\_2463012\_2476510\_2315 | 756 | NA | NA | NA |
| NC\_000913.3\|provirus\_2463012\_2476510\_2316 | 933 | NA | NA | NA |
| NC\_000913.3\|provirus\_2463012\_2476510\_2317 | 1158 | NA | NA | NA |
| NC\_000913.3\|provirus\_2463012\_2476510\_2318 | 921 | NA | NA | NA |
| NC\_000913.3\|provirus\_2463012\_2476510\_2319 | 1332 | NA | NA | NA |
| NC\_000913.3\|provirus\_2463012\_2476510\_2320 | 303 | GENOMAD.167268.VP | PF02413 | Caudovirales tail fibre assembly protein, lambda gpK |
| NC\_000913.3\|provirus\_2463012\_2476510\_2321 | 441 | GENOMAD.045203.VV | PF02413 | Caudovirales tail fibre assembly protein, lambda gpK |
| NC\_000913.3\|provirus\_2463012\_2476510\_2322 | 603 | GENOMAD.044962.VV | PF21882 | Putative tail fiber protein gp53-like, C-terminal |
| NC\_000913.3\|provirus\_2463012\_2476510\_2323 | 276 | GENOMAD.178309.VP | PF05869;TIGR01712 | phage N-6-adenine-methyltransferase |
| NC\_000913.3\|provirus\_2463012\_2476510\_2324 | 495 | GENOMAD.225234.VP | PF06069 | PerC transcriptional activator |
| NC\_000913.3\|provirus\_2463012\_2476510\_2325 | 369 | GENOMAD.086968.VV | NA | NA |
| NC\_000913.3\|provirus\_2463012\_2476510\_2326 | 363 | GENOMAD.056235.VV | NA | NA |
| NC\_000913.3\|provirus\_2463012\_2476510\_2327 | 825 | GENOMAD.052651.VV | PF10065;COG5532 | Uncharacterized conserved protein YfdQ, DUF2303 family |
| NC\_000913.3\|provirus\_2463012\_2476510\_2328 | 537 | GENOMAD.172054.VC | NA | NA |
| NC\_000913.3\|provirus\_2463012\_2476510\_2329 | 363 | GENOMAD.067358.VV | NA | NA |
| NC\_000913.3\|provirus\_2463012\_2476510\_2330 | 306 | GENOMAD.222350.VP | NA | NA |
| NC\_000913.3\|provirus\_2463012\_2476510\_2331 | 201 | GENOMAD.195460.VP | COG2452 | NA |

###### 2.1.2.4 Phage ID: NC\_000913.3|provirus\_563848\_584430

**Phage-Host Genome Ideogram:**

**Genes Annotation (geNomad):**

| gene | length | marker | annotation\_accessions | annotation\_description |
| --- | --- | --- | --- | --- |
| NC\_000913.3\|provirus\_563848\_584430\_520 | 633 | NA | NA | NA |
| NC\_000913.3\|provirus\_563848\_584430\_521 | 1164 | GENOMAD.133037.VP | NA | NA |
| NC\_000913.3\|provirus\_563848\_584430\_522 | 264 | GENOMAD.095346.VP | PF09588;K01143;COG5377;TIGR03033 | Phage-related protein, predicted endonuclease |
| NC\_000913.3\|provirus\_563848\_584430\_523 | 96 | GENOMAD.086130.VV | PF08222;TIGR00373;COG4519 | CodY helix-turn-helix domain |
| NC\_000913.3\|provirus\_563848\_584430\_524 | 300 | NA | NA | NA |
| NC\_000913.3\|provirus\_563848\_584430\_525 | 867 | NA | NA | NA |
| NC\_000913.3\|provirus\_563848\_584430\_526 | 333 | NA | NA | NA |
| NC\_000913.3\|provirus\_563848\_584430\_527 | 1527 | NA | NA | NA |
| NC\_000913.3\|provirus\_563848\_584430\_528 | 552 | GENOMAD.189584.CP | PF01161;COG1881;TIGR00481 | Phosphatidylethanolamine-binding protein |
| NC\_000913.3\|provirus\_563848\_584430\_529 | 798 | GENOMAD.166602.PC | PF06719;PF12833;TIGR02297;K18991;COG2207 | 4-hydroxyphenylacetate catabolism regulatory protein HpaA |
| NC\_000913.3\|provirus\_563848\_584430\_530 | 456 | GENOMAD.022699.VV | PF05772 | NinB protein |
| NC\_000913.3\|provirus\_563848\_584430\_531 | 171 | GENOMAD.177576.VP | PF05322 | NINE Protein |
| NC\_000913.3\|provirus\_563848\_584430\_532 | 291 | GENOMAD.038211.VV | PF07102 | Putative nuclease YbcO |
| NC\_000913.3\|provirus\_563848\_584430\_533 | 363 | GENOMAD.154527.VP | PF05866;K01160;COG4570 | Holliday junction resolvase RusA (prophage-encoded endonuclease) |
| NC\_000913.3\|provirus\_563848\_584430\_534 | 138 | GENOMAD.073918.VV | NA | NA |
| NC\_000913.3\|provirus\_563848\_584430\_535 | 384 | GENOMAD.054312.VV | PF06576 | Phage antitermination protein Q |
| NC\_000913.3\|provirus\_563848\_584430\_536 | 219 | NA | NA | NA |
| NC\_000913.3\|provirus\_563848\_584430\_537 | 981 | NA | NA | NA |
| NC\_000913.3\|provirus\_563848\_584430\_538 | 1068 | NA | NA | NA |
| NC\_000913.3\|provirus\_563848\_584430\_539 | 216 | GENOMAD.208385.VP | PF04971 | Bacteriophage P21 holin S |
| NC\_000913.3\|provirus\_563848\_584430\_540 | 498 | GENOMAD.171676.VP | PF00959;COG3772 | Phage-related lysozyme (muramidase), GH24 family |
| NC\_000913.3\|provirus\_563848\_584430\_541 | 462 | GENOMAD.219285.VP | PF03245 | Bacteriophage Rz lysis protein |
| NC\_000913.3\|provirus\_563848\_584430\_542 | 294 | NA | NA | NA |
| NC\_000913.3\|provirus\_563848\_584430\_543 | 411 | GENOMAD.227189.VP | NA | NA |
| NC\_000913.3\|provirus\_563848\_584430\_544 | 207 | GENOMAD.227340.VP | NA | NA |
| NC\_000913.3\|provirus\_563848\_584430\_545 | 546 | GENOMAD.161251.VP | PF07471;K22014;COG4220 | Phage DNA packaging protein, Nu1 subunit of terminase |
| NC\_000913.3\|provirus\_563848\_584430\_546 | 744 | GENOMAD.167268.VP | PF02413 | Caudovirales tail fibre assembly protein, lambda gpK |
| NC\_000913.3\|provirus\_563848\_584430\_547 | 432 | NA | NA | NA |
| NC\_000913.3\|provirus\_563848\_584430\_548 | 186 | GENOMAD.178075.VP | PF02413 | Caudovirales tail fibre assembly protein, lambda gpK |
| NC\_000913.3\|provirus\_563848\_584430\_549 | 750 | NA | NA | NA |

###### 2.1.2.5 Phage ID: NC\_000913.3|provirus\_1627517\_1656149

**Phage-Host Genome Ideogram:**

**Genes Annotation (geNomad):**

| gene | length | marker | annotation\_accessions | annotation\_description |
| --- | --- | --- | --- | --- |
| NC\_000913.3\|provirus\_1627517\_1656149\_1513 | 747 | NA | NA | NA |
| NC\_000913.3\|provirus\_1627517\_1656149\_1514 | 687 | NA | NA | NA |
| NC\_000913.3\|provirus\_1627517\_1656149\_1515 | 204 | NA | NA | NA |
| NC\_000913.3\|provirus\_1627517\_1656149\_1516 | 1461 | NA | NA | NA |
| NC\_000913.3\|provirus\_1627517\_1656149\_1517 | 1284 | NA | NA | NA |
| NC\_000913.3\|provirus\_1627517\_1656149\_1518 | 234 | GENOMAD.221767.PV | PF10965 | Biofilm development protein YmgB/AriR |
| NC\_000913.3\|provirus\_1627517\_1656149\_1519 | 591 | NA | NA | NA |
| NC\_000913.3\|provirus\_1627517\_1656149\_1520 | 576 | GENOMAD.167268.VP | PF02413 | Caudovirales tail fibre assembly protein, lambda gpK |
| NC\_000913.3\|provirus\_1627517\_1656149\_1521 | 963 | GENOMAD.211480.VP | COG5301 | Phage-related tail fibre protein |
| NC\_000913.3\|provirus\_1627517\_1656149\_1522 | 570 | GENOMAD.161251.VP | PF07471;K22014;COG4220 | Phage DNA packaging protein, Nu1 subunit of terminase |
| NC\_000913.3\|provirus\_1627517\_1656149\_1523 | 411 | GENOMAD.227189.VP | NA | NA |
| NC\_000913.3\|provirus\_1627517\_1656149\_1524 | 174 | NA | NA | NA |
| NC\_000913.3\|provirus\_1627517\_1656149\_1525 | 213 | NA | NA | NA |
| NC\_000913.3\|provirus\_1627517\_1656149\_1526 | 498 | GENOMAD.062288.VV | PF10721 | NA |
| NC\_000913.3\|provirus\_1627517\_1656149\_1527 | 534 | GENOMAD.171676.VP | PF00959;COG3772 | Phage-related lysozyme (muramidase), GH24 family |
| NC\_000913.3\|provirus\_1627517\_1656149\_1528 | 312 | GENOMAD.225519.VP | PF07041 | NA |
| NC\_000913.3\|provirus\_1627517\_1656149\_1529 | 207 | GENOMAD.208385.VP | PF04971 | Bacteriophage P21 holin S |
| NC\_000913.3\|provirus\_1627517\_1656149\_1530 | 120 | NA | NA | NA |
| NC\_000913.3\|provirus\_1627517\_1656149\_1531 | 216 | NA | NA | NA |
| NC\_000913.3\|provirus\_1627517\_1656149\_1532 | 213 | NA | NA | NA |
| NC\_000913.3\|provirus\_1627517\_1656149\_1533 | 753 | GENOMAD.136613.VP | PF03589;TIGR02642 | Antitermination protein |
| NC\_000913.3\|provirus\_1627517\_1656149\_1534 | 1050 | GENOMAD.196019.VP | PF06147 | dATP/dGTP pyrophosphohydrolase |
| NC\_000913.3\|provirus\_1627517\_1656149\_1535 | 252 | NA | NA | NA |
| NC\_000913.3\|provirus\_1627517\_1656149\_1536 | 156 | GENOMAD.136858.PV | PF01848;K18919 | Hok/gef family |
| NC\_000913.3\|provirus\_1627517\_1656149\_1537 | 288 | NA | NA | NA |
| NC\_000913.3\|provirus\_1627517\_1656149\_1538 | 240 | GENOMAD.151401.PV | PF04221;TIGR02384;COG3077;K07473 | addiction module antitoxin, RelB/DinJ family |
| NC\_000913.3\|provirus\_1627517\_1656149\_1539 | 333 | GENOMAD.078672.PP | PF14282 | FlxA-like protein |
| NC\_000913.3\|provirus\_1627517\_1656149\_1540 | 228 | NA | NA | NA |
| NC\_000913.3\|provirus\_1627517\_1656149\_1541 | 291 | GENOMAD.150946.VV | PF06254 | Putative bacterial toxin ydaT |
| NC\_000913.3\|provirus\_1627517\_1656149\_1542 | 231 | GENOMAD.108381.VV | K22302 | NA |
| NC\_000913.3\|provirus\_1627517\_1656149\_1543 | 408 | NA | NA | NA |
| NC\_000913.3\|provirus\_1627517\_1656149\_1544 | 156 | GENOMAD.222124.VP | PF07151 | NA |
| NC\_000913.3\|provirus\_1627517\_1656149\_1545 | 129 | GENOMAD.136713.VV | NA | NA |
| NC\_000913.3\|provirus\_1627517\_1656149\_1546 | 219 | GENOMAD.180239.VV | NA | NA |
| NC\_000913.3\|provirus\_1627517\_1656149\_1547 | 165 | NA | NA | NA |
| NC\_000913.3\|provirus\_1627517\_1656149\_1548 | 189 | GENOMAD.189052.VV | PF05358;K22304 | DicB protein |
| NC\_000913.3\|provirus\_1627517\_1656149\_1549 | 192 | GENOMAD.094923.VV | PF07358 | NA |
| NC\_000913.3\|provirus\_1627517\_1656149\_1550 | 921 | GENOMAD.075680.VV | K10906 | NA |
| NC\_000913.3\|provirus\_1627517\_1656149\_1551 | 543 | NA | NA | NA |
| NC\_000913.3\|provirus\_1627517\_1656149\_1552 | 1197 | GENOMAD.099267.VV | PF00589;TIGR02224;COG4973;K03733 | tyrosine recombinase XerC |
| NC\_000913.3\|provirus\_1627517\_1656149\_1553 | 111 | NA | NA | NA |
| NC\_000913.3\|provirus\_1627517\_1656149\_1554 | 1020 | NA | NA | NA |
| NC\_000913.3\|provirus\_1627517\_1656149\_1555 | 1215 | NA | NA | NA |
| NC\_000913.3\|provirus\_1627517\_1656149\_1556 | 327 | NA | NA | NA |
| NC\_000913.3\|provirus\_1627517\_1656149\_1557 | 342 | NA | NA | NA |

##### 2.1.3 vOTUs

When more than 1 phage is detected in the host genome, we perform a
clustering step using the tool dRep.

A threshold of 0.95 was applied to the ANI similarity index to define
clusters of virus operational taxonomic units (vOTUs).

**Final cluster designations**

| genomad\_summary$seq\_name | genome | secondary\_cluster | threshold | cluster\_method | comparison\_algorithm | primary\_cluster |
| --- | --- | --- | --- | --- | --- | --- |
| NC\_000913.3\|provirus\_1196867\_1213107 | sequence\_000002.fasta.fasta | 1\_0 | 0.05 | average | ANImf | 1 |
| NC\_000913.3\|provirus\_1412000\_1434224 | sequence\_000003.fasta.fasta | 2\_0 | 0.05 | average | ANImf | 2 |
| NC\_000913.3\|provirus\_2463012\_2476510 | sequence\_000000.fasta.fasta | 3\_0 | 0.05 | average | ANImf | 3 |
| NC\_000913.3\|provirus\_563848\_584430 | sequence\_000001.fasta.fasta | 4\_0 | 0.05 | average | ANImf | 4 |
| NC\_000913.3\|provirus\_1627517\_1656149 | sequence\_000004.fasta.fasta | 5\_0 | 0.05 | average | ANImf | 5 |

**Primary clustering plot**

##### 2.1.4 Virulence Genes

Screening of virulence genes present in the prophage contigs.

| sequence | start | end | strand | gene | coverage | coverage\_map | gaps | percent\_coverage | percent\_identity | database | accession | product | resistance |
| --- | --- | --- | --- | --- | --- | --- | --- | --- | --- | --- | --- | --- | --- |
| NC\_000913.3\|provirus\_2463012\_2476510 | 4844 | 5206 |  | gtrA | 1-363/363 | =============== | 0/0 | 100.00 | 88.70 | vfdb | NP\_706257 | (gtrA) bactoprenol-linked glucose translocase/flippase [LPS (VF0124)] [Shigella flexneri 2a str. 301] | NA |
| NC\_000913.3\|provirus\_2463012\_2476510 | 5203 | 6107 |  | gtrB | 1-905/930 | =============== | 0/0 | 97.31 | 80.66 | vfdb | NP\_706258 | (gtrB) bactoprenol glucosyl transferase [LPS (VF0124)] [Shigella flexneri 2a str. 301] | NA |


##### 2.1.5 Prophage-Host Prediction

Prediction of potential hosts for the prophage contigs.

| virus | host\_genome | host\_taxonomy | main\_method | confidence\_score | additional\_methods |
| --- | --- | --- | --- | --- | --- |
| NC\_000913.3\|provirus\_1196867\_1213107 | GB\_GCA\_021307345.1 | d\_\_Bacteria;p\_\_Pseudomonadota;c\_\_Gammaproteobacteria;o\_\_Enterobacterales;f\_\_Enterobacteriaceae;g\_\_Escherichia;s\_\_Escherichia ruysiae | blast | 96.9 | iPHoP-RF;92.80 |
| NC\_000913.3\|provirus\_1196867\_1213107 | RS\_GCF\_000026225.1 | d\_\_Bacteria;p\_\_Pseudomonadota;c\_\_Gammaproteobacteria;o\_\_Enterobacterales;f\_\_Enterobacteriaceae;g\_\_Escherichia;s\_\_Escherichia fergusonii | blast | 96.9 | iPHoP-RF;92.40 |
| NC\_000913.3\|provirus\_1196867\_1213107 | RS\_GCF\_000759775.1 | d\_\_Bacteria;p\_\_Pseudomonadota;c\_\_Gammaproteobacteria;o\_\_Enterobacterales;f\_\_Enterobacteriaceae;g\_\_Escherichia;s\_\_Escherichia albertii | blast | 96.9 | iPHoP-RF;93.40 |
| NC\_000913.3\|provirus\_1196867\_1213107 | RS\_GCF\_002900365.1 | d\_\_Bacteria;p\_\_Pseudomonadota;c\_\_Gammaproteobacteria;o\_\_Enterobacterales;f\_\_Enterobacteriaceae;g\_\_Escherichia;s\_\_Escherichia marmotae | blast | 96.9 | iPHoP-RF;96.40 |
| NC\_000913.3\|provirus\_1196867\_1213107 | RS\_GCF\_003697165.2 | d\_\_Bacteria;p\_\_Pseudomonadota;c\_\_Gammaproteobacteria;o\_\_Enterobacterales;f\_\_Enterobacteriaceae;g\_\_Escherichia;s\_\_Escherichia coli | blast | 96.9 | iPHoP-RF;93.70 |
| NC\_000913.3\|provirus\_1196867\_1213107 | RS\_GCF\_004211955.1 | d\_\_Bacteria;p\_\_Pseudomonadota;c\_\_Gammaproteobacteria;o\_\_Enterobacterales;f\_\_Enterobacteriaceae;g\_\_Escherichia;s\_\_Escherichia sp004211955 | iPHoP-RF | 96.4 | None |
| NC\_000913.3\|provirus\_1196867\_1213107 | RS\_GCF\_005843885.1 | d\_\_Bacteria;p\_\_Pseudomonadota;c\_\_Gammaproteobacteria;o\_\_Enterobacterales;f\_\_Enterobacteriaceae;g\_\_Escherichia;s\_\_Escherichia sp005843885 | iPHoP-RF | 96.4 | None |
| NC\_000913.3\|provirus\_1196867\_1213107 | RS\_GCF\_011881725.1 | d\_\_Bacteria;p\_\_Pseudomonadota;c\_\_Gammaproteobacteria;o\_\_Enterobacterales;f\_\_Enterobacteriaceae;g\_\_Escherichia;s\_\_Escherichia coli\_E | iPHoP-RF | 96.4 | None |
| NC\_000913.3\|provirus\_1196867\_1213107 | RS\_GCF\_002965065.1 | d\_\_Bacteria;p\_\_Pseudomonadota;c\_\_Gammaproteobacteria;o\_\_Enterobacterales;f\_\_Enterobacteriaceae;g\_\_Escherichia;s\_\_Escherichia sp002965065 | iPHoP-RF | 96.1 | None |
| NC\_000913.3\|provirus\_1196867\_1213107 | RS\_GCF\_014836715.1 | d\_\_Bacteria;p\_\_Pseudomonadota;c\_\_Gammaproteobacteria;o\_\_Enterobacterales;f\_\_Enterobacteriaceae;g\_\_Escherichia;s\_\_Escherichia whittamii | iPHoP-RF | 95.1 | None |
| NC\_000913.3\|provirus\_1412000\_1434224 | GB\_GCA\_021307345.1 | d\_\_Bacteria;p\_\_Pseudomonadota;c\_\_Gammaproteobacteria;o\_\_Enterobacterales;f\_\_Enterobacteriaceae;g\_\_Escherichia;s\_\_Escherichia ruysiae | blast | 96.9 | iPHoP-RF;93.70 |
| NC\_000913.3\|provirus\_1412000\_1434224 | RS\_GCF\_000759775.1 | d\_\_Bacteria;p\_\_Pseudomonadota;c\_\_Gammaproteobacteria;o\_\_Enterobacterales;f\_\_Enterobacteriaceae;g\_\_Escherichia;s\_\_Escherichia albertii | blast | 96.9 | iPHoP-RF;93.10 |
| NC\_000913.3\|provirus\_1412000\_1434224 | RS\_GCF\_002075345.1 | d\_\_Bacteria;p\_\_Pseudomonadota;c\_\_Gammaproteobacteria;o\_\_Enterobacterales;f\_\_Enterobacteriaceae;g\_\_Citrobacter;s\_\_Citrobacter braakii | blast | 96.9 | None |
| NC\_000913.3\|provirus\_1412000\_1434224 | RS\_GCF\_002900365.1 | d\_\_Bacteria;p\_\_Pseudomonadota;c\_\_Gammaproteobacteria;o\_\_Enterobacterales;f\_\_Enterobacteriaceae;g\_\_Escherichia;s\_\_Escherichia marmotae | blast | 96.9 | iPHoP-RF;95.40 |
| NC\_000913.3\|provirus\_1412000\_1434224 | RS\_GCF\_002925905.1 | d\_\_Bacteria;p\_\_Pseudomonadota;c\_\_Gammaproteobacteria;o\_\_Enterobacterales;f\_\_Enterobacteriaceae;g\_\_Klebsiella;s\_\_Klebsiella michiganensis | blast | 96.9 | None |
| NC\_000913.3\|provirus\_1412000\_1434224 | RS\_GCF\_003697165.2 | d\_\_Bacteria;p\_\_Pseudomonadota;c\_\_Gammaproteobacteria;o\_\_Enterobacterales;f\_\_Enterobacteriaceae;g\_\_Escherichia;s\_\_Escherichia coli | blast | 96.9 | iPHoP-RF;95.40 |
| NC\_000913.3\|provirus\_1412000\_1434224 | RS\_GCF\_005843885.1 | d\_\_Bacteria;p\_\_Pseudomonadota;c\_\_Gammaproteobacteria;o\_\_Enterobacterales;f\_\_Enterobacteriaceae;g\_\_Escherichia;s\_\_Escherichia sp005843885 | iPHoP-RF | 96.1 | None |
| NC\_000913.3\|provirus\_1412000\_1434224 | RS\_GCF\_002965065.1 | d\_\_Bacteria;p\_\_Pseudomonadota;c\_\_Gammaproteobacteria;o\_\_Enterobacterales;f\_\_Enterobacteriaceae;g\_\_Escherichia;s\_\_Escherichia sp002965065 | iPHoP-RF | 95.4 | None |
| NC\_000913.3\|provirus\_1412000\_1434224 | RS\_GCF\_014836715.1 | d\_\_Bacteria;p\_\_Pseudomonadota;c\_\_Gammaproteobacteria;o\_\_Enterobacterales;f\_\_Enterobacteriaceae;g\_\_Escherichia;s\_\_Escherichia whittamii | iPHoP-RF | 95.4 | None |
| NC\_000913.3\|provirus\_1412000\_1434224 | RS\_GCF\_000026225.1 | d\_\_Bacteria;p\_\_Pseudomonadota;c\_\_Gammaproteobacteria;o\_\_Enterobacterales;f\_\_Enterobacteriaceae;g\_\_Escherichia;s\_\_Escherichia fergusonii | iPHoP-RF | 95.1 | None |
| NC\_000913.3\|provirus\_1412000\_1434224 | RS\_GCF\_011881725.1 | d\_\_Bacteria;p\_\_Pseudomonadota;c\_\_Gammaproteobacteria;o\_\_Enterobacterales;f\_\_Enterobacteriaceae;g\_\_Escherichia;s\_\_Escherichia coli\_E | iPHoP-RF | 95.1 | None |
| NC\_000913.3\|provirus\_1412000\_1434224 | RS\_GCF\_002042885.1 | d\_\_Bacteria;p\_\_Pseudomonadota;c\_\_Gammaproteobacteria;o\_\_Enterobacterales;f\_\_Enterobacteriaceae;g\_\_Citrobacter;s\_\_Citrobacter portucalensis | blast | 94.9 | None |
| NC\_000913.3\|provirus\_1412000\_1434224 | RS\_GCF\_004211955.1 | d\_\_Bacteria;p\_\_Pseudomonadota;c\_\_Gammaproteobacteria;o\_\_Enterobacterales;f\_\_Enterobacteriaceae;g\_\_Escherichia;s\_\_Escherichia sp004211955 | iPHoP-RF | 92.4 | None |
| NC\_000913.3\|provirus\_1627517\_1656149 | GB\_GCA\_021307345.1 | d\_\_Bacteria;p\_\_Pseudomonadota;c\_\_Gammaproteobacteria;o\_\_Enterobacterales;f\_\_Enterobacteriaceae;g\_\_Escherichia;s\_\_Escherichia ruysiae | blast | 96.9 | iPHoP-RF;93.10 |
| NC\_000913.3\|provirus\_1627517\_1656149 | RS\_GCF\_000759775.1 | d\_\_Bacteria;p\_\_Pseudomonadota;c\_\_Gammaproteobacteria;o\_\_Enterobacterales;f\_\_Enterobacteriaceae;g\_\_Escherichia;s\_\_Escherichia albertii | blast | 96.9 | iPHoP-RF;93.10 |
| NC\_000913.3\|provirus\_1627517\_1656149 | RS\_GCF\_002900365.1 | d\_\_Bacteria;p\_\_Pseudomonadota;c\_\_Gammaproteobacteria;o\_\_Enterobacterales;f\_\_Enterobacteriaceae;g\_\_Escherichia;s\_\_Escherichia marmotae | blast | 96.9 | iPHoP-RF;96.40 |
| NC\_000913.3\|provirus\_1627517\_1656149 | RS\_GCF\_003697165.2 | d\_\_Bacteria;p\_\_Pseudomonadota;c\_\_Gammaproteobacteria;o\_\_Enterobacterales;f\_\_Enterobacteriaceae;g\_\_Escherichia;s\_\_Escherichia coli | blast | 96.9 | iPHoP-RF;95.70 |
| NC\_000913.3\|provirus\_1627517\_1656149 | RS\_GCF\_011881725.1 | d\_\_Bacteria;p\_\_Pseudomonadota;c\_\_Gammaproteobacteria;o\_\_Enterobacterales;f\_\_Enterobacteriaceae;g\_\_Escherichia;s\_\_Escherichia coli\_E | blast | 96.9 | iPHoP-RF;95.40 |
| NC\_000913.3\|provirus\_1627517\_1656149 | RS\_GCF\_005843885.1 | d\_\_Bacteria;p\_\_Pseudomonadota;c\_\_Gammaproteobacteria;o\_\_Enterobacterales;f\_\_Enterobacteriaceae;g\_\_Escherichia;s\_\_Escherichia sp005843885 | iPHoP-RF | 96.1 | None |
| NC\_000913.3\|provirus\_1627517\_1656149 | RS\_GCF\_002965065.1 | d\_\_Bacteria;p\_\_Pseudomonadota;c\_\_Gammaproteobacteria;o\_\_Enterobacterales;f\_\_Enterobacteriaceae;g\_\_Escherichia;s\_\_Escherichia sp002965065 | iPHoP-RF | 95.7 | None |
| NC\_000913.3\|provirus\_1627517\_1656149 | RS\_GCF\_004211955.1 | d\_\_Bacteria;p\_\_Pseudomonadota;c\_\_Gammaproteobacteria;o\_\_Enterobacterales;f\_\_Enterobacteriaceae;g\_\_Escherichia;s\_\_Escherichia sp004211955 | iPHoP-RF | 95.1 | None |
| NC\_000913.3\|provirus\_1627517\_1656149 | RS\_GCF\_000026225.1 | d\_\_Bacteria;p\_\_Pseudomonadota;c\_\_Gammaproteobacteria;o\_\_Enterobacterales;f\_\_Enterobacteriaceae;g\_\_Escherichia;s\_\_Escherichia fergusonii | iPHoP-RF | 94.4 | None |
| NC\_000913.3\|provirus\_1627517\_1656149 | RS\_GCF\_014836715.1 | d\_\_Bacteria;p\_\_Pseudomonadota;c\_\_Gammaproteobacteria;o\_\_Enterobacterales;f\_\_Enterobacteriaceae;g\_\_Escherichia;s\_\_Escherichia whittamii | iPHoP-RF | 92.8 | None |
| NC\_000913.3\|provirus\_2463012\_2476510 | GB\_GCA\_021307345.1 | d\_\_Bacteria;p\_\_Pseudomonadota;c\_\_Gammaproteobacteria;o\_\_Enterobacterales;f\_\_Enterobacteriaceae;g\_\_Escherichia;s\_\_Escherichia ruysiae | blast | 96.9 | iPHoP-RF;93.40 |
| NC\_000913.3\|provirus\_2463012\_2476510 | RS\_GCF\_000026225.1 | d\_\_Bacteria;p\_\_Pseudomonadota;c\_\_Gammaproteobacteria;o\_\_Enterobacterales;f\_\_Enterobacteriaceae;g\_\_Escherichia;s\_\_Escherichia fergusonii | blast | 96.9 | iPHoP-RF;92.10 |
| NC\_000913.3\|provirus\_2463012\_2476510 | RS\_GCF\_000759775.1 | d\_\_Bacteria;p\_\_Pseudomonadota;c\_\_Gammaproteobacteria;o\_\_Enterobacterales;f\_\_Enterobacteriaceae;g\_\_Escherichia;s\_\_Escherichia albertii | blast | 96.9 | iPHoP-RF;96.70 |
| NC\_000913.3\|provirus\_2463012\_2476510 | RS\_GCF\_002900365.1 | d\_\_Bacteria;p\_\_Pseudomonadota;c\_\_Gammaproteobacteria;o\_\_Enterobacterales;f\_\_Enterobacteriaceae;g\_\_Escherichia;s\_\_Escherichia marmotae | blast | 96.9 | iPHoP-RF;94.70 |
| NC\_000913.3\|provirus\_2463012\_2476510 | RS\_GCF\_003697165.2 | d\_\_Bacteria;p\_\_Pseudomonadota;c\_\_Gammaproteobacteria;o\_\_Enterobacterales;f\_\_Enterobacteriaceae;g\_\_Escherichia;s\_\_Escherichia coli | blast | 96.9 | iPHoP-RF;94.10 |
| NC\_000913.3\|provirus\_2463012\_2476510 | RS\_GCF\_004211955.1 | d\_\_Bacteria;p\_\_Pseudomonadota;c\_\_Gammaproteobacteria;o\_\_Enterobacterales;f\_\_Enterobacteriaceae;g\_\_Escherichia;s\_\_Escherichia sp004211955 | blast | 96.9 | iPHoP-RF;94.70 |
| NC\_000913.3\|provirus\_2463012\_2476510 | RS\_GCF\_011881725.1 | d\_\_Bacteria;p\_\_Pseudomonadota;c\_\_Gammaproteobacteria;o\_\_Enterobacterales;f\_\_Enterobacteriaceae;g\_\_Escherichia;s\_\_Escherichia coli\_E | blast | 96.9 | iPHoP-RF;94.10 |
| NC\_000913.3\|provirus\_2463012\_2476510 | RS\_GCF\_005843885.1 | d\_\_Bacteria;p\_\_Pseudomonadota;c\_\_Gammaproteobacteria;o\_\_Enterobacterales;f\_\_Enterobacteriaceae;g\_\_Escherichia;s\_\_Escherichia sp005843885 | iPHoP-RF | 94.7 | None |
| NC\_000913.3\|provirus\_2463012\_2476510 | RS\_GCF\_002965065.1 | d\_\_Bacteria;p\_\_Pseudomonadota;c\_\_Gammaproteobacteria;o\_\_Enterobacterales;f\_\_Enterobacteriaceae;g\_\_Escherichia;s\_\_Escherichia sp002965065 | iPHoP-RF | 93.7 | None |
| NC\_000913.3\|provirus\_2463012\_2476510 | RS\_GCF\_014836715.1 | d\_\_Bacteria;p\_\_Pseudomonadota;c\_\_Gammaproteobacteria;o\_\_Enterobacterales;f\_\_Enterobacteriaceae;g\_\_Escherichia;s\_\_Escherichia whittamii | iPHoP-RF | 92.8 | None |
| NC\_000913.3\|provirus\_563848\_584430 | GB\_GCA\_021307345.1 | d\_\_Bacteria;p\_\_Pseudomonadota;c\_\_Gammaproteobacteria;o\_\_Enterobacterales;f\_\_Enterobacteriaceae;g\_\_Escherichia;s\_\_Escherichia ruysiae | blast | 96.9 | iPHoP-RF;88.20 |
| NC\_000913.3\|provirus\_563848\_584430 | RS\_GCF\_000759775.1 | d\_\_Bacteria;p\_\_Pseudomonadota;c\_\_Gammaproteobacteria;o\_\_Enterobacterales;f\_\_Enterobacteriaceae;g\_\_Escherichia;s\_\_Escherichia albertii | blast | 96.9 | iPHoP-RF;89.20 |
| NC\_000913.3\|provirus\_563848\_584430 | RS\_GCF\_002900365.1 | d\_\_Bacteria;p\_\_Pseudomonadota;c\_\_Gammaproteobacteria;o\_\_Enterobacterales;f\_\_Enterobacteriaceae;g\_\_Escherichia;s\_\_Escherichia marmotae | blast | 96.9 | iPHoP-RF;88.50 |
| NC\_000913.3\|provirus\_563848\_584430 | RS\_GCF\_002925905.1 | d\_\_Bacteria;p\_\_Pseudomonadota;c\_\_Gammaproteobacteria;o\_\_Enterobacterales;f\_\_Enterobacteriaceae;g\_\_Klebsiella;s\_\_Klebsiella michiganensis | blast | 96.9 | None |
| NC\_000913.3\|provirus\_563848\_584430 | RS\_GCF\_003697165.2 | d\_\_Bacteria;p\_\_Pseudomonadota;c\_\_Gammaproteobacteria;o\_\_Enterobacterales;f\_\_Enterobacteriaceae;g\_\_Escherichia;s\_\_Escherichia coli | blast | 96.9 | iPHoP-RF;87.00 |
| NC\_000913.3\|provirus\_563848\_584430 | RS\_GCF\_002075345.1 | d\_\_Bacteria;p\_\_Pseudomonadota;c\_\_Gammaproteobacteria;o\_\_Enterobacterales;f\_\_Enterobacteriaceae;g\_\_Citrobacter;s\_\_Citrobacter braakii | blast | 96.5 | None |

##### 2.1.6 AMG Predictions

Prediction of auxiliary metabolic genes in the prophage contigs.

| protein | scaffold | amg\_ko | amg\_ko\_name | pfam | pfam\_name |
| --- | --- | --- | --- | --- | --- |
| NC\_000913.3\|provirus\_1196867\_1213107\_23 | NC\_000913.3\|provirus\_1196867\_1213107 | K00031 | IDH1, IDH2, icd; isocitrate dehydrogenase [EC:1.1.1.42] | PF00180.20 | Isocitrate/isopropylmalate dehydrogenase |

## 2.2 NC\_002737

##### 2.2.1 Host Genome

**GTDB-Tk taxonomy**:

classification

d\_\_Bacteria;p\_\_Bacillota;c\_\_Bacilli;o\_\_Lactobacillales;f\_\_Streptococcaceae;g\_\_Streptococcus;s\_\_Streptococcus
pyogenes

**CheckM2 Summary**:

| total\_contigs | contig\_n50 | completeness | contamination |
| --- | --- | --- | --- |
| 1 | 1852433 | 99.99 | 0.48 |

**Defense-Finder Systems**:

| sys\_id | type | subtype | sys\_beg | sys\_end | protein\_in\_syst | genes\_count | name\_of\_profiles\_in\_sys |
| --- | --- | --- | --- | --- | --- | --- | --- |
| NC\_002737.2\_AbiAlpha\_1 | AbiAlpha | AbiAlpha | NC\_002737.2\_745 | NC\_002737.2\_745 | NC\_002737.2\_745 | 1 | AbiAlpha\_\_AbiAlpha |
| NC\_002737.2\_VP1853\_2 | VP1853 | VP1853 | NC\_002737.2\_750 | NC\_002737.2\_750 | NC\_002737.2\_750 | 1 | VP1853\_\_VP1853 |
| NC\_002737.2\_CAS\_Class2-Subtype-II-A\_7 | Cas | CAS\_Class2-Subtype-II-A | NC\_002737.2\_832 | NC\_002737.2\_835 | NC\_002737.2\_832,NC\_002737.2\_833,NC\_002737.2\_834,NC\_002737.2\_835 | 4 | cas1\_I\_II\_III\_IV\_V\_VI\_5,cas2\_I\_II\_III\_IV\_V\_VI\_6,cas9\_II-A\_II-B\_II-C\_3,csn2\_II-A\_4 |
| NC\_002737.2\_CAS\_Class1-Subtype-I-C\_5 | Cas | CAS\_Class1-Subtype-I-C | NC\_002737.2\_1263 | NC\_002737.2\_1269 | NC\_002737.2\_1263,NC\_002737.2\_1264,NC\_002737.2\_1265,NC\_002737.2\_1266,NC\_002737.2\_1267,NC\_002737.2\_1268,NC\_002737.2\_1269 | 7 | cas1\_I\_II\_III\_IV\_V\_VI\_6,cas2\_I\_II\_III\_IV\_V\_VI\_5,cas3\_I\_5,cas4\_I\_II\_III\_IV\_V\_VI\_4,cas5\_I-C\_11,cas7\_I-C\_13,cas8c\_I-C\_1 |
| NC\_002737.2\_RM\_Type\_I\_4 | RM | RM\_Type\_I | NC\_002737.2\_1533 | NC\_002737.2\_1535 | NC\_002737.2\_1533,NC\_002737.2\_1534,NC\_002737.2\_1535 | 3 | RM\_\_Type\_I\_MTases\_FAM\_0,RM\_\_Type\_I\_REases\_FAM\_0.einsi\_trimmed,RM\_\_Type\_I\_S\_51 |

**Genomad and CheckV Summary**:

| seq\_name | length | gene\_count | viral\_genes | checkv\_quality | miuvig\_quality | taxonomy | topology | coordinates |
| --- | --- | --- | --- | --- | --- | --- | --- | --- |
| NC\_002737.2\|provirus\_529627\_569283 | 39657 | 46 | 41 | High-quality | High-quality | Viruses;Duplodnaviria;Heunggongvirae;Uroviricota;Caudoviricetes;; | Provirus | 529627-569283 |
| NC\_002737.2\|provirus\_777501\_820593 | 43093 | 65 | 42 | High-quality | High-quality | Viruses;Duplodnaviria;Heunggongvirae;Uroviricota;Caudoviricetes;; | Provirus | 777501-820593 |
| NC\_002737.2\|provirus\_1186916\_1222544 | 35629 | 52 | 39 | High-quality | High-quality | Viruses;Duplodnaviria;Heunggongvirae;Uroviricota;Caudoviricetes;; | Provirus | 1186916-1222544 |

**Host Genome Ideogram with Phages**:

In
this circular plot, **“c”** indicates the contig, and the
number that follows (e.g., **c1**) represents the contig
number. If multiple contigs are present in the genome, each will be
shown with a distinct label (e.g., **c1**,
**c2**, etc.).

##### 2.2.2 Prophages

**Select prophage to show:**

###### 2.2.2.1 Phage ID: NC\_002737.2|provirus\_529627\_569283

**Phage-Host Genome Ideogram:**

**Genes Annotation (geNomad):**

| gene | length | marker | annotation\_accessions | annotation\_description |
| --- | --- | --- | --- | --- |
| NC\_002737.2\|provirus\_529627\_569283\_517 | 1416 | GENOMAD.022372.VV | PF04708;COG1961;K14060 | Site-specific DNA recombinase related to the DNA invertase Pin |
| NC\_002737.2\|provirus\_529627\_569283\_518 | 366 | GENOMAD.142683.VV | PF11446;TIGR04165;COG4888 | Cys-rich peptide, TIGR04165 family |
| NC\_002737.2\|provirus\_529627\_569283\_519 | 762 | GENOMAD.108504.VP | PF05866;PF00717;COG4570;TIGR02754 | Endodeoxyribonuclease RusA; Peptidase S24-like |
| NC\_002737.2\|provirus\_529627\_569283\_520 | 231 | NA | NA | NA |
| NC\_002737.2\|provirus\_529627\_569283\_521 | 138 | NA | NA | NA |
| NC\_002737.2\|provirus\_529627\_569283\_522 | 126 | GENOMAD.129654.VV | NA | NA |
| NC\_002737.2\|provirus\_529627\_569283\_523 | 228 | GENOMAD.108243.VV | NA | NA |
| NC\_002737.2\|provirus\_529627\_569283\_524 | 1320 | GENOMAD.010825.VV | PF13175;TIGR00634;COG4717;K03546 | DNA repair protein RecN |
| NC\_002737.2\|provirus\_529627\_569283\_525 | 1092 | GENOMAD.141036.VV | PF13479;TIGR01618;K04484;COG0468 | phage nucleotide-binding protein |
| NC\_002737.2\|provirus\_529627\_569283\_526 | 423 | GENOMAD.092954.VV | NA | NA |
| NC\_002737.2\|provirus\_529627\_569283\_527 | 735 | GENOMAD.014438.VV | NA | NA |
| NC\_002737.2\|provirus\_529627\_569283\_528 | 636 | GENOMAD.005320.VV | PF05037 | NA |
| NC\_002737.2\|provirus\_529627\_569283\_529 | 1584 | NA | NA | NA |
| NC\_002737.2\|provirus\_529627\_569283\_530 | 630 | GENOMAD.116271.VP | PF07768 | PVL ORF-50-like family |
| NC\_002737.2\|provirus\_529627\_569283\_531 | 2274 | GENOMAD.168744.VP | K07505 | NA |
| NC\_002737.2\|provirus\_529627\_569283\_532 | 219 | GENOMAD.004447.VV | PF21825 | crAss001\_48 related protein |
| NC\_002737.2\|provirus\_529627\_569283\_533 | 396 | GENOMAD.028440.VV | PF05866;COG4570;K01160 | Holliday junction resolvase RusA (prophage-encoded endonuclease) |
| NC\_002737.2\|provirus\_529627\_569283\_534 | 222 | GENOMAD.053518.VV | NA | NA |
| NC\_002737.2\|provirus\_529627\_569283\_535 | 273 | GENOMAD.072347.VV | TIGR02209 | NA |
| NC\_002737.2\|provirus\_529627\_569283\_536 | 636 | GENOMAD.138295.VP | TIGR02987;COG0827 | NA |
| NC\_002737.2\|provirus\_529627\_569283\_537 | 435 | GENOMAD.111673.VV | PF05263;TIGR01637 | phage transcriptional regulator, ArpU family |
| NC\_002737.2\|provirus\_529627\_569283\_538 | 1167 | GENOMAD.038338.VV | COG3392 | NA |
| NC\_002737.2\|provirus\_529627\_569283\_539 | 477 | GENOMAD.003239.VV | PF05133;TIGR01538 | phage portal protein, SPP1 family |
| NC\_002737.2\|provirus\_529627\_569283\_540 | 1212 | GENOMAD.064889.VV | PF04466;PF17288;TIGR01547;COG1783;K06909 | phage terminase, large subunit, PBSX family |
| NC\_002737.2\|provirus\_529627\_569283\_541 | 1503 | GENOMAD.003916.VV | PF05133;TIGR01542 | phage portal protein, putative, A118 family |
| NC\_002737.2\|provirus\_529627\_569283\_542 | 1494 | GENOMAD.063181.VV | PF06152;TIGR01641 | Phage minor capsid protein 2 |
| NC\_002737.2\|provirus\_529627\_569283\_543 | 228 | GENOMAD.148778.VV | NA | NA |
| NC\_002737.2\|provirus\_529627\_569283\_544 | 267 | GENOMAD.076597.VV | NA | NA |
| NC\_002737.2\|provirus\_529627\_569283\_545 | 609 | GENOMAD.041104.VV | NA | NA |
| NC\_002737.2\|provirus\_529627\_569283\_546 | 819 | GENOMAD.013974.VV | PF20036;TIGR04387 | major capsid protein, N4-gp56 family |
| NC\_002737.2\|provirus\_529627\_569283\_547 | 417 | GENOMAD.034545.VV | PF11436 | NA |
| NC\_002737.2\|provirus\_529627\_569283\_548 | 333 | GENOMAD.114565.VV | PF10665 | Minor capsid protein |
| NC\_002737.2\|provirus\_529627\_569283\_549 | 357 | GENOMAD.008261.VV | PF11114 | Minor capsid protein |
| NC\_002737.2\|provirus\_529627\_569283\_550 | 399 | GENOMAD.034546.VV | PF12691 | Bacteriophage minor capsid protein |
| NC\_002737.2\|provirus\_529627\_569283\_551 | 471 | GENOMAD.028171.VV | PF16461;COG5437 | Lambda phage tail tube protein, TTP |
| NC\_002737.2\|provirus\_529627\_569283\_552 | 435 | GENOMAD.064813.VV | PF10666 | Phage tail assembly chaperone protein Gp14 ()A118 |
| NC\_002737.2\|provirus\_529627\_569283\_553 | 582 | GENOMAD.013935.VV | PF06854 | Bacteriophage Gp15 protein |
| NC\_002737.2\|provirus\_529627\_569283\_554 | 3261 | GENOMAD.002456.VV | COG3941 | NA |
| NC\_002737.2\|provirus\_529627\_569283\_555 | 717 | GENOMAD.006390.VV | PF20195;COG4722;TIGR01633 | Phage-related protein |
| NC\_002737.2\|provirus\_529627\_569283\_556 | 2145 | GENOMAD.049306.VV | COG4926 | Phage-related protein |
| NC\_002737.2\|provirus\_529627\_569283\_557 | 1014 | GENOMAD.093524.VV | PF07212 | Hyaluronidase protein (HylP) |
| NC\_002737.2\|provirus\_529627\_569283\_558 | 1887 | GENOMAD.033709.VV | PF07902 | gp58-like protein |
| NC\_002737.2\|provirus\_529627\_569283\_559 | 432 | GENOMAD.073534.VV | PF07761 | NA |
| NC\_002737.2\|provirus\_529627\_569283\_560 | 618 | GENOMAD.024672.VV | PF07104 | NA |
| NC\_002737.2\|provirus\_529627\_569283\_561 | 276 | GENOMAD.039653.VV | PF24073 | NA |
| NC\_002737.2\|provirus\_529627\_569283\_562 | 228 | GENOMAD.126098.VV | COG5546 | NA |
| NC\_002737.2\|provirus\_529627\_569283\_563 | 111 | GENOMAD.102275.VV | NA | NA |
| NC\_002737.2\|provirus\_529627\_569283\_564 | 1203 | GENOMAD.133734.VP | NA | NA |
| NC\_002737.2\|provirus\_529627\_569283\_565 | 708 | GENOMAD.223462.VP | PF02876;PF01123;K11040 | Staphylococcal/Streptococcal toxin, beta-grasp domain; Staphylococcal/Streptococcal toxin, OB-fold domain |

###### 2.2.2.2 Phage ID: NC\_002737.2|provirus\_777501\_820593

**Phage-Host Genome Ideogram:**

**Genes Annotation (geNomad):**

| gene | length | marker | annotation\_accessions | annotation\_description |
| --- | --- | --- | --- | --- |
| NC\_002737.2\|provirus\_777501\_820593\_756 | 1041 | NA | NA | NA |
| NC\_002737.2\|provirus\_777501\_820593\_757 | 1140 | GENOMAD.111131.VV | NA | NA |
| NC\_002737.2\|provirus\_777501\_820593\_758 | 810 | GENOMAD.221704.VP | PF02452;COG3692;K18841 | Uncharacterized protein YifN, PemK superfamily |
| NC\_002737.2\|provirus\_777501\_820593\_759 | 747 | NA | NA | NA |
| NC\_002737.2\|provirus\_777501\_820593\_760 | 159 | GENOMAD.097925.VV | NA | NA |
| NC\_002737.2\|provirus\_777501\_820593\_761 | 168 | GENOMAD.116752.VV | NA | NA |
| NC\_002737.2\|provirus\_777501\_820593\_762 | 840 | NA | NA | NA |
| NC\_002737.2\|provirus\_777501\_820593\_763 | 228 | NA | NA | NA |
| NC\_002737.2\|provirus\_777501\_820593\_764 | 207 | NA | NA | NA |
| NC\_002737.2\|provirus\_777501\_820593\_765 | 729 | GENOMAD.133073.VP | PF03374;COG3645 | Phage antirepressor protein YoqD, KilAC domain |
| NC\_002737.2\|provirus\_777501\_820593\_766 | 267 | GENOMAD.133830.VV | NA | NA |
| NC\_002737.2\|provirus\_777501\_820593\_767 | 807 | NA | NA | NA |
| NC\_002737.2\|provirus\_777501\_820593\_768 | 237 | GENOMAD.153962.VV | NA | NA |
| NC\_002737.2\|provirus\_777501\_820593\_769 | 240 | GENOMAD.050216.VV | NA | NA |
| NC\_002737.2\|provirus\_777501\_820593\_770 | 186 | GENOMAD.105558.VV | NA | NA |
| NC\_002737.2\|provirus\_777501\_820593\_771 | 240 | GENOMAD.136142.VP | PF13730;TIGR00373;COG4738;K07721 | transcription factor E |
| NC\_002737.2\|provirus\_777501\_820593\_772 | 387 | NA | NA | NA |
| NC\_002737.2\|provirus\_777501\_820593\_773 | 234 | GENOMAD.100896.VV | NA | NA |
| NC\_002737.2\|provirus\_777501\_820593\_774 | 141 | GENOMAD.079484.VV | NA | NA |
| NC\_002737.2\|provirus\_777501\_820593\_775 | 207 | GENOMAD.114751.VV | NA | NA |
| NC\_002737.2\|provirus\_777501\_820593\_776 | 330 | GENOMAD.040347.VV | NA | NA |
| NC\_002737.2\|provirus\_777501\_820593\_777 | 792 | GENOMAD.177891.VP | PF03837;TIGR01913;COG3723 | phage recombination protein Bet |
| NC\_002737.2\|provirus\_777501\_820593\_778 | 666 | GENOMAD.012371.VV | PF07083 | NA |
| NC\_002737.2\|provirus\_777501\_820593\_779 | 360 | GENOMAD.012371.VV | PF07083 | NA |
| NC\_002737.2\|provirus\_777501\_820593\_780 | 339 | GENOMAD.060064.VV | NA | NA |
| NC\_002737.2\|provirus\_777501\_820593\_781 | 513 | NA | NA | NA |
| NC\_002737.2\|provirus\_777501\_820593\_782 | 195 | GENOMAD.100776.VV | NA | NA |
| NC\_002737.2\|provirus\_777501\_820593\_783 | 246 | GENOMAD.105486.VP | PF09643;TIGR01671 | phage uncharacterized protein TIGR01671 |
| NC\_002737.2\|provirus\_777501\_820593\_784 | 462 | GENOMAD.003865.VV | PF11753;PF07852 | Protein of unknwon function (DUF3310) |
| NC\_002737.2\|provirus\_777501\_820593\_785 | 171 | GENOMAD.094280.VV | NA | NA |
| NC\_002737.2\|provirus\_777501\_820593\_786 | 441 | GENOMAD.111673.VV | PF05263;TIGR01637 | phage transcriptional regulator, ArpU family |
| NC\_002737.2\|provirus\_777501\_820593\_787 | 204 | NA | NA | NA |
| NC\_002737.2\|provirus\_777501\_820593\_788 | 915 | GENOMAD.141545.VV | PF20505 | Intracellular sensor of Lambda phage, Abi component |
| NC\_002737.2\|provirus\_777501\_820593\_789 | 462 | GENOMAD.146805.VV | NA | NA |
| NC\_002737.2\|provirus\_777501\_820593\_790 | 483 | GENOMAD.160265.VP | PF03592;COG3728;K07474 | Phage terminase, small subunit |
| NC\_002737.2\|provirus\_777501\_820593\_791 | 1290 | GENOMAD.113336.VP | PF04466;TIGR01547;COG1783 | phage terminase, large subunit, PBSX family |
| NC\_002737.2\|provirus\_777501\_820593\_792 | 363 | GENOMAD.033166.VV | PF05133;TIGR01538 | phage portal protein, SPP1 family |
| NC\_002737.2\|provirus\_777501\_820593\_793 | 1029 | GENOMAD.100877.VP | NA | NA |
| NC\_002737.2\|provirus\_777501\_820593\_794 | 1626 | GENOMAD.004769.VV | PF04233;COG5585;TIGR01641 | NAD+–asparagine ADP-ribosyltransferase |
| NC\_002737.2\|provirus\_777501\_820593\_795 | 180 | GENOMAD.137178.VV | NA | NA |
| NC\_002737.2\|provirus\_777501\_820593\_796 | 270 | GENOMAD.076597.VV | NA | NA |
| NC\_002737.2\|provirus\_777501\_820593\_797 | 171 | GENOMAD.179672.VV | PF16777;COG4453;TIGR04563 | Transcriptional regulator, RHH-like, CopG |
| NC\_002737.2\|provirus\_777501\_820593\_798 | 249 | GENOMAD.106940.VV | NA | NA |
| NC\_002737.2\|provirus\_777501\_820593\_799 | 747 | NA | NA | NA |
| NC\_002737.2\|provirus\_777501\_820593\_800 | 534 | GENOMAD.010937.VV | PF14265 | NA |
| NC\_002737.2\|provirus\_777501\_820593\_801 | 381 | GENOMAD.010435.VV | PF02924 | NA |
| NC\_002737.2\|provirus\_777501\_820593\_802 | 1050 | GENOMAD.004151.VV | PF03864 | Phage major capsid protein E |
| NC\_002737.2\|provirus\_777501\_820593\_803 | 267 | NA | NA | NA |
| NC\_002737.2\|provirus\_777501\_820593\_804 | 354 | GENOMAD.020971.VV | PF11436;TIGR01560 | NA |
| NC\_002737.2\|provirus\_777501\_820593\_805 | 309 | GENOMAD.042575.VV | NA | NA |
| NC\_002737.2\|provirus\_777501\_820593\_806 | 366 | GENOMAD.011013.VV | PF11114;TIGR01725;COG5005 | phage protein, HK97 gp10 family |
| NC\_002737.2\|provirus\_777501\_820593\_807 | 390 | GENOMAD.023411.VV | PF16807 | NA |
| NC\_002737.2\|provirus\_777501\_820593\_808 | 654 | GENOMAD.018762.VV | TIGR02126;COG5437 | phage major tail protein, TP901-1 family |
| NC\_002737.2\|provirus\_777501\_820593\_809 | 354 | GENOMAD.019874.VV | PF12363 | Phage tail assembly chaperone protein, TAC |
| NC\_002737.2\|provirus\_777501\_820593\_810 | 330 | GENOMAD.025799.VV | NA | NA |
| NC\_002737.2\|provirus\_777501\_820593\_811 | 3636 | GENOMAD.013432.VV | NA | NA |
| NC\_002737.2\|provirus\_777501\_820593\_812 | 780 | GENOMAD.016670.VV | PF20195;TIGR01633;COG4722 | putative phage tail component, N-terminal domain |
| NC\_002737.2\|provirus\_777501\_820593\_813 | 2049 | GENOMAD.130674.VP | COG4926 | Phage-related protein |
| NC\_002737.2\|provirus\_777501\_820593\_814 | 1119 | GENOMAD.093524.VV | PF07212 | Hyaluronidase protein (HylP) |
| NC\_002737.2\|provirus\_777501\_820593\_815 | 1911 | GENOMAD.033709.VV | PF07902 | gp58-like protein |
| NC\_002737.2\|provirus\_777501\_820593\_816 | 162 | GENOMAD.181912.VC | NA | NA |
| NC\_002737.2\|provirus\_777501\_820593\_817 | 612 | GENOMAD.024672.VV | PF07104 | NA |
| NC\_002737.2\|provirus\_777501\_820593\_818 | 297 | GENOMAD.084276.VV | NA | NA |
| NC\_002737.2\|provirus\_777501\_820593\_819 | 186 | GENOMAD.004934.VV | PF16945 | Putative lactococcus lactis phage r1t holin |
| NC\_002737.2\|provirus\_777501\_820593\_820 | 108 | GENOMAD.102275.VV | NA | NA |
| NC\_002737.2\|provirus\_777501\_820593\_821 | 1335 | GENOMAD.006162.VV | K07273;COG3757 | Lyzozyme M1 (1,4-beta-N-acetylmuramidase), GH25 family |
| NC\_002737.2\|provirus\_777501\_820593\_822 | 678 | GENOMAD.223611.VP | PF02876;PF01123;K11040 | Staphylococcal/Streptococcal toxin, beta-grasp domain; Staphylococcal/Streptococcal toxin, OB-fold domain |
| NC\_002737.2\|provirus\_777501\_820593\_823 | 711 | GENOMAD.224010.VP | PF02876 | Staphylococcal/Streptococcal toxin, beta-grasp domain |

###### 2.2.2.3 Phage ID: NC\_002737.2|provirus\_1186916\_1222544

**Phage-Host Genome Ideogram:**

**Genes Annotation (geNomad):**

| gene | length | marker | annotation\_accessions | annotation\_description |
| --- | --- | --- | --- | --- |
| NC\_002737.2\|provirus\_1186916\_1222544\_1181 | 1863 | NA | NA | NA |
| NC\_002737.2\|provirus\_1186916\_1222544\_1182 | 183 | GENOMAD.222241.VP | PF24313 | Paratox |
| NC\_002737.2\|provirus\_1186916\_1222544\_1183 | 801 | NA | NA | NA |
| NC\_002737.2\|provirus\_1186916\_1222544\_1184 | 486 | NA | NA | NA |
| NC\_002737.2\|provirus\_1186916\_1222544\_1185 | 1206 | GENOMAD.133734.VP | NA | NA |
| NC\_002737.2\|provirus\_1186916\_1222544\_1186 | 111 | GENOMAD.102275.VV | NA | NA |
| NC\_002737.2\|provirus\_1186916\_1222544\_1187 | 228 | GENOMAD.126098.VV | COG5546 | NA |
| NC\_002737.2\|provirus\_1186916\_1222544\_1188 | 276 | GENOMAD.039653.VV | PF24073 | NA |
| NC\_002737.2\|provirus\_1186916\_1222544\_1189 | 618 | GENOMAD.024672.VV | PF07104 | NA |
| NC\_002737.2\|provirus\_1186916\_1222544\_1190 | 432 | GENOMAD.073534.VV | PF07761 | NA |
| NC\_002737.2\|provirus\_1186916\_1222544\_1191 | 1785 | GENOMAD.033709.VV | PF07902 | gp58-like protein |
| NC\_002737.2\|provirus\_1186916\_1222544\_1192 | 1113 | GENOMAD.093524.VV | PF07212 | Hyaluronidase protein (HylP) |
| NC\_002737.2\|provirus\_1186916\_1222544\_1193 | 1959 | GENOMAD.010194.VV | PF06605;COG4926 | Phage-related protein |
| NC\_002737.2\|provirus\_1186916\_1222544\_1194 | 696 | GENOMAD.004911.VV | PF20195;COG4722;TIGR01633 | Phage-related protein |
| NC\_002737.2\|provirus\_1186916\_1222544\_1195 | 2358 | GENOMAD.003531.VV | COG5412 | Phage-related protein |
| NC\_002737.2\|provirus\_1186916\_1222544\_1196 | 372 | GENOMAD.024158.VV | PF17318 | NA |
| NC\_002737.2\|provirus\_1186916\_1222544\_1197 | 264 | GENOMAD.021224.VV | NA | NA |
| NC\_002737.2\|provirus\_1186916\_1222544\_1198 | 594 | GENOMAD.073691.VV | NA | NA |
| NC\_002737.2\|provirus\_1186916\_1222544\_1199 | 336 | GENOMAD.019860.VV | PF16807 | NA |
| NC\_002737.2\|provirus\_1186916\_1222544\_1200 | 237 | GENOMAD.017784.VV | NA | NA |
| NC\_002737.2\|provirus\_1186916\_1222544\_1201 | 339 | GENOMAD.018809.VV | PF10665 | Minor capsid protein |
| NC\_002737.2\|provirus\_1186916\_1222544\_1202 | 423 | GENOMAD.076853.VV | PF09355 | Phage protein Gp19/Gp15/Gp42 |
| NC\_002737.2\|provirus\_1186916\_1222544\_1203 | 201 | GENOMAD.069443.VV | PF09124 | T4 recombination endonuclease VII, dimerisation |
| NC\_002737.2\|provirus\_1186916\_1222544\_1204 | 912 | GENOMAD.013457.VV | PF05065;TIGR01554;COG4653 | Phage capsid family |
| NC\_002737.2\|provirus\_1186916\_1222544\_1205 | 462 | GENOMAD.083647.VV | PF14265 | NA |
| NC\_002737.2\|provirus\_1186916\_1222544\_1206 | 1416 | GENOMAD.003419.VV | PF20441;PF03354;COG4626;TIGR01547;K17677 | Phage terminase-like protein, large subunit, contains N-terminal HTH domain |
| NC\_002737.2\|provirus\_1186916\_1222544\_1207 | 267 | GENOMAD.076597.VV | NA | NA |
| NC\_002737.2\|provirus\_1186916\_1222544\_1208 | 180 | NA | NA | NA |
| NC\_002737.2\|provirus\_1186916\_1222544\_1209 | 225 | GENOMAD.193246.VV | NA | NA |
| NC\_002737.2\|provirus\_1186916\_1222544\_1210 | 1494 | GENOMAD.003531.VV | COG5412 | Phage-related protein |
| NC\_002737.2\|provirus\_1186916\_1222544\_1211 | 1269 | GENOMAD.076853.VV | PF09355 | Phage protein Gp19/Gp15/Gp42 |
| NC\_002737.2\|provirus\_1186916\_1222544\_1212 | 357 | GENOMAD.072475.VV | PF12855;COG4338 | ECL1/2/3 zinc binding proteins |
| NC\_002737.2\|provirus\_1186916\_1222544\_1213 | 345 | GENOMAD.108097.VV | PF13395 | HNH endonuclease |
| NC\_002737.2\|provirus\_1186916\_1222544\_1214 | 420 | GENOMAD.024790.VV | PF07374;TIGR01636;COG2739 | phage transcriptional activator, RinA family |
| NC\_002737.2\|provirus\_1186916\_1222544\_1215 | 636 | GENOMAD.138295.VP | TIGR02987;COG0827 | NA |
| NC\_002737.2\|provirus\_1186916\_1222544\_1216 | 270 | GENOMAD.072347.VV | TIGR02209 | NA |
| NC\_002737.2\|provirus\_1186916\_1222544\_1217 | 513 | NA | NA | NA |
| NC\_002737.2\|provirus\_1186916\_1222544\_1218 | 393 | GENOMAD.060064.VV | NA | NA |
| NC\_002737.2\|provirus\_1186916\_1222544\_1219 | 798 | GENOMAD.161534.VP | PF12684;TIGR01896 | PDDEXK-like domain of unknown function (DUF3799) |
| NC\_002737.2\|provirus\_1186916\_1222544\_1220 | 201 | GENOMAD.148606.VV | PF06289 | NA |
| NC\_002737.2\|provirus\_1186916\_1222544\_1221 | 990 | GENOMAD.091877.VP | PF03837;COG3723;K07455;TIGR00616 | Recombinational DNA repair protein RecT |
| NC\_002737.2\|provirus\_1186916\_1222544\_1222 | 333 | GENOMAD.040347.VV | NA | NA |
| NC\_002737.2\|provirus\_1186916\_1222544\_1223 | 207 | GENOMAD.114751.VV | NA | NA |
| NC\_002737.2\|provirus\_1186916\_1222544\_1224 | 141 | GENOMAD.079484.VV | NA | NA |
| NC\_002737.2\|provirus\_1186916\_1222544\_1225 | 234 | GENOMAD.100896.VV | NA | NA |
| NC\_002737.2\|provirus\_1186916\_1222544\_1226 | 387 | NA | NA | NA |
| NC\_002737.2\|provirus\_1186916\_1222544\_1227 | 255 | GENOMAD.136142.VP | PF13730;TIGR00373;COG4738;K07721 | transcription factor E |
| NC\_002737.2\|provirus\_1186916\_1222544\_1228 | 186 | GENOMAD.105558.VV | NA | NA |
| NC\_002737.2\|provirus\_1186916\_1222544\_1229 | 312 | GENOMAD.050216.VV | NA | NA |
| NC\_002737.2\|provirus\_1186916\_1222544\_1230 | 132 | GENOMAD.114759.VV | NA | NA |
| NC\_002737.2\|provirus\_1186916\_1222544\_1231 | 213 | GENOMAD.225776.VP | PF05339;TIGR00673;COG3620;K22299 | cyanase |
| NC\_002737.2\|provirus\_1186916\_1222544\_1232 | 756 | GENOMAD.091877.VP | PF03837;COG3723;K07455;TIGR00616 | Recombinational DNA repair protein RecT |
| NC\_002737.2\|provirus\_1186916\_1222544\_1233 | 519 | GENOMAD.064488.VV | PF08797 | HIRAN domain |
| NC\_002737.2\|provirus\_1186916\_1222544\_1234 | 1143 | GENOMAD.111131.VV | NA | NA |

##### 2.2.3 vOTUs

When more than 1 phage is detected in the host genome, we perform a
clustering step using the tool dRep.

A threshold of 0.95 was applied to the ANI similarity index to define
clusters of virus operational taxonomic units (vOTUs).

**Final cluster designations**

| genomad\_summary$seq\_name | genome | secondary\_cluster | threshold | cluster\_method | comparison\_algorithm | primary\_cluster |
| --- | --- | --- | --- | --- | --- | --- |
| NC\_002737.2\|provirus\_529627\_569283 | sequence\_000001.fasta.fasta | 1\_0 | 0.05 | average | ANImf | 1 |
| NC\_002737.2\|provirus\_777501\_820593 | sequence\_000002.fasta.fasta | 2\_0 | 0.05 | average | ANImf | 2 |
| NC\_002737.2\|provirus\_1186916\_1222544 | sequence\_000000.fasta.fasta | 3\_0 | 0.05 | average | ANImf | 3 |

**Primary clustering plot**

##### 2.2.4 Virulence Genes

Screening of virulence genes present in the prophage contigs.

| sequence | start | end | strand | gene | coverage | coverage\_map | gaps | percent\_coverage | percent\_identity | database | accession | product | resistance |
| --- | --- | --- | --- | --- | --- | --- | --- | --- | --- | --- | --- | --- | --- |
| NC\_002737.2\|provirus\_1186916\_1222544 | 2808 | 3614 |  | mf3 | 1-807/807 | =============== | 0/0 | 100 | 100.00 | vfdb | NP\_269520 | (mf3) deoxyribonuclease [DNase (VF0252)] [Streptococcus pyogenes M1 GAS] | NA |
| NC\_002737.2\|provirus\_1186916\_1222544 | 9081 | 10193 |  | hylP | 1-1113/1113 | =============== | 0/0 | 100 | 100.00 | vfdb | NP\_269528 | (hylP) hyaluronidase phage associated [Hyaluronidase (VF0246)] [Streptococcus pyogenes M1 GAS] | NA |
| NC\_002737.2\|provirus\_529627\_569283 | 33078 | 34091 |  | hylP | 1-1014/1014 | =============== | 0/0 | 100 | 100.00 | vfdb | NP\_268936 | (hylP) hyaluronidase phage associated [Hyaluronidase (VF0246)] [Streptococcus pyogenes M1 GAS] | NA |
| NC\_002737.2\|provirus\_529627\_569283 | 38950 | 39657 |  | spec | 1-708/708 | =============== | 0/0 | 100 | 100.00 | vfdb | NP\_268943 | (spec) streptococcal exotoxin C precursor phage associated [Spes (VF0248)] [Streptococcus pyogenes M1 GAS] | NA |
| NC\_002737.2\|provirus\_777501\_820593 | 35641 | 36759 |  | hylP | 1-1119/1119 | =============== | 0/0 | 100 | 100.00 | vfdb | NP\_269179 | (hylP) hyaluronidase phage associated [Hyaluronidase (VF0246)] [Streptococcus pyogenes M1 GAS] | NA |
| NC\_002737.2\|provirus\_777501\_820593 | 41680 | 42357 |  | spei | 1-678/678 | =============== | 0/0 | 100 | 99.85 | vfdb | NP\_269185 | (spei) streptococcal exotoxin I precursor [Spes (VF0248)] [Streptococcus pyogenes M1 GAS] | NA |
| NC\_002737.2\|provirus\_777501\_820593 | 42383 | 43093 |  | speh | 1-711/711 | =============== | 0/0 | 100 | 100.00 | vfdb | NP\_269186 | (speh) streptococcal exotoxin H precursor [Spes (VF0248)] [Streptococcus pyogenes M1 GAS] | NA |


##### 2.2.5 Prophage-Host Prediction

Prediction of potential hosts for the prophage contigs.

| virus | host\_genome | host\_taxonomy | main\_method | confidence\_score | additional\_methods |
| --- | --- | --- | --- | --- | --- |
| NC\_002737.2\|provirus\_1186916\_1222544 | RS\_GCF\_000186445.1 | d\_\_Bacteria;p\_\_Bacillota;c\_\_Bacilli;o\_\_Lactobacillales;f\_\_Streptococcaceae;g\_\_Streptococcus;s\_\_Streptococcus agalactiae | CRISPR | 97.8 | blast;91.00 iPHoP-RF;88.90 |
| NC\_002737.2\|provirus\_1186916\_1222544 | RS\_GCF\_900459225.1 | d\_\_Bacteria;p\_\_Bacillota;c\_\_Bacilli;o\_\_Lactobacillales;f\_\_Streptococcaceae;g\_\_Streptococcus;s\_\_Streptococcus dysgalactiae | iPHoP-RF | 97.1 | blast;96.90 |
| NC\_002737.2\|provirus\_1186916\_1222544 | RS\_GCF\_002055535.1 | d\_\_Bacteria;p\_\_Bacillota;c\_\_Bacilli;o\_\_Lactobacillales;f\_\_Streptococcaceae;g\_\_Streptococcus;s\_\_Streptococcus pyogenes | blast | 96.9 | iPHoP-RF;96.40 |
| NC\_002737.2\|provirus\_1186916\_1222544 | RS\_GCF\_022354845.1 | d\_\_Bacteria;p\_\_Bacillota;c\_\_Bacilli;o\_\_Lactobacillales;f\_\_Streptococcaceae;g\_\_Streptococcus;s\_\_Streptococcus suis\_AA | CRISPR | 96.8 | None |
| NC\_002737.2\|provirus\_1186916\_1222544 | RS\_GCF\_900459405.1 | d\_\_Bacteria;p\_\_Bacillota;c\_\_Bacilli;o\_\_Lactobacillales;f\_\_Streptococcaceae;g\_\_Streptococcus;s\_\_Streptococcus hyointestinalis | CRISPR | 96.8 | iPHoP-RF;89.50 |
| NC\_002737.2\|provirus\_1186916\_1222544 | RS\_GCF\_900636575.1 | d\_\_Bacteria;p\_\_Bacillota;c\_\_Bacilli;o\_\_Lactobacillales;f\_\_Streptococcaceae;g\_\_Streptococcus;s\_\_Streptococcus canis | iPHoP-RF | 96.7 | blast;96.60 |
| NC\_002737.2\|provirus\_1186916\_1222544 | RS\_GCF\_000425025.1 | d\_\_Bacteria;p\_\_Bacillota;c\_\_Bacilli;o\_\_Lactobacillales;f\_\_Streptococcaceae;g\_\_Streptococcus;s\_\_Streptococcus castoreus | iPHoP-RF | 94.7 | None |
| NC\_002737.2\|provirus\_1186916\_1222544 | RS\_GCF\_001598035.1 | d\_\_Bacteria;p\_\_Bacillota;c\_\_Bacilli;o\_\_Lactobacillales;f\_\_Streptococcaceae;g\_\_Streptococcus;s\_\_Streptococcus halotolerans | iPHoP-RF | 93.7 | None |
| NC\_002737.2\|provirus\_1186916\_1222544 | RS\_GCF\_000154985.1 | d\_\_Bacteria;p\_\_Bacillota;c\_\_Bacilli;o\_\_Lactobacillales;f\_\_Streptococcaceae;g\_\_Streptococcus;s\_\_Streptococcus infantarius | iPHoP-RF | 93.4 | None |
| NC\_002737.2\|provirus\_1186916\_1222544 | RS\_GCF\_900475675.1 | d\_\_Bacteria;p\_\_Bacillota;c\_\_Bacilli;o\_\_Lactobacillales;f\_\_Streptococcaceae;g\_\_Streptococcus;s\_\_Streptococcus lutetiensis | iPHoP-RF | 93.1 | None |
| NC\_002737.2\|provirus\_1186916\_1222544 | RS\_GCF\_000187265.1 | d\_\_Bacteria;p\_\_Bacillota;c\_\_Bacilli;o\_\_Lactobacillales;f\_\_Streptococcaceae;g\_\_Streptococcus;s\_\_Streptococcus equinus | iPHoP-RF | 92.8 | None |
| NC\_002737.2\|provirus\_1186916\_1222544 | RS\_GCF\_900101445.1 | d\_\_Bacteria;p\_\_Bacillota;c\_\_Bacilli;o\_\_Lactobacillales;f\_\_Streptococcaceae;g\_\_Streptococcus;s\_\_Streptococcus equinus\_B | iPHoP-RF | 92.8 | None |
| NC\_002737.2\|provirus\_1186916\_1222544 | GB\_GCA\_000283635.1 | d\_\_Bacteria;p\_\_Bacillota;c\_\_Bacilli;o\_\_Lactobacillales;f\_\_Streptococcaceae;g\_\_Streptococcus;s\_\_Streptococcus macedonicus | iPHoP-RF | 92.4 | None |
| NC\_002737.2\|provirus\_1186916\_1222544 | GB\_GCA\_934196125.1 | d\_\_Bacteria;p\_\_Bacillota;c\_\_Bacilli;o\_\_Lactobacillales;f\_\_Streptococcaceae;g\_\_Streptococcus;s\_\_Streptococcus sp934196125 | iPHoP-RF | 92.4 | None |
| NC\_002737.2\|provirus\_1186916\_1222544 | GB\_GCA\_900637675.1 | d\_\_Bacteria;p\_\_Bacillota;c\_\_Bacilli;o\_\_Lactobacillales;f\_\_Streptococcaceae;g\_\_Streptococcus;s\_\_Streptococcus equi | iPHoP-RF | 92.1 | blast;62.80 |
| NC\_002737.2\|provirus\_1186916\_1222544 | RS\_GCF\_004843545.1 | d\_\_Bacteria;p\_\_Bacillota;c\_\_Bacilli;o\_\_Lactobacillales;f\_\_Streptococcaceae;g\_\_Streptococcus;s\_\_Streptococcus pasteurianus | iPHoP-RF | 91.8 | None |
| NC\_002737.2\|provirus\_1186916\_1222544 | RS\_GCF\_003337175.1 | d\_\_Bacteria;p\_\_Bacillota;c\_\_Bacilli;o\_\_Lactobacillales;f\_\_Streptococcaceae;g\_\_Streptococcus;s\_\_Streptococcus gallolyticus\_B | iPHoP-RF | 91.4 | None |
| NC\_002737.2\|provirus\_1186916\_1222544 | RS\_GCF\_009870755.1 | d\_\_Bacteria;p\_\_Bacillota;c\_\_Bacilli;o\_\_Lactobacillales;f\_\_Streptococcaceae;g\_\_Streptococcus;s\_\_Streptococcus halichoeri | iPHoP-RF | 91.1 | None |
| NC\_002737.2\|provirus\_1186916\_1222544 | RS\_GCF\_002000985.1 | d\_\_Bacteria;p\_\_Bacillota;c\_\_Bacilli;o\_\_Lactobacillales;f\_\_Streptococcaceae;g\_\_Streptococcus;s\_\_Streptococcus gallolyticus | iPHoP-RF | 90.8 | None |
| NC\_002737.2\|provirus\_1186916\_1222544 | RS\_GCF\_000188015.2 | d\_\_Bacteria;p\_\_Bacillota;c\_\_Bacilli;o\_\_Lactobacillales;f\_\_Streptococcaceae;g\_\_Streptococcus;s\_\_Streptococcus ictaluri | iPHoP-RF | 90.5 | blast;62.80 |
| NC\_002737.2\|provirus\_1186916\_1222544 | RS\_GCF\_000188055.2 | d\_\_Bacteria;p\_\_Bacillota;c\_\_Bacilli;o\_\_Lactobacillales;f\_\_Streptococcaceae;g\_\_Streptococcus;s\_\_Streptococcus urinalis | iPHoP-RF | 90.5 | None |
| NC\_002737.2\|provirus\_529627\_569283 | RS\_GCF\_000960035.1 | d\_\_Bacteria;p\_\_Bacillota;c\_\_Bacilli;o\_\_Lactobacillales;f\_\_Streptococcaceae;g\_\_Streptococcus;s\_\_Streptococcus oralis\_G | CRISPR | 98.6 | iPHoP-RF;97.10 |
| NC\_002737.2\|provirus\_529627\_569283 | RS\_GCF\_002055535.1 | d\_\_Bacteria;p\_\_Bacillota;c\_\_Bacilli;o\_\_Lactobacillales;f\_\_Streptococcaceae;g\_\_Streptococcus;s\_\_Streptococcus pyogenes | CRISPR | 98.6 | blast;97.50 iPHoP-RF;96.10 |
| NC\_002737.2\|provirus\_529627\_569283 | RS\_GCF\_002093545.1 | d\_\_Bacteria;p\_\_Bacillota;c\_\_Bacilli;o\_\_Lactobacillales;f\_\_Streptococcaceae;g\_\_Streptococcus;s\_\_Streptococcus oralis\_C | CRISPR | 98.6 | iPHoP-RF;96.10 |
| NC\_002737.2\|provirus\_529627\_569283 | RS\_GCF\_900636575.1 | d\_\_Bacteria;p\_\_Bacillota;c\_\_Bacilli;o\_\_Lactobacillales;f\_\_Streptococcaceae;g\_\_Streptococcus;s\_\_Streptococcus canis | CRISPR | 98.4 | iPHoP-RF;97.10 blast;96.70 |
| NC\_002737.2\|provirus\_529627\_569283 | RS\_GCF\_001937065.1 | d\_\_Bacteria;p\_\_Bacillota;c\_\_Bacilli;o\_\_Lactobacillales;f\_\_Streptococcaceae;g\_\_Streptococcus;s\_\_Streptococcus sp001937065 | CRISPR | 98.2 | iPHoP-RF;96.70 |
| NC\_002737.2\|provirus\_529627\_569283 | RS\_GCF\_902729355.1 | d\_\_Bacteria;p\_\_Bacillota;c\_\_Bacilli;o\_\_Lactobacillales;f\_\_Streptococcaceae;g\_\_Streptococcus;s\_\_Streptococcus sp902729355 | CRISPR | 98.1 | iPHoP-RF;95.40 |
| NC\_002737.2\|provirus\_529627\_569283 | RS\_GCF\_000186445.1 | d\_\_Bacteria;p\_\_Bacillota;c\_\_Bacilli;o\_\_Lactobacillales;f\_\_Streptococcaceae;g\_\_Streptococcus;s\_\_Streptococcus agalactiae | CRISPR | 97.6 | iPHoP-RF;91.40 blast;67.70 |
| NC\_002737.2\|provirus\_529627\_569283 | RS\_GCF\_000188035.1 | d\_\_Bacteria;p\_\_Bacillota;c\_\_Bacilli;o\_\_Lactobacillales;f\_\_Streptococcaceae;g\_\_Streptococcus;s\_\_Streptococcus pseudoporcinus | CRISPR | 97.6 | iPHoP-RF;96.40 |
| NC\_002737.2\|provirus\_529627\_569283 | RS\_GCF\_000220065.1 | d\_\_Bacteria;p\_\_Bacillota;c\_\_Bacilli;o\_\_Lactobacillales;f\_\_Streptococcaceae;g\_\_Streptococcus;s\_\_Streptococcus sp000220065 | iPHoP-RF | 97.1 | None |
| NC\_002737.2\|provirus\_529627\_569283 | RS\_GCF\_000379985.1 | d\_\_Bacteria;p\_\_Bacillota;c\_\_Bacilli;o\_\_Lactobacillales;f\_\_Streptococcaceae;g\_\_Streptococcus;s\_\_Streptococcus caballi | iPHoP-RF | 97.1 | None |
| NC\_002737.2\|provirus\_529627\_569283 | RS\_GCF\_000380105.1 | d\_\_Bacteria;p\_\_Bacillota;c\_\_Bacilli;o\_\_Lactobacillales;f\_\_Streptococcaceae;g\_\_Streptococcus;s\_\_Streptococcus orisratti | iPHoP-RF | 97.1 | None |
| NC\_002737.2\|provirus\_529627\_569283 | RS\_GCF\_000380125.1 | d\_\_Bacteria;p\_\_Bacillota;c\_\_Bacilli;o\_\_Lactobacillales;f\_\_Streptococcaceae;g\_\_Streptococcus;s\_\_Streptococcus ovis | iPHoP-RF | 97.1 | None |
| NC\_002737.2\|provirus\_529627\_569283 | RS\_GCF\_900459225.1 | d\_\_Bacteria;p\_\_Bacillota;c\_\_Bacilli;o\_\_Lactobacillales;f\_\_Streptococcaceae;g\_\_Streptococcus;s\_\_Streptococcus dysgalactiae | blast | 96.9 | iPHoP-RF;93.10 |
| NC\_002737.2\|provirus\_529627\_569283 | RS\_GCF\_000423745.1 | d\_\_Bacteria;p\_\_Bacillota;c\_\_Bacilli;o\_\_Lactobacillales;f\_\_Streptococcaceae;g\_\_Streptococcus;s\_\_Streptococcus plurextorum | iPHoP-RF | 96.7 | None |
| NC\_002737.2\|provirus\_529627\_569283 | RS\_GCF\_002953735.1 | d\_\_Bacteria;p\_\_Bacillota;c\_\_Bacilli;o\_\_Lactobacillales;f\_\_Streptococcaceae;g\_\_Streptococcus;s\_\_Streptococcus pluranimalium | iPHoP-RF | 96.7 | None |
| NC\_002737.2\|provirus\_529627\_569283 | RS\_GCF\_006739205.1 | d\_\_Bacteria;p\_\_Bacillota;c\_\_Bacilli;o\_\_Lactobacillales;f\_\_Streptococcaceae;g\_\_Streptococcus;s\_\_Streptococcus mutans | iPHoP-RF | 96.7 | None |
| NC\_002737.2\|provirus\_529627\_569283 | GB\_GCA\_945876895.1 | d\_\_Bacteria;p\_\_Bacillota;c\_\_Bacilli;o\_\_Lactobacillales;f\_\_Streptococcaceae;g\_\_Streptococcus;s\_\_Streptococcus sp945876895 | iPHoP-RF | 96.4 | None |
| NC\_002737.2\|provirus\_529627\_569283 | RS\_GCF\_000425025.1 | d\_\_Bacteria;p\_\_Bacillota;c\_\_Bacilli;o\_\_Lactobacillales;f\_\_Streptococcaceae;g\_\_Streptococcus;s\_\_Streptococcus castoreus | iPHoP-RF | 96.4 | None |
| NC\_002737.2\|provirus\_529627\_569283 | RS\_GCF\_002000985.1 | d\_\_Bacteria;p\_\_Bacillota;c\_\_Bacilli;o\_\_Lactobacillales;f\_\_Streptococcaceae;g\_\_Streptococcus;s\_\_Streptococcus gallolyticus | iPHoP-RF | 96.4 | None |
| NC\_002737.2\|provirus\_529627\_569283 | RS\_GCF\_002355215.1 | d\_\_Bacteria;p\_\_Bacillota;c\_\_Bacilli;o\_\_Lactobacillales;f\_\_Streptococcaceae;g\_\_Streptococcus;s\_\_Streptococcus troglodytae | iPHoP-RF | 96.4 | None |
| NC\_002737.2\|provirus\_529627\_569283 | RS\_GCF\_003337175.1 | d\_\_Bacteria;p\_\_Bacillota;c\_\_Bacilli;o\_\_Lactobacillales;f\_\_Streptococcaceae;g\_\_Streptococcus;s\_\_Streptococcus gallolyticus\_B | iPHoP-RF | 96.4 | None |
| NC\_002737.2\|provirus\_529627\_569283 | RS\_GCF\_900475415.1 | d\_\_Bacteria;p\_\_Bacillota;c\_\_Bacilli;o\_\_Lactobacillales;f\_\_Streptococcaceae;g\_\_Streptococcus;s\_\_Streptococcus porcinus | iPHoP-RF | 96.4 | None |
| NC\_002737.2\|provirus\_529627\_569283 | GB\_GCA\_900637675.1 | d\_\_Bacteria;p\_\_Bacillota;c\_\_Bacilli;o\_\_Lactobacillales;f\_\_Streptococcaceae;g\_\_Streptococcus;s\_\_Streptococcus equi | iPHoP-RF | 96.1 | blast;78.70 |
| NC\_002737.2\|provirus\_529627\_569283 | GB\_GCA\_934196125.1 | d\_\_Bacteria;p\_\_Bacillota;c\_\_Bacilli;o\_\_Lactobacillales;f\_\_Streptococcaceae;g\_\_Streptococcus;s\_\_Streptococcus sp934196125 | iPHoP-RF | 96.1 | None |
| NC\_002737.2\|provirus\_529627\_569283 | RS\_GCF\_000420785.1 | d\_\_Bacteria;p\_\_Bacillota;c\_\_Bacilli;o\_\_Lactobacillales;f\_\_Streptococcaceae;g\_\_Streptococcus;s\_\_Streptococcus hyovaginalis | iPHoP-RF | 96.1 | None |
| NC\_002737.2\|provirus\_529627\_569283 | RS\_GCF\_004843545.1 | d\_\_Bacteria;p\_\_Bacillota;c\_\_Bacilli;o\_\_Lactobacillales;f\_\_Streptococcaceae;g\_\_Streptococcus;s\_\_Streptococcus pasteurianus | iPHoP-RF | 96.1 | None |
| NC\_002737.2\|provirus\_529627\_569283 | GB\_GCA\_000283635.1 | d\_\_Bacteria;p\_\_Bacillota;c\_\_Bacilli;o\_\_Lactobacillales;f\_\_Streptococcaceae;g\_\_Streptococcus;s\_\_Streptococcus macedonicus | iPHoP-RF | 95.7 | None |
| NC\_002737.2\|provirus\_529627\_569283 | RS\_GCF\_000423765.1 | d\_\_Bacteria;p\_\_Bacillota;c\_\_Bacilli;o\_\_Lactobacillales;f\_\_Streptococcaceae;g\_\_Streptococcus;s\_\_Streptococcus porci | iPHoP-RF | 95.7 | blast;72.00 |
| NC\_002737.2\|provirus\_529627\_569283 | RS\_GCF\_003686955.1 | d\_\_Bacteria;p\_\_Bacillota;c\_\_Bacilli;o\_\_Lactobacillales;f\_\_Streptococcaceae;g\_\_Streptococcus;s\_\_Streptococcus hillyeri | iPHoP-RF | 95.7 | None |
| NC\_002737.2\|provirus\_529627\_569283 | RS\_GCF\_011039275.1 | d\_\_Bacteria;p\_\_Bacillota;c\_\_Bacilli;o\_\_Lactobacillales;f\_\_Streptococcaceae;g\_\_Streptococcus;s\_\_Streptococcus hyointestinalis\_A | iPHoP-RF | 95.7 | None |
| NC\_002737.2\|provirus\_529627\_569283 | RS\_GCF\_012277075.1 | d\_\_Bacteria;p\_\_Bacillota;c\_\_Bacilli;o\_\_Lactobacillales;f\_\_Streptococcaceae;g\_\_Streptococcus;s\_\_Streptococcus alactolyticus | iPHoP-RF | 95.7 | None |
| NC\_002737.2\|provirus\_529627\_569283 | RS\_GCF\_900459405.1 | d\_\_Bacteria;p\_\_Bacillota;c\_\_Bacilli;o\_\_Lactobacillales;f\_\_Streptococcaceae;g\_\_Streptococcus;s\_\_Streptococcus hyointestinalis | iPHoP-RF | 95.7 | None |
| NC\_002737.2\|provirus\_529627\_569283 | RS\_GCF\_901542335.1 | d\_\_Bacteria;p\_\_Bacillota;c\_\_Bacilli;o\_\_Lactobacillales;f\_\_Streptococcaceae;g\_\_Streptococcus;s\_\_Streptococcus porcinus\_A | iPHoP-RF | 95.7 | None |
| NC\_002737.2\|provirus\_529627\_569283 | RS\_GCF\_000187995.2 | d\_\_Bacteria;p\_\_Bacillota;c\_\_Bacilli;o\_\_Lactobacillales;f\_\_Streptococcaceae;g\_\_Streptococcus;s\_\_Streptococcus macacae | iPHoP-RF | 95.4 | None |
| NC\_002737.2\|provirus\_529627\_569283 | RS\_GCF\_000380145.1 | d\_\_Bacteria;p\_\_Bacillota;c\_\_Bacilli;o\_\_Lactobacillales;f\_\_Streptococcaceae;g\_\_Streptococcus;s\_\_Streptococcus thoraltensis | iPHoP-RF | 95.4 | None |
| NC\_002737.2\|provirus\_529627\_569283 | RS\_GCF\_012396585.1 | d\_\_Bacteria;p\_\_Bacillota;c\_\_Bacilli;o\_\_Lactobacillales;f\_\_Streptococcaceae;g\_\_Streptococcus;s\_\_Streptococcus ovuberis | iPHoP-RF | 95.4 | None |
| NC\_002737.2\|provirus\_529627\_569283 | RS\_GCF\_016908655.1 | d\_\_Bacteria;p\_\_Bacillota;c\_\_Bacilli;o\_\_Lactobacillales;f\_\_Streptococcaceae;g\_\_Streptococcus;s\_\_Streptococcus saliviloxodontae | iPHoP-RF | 95.4 | None |
| NC\_002737.2\|provirus\_529627\_569283 | RS\_GCF\_000188055.2 | d\_\_Bacteria;p\_\_Bacillota;c\_\_Bacilli;o\_\_Lactobacillales;f\_\_Streptococcaceae;g\_\_Streptococcus;s\_\_Streptococcus urinalis | iPHoP-RF | 95.1 | blast;66.30 |
| NC\_002737.2\|provirus\_529627\_569283 | RS\_GCF\_001598035.1 | d\_\_Bacteria;p\_\_Bacillota;c\_\_Bacilli;o\_\_Lactobacillales;f\_\_Streptococcaceae;g\_\_Streptococcus;s\_\_Streptococcus halotolerans | iPHoP-RF | 95.1 | None |
| NC\_002737.2\|provirus\_529627\_569283 | RS\_GCF\_009870755.1 | d\_\_Bacteria;p\_\_Bacillota;c\_\_Bacilli;o\_\_Lactobacillales;f\_\_Streptococcaceae;g\_\_Streptococcus;s\_\_Streptococcus halichoeri | iPHoP-RF | 95.1 | None |
| NC\_002737.2\|provirus\_529627\_569283 | RS\_GCF\_024814375.1 | d\_\_Bacteria;p\_\_Bacillota;c\_\_Bacilli;o\_\_Lactobacillales;f\_\_Streptococcaceae;g\_\_Streptococcus;s\_\_Streptococcus sp024814375 | iPHoP-RF | 95.1 | None |
| NC\_002737.2\|provirus\_529627\_569283 | RS\_GCF\_011421425.1 | d\_\_Bacteria;p\_\_Bacillota;c\_\_Bacilli;o\_\_Lactobacillales;f\_\_Streptococcaceae;g\_\_Streptococcus;s\_\_Streptococcus catagoni | iPHoP-RF | 94.7 | blast;66.30 |
| NC\_002737.2\|provirus\_529627\_569283 | RS\_GCF\_000380085.1 | d\_\_Bacteria;p\_\_Bacillota;c\_\_Bacilli;o\_\_Lactobacillales;f\_\_Streptococcaceae;g\_\_Streptococcus;s\_\_Streptococcus merionis | iPHoP-RF | 93.7 | None |
| NC\_002737.2\|provirus\_529627\_569283 | RS\_GCF\_000785785.1 | d\_\_Bacteria;p\_\_Bacillota;c\_\_Bacilli;o\_\_Lactobacillales;f\_\_Streptococcaceae;g\_\_Streptococcus;s\_\_Streptococcus uberis\_A | iPHoP-RF | 93.7 | None |
| NC\_002737.2\|provirus\_529627\_569283 | RS\_GCF\_003674745.1 | d\_\_Bacteria;p\_\_Bacillota;c\_\_Bacilli;o\_\_Lactobacillales;f\_\_Streptococcaceae;g\_\_Streptococcus;s\_\_Streptococcus iniae | iPHoP-RF | 93.7 | None |
| NC\_002737.2\|provirus\_529627\_569283 | RS\_GCF\_900475595.1 | d\_\_Bacteria;p\_\_Bacillota;c\_\_Bacilli;o\_\_Lactobacillales;f\_\_Streptococcaceae;g\_\_Streptococcus;s\_\_Streptococcus uberis | iPHoP-RF | 93.7 | None |
| NC\_002737.2\|provirus\_529627\_569283 | RS\_GCF\_019794555.1 | d\_\_Bacteria;p\_\_Bacillota;c\_\_Bacilli;o\_\_Lactobacillales;f\_\_Streptococcaceae;g\_\_Streptococcus;s\_\_Streptococcus suis\_AB | iPHoP-RF | 91.1 | None |
| NC\_002737.2\|provirus\_777501\_820593 | MGYG000003717 | d\_\_Bacteria;p\_\_Bacillota;c\_\_Bacilli;o\_\_Lactobacillales;f\_\_Streptococcaceae;g**Streptococcus;s** | CRISPR | 98.4 | iPHoP-RF;96.70 |
| NC\_002737.2\|provirus\_777501\_820593 | RS\_GCF\_004843545.1 | d\_\_Bacteria;p\_\_Bacillota;c\_\_Bacilli;o\_\_Lactobacillales;f\_\_Streptococcaceae;g\_\_Streptococcus;s\_\_Streptococcus pasteurianus | CRISPR | 98.4 | iPHoP-RF;94.10 |
| NC\_002737.2\|provirus\_777501\_820593 | GB\_GCA\_000283635.1 | d\_\_Bacteria;p\_\_Bacillota;c\_\_Bacilli;o\_\_Lactobacillales;f\_\_Streptococcaceae;g\_\_Streptococcus;s\_\_Streptococcus macedonicus | CRISPR | 98.3 | iPHoP-RF;88.50 |
| NC\_002737.2\|provirus\_777501\_820593 | RS\_GCF\_000186445.1 | d\_\_Bacteria;p\_\_Bacillota;c\_\_Bacilli;o\_\_Lactobacillales;f\_\_Streptococcaceae;g\_\_Streptococcus;s\_\_Streptococcus agalactiae | CRISPR | 98.3 | iPHoP-RF;80.30 blast;76.70 |
| NC\_002737.2\|provirus\_777501\_820593 | RS\_GCF\_002055535.1 | d\_\_Bacteria;p\_\_Bacillota;c\_\_Bacilli;o\_\_Lactobacillales;f\_\_Streptococcaceae;g\_\_Streptococcus;s\_\_Streptococcus pyogenes | blast | 98.0 | iPHoP-RF;93.70 |
| NC\_002737.2\|provirus\_777501\_820593 | RS\_GCF\_900636575.1 | d\_\_Bacteria;p\_\_Bacillota;c\_\_Bacilli;o\_\_Lactobacillales;f\_\_Streptococcaceae;g\_\_Streptococcus;s\_\_Streptococcus canis | CRISPR | 98.0 | blast;96.50 iPHoP-RF;95.40 |
| NC\_002737.2\|provirus\_777501\_820593 | RS\_GCF\_010120595.1 | d\_\_Bacteria;p\_\_Bacillota;c\_\_Bacilli;o\_\_Lactobacillales;f\_\_Streptococcaceae;g\_\_Streptococcus;s\_\_Streptococcus thermophilus | CRISPR | 97.9 | iPHoP-RF;96.10 |
| NC\_002737.2\|provirus\_777501\_820593 | RS\_GCF\_000220065.1 | d\_\_Bacteria;p\_\_Bacillota;c\_\_Bacilli;o\_\_Lactobacillales;f\_\_Streptococcaceae;g\_\_Streptococcus;s\_\_Streptococcus sp000220065 | iPHoP-RF | 97.1 | None |
| NC\_002737.2\|provirus\_777501\_820593 | RS\_GCF\_001697145.1 | d\_\_Bacteria;p\_\_Bacillota;c\_\_Bacilli;o\_\_Lactobacillales;f\_\_Streptococcaceae;g\_\_Streptococcus;s\_\_Streptococcus anginosus\_C | iPHoP-RF | 97.1 | None |
| NC\_002737.2\|provirus\_777501\_820593 | RS\_GCF\_009717815.1 | d\_\_Bacteria;p\_\_Bacillota;c\_\_Bacilli;o\_\_Lactobacillales;f\_\_Streptococcaceae;g\_\_Streptococcus;s\_\_Streptococcus parasanguinis\_F | iPHoP-RF | 97.1 | blast;83.70 |
| NC\_002737.2\|provirus\_777501\_820593 | RS\_GCF\_016648925.1 | d\_\_Bacteria;p\_\_Bacillota;c\_\_Bacilli;o\_\_Lactobacillales;f\_\_Streptococcaceae;g\_\_Streptococcus;s\_\_Streptococcus sp900766505 | iPHoP-RF | 97.1 | blast;83.70 |
| NC\_002737.2\|provirus\_777501\_820593 | RS\_GCF\_900636475.1 | d\_\_Bacteria;p\_\_Bacillota;c\_\_Bacilli;o\_\_Lactobacillales;f\_\_Streptococcaceae;g\_\_Streptococcus;s\_\_Streptococcus anginosus | iPHoP-RF | 97.1 | None |
| NC\_002737.2\|provirus\_777501\_820593 | RS\_GCF\_902167705.1 | d\_\_Bacteria;p\_\_Bacillota;c\_\_Bacilli;o\_\_Lactobacillales;f\_\_Streptococcaceae;g\_\_Streptococcus;s\_\_Streptococcus constellatus\_A | iPHoP-RF | 97.1 | None |
| NC\_002737.2\|provirus\_777501\_820593 | RS\_GCF\_943193075.1 | d\_\_Bacteria;p\_\_Bacillota;c\_\_Bacilli;o\_\_Lactobacillales;f\_\_Streptococcaceae;g\_\_Streptococcus;s\_\_Streptococcus parasanguinis\_E | iPHoP-RF | 97.1 | blast;83.70 |
| NC\_002737.2\|provirus\_777501\_820593 | RS\_GCF\_900459225.1 | d\_\_Bacteria;p\_\_Bacillota;c\_\_Bacilli;o\_\_Lactobacillales;f\_\_Streptococcaceae;g\_\_Streptococcus;s\_\_Streptococcus dysgalactiae | blast | 96.9 | iPHoP-RF;95.10 |
| NC\_002737.2\|provirus\_777501\_820593 | RS\_GCF\_000380125.1 | d\_\_Bacteria;p\_\_Bacillota;c\_\_Bacilli;o\_\_Lactobacillales;f\_\_Streptococcaceae;g\_\_Streptococcus;s\_\_Streptococcus ovis | iPHoP-RF | 96.7 | None |
| NC\_002737.2\|provirus\_777501\_820593 | RS\_GCF\_000423765.1 | d\_\_Bacteria;p\_\_Bacillota;c\_\_Bacilli;o\_\_Lactobacillales;f\_\_Streptococcaceae;g\_\_Streptococcus;s\_\_Streptococcus porci | iPHoP-RF | 96.7 | None |
| NC\_002737.2\|provirus\_777501\_820593 | RS\_GCF\_001598035.1 | d\_\_Bacteria;p\_\_Bacillota;c\_\_Bacilli;o\_\_Lactobacillales;f\_\_Streptococcaceae;g\_\_Streptococcus;s\_\_Streptococcus halotolerans | iPHoP-RF | 96.7 | None |
| NC\_002737.2\|provirus\_777501\_820593 | RS\_GCF\_023109675.1 | d\_\_Bacteria;p\_\_Bacillota;c\_\_Bacilli;o\_\_Lactobacillales;f\_\_Streptococcaceae;g\_\_Streptococcus;s\_\_Streptococcus parasanguinis\_I | iPHoP-RF | 96.7 | blast;83.70 |
| NC\_002737.2\|provirus\_777501\_820593 | RS\_GCF\_023167545.1 | d\_\_Bacteria;p\_\_Bacillota;c\_\_Bacilli;o\_\_Lactobacillales;f\_\_Streptococcaceae;g\_\_Streptococcus;s\_\_Streptococcus constellatus | iPHoP-RF | 96.7 | None |
| NC\_002737.2\|provirus\_777501\_820593 | RS\_GCF\_000423745.1 | d\_\_Bacteria;p\_\_Bacillota;c\_\_Bacilli;o\_\_Lactobacillales;f\_\_Streptococcaceae;g\_\_Streptococcus;s\_\_Streptococcus plurextorum | iPHoP-RF | 96.4 | None |
| NC\_002737.2\|provirus\_777501\_820593 | RS\_GCF\_003686955.1 | d\_\_Bacteria;p\_\_Bacillota;c\_\_Bacilli;o\_\_Lactobacillales;f\_\_Streptococcaceae;g\_\_Streptococcus;s\_\_Streptococcus hillyeri | iPHoP-RF | 96.4 | blast;70.10 |
| NC\_002737.2\|provirus\_777501\_820593 | RS\_GCF\_011039275.1 | d\_\_Bacteria;p\_\_Bacillota;c\_\_Bacilli;o\_\_Lactobacillales;f\_\_Streptococcaceae;g\_\_Streptococcus;s\_\_Streptococcus hyointestinalis\_A | iPHoP-RF | 96.4 | None |
| NC\_002737.2\|provirus\_777501\_820593 | RS\_GCF\_000380025.1 | d\_\_Bacteria;p\_\_Bacillota;c\_\_Bacilli;o\_\_Lactobacillales;f\_\_Streptococcaceae;g\_\_Streptococcus;s\_\_Streptococcus entericus | iPHoP-RF | 96.1 | None |
| NC\_002737.2\|provirus\_777501\_820593 | RS\_GCF\_000380145.1 | d\_\_Bacteria;p\_\_Bacillota;c\_\_Bacilli;o\_\_Lactobacillales;f\_\_Streptococcaceae;g\_\_Streptococcus;s\_\_Streptococcus thoraltensis | iPHoP-RF | 96.1 | None |
| NC\_002737.2\|provirus\_777501\_820593 | RS\_GCF\_900475675.1 | d\_\_Bacteria;p\_\_Bacillota;c\_\_Bacilli;o\_\_Lactobacillales;f\_\_Streptococcaceae;g\_\_Streptococcus;s\_\_Streptococcus lutetiensis | iPHoP-RF | 96.1 | blast;68.90 |
| NC\_002737.2\|provirus\_777501\_820593 | RS\_GCF\_000154985.1 | d\_\_Bacteria;p\_\_Bacillota;c\_\_Bacilli;o\_\_Lactobacillales;f\_\_Streptococcaceae;g\_\_Streptococcus;s\_\_Streptococcus infantarius | iPHoP-RF | 95.7 | None |
| NC\_002737.2\|provirus\_777501\_820593 | RS\_GCF\_000420785.1 | d\_\_Bacteria;p\_\_Bacillota;c\_\_Bacilli;o\_\_Lactobacillales;f\_\_Streptococcaceae;g\_\_Streptococcus;s\_\_Streptococcus hyovaginalis | iPHoP-RF | 95.7 | None |
| NC\_002737.2\|provirus\_777501\_820593 | RS\_GCF\_000425025.1 | d\_\_Bacteria;p\_\_Bacillota;c\_\_Bacilli;o\_\_Lactobacillales;f\_\_Streptococcaceae;g\_\_Streptococcus;s\_\_Streptococcus castoreus | iPHoP-RF | 95.4 | None |
| NC\_002737.2\|provirus\_777501\_820593 | RS\_GCF\_002000985.1 | d\_\_Bacteria;p\_\_Bacillota;c\_\_Bacilli;o\_\_Lactobacillales;f\_\_Streptococcaceae;g\_\_Streptococcus;s\_\_Streptococcus gallolyticus | iPHoP-RF | 95.4 | blast;68.90 |
| NC\_002737.2\|provirus\_777501\_820593 | GB\_GCA\_000440235.1 | d\_\_Bacteria;p\_\_Bacillota;c\_\_Bacilli;o\_\_Lactobacillales;f\_\_Streptococcaceae;g\_\_Streptococcus;s\_\_Streptococcus suis\_F | iPHoP-RF | 95.1 | None |
| NC\_002737.2\|provirus\_777501\_820593 | GB\_GCA\_934196125.1 | d\_\_Bacteria;p\_\_Bacillota;c\_\_Bacilli;o\_\_Lactobacillales;f\_\_Streptococcaceae;g\_\_Streptococcus;s\_\_Streptococcus sp934196125 | iPHoP-RF | 95.1 | None |
| NC\_002737.2\|provirus\_777501\_820593 | GB\_GCA\_002831545.1 | d\_\_Bacteria;p\_\_Bacillota;c\_\_Bacilli;o\_\_Lactobacillales;f\_\_Streptococcaceae;g\_\_Streptococcus;s\_\_Streptococcus suis\_P | iPHoP-RF | 94.7 | None |
| NC\_002737.2\|provirus\_777501\_820593 | RS\_GCF\_000785515.1 | d\_\_Bacteria;p\_\_Bacillota;c\_\_Bacilli;o\_\_Lactobacillales;f\_\_Streptococcaceae;g\_\_Streptococcus;s\_\_Streptococcus salivarius | iPHoP-RF | 94.7 | blast;68.90 |
| NC\_002737.2\|provirus\_777501\_820593 | RS\_GCF\_003337175.1 | d\_\_Bacteria;p\_\_Bacillota;c\_\_Bacilli;o\_\_Lactobacillales;f\_\_Streptococcaceae;g\_\_Streptococcus;s\_\_Streptococcus gallolyticus\_B | iPHoP-RF | 94.7 | None |
| NC\_002737.2\|provirus\_777501\_820593 | RS\_GCF\_900101445.1 | d\_\_Bacteria;p\_\_Bacillota;c\_\_Bacilli;o\_\_Lactobacillales;f\_\_Streptococcaceae;g\_\_Streptococcus;s\_\_Streptococcus equinus\_B | iPHoP-RF | 94.7 | None |
| NC\_002737.2\|provirus\_777501\_820593 | GB\_GCA\_945876895.1 | d\_\_Bacteria;p\_\_Bacillota;c\_\_Bacilli;o\_\_Lactobacillales;f\_\_Streptococcaceae;g\_\_Streptococcus;s\_\_Streptococcus sp945876895 | iPHoP-RF | 94.4 | None |
| NC\_002737.2\|provirus\_777501\_820593 | RS\_GCF\_002760245.1 | d\_\_Bacteria;p\_\_Bacillota;c\_\_Bacilli;o\_\_Lactobacillales;f\_\_Streptococcaceae;g\_\_Streptococcus;s\_\_Streptococcus suis\_I | iPHoP-RF | 94.4 | None |
| NC\_002737.2\|provirus\_777501\_820593 | RS\_GCF\_002953735.1 | d\_\_Bacteria;p\_\_Bacillota;c\_\_Bacilli;o\_\_Lactobacillales;f\_\_Streptococcaceae;g\_\_Streptococcus;s\_\_Streptococcus pluranimalium | iPHoP-RF | 94.4 | None |
| NC\_002737.2\|provirus\_777501\_820593 | RS\_GCF\_000380105.1 | d\_\_Bacteria;p\_\_Bacillota;c\_\_Bacilli;o\_\_Lactobacillales;f\_\_Streptococcaceae;g\_\_Streptococcus;s\_\_Streptococcus orisratti | iPHoP-RF | 94.1 | None |
| NC\_002737.2\|provirus\_777501\_820593 | RS\_GCF\_902702775.1 | d\_\_Bacteria;p\_\_Bacillota;c\_\_Bacilli;o\_\_Lactobacillales;f\_\_Streptococcaceae;g\_\_Streptococcus;s\_\_Streptococcus suis\_W | iPHoP-RF | 94.1 | None |
| NC\_002737.2\|provirus\_777501\_820593 | RS\_GCF\_902729355.1 | d\_\_Bacteria;p\_\_Bacillota;c\_\_Bacilli;o\_\_Lactobacillales;f\_\_Streptococcaceae;g\_\_Streptococcus;s\_\_Streptococcus sp902729355 | iPHoP-RF | 94.1 | None |
| NC\_002737.2\|provirus\_777501\_820593 | RS\_GCF\_000376985.1 | d\_\_Bacteria;p\_\_Bacillota;c\_\_Bacilli;o\_\_Lactobacillales;f\_\_Streptococcaceae;g\_\_Streptococcus;s\_\_Streptococcus henryi | iPHoP-RF | 93.7 | None |
| NC\_002737.2\|provirus\_777501\_820593 | RS\_GCF\_012396585.1 | d\_\_Bacteria;p\_\_Bacillota;c\_\_Bacilli;o\_\_Lactobacillales;f\_\_Streptococcaceae;g\_\_Streptococcus;s\_\_Streptococcus ovuberis | iPHoP-RF | 93.7 | None |
| NC\_002737.2\|provirus\_777501\_820593 | RS\_GCF\_016743335.1 | d\_\_Bacteria;p\_\_Bacillota;c\_\_Bacilli;o\_\_Lactobacillales;f\_\_Streptococcaceae;g\_\_Streptococcus;s\_\_Streptococcus suis\_Y | iPHoP-RF | 93.7 | None |
| NC\_002737.2\|provirus\_777501\_820593 | RS\_GCF\_900459405.1 | d\_\_Bacteria;p\_\_Bacillota;c\_\_Bacilli;o\_\_Lactobacillales;f\_\_Streptococcaceae;g\_\_Streptococcus;s\_\_Streptococcus hyointestinalis | iPHoP-RF | 93.7 | None |
| NC\_002737.2\|provirus\_777501\_820593 | RS\_GCF\_000294495.1 | d\_\_Bacteria;p\_\_Bacillota;c\_\_Bacilli;o\_\_Lactobacillales;f\_\_Streptococcaceae;g\_\_Streptococcus;s\_\_Streptococcus suis | iPHoP-RF | 93.4 | blast;65.20 |
| NC\_002737.2\|provirus\_777501\_820593 | RS\_GCF\_000380085.1 | d\_\_Bacteria;p\_\_Bacillota;c\_\_Bacilli;o\_\_Lactobacillales;f\_\_Streptococcaceae;g\_\_Streptococcus;s\_\_Streptococcus merionis | iPHoP-RF | 93.4 | None |
| NC\_002737.2\|provirus\_777501\_820593 | RS\_GCF\_001302265.1 | d\_\_Bacteria;p\_\_Bacillota;c\_\_Bacilli;o\_\_Lactobacillales;f\_\_Streptococcaceae;g\_\_Streptococcus;s\_\_Streptococcus phocae | iPHoP-RF | 93.4 | None |
| NC\_002737.2\|provirus\_777501\_820593 | RS\_GCF\_016908655.1 | d\_\_Bacteria;p\_\_Bacillota;c\_\_Bacilli;o\_\_Lactobacillales;f\_\_Streptococcaceae;g\_\_Streptococcus;s\_\_Streptococcus saliviloxodontae | iPHoP-RF | 93.4 | None |
| NC\_002737.2\|provirus\_777501\_820593 | RS\_GCF\_019794555.1 | d\_\_Bacteria;p\_\_Bacillota;c\_\_Bacilli;o\_\_Lactobacillales;f\_\_Streptococcaceae;g\_\_Streptococcus;s\_\_Streptococcus suis\_AB | iPHoP-RF | 93.4 | None |
| NC\_002737.2\|provirus\_777501\_820593 | RS\_GCF\_021654455.1 | d\_\_Bacteria;p\_\_Bacillota;c\_\_Bacilli;o\_\_Lactobacillales;f\_\_Streptococcaceae;g\_\_Streptococcus;s\_\_Streptococcus parasuis | iPHoP-RF | 93.4 | None |
| NC\_002737.2\|provirus\_777501\_820593 | GB\_GCA\_900637675.1 | d\_\_Bacteria;p\_\_Bacillota;c\_\_Bacilli;o\_\_Lactobacillales;f\_\_Streptococcaceae;g\_\_Streptococcus;s\_\_Streptococcus equi | iPHoP-RF | 93.1 | blast;70.70 |
| NC\_002737.2\|provirus\_777501\_820593 | RS\_GCF\_000440115.1 | d\_\_Bacteria;p\_\_Bacillota;c\_\_Bacilli;o\_\_Lactobacillales;f\_\_Streptococcaceae;g\_\_Streptococcus;s\_\_Streptococcus suis\_L | iPHoP-RF | 93.1 | None |
| NC\_002737.2\|provirus\_777501\_820593 | RS\_GCF\_003595525.1 | d\_\_Bacteria;p\_\_Bacillota;c\_\_Bacilli;o\_\_Lactobacillales;f\_\_Streptococcaceae;g\_\_Streptococcus;s\_\_Streptococcus respiraculi | iPHoP-RF | 93.1 | None |
| NC\_002737.2\|provirus\_777501\_820593 | RS\_GCF\_015594605.1 | d\_\_Bacteria;p\_\_Bacillota;c\_\_Bacilli;o\_\_Lactobacillales;f\_\_Streptococcaceae;g\_\_Streptococcus;s\_\_Streptococcus sp015594605 | iPHoP-RF | 92.8 | blast;68.90 |
| NC\_002737.2\|provirus\_777501\_820593 | RS\_GCF\_000188055.2 | d\_\_Bacteria;p\_\_Bacillota;c\_\_Bacilli;o\_\_Lactobacillales;f\_\_Streptococcaceae;g\_\_Streptococcus;s\_\_Streptococcus urinalis | iPHoP-RF | 92.1 | None |
| NC\_002737.2\|provirus\_777501\_820593 | RS\_GCF\_012277075.1 | d\_\_Bacteria;p\_\_Bacillota;c\_\_Bacilli;o\_\_Lactobacillales;f\_\_Streptococcaceae;g\_\_Streptococcus;s\_\_Streptococcus alactolyticus | iPHoP-RF | 92.1 | None |
| NC\_002737.2\|provirus\_777501\_820593 | RS\_GCF\_001578805.1 | d\_\_Bacteria;p\_\_Bacillota;c\_\_Bacilli;o\_\_Lactobacillales;f\_\_Streptococcaceae;g\_\_Streptococcus;s\_\_Streptococcus sp001578805 | iPHoP-RF | 91.8 | None |
| NC\_002737.2\|provirus\_777501\_820593 | RS\_GCF\_000785785.1 | d\_\_Bacteria;p\_\_Bacillota;c\_\_Bacilli;o\_\_Lactobacillales;f\_\_Streptococcaceae;g\_\_Streptococcus;s\_\_Streptococcus uberis\_A | iPHoP-RF | 91.4 | None |
| NC\_002737.2\|provirus\_777501\_820593 | RS\_GCF\_022354845.1 | d\_\_Bacteria;p\_\_Bacillota;c\_\_Bacilli;o\_\_Lactobacillales;f\_\_Streptococcaceae;g\_\_Streptococcus;s\_\_Streptococcus suis\_AA | iPHoP-RF | 91.4 | None |
| NC\_002737.2\|provirus\_777501\_820593 | RS\_GCF\_900475595.1 | d\_\_Bacteria;p\_\_Bacillota;c\_\_Bacilli;o\_\_Lactobacillales;f\_\_Streptococcaceae;g\_\_Streptococcus;s\_\_Streptococcus uberis | iPHoP-RF | 91.1 | None |
| NC\_002737.2\|provirus\_777501\_820593 | RS\_GCF\_003674745.1 | d\_\_Bacteria;p\_\_Bacillota;c\_\_Bacilli;o\_\_Lactobacillales;f\_\_Streptococcaceae;g\_\_Streptococcus;s\_\_Streptococcus iniae | iPHoP-RF | 90.8 | None |
| NC\_002737.2\|provirus\_777501\_820593 | RS\_GCF\_000188035.1 | d\_\_Bacteria;p\_\_Bacillota;c\_\_Bacilli;o\_\_Lactobacillales;f\_\_Streptococcaceae;g\_\_Streptococcus;s\_\_Streptococcus pseudoporcinus | iPHoP-RF | 90.5 | None |

##### 2.2.6 AMG Predictions

Prediction of auxiliary metabolic genes in the prophage contigs.

| protein | scaffold | amg\_ko | amg\_ko\_name | pfam | pfam\_name |
| --- | --- | --- | --- | --- | --- |
| NC\_002737.2\|provirus\_777501\_820593\_1 | NC\_002737.2\|provirus\_777501\_820593 | K01710 | E4.2.1.46, rfbB, rffG; dTDP-glucose 4,6-dehydratase [EC:4.2.1.46] | PF16363.5 | GDP-mannose 4,6 dehydratase |

## 2.3 NC\_003450

##### 2.3.1 Host Genome

**GTDB-Tk taxonomy**:

classification

d\_\_Bacteria;p\_\_Actinomycetota;c\_\_Actinomycetes;o\_\_Mycobacteriales;f\_\_Mycobacteriaceae;g\_\_Corynebacterium;s\_\_Corynebacterium
glutamicum

**CheckM2 Summary**:

| total\_contigs | contig\_n50 | completeness | contamination |
| --- | --- | --- | --- |
| 1 | 3309401 | 100 | 0.29 |

**Defense-Finder Systems**:

| sys\_id | type | subtype | sys\_beg | sys\_end | protein\_in\_syst | genes\_count | name\_of\_profiles\_in\_sys |
| --- | --- | --- | --- | --- | --- | --- | --- |
| NC\_003450.3\_RM\_Type\_IIG\_2\_7 | RM | RM\_Type\_IIG\_2 | NC\_003450.3\_730 | NC\_003450.3\_730 | NC\_003450.3\_730 | 1 | RM\_Type\_IIG\_\_Type\_IIG\_FAM\_0.einsi\_trimmed |
| NC\_003450.3\_Uzume\_1 | Uzume | Uzume | NC\_003450.3\_1364 | NC\_003450.3\_1364 | NC\_003450.3\_1364 | 1 | Uzume\_\_UzuA |
| NC\_003450.3\_RM\_Type\_II\_4 | RM | RM\_Type\_II | NC\_003450.3\_1762 | NC\_003450.3\_1764 | NC\_003450.3\_1762,NC\_003450.3\_1763,NC\_003450.3\_1764 | 3 | RM\_Type\_II\_\_Type\_II\_MTases\_FAM\_0,RM\_Type\_II\_\_Type\_II\_REase29,RM\_Type\_II\_\_Type\_II\_REase38 |
| NC\_003450.3\_Wadjet\_I\_3 | Wadjet | Wadjet\_I | NC\_003450.3\_2782 | NC\_003450.3\_2785 | NC\_003450.3\_2782,NC\_003450.3\_2783,NC\_003450.3\_2784,NC\_003450.3\_2785 | 4 | Wadjet\_\_JetA\_I,Wadjet\_\_JetB\_I,Wadjet\_\_JetC\_I,Wadjet\_\_JetD\_I |
| NC\_003450.3\_RM\_Type\_IIG\_6 | RM | RM\_Type\_IIG | NC\_003450.3\_3041 | NC\_003450.3\_3041 | NC\_003450.3\_3041 | 1 | RM\_Type\_IIG\_\_Type\_IIG\_FAM\_2.einsi\_trimmed |

**Genomad and CheckV Summary**:

| seq\_name | length | gene\_count | viral\_genes | checkv\_quality | miuvig\_quality | taxonomy | topology | coordinates |
| --- | --- | --- | --- | --- | --- | --- | --- | --- |
| NC\_003450.3\|provirus\_1777332\_1987821 | 210490 | 195 | 13 | High-quality | High-quality | Viruses;Duplodnaviria;Heunggongvirae;Uroviricota;Caudoviricetes;;Herelleviridae | Provirus | 1777332-1987821 |

**Host Genome Ideogram with Phages**:

In
this circular plot, **“c”** indicates the contig, and the
number that follows (e.g., **c1**) represents the contig
number. If multiple contigs are present in the genome, each will be
shown with a distinct label (e.g., **c1**,
**c2**, etc.).

##### 2.3.2 Prophages

**Select prophage to show:**

###### 2.3.2.1 Phage ID: NC\_003450.3|provirus\_1777332\_1987821

**Phage-Host Genome Ideogram:**

**Genes Annotation (geNomad):**

| gene | length | marker | annotation\_accessions | annotation\_description |
| --- | --- | --- | --- | --- |
| NC\_003450.3\|provirus\_1777332\_1987821\_1641 | 318 | NA | NA | NA |
| NC\_003450.3\|provirus\_1777332\_1987821\_1642 | 126 | NA | NA | NA |
| NC\_003450.3\|provirus\_1777332\_1987821\_1643 | 1410 | NA | NA | NA |
| NC\_003450.3\|provirus\_1777332\_1987821\_1644 | 564 | NA | NA | NA |
| NC\_003450.3\|provirus\_1777332\_1987821\_1645 | 174 | NA | NA | NA |
| NC\_003450.3\|provirus\_1777332\_1987821\_1646 | 564 | NA | NA | NA |
| NC\_003450.3\|provirus\_1777332\_1987821\_1647 | 870 | NA | NA | NA |
| NC\_003450.3\|provirus\_1777332\_1987821\_1648 | 2583 | GENOMAD.055261.VV | NA | NA |
| NC\_003450.3\|provirus\_1777332\_1987821\_1649 | 1116 | GENOMAD.055261.VV | NA | NA |
| NC\_003450.3\|provirus\_1777332\_1987821\_1650 | 1926 | GENOMAD.100223.VV | NA | NA |
| NC\_003450.3\|provirus\_1777332\_1987821\_1651 | 486 | NA | NA | NA |
| NC\_003450.3\|provirus\_1777332\_1987821\_1652 | 192 | NA | NA | NA |
| NC\_003450.3\|provirus\_1777332\_1987821\_1653 | 228 | GENOMAD.189071.VV | NA | NA |
| NC\_003450.3\|provirus\_1777332\_1987821\_1654 | 432 | NA | NA | NA |
| NC\_003450.3\|provirus\_1777332\_1987821\_1655 | 600 | NA | NA | NA |
| NC\_003450.3\|provirus\_1777332\_1987821\_1656 | 1002 | NA | NA | NA |
| NC\_003450.3\|provirus\_1777332\_1987821\_1657 | 156 | NA | NA | NA |
| NC\_003450.3\|provirus\_1777332\_1987821\_1658 | 1110 | NA | NA | NA |
| NC\_003450.3\|provirus\_1777332\_1987821\_1659 | 423 | NA | NA | NA |
| NC\_003450.3\|provirus\_1777332\_1987821\_1660 | 561 | NA | NA | NA |
| NC\_003450.3\|provirus\_1777332\_1987821\_1661 | 867 | NA | NA | NA |
| NC\_003450.3\|provirus\_1777332\_1987821\_1662 | 423 | NA | NA | NA |
| NC\_003450.3\|provirus\_1777332\_1987821\_1663 | 738 | GENOMAD.207798.PC | NA | NA |
| NC\_003450.3\|provirus\_1777332\_1987821\_1664 | 252 | NA | NA | NA |
| NC\_003450.3\|provirus\_1777332\_1987821\_1665 | 228 | NA | NA | NA |
| NC\_003450.3\|provirus\_1777332\_1987821\_1666 | 897 | GENOMAD.222424.VP | PF14243 | NA |
| NC\_003450.3\|provirus\_1777332\_1987821\_1667 | 141 | NA | NA | NA |
| NC\_003450.3\|provirus\_1777332\_1987821\_1668 | 132 | NA | NA | NA |
| NC\_003450.3\|provirus\_1777332\_1987821\_1669 | 438 | NA | NA | NA |
| NC\_003450.3\|provirus\_1777332\_1987821\_1670 | 600 | NA | NA | NA |
| NC\_003450.3\|provirus\_1777332\_1987821\_1671 | 426 | NA | NA | NA |
| NC\_003450.3\|provirus\_1777332\_1987821\_1672 | 534 | NA | NA | NA |
| NC\_003450.3\|provirus\_1777332\_1987821\_1673 | 336 | NA | NA | NA |
| NC\_003450.3\|provirus\_1777332\_1987821\_1674 | 468 | NA | NA | NA |
| NC\_003450.3\|provirus\_1777332\_1987821\_1675 | 615 | NA | NA | NA |
| NC\_003450.3\|provirus\_1777332\_1987821\_1676 | 915 | GENOMAD.138073.VV | NA | NA |
| NC\_003450.3\|provirus\_1777332\_1987821\_1677 | 156 | NA | NA | NA |
| NC\_003450.3\|provirus\_1777332\_1987821\_1678 | 459 | NA | NA | NA |
| NC\_003450.3\|provirus\_1777332\_1987821\_1679 | 684 | GENOMAD.094320.VV | NA | NA |
| NC\_003450.3\|provirus\_1777332\_1987821\_1680 | 288 | NA | NA | NA |
| NC\_003450.3\|provirus\_1777332\_1987821\_1681 | 336 | NA | NA | NA |
| NC\_003450.3\|provirus\_1777332\_1987821\_1682 | 615 | NA | NA | NA |
| NC\_003450.3\|provirus\_1777332\_1987821\_1683 | 996 | NA | NA | NA |
| NC\_003450.3\|provirus\_1777332\_1987821\_1684 | 378 | NA | NA | NA |
| NC\_003450.3\|provirus\_1777332\_1987821\_1685 | 480 | NA | NA | NA |
| NC\_003450.3\|provirus\_1777332\_1987821\_1686 | 729 | NA | NA | NA |
| NC\_003450.3\|provirus\_1777332\_1987821\_1687 | 741 | NA | NA | NA |
| NC\_003450.3\|provirus\_1777332\_1987821\_1688 | 768 | GENOMAD.215962.PC | PF00110 | wnt family |
| NC\_003450.3\|provirus\_1777332\_1987821\_1689 | 459 | NA | NA | NA |
| NC\_003450.3\|provirus\_1777332\_1987821\_1690 | 189 | NA | NA | NA |
| NC\_003450.3\|provirus\_1777332\_1987821\_1691 | 675 | NA | NA | NA |
| NC\_003450.3\|provirus\_1777332\_1987821\_1692 | 420 | NA | NA | NA |
| NC\_003450.3\|provirus\_1777332\_1987821\_1693 | 306 | NA | NA | NA |
| NC\_003450.3\|provirus\_1777332\_1987821\_1694 | 372 | NA | NA | NA |
| NC\_003450.3\|provirus\_1777332\_1987821\_1695 | 210 | NA | NA | NA |
| NC\_003450.3\|provirus\_1777332\_1987821\_1696 | 2205 | NA | NA | NA |
| NC\_003450.3\|provirus\_1777332\_1987821\_1697 | 1698 | NA | NA | NA |
| NC\_003450.3\|provirus\_1777332\_1987821\_1698 | 222 | NA | NA | NA |
| NC\_003450.3\|provirus\_1777332\_1987821\_1699 | 432 | NA | NA | NA |
| NC\_003450.3\|provirus\_1777332\_1987821\_1700 | 897 | NA | NA | NA |
| NC\_003450.3\|provirus\_1777332\_1987821\_1701 | 297 | NA | NA | NA |
| NC\_003450.3\|provirus\_1777332\_1987821\_1702 | 216 | NA | NA | NA |
| NC\_003450.3\|provirus\_1777332\_1987821\_1703 | 1302 | NA | NA | NA |
| NC\_003450.3\|provirus\_1777332\_1987821\_1704 | 1863 | NA | NA | NA |
| NC\_003450.3\|provirus\_1777332\_1987821\_1705 | 783 | NA | NA | NA |
| NC\_003450.3\|provirus\_1777332\_1987821\_1706 | 1776 | NA | NA | NA |
| NC\_003450.3\|provirus\_1777332\_1987821\_1707 | 3792 | GENOMAD.224017.VP | NA | NA |
| NC\_003450.3\|provirus\_1777332\_1987821\_1708 | 450 | NA | NA | NA |
| NC\_003450.3\|provirus\_1777332\_1987821\_1709 | 420 | NA | NA | NA |
| NC\_003450.3\|provirus\_1777332\_1987821\_1710 | 1842 | GENOMAD.007224.VV | PF00176;K14440;COG1061;TIGR04095 | Superfamily II DNA or RNA helicase |
| NC\_003450.3\|provirus\_1777332\_1987821\_1711 | 378 | GENOMAD.222568.VP | NA | NA |
| NC\_003450.3\|provirus\_1777332\_1987821\_1712 | 339 | GENOMAD.168436.VP | PF14359;COG3613;TIGR03646 | Nucleoside 2-deoxyribosyltransferase |
| NC\_003450.3\|provirus\_1777332\_1987821\_1713 | 351 | GENOMAD.224423.VP | PF11774 | Lsr2 |
| NC\_003450.3\|provirus\_1777332\_1987821\_1714 | 621 | GENOMAD.221383.VP | PF07443 | HepA-related protein (HARP) |
| NC\_003450.3\|provirus\_1777332\_1987821\_1715 | 540 | NA | NA | NA |
| NC\_003450.3\|provirus\_1777332\_1987821\_1716 | 531 | NA | NA | NA |
| NC\_003450.3\|provirus\_1777332\_1987821\_1717 | 750 | NA | NA | NA |
| NC\_003450.3\|provirus\_1777332\_1987821\_1718 | 189 | NA | NA | NA |
| NC\_003450.3\|provirus\_1777332\_1987821\_1719 | 336 | NA | NA | NA |
| NC\_003450.3\|provirus\_1777332\_1987821\_1720 | 579 | NA | NA | NA |
| NC\_003450.3\|provirus\_1777332\_1987821\_1721 | 1224 | NA | NA | NA |
| NC\_003450.3\|provirus\_1777332\_1987821\_1722 | 1281 | GENOMAD.226101.VP | PF08800;COG4983 | Uncharacterized protein, contains Primase-polymerase (Primpol) domain |
| NC\_003450.3\|provirus\_1777332\_1987821\_1723 | 576 | NA | NA | NA |
| NC\_003450.3\|provirus\_1777332\_1987821\_1724 | 183 | NA | NA | NA |
| NC\_003450.3\|provirus\_1777332\_1987821\_1725 | 1260 | NA | NA | NA |
| NC\_003450.3\|provirus\_1777332\_1987821\_1726 | 1827 | NA | NA | NA |
| NC\_003450.3\|provirus\_1777332\_1987821\_1727 | 1968 | NA | NA | NA |
| NC\_003450.3\|provirus\_1777332\_1987821\_1728 | 477 | GENOMAD.159467.VP | NA | NA |
| NC\_003450.3\|provirus\_1777332\_1987821\_1729 | 3768 | NA | NA | NA |
| NC\_003450.3\|provirus\_1777332\_1987821\_1730 | 378 | NA | NA | NA |
| NC\_003450.3\|provirus\_1777332\_1987821\_1731 | 381 | NA | NA | NA |
| NC\_003450.3\|provirus\_1777332\_1987821\_1732 | 468 | GENOMAD.218177.VP | PF06892;K22299;TIGR00673;COG5606 | cyanase |
| NC\_003450.3\|provirus\_1777332\_1987821\_1733 | 267 | NA | NA | NA |
| NC\_003450.3\|provirus\_1777332\_1987821\_1734 | 780 | GENOMAD.162261.CP | PF16262;K13355;COG4119;TIGR02150 | Predicted NTP pyrophosphohydrolase, NUDIX family |
| NC\_003450.3\|provirus\_1777332\_1987821\_1735 | 705 | GENOMAD.148006.VV | PF04275;K19797;TIGR01223;COG1102 | phosphomevalonate kinase, animal type |
| NC\_003450.3\|provirus\_1777332\_1987821\_1736 | 228 | NA | NA | NA |
| NC\_003450.3\|provirus\_1777332\_1987821\_1737 | 2160 | NA | NA | NA |
| NC\_003450.3\|provirus\_1777332\_1987821\_1738 | 234 | NA | NA | NA |
| NC\_003450.3\|provirus\_1777332\_1987821\_1739 | 6510 | NA | NA | NA |
| NC\_003450.3\|provirus\_1777332\_1987821\_1740 | 1092 | GENOMAD.136632.VV | PF00145;TIGR00675;K00558;COG0270 | DNA-methyltransferase (dcm) |
| NC\_003450.3\|provirus\_1777332\_1987821\_1741 | 1077 | NA | NA | NA |
| NC\_003450.3\|provirus\_1777332\_1987821\_1742 | 1899 | NA | NA | NA |
| NC\_003450.3\|provirus\_1777332\_1987821\_1743 | 1524 | NA | NA | NA |
| NC\_003450.3\|provirus\_1777332\_1987821\_1744 | 672 | NA | NA | NA |
| NC\_003450.3\|provirus\_1777332\_1987821\_1745 | 1821 | GENOMAD.218864.VV | NA | NA |
| NC\_003450.3\|provirus\_1777332\_1987821\_1746 | 189 | NA | NA | NA |
| NC\_003450.3\|provirus\_1777332\_1987821\_1747 | 354 | NA | NA | NA |
| NC\_003450.3\|provirus\_1777332\_1987821\_1748 | 1218 | GENOMAD.189350.VP | NA | NA |
| NC\_003450.3\|provirus\_1777332\_1987821\_1749 | 1674 | NA | NA | NA |
| NC\_003450.3\|provirus\_1777332\_1987821\_1750 | 1209 | GENOMAD.080614.VV | PF08722 | NA |
| NC\_003450.3\|provirus\_1777332\_1987821\_1751 | 1248 | NA | NA | NA |
| NC\_003450.3\|provirus\_1777332\_1987821\_1752 | 2496 | NA | NA | NA |
| NC\_003450.3\|provirus\_1777332\_1987821\_1753 | 1788 | NA | NA | NA |
| NC\_003450.3\|provirus\_1777332\_1987821\_1754 | 579 | NA | NA | NA |
| NC\_003450.3\|provirus\_1777332\_1987821\_1755 | 1116 | NA | NA | NA |
| NC\_003450.3\|provirus\_1777332\_1987821\_1756 | 849 | NA | NA | NA |
| NC\_003450.3\|provirus\_1777332\_1987821\_1757 | 882 | NA | NA | NA |
| NC\_003450.3\|provirus\_1777332\_1987821\_1758 | 2769 | GENOMAD.161127.PV | PF18555 | MobL relaxases |
| NC\_003450.3\|provirus\_1777332\_1987821\_1759 | 603 | NA | NA | NA |
| NC\_003450.3\|provirus\_1777332\_1987821\_1760 | 1254 | NA | NA | NA |
| NC\_003450.3\|provirus\_1777332\_1987821\_1761 | 699 | NA | NA | NA |
| NC\_003450.3\|provirus\_1777332\_1987821\_1762 | 717 | NA | NA | NA |
| NC\_003450.3\|provirus\_1777332\_1987821\_1763 | 1011 | NA | NA | NA |
| NC\_003450.3\|provirus\_1777332\_1987821\_1764 | 1662 | GENOMAD.214492.VP | PF21822 | Phage tail assembly chaperone protein, TAC |
| NC\_003450.3\|provirus\_1777332\_1987821\_1765 | 1464 | NA | NA | NA |
| NC\_003450.3\|provirus\_1777332\_1987821\_1766 | 360 | GENOMAD.222537.PV | NA | NA |
| NC\_003450.3\|provirus\_1777332\_1987821\_1767 | 1512 | GENOMAD.179757.VP | PF01541;COG2827;TIGR01453 | Predicted endonuclease, GIY-YIG superfamily |
| NC\_003450.3\|provirus\_1777332\_1987821\_1768 | 363 | NA | NA | NA |
| NC\_003450.3\|provirus\_1777332\_1987821\_1769 | 225 | GENOMAD.098687.PP | PF09954 | NA |
| NC\_003450.3\|provirus\_1777332\_1987821\_1770 | 294 | NA | NA | NA |
| NC\_003450.3\|provirus\_1777332\_1987821\_1771 | 510 | NA | NA | NA |
| NC\_003450.3\|provirus\_1777332\_1987821\_1772 | 762 | NA | NA | NA |
| NC\_003450.3\|provirus\_1777332\_1987821\_1773 | 552 | NA | NA | NA |
| NC\_003450.3\|provirus\_1777332\_1987821\_1774 | 933 | GENOMAD.225221.VP | PF12686 | NA |
| NC\_003450.3\|provirus\_1777332\_1987821\_1775 | 4467 | NA | NA | NA |
| NC\_003450.3\|provirus\_1777332\_1987821\_1776 | 582 | NA | NA | NA |
| NC\_003450.3\|provirus\_1777332\_1987821\_1777 | 936 | NA | NA | NA |
| NC\_003450.3\|provirus\_1777332\_1987821\_1778 | 138 | NA | NA | NA |
| NC\_003450.3\|provirus\_1777332\_1987821\_1779 | 378 | NA | NA | NA |
| NC\_003450.3\|provirus\_1777332\_1987821\_1780 | 1824 | NA | NA | NA |
| NC\_003450.3\|provirus\_1777332\_1987821\_1781 | 204 | NA | NA | NA |
| NC\_003450.3\|provirus\_1777332\_1987821\_1782 | 462 | GENOMAD.223777.PV | PF02556;TIGR00809;COG1952;K03071 | protein-export chaperone SecB |
| NC\_003450.3\|provirus\_1777332\_1987821\_1783 | 360 | NA | NA | NA |
| NC\_003450.3\|provirus\_1777332\_1987821\_1784 | 408 | NA | NA | NA |
| NC\_003450.3\|provirus\_1777332\_1987821\_1785 | 732 | NA | NA | NA |
| NC\_003450.3\|provirus\_1777332\_1987821\_1786 | 945 | NA | NA | NA |
| NC\_003450.3\|provirus\_1777332\_1987821\_1787 | 561 | NA | NA | NA |
| NC\_003450.3\|provirus\_1777332\_1987821\_1788 | 213 | NA | NA | NA |
| NC\_003450.3\|provirus\_1777332\_1987821\_1789 | 537 | NA | NA | NA |
| NC\_003450.3\|provirus\_1777332\_1987821\_1790 | 537 | NA | NA | NA |
| NC\_003450.3\|provirus\_1777332\_1987821\_1791 | 945 | NA | NA | NA |
| NC\_003450.3\|provirus\_1777332\_1987821\_1792 | 591 | NA | NA | NA |
| NC\_003450.3\|provirus\_1777332\_1987821\_1793 | 447 | NA | NA | NA |
| NC\_003450.3\|provirus\_1777332\_1987821\_1794 | 702 | NA | NA | NA |
| NC\_003450.3\|provirus\_1777332\_1987821\_1795 | 297 | NA | NA | NA |
| NC\_003450.3\|provirus\_1777332\_1987821\_1796 | 219 | NA | NA | NA |
| NC\_003450.3\|provirus\_1777332\_1987821\_1797 | 135 | NA | NA | NA |
| NC\_003450.3\|provirus\_1777332\_1987821\_1798 | 777 | NA | NA | NA |
| NC\_003450.3\|provirus\_1777332\_1987821\_1799 | 831 | NA | NA | NA |
| NC\_003450.3\|provirus\_1777332\_1987821\_1800 | 276 | NA | NA | NA |
| NC\_003450.3\|provirus\_1777332\_1987821\_1801 | 381 | NA | NA | NA |
| NC\_003450.3\|provirus\_1777332\_1987821\_1802 | 432 | NA | NA | NA |
| NC\_003450.3\|provirus\_1777332\_1987821\_1803 | 1584 | NA | NA | NA |
| NC\_003450.3\|provirus\_1777332\_1987821\_1804 | 2433 | NA | NA | NA |
| NC\_003450.3\|provirus\_1777332\_1987821\_1805 | 870 | GENOMAD.207947.PV | NA | NA |
| NC\_003450.3\|provirus\_1777332\_1987821\_1806 | 2280 | NA | NA | NA |
| NC\_003450.3\|provirus\_1777332\_1987821\_1807 | 2088 | NA | NA | NA |
| NC\_003450.3\|provirus\_1777332\_1987821\_1808 | 894 | NA | NA | NA |
| NC\_003450.3\|provirus\_1777332\_1987821\_1809 | 435 | NA | NA | NA |
| NC\_003450.3\|provirus\_1777332\_1987821\_1810 | 747 | NA | NA | NA |
| NC\_003450.3\|provirus\_1777332\_1987821\_1811 | 1857 | NA | NA | NA |
| NC\_003450.3\|provirus\_1777332\_1987821\_1812 | 294 | NA | NA | NA |
| NC\_003450.3\|provirus\_1777332\_1987821\_1813 | 1221 | NA | NA | NA |
| NC\_003450.3\|provirus\_1777332\_1987821\_1814 | 1179 | NA | NA | NA |
| NC\_003450.3\|provirus\_1777332\_1987821\_1815 | 333 | NA | NA | NA |
| NC\_003450.3\|provirus\_1777332\_1987821\_1816 | 639 | NA | NA | NA |
| NC\_003450.3\|provirus\_1777332\_1987821\_1817 | 120 | NA | NA | NA |
| NC\_003450.3\|provirus\_1777332\_1987821\_1818 | 144 | NA | NA | NA |
| NC\_003450.3\|provirus\_1777332\_1987821\_1819 | 567 | NA | NA | NA |
| NC\_003450.3\|provirus\_1777332\_1987821\_1820 | 1455 | NA | NA | NA |
| NC\_003450.3\|provirus\_1777332\_1987821\_1821 | 462 | NA | NA | NA |
| NC\_003450.3\|provirus\_1777332\_1987821\_1822 | 1143 | NA | NA | NA |
| NC\_003450.3\|provirus\_1777332\_1987821\_1823 | 150 | NA | NA | NA |
| NC\_003450.3\|provirus\_1777332\_1987821\_1824 | 1422 | NA | NA | NA |
| NC\_003450.3\|provirus\_1777332\_1987821\_1825 | 594 | NA | NA | NA |
| NC\_003450.3\|provirus\_1777332\_1987821\_1826 | 399 | NA | NA | NA |
| NC\_003450.3\|provirus\_1777332\_1987821\_1827 | 240 | NA | NA | NA |
| NC\_003450.3\|provirus\_1777332\_1987821\_1828 | 627 | NA | NA | NA |
| NC\_003450.3\|provirus\_1777332\_1987821\_1829 | 582 | NA | NA | NA |
| NC\_003450.3\|provirus\_1777332\_1987821\_1830 | 465 | NA | NA | NA |
| NC\_003450.3\|provirus\_1777332\_1987821\_1831 | 510 | NA | NA | NA |
| NC\_003450.3\|provirus\_1777332\_1987821\_1832 | 561 | NA | NA | NA |
| NC\_003450.3\|provirus\_1777332\_1987821\_1833 | 336 | NA | NA | NA |
| NC\_003450.3\|provirus\_1777332\_1987821\_1834 | 483 | NA | NA | NA |
| NC\_003450.3\|provirus\_1777332\_1987821\_1835 | 573 | NA | NA | NA |
| NC\_003450.3\|provirus\_1777332\_1987821\_1836 | 915 | NA | NA | NA |
| NC\_003450.3\|provirus\_1777332\_1987821\_1837 | 696 | GENOMAD.178028.VP | NA | NA |
| NC\_003450.3\|provirus\_1777332\_1987821\_1838 | 369 | NA | NA | NA |
| NC\_003450.3\|provirus\_1777332\_1987821\_1839 | 645 | NA | NA | NA |
| NC\_003450.3\|provirus\_1777332\_1987821\_1840 | 366 | NA | NA | NA |
| NC\_003450.3\|provirus\_1777332\_1987821\_1841 | 276 | NA | NA | NA |
| NC\_003450.3\|provirus\_1777332\_1987821\_1842 | 267 | NA | NA | NA |
| NC\_003450.3\|provirus\_1777332\_1987821\_1843 | 237 | NA | NA | NA |
| NC\_003450.3\|provirus\_1777332\_1987821\_1844 | 237 | GENOMAD.082277.VV | NA | NA |
| NC\_003450.3\|provirus\_1777332\_1987821\_1845 | 276 | NA | NA | NA |
| NC\_003450.3\|provirus\_1777332\_1987821\_1846 | 1086 | NA | NA | NA |
| NC\_003450.3\|provirus\_1777332\_1987821\_1847 | 318 | NA | NA | NA |

##### 2.3.3 Prophage-Host Prediction

Prediction of potential hosts for the prophage contigs.

| virus | host\_genome | host\_taxonomy | main\_method | confidence\_score | additional\_methods |
| --- | --- | --- | --- | --- | --- |
| NC\_003450.3\|provirus\_1777332\_1987821 | RS\_GCF\_000011325.1 | d\_\_Bacteria;p\_\_Actinomycetota;c\_\_Actinomycetia;o\_\_Mycobacteriales;f\_\_Mycobacteriaceae;g\_\_Corynebacterium;s\_\_Corynebacterium glutamicum | blast | 96.8 | iPHoP-RF;52.80 |
| NC\_003450.3\|provirus\_1777332\_1987821 | RS\_GCF\_002355155.1 | d\_\_Bacteria;p\_\_Actinomycetota;c\_\_Actinomycetia;o\_\_Mycobacteriales;f\_\_Mycobacteriaceae;g\_\_Corynebacterium;s\_\_Corynebacterium suranareeae | blast | 96.3 | None |
| NC\_003450.3\|provirus\_1777332\_1987821 | RS\_GCF\_001643015.1 | d\_\_Bacteria;p\_\_Actinomycetota;c\_\_Actinomycetia;o\_\_Mycobacteriales;f\_\_Mycobacteriaceae;g\_\_Corynebacterium;s\_\_Corynebacterium crudilactis | blast | 95.9 | None |

## 2.4 NC\_008261

##### 2.4.1 Host Genome

**GTDB-Tk taxonomy**:

classification

d\_\_Bacteria;p\_\_Bacillota\_A;c\_\_Clostridia;o\_\_Clostridiales;f\_\_Clostridiaceae;g\_\_Sarcina;s\_\_Sarcina
perfringens

**CheckM2 Summary**:

| total\_contigs | contig\_n50 | completeness | contamination |
| --- | --- | --- | --- |
| 1 | 3256683 | 100 | 0.14 |

**Defense-Finder Systems**:

| sys\_id | type | subtype | sys\_beg | sys\_end | protein\_in\_syst | genes\_count | name\_of\_profiles\_in\_sys |
| --- | --- | --- | --- | --- | --- | --- | --- |
| NC\_008261.1\_RM\_Type\_II\_5 | RM | RM\_Type\_II | NC\_008261.1\_127 | NC\_008261.1\_128 | NC\_008261.1\_127,NC\_008261.1\_128 | 2 | RM\_Type\_II\_\_Type\_II\_MTases\_FAM\_0,RM\_Type\_II\_\_Type\_II\_REase18 |
| NC\_008261.1\_RM\_Type\_III\_7 | RM | RM\_Type\_III | NC\_008261.1\_340 | NC\_008261.1\_341 | NC\_008261.1\_340,NC\_008261.1\_341 | 2 | RM\_Type\_III\_\_Type\_III\_MTases\_FAM\_0,RM\_Type\_III\_\_Type\_III\_REases\_FAM\_0.einsi\_trimmed |
| NC\_008261.1\_RM\_Type\_IV\_8 | RM | RM\_Type\_IV | NC\_008261.1\_962 | NC\_008261.1\_963 | NC\_008261.1\_962,NC\_008261.1\_963 | 2 | RM\_Type\_IV\_\_FAM\_1,RM\_Type\_IV\_\_FAM\_2 |
| NC\_008261.1\_DRT\_2\_1 | DRT | DRT\_2 | NC\_008261.1\_983 | NC\_008261.1\_983 | NC\_008261.1\_983 | 1 | DRT2\_\_DRT2 |
| NC\_008261.1\_RM\_Type\_II\_6 | RM | RM\_Type\_II | NC\_008261.1\_985 | NC\_008261.1\_987 | NC\_008261.1\_985,NC\_008261.1\_986,NC\_008261.1\_987 | 3 | RM\_Type\_II\_\_Type\_II\_MTases\_FAM\_0,RM\_Type\_II\_\_Type\_II\_MTases\_FAM\_22,RM\_Type\_II\_\_Type\_II\_REase34 |
| NC\_008261.1\_PD-Lambda-5\_2 | PD-Lambda-5 | PD-Lambda-5 | NC\_008261.1\_1563 | NC\_008261.1\_1564 | NC\_008261.1\_1563,NC\_008261.1\_1564 | 2 | PD-Lambda-5\_\_PD-Lambda-5\_A,PD-Lambda-5\_\_PD-Lambda-5\_B |
| NC\_008261.1\_RM\_Type\_I\_4 | RM | RM\_Type\_I | NC\_008261.1\_2512 | NC\_008261.1\_2515 | NC\_008261.1\_2512,NC\_008261.1\_2514,NC\_008261.1\_2515 | 3 | RM\_\_Type\_I\_MTases\_FAM\_0,RM\_\_Type\_I\_REases\_FAM\_0.einsi\_trimmed,RM\_\_Type\_I\_S\_51 |

**Genomad and CheckV Summary**:

| seq\_name | length | gene\_count | viral\_genes | checkv\_quality | miuvig\_quality | taxonomy | topology | coordinates |
| --- | --- | --- | --- | --- | --- | --- | --- | --- |
| NC\_008261.1\|provirus\_1084498\_1127691 | 43194 | 55 | 25 | High-quality | High-quality | Viruses;Duplodnaviria;Heunggongvirae;Uroviricota;Caudoviricetes;; | Provirus | 1084498-1127691 |
| NC\_008261.1\|provirus\_1784095\_1821197 | 37103 | 44 | 21 | Medium-quality | Genome-fragment | Viruses;Duplodnaviria;Heunggongvirae;Uroviricota;Caudoviricetes;; | Provirus | 1784095-1821197 |

**Host Genome Ideogram with Phages**:

In
this circular plot, **“c”** indicates the contig, and the
number that follows (e.g., **c1**) represents the contig
number. If multiple contigs are present in the genome, each will be
shown with a distinct label (e.g., **c1**,
**c2**, etc.).

##### 2.4.2 Prophages

**Select prophage to show:**

###### 2.4.2.1 Phage ID: NC\_008261.1|provirus\_1084498\_1127691

**Phage-Host Genome Ideogram:**

**Genes Annotation (geNomad):**

| gene | length | marker | annotation\_accessions | annotation\_description |
| --- | --- | --- | --- | --- |
| NC\_008261.1\|provirus\_1084498\_1127691\_890 | 489 | NA | NA | NA |
| NC\_008261.1\|provirus\_1084498\_1127691\_891 | 1761 | NA | NA | NA |
| NC\_008261.1\|provirus\_1084498\_1127691\_892 | 459 | NA | NA | NA |
| NC\_008261.1\|provirus\_1084498\_1127691\_893 | 723 | NA | NA | NA |
| NC\_008261.1\|provirus\_1084498\_1127691\_894 | 219 | NA | NA | NA |
| NC\_008261.1\|provirus\_1084498\_1127691\_895 | 1065 | NA | NA | NA |
| NC\_008261.1\|provirus\_1084498\_1127691\_896 | 465 | GENOMAD.123021.VV | PF06114;COG2856 | IrrE N-terminal-like domain |
| NC\_008261.1\|provirus\_1084498\_1127691\_897 | 504 | NA | NA | NA |
| NC\_008261.1\|provirus\_1084498\_1127691\_898 | 237 | GENOMAD.069056.VV | PF06892;TIGR00673;COG1974;K07727 | cyanase |
| NC\_008261.1\|provirus\_1084498\_1127691\_899 | 732 | GENOMAD.119058.VP | PF10552 | ORF6C domain |
| NC\_008261.1\|provirus\_1084498\_1127691\_900 | 255 | NA | NA | NA |
| NC\_008261.1\|provirus\_1084498\_1127691\_901 | 672 | NA | NA | NA |
| NC\_008261.1\|provirus\_1084498\_1127691\_902 | 303 | GENOMAD.061044.VV | PF13411;COG4220;TIGR02054 | Phage DNA packaging protein, Nu1 subunit of terminase |
| NC\_008261.1\|provirus\_1084498\_1127691\_903 | 180 | GENOMAD.164182.VV | NA | NA |
| NC\_008261.1\|provirus\_1084498\_1127691\_904 | 183 | GENOMAD.117872.VV | NA | NA |
| NC\_008261.1\|provirus\_1084498\_1127691\_905 | 108 | NA | NA | NA |
| NC\_008261.1\|provirus\_1084498\_1127691\_906 | 186 | GENOMAD.172161.VC | NA | NA |
| NC\_008261.1\|provirus\_1084498\_1127691\_907 | 912 | GENOMAD.077205.VV | NA | NA |
| NC\_008261.1\|provirus\_1084498\_1127691\_908 | 870 | GENOMAD.100145.VP | PF03837;COG3723;K07455;TIGR00616 | Recombinational DNA repair protein RecT |
| NC\_008261.1\|provirus\_1084498\_1127691\_909 | 714 | GENOMAD.178290.VP | PF04492;TIGR01714;COG2188 | Bacteriophage replication protein O |
| NC\_008261.1\|provirus\_1084498\_1127691\_910 | 174 | NA | NA | NA |
| NC\_008261.1\|provirus\_1084498\_1127691\_911 | 387 | GENOMAD.177634.VP | PF05866;COG4570 | Endodeoxyribonuclease RusA |
| NC\_008261.1\|provirus\_1084498\_1127691\_912 | 648 | NA | NA | NA |
| NC\_008261.1\|provirus\_1084498\_1127691\_913 | 474 | GENOMAD.062141.VV | PF05263;TIGR01636 | phage transcriptional activator, RinA family |
| NC\_008261.1\|provirus\_1084498\_1127691\_914 | 1176 | NA | NA | NA |
| NC\_008261.1\|provirus\_1084498\_1127691\_915 | 468 | GENOMAD.026460.VV | PF03592;COG3728;K07474 | Phage terminase, small subunit |
| NC\_008261.1\|provirus\_1084498\_1127691\_916 | 1365 | GENOMAD.017640.VV | NA | NA |
| NC\_008261.1\|provirus\_1084498\_1127691\_917 | 768 | NA | NA | NA |
| NC\_008261.1\|provirus\_1084498\_1127691\_918 | 1521 | GENOMAD.016130.VV | PF05133;TIGR01542 | phage portal protein, putative, A118 family |
| NC\_008261.1\|provirus\_1084498\_1127691\_919 | 1638 | GENOMAD.015690.VV | PF06152;TIGR01641 | Phage minor capsid protein 2 |
| NC\_008261.1\|provirus\_1084498\_1127691\_920 | 225 | GENOMAD.093339.VV | NA | NA |
| NC\_008261.1\|provirus\_1084498\_1127691\_921 | 618 | NA | NA | NA |
| NC\_008261.1\|provirus\_1084498\_1127691\_922 | 606 | GENOMAD.041104.VV | NA | NA |
| NC\_008261.1\|provirus\_1084498\_1127691\_923 | 906 | GENOMAD.075076.VV | NA | NA |
| NC\_008261.1\|provirus\_1084498\_1127691\_924 | 270 | GENOMAD.050526.VV | NA | NA |
| NC\_008261.1\|provirus\_1084498\_1127691\_925 | 363 | GENOMAD.020431.VV | PF11436 | Putative DnaT-like ssDNA binding protein |
| NC\_008261.1\|provirus\_1084498\_1127691\_926 | 327 | GENOMAD.024076.VV | PF10665 | Minor capsid protein |
| NC\_008261.1\|provirus\_1084498\_1127691\_927 | 384 | GENOMAD.019595.VV | PF11114 | Minor capsid protein |
| NC\_008261.1\|provirus\_1084498\_1127691\_928 | 387 | GENOMAD.028696.VV | NA | NA |
| NC\_008261.1\|provirus\_1084498\_1127691\_929 | 459 | GENOMAD.019307.VV | PF16461;COG5437 | Predicted secreted protein |
| NC\_008261.1\|provirus\_1084498\_1127691\_930 | 369 | GENOMAD.052890.VV | NA | NA |
| NC\_008261.1\|provirus\_1084498\_1127691\_931 | 363 | GENOMAD.015650.VV | NA | NA |
| NC\_008261.1\|provirus\_1084498\_1127691\_932 | 3213 | GENOMAD.171590.VP | NA | NA |
| NC\_008261.1\|provirus\_1084498\_1127691\_933 | 354 | GENOMAD.012935.VV | PF20458 | NA |
| NC\_008261.1\|provirus\_1084498\_1127691\_934 | 267 | GENOMAD.163907.VV | NA | NA |
| NC\_008261.1\|provirus\_1084498\_1127691\_935 | 381 | GENOMAD.016019.VV | PF07761 | NA |
| NC\_008261.1\|provirus\_1084498\_1127691\_936 | 5109 | GENOMAD.141397.VP | PF06605;PF00149;TIGR01665;K01517;COG4926 | phage minor structural protein, N-terminal region |
| NC\_008261.1\|provirus\_1084498\_1127691\_937 | 249 | GENOMAD.113645.VV | NA | NA |
| NC\_008261.1\|provirus\_1084498\_1127691\_938 | 417 | NA | NA | NA |
| NC\_008261.1\|provirus\_1084498\_1127691\_939 | 417 | GENOMAD.224084.VP | NA | NA |
| NC\_008261.1\|provirus\_1084498\_1127691\_940 | 1029 | GENOMAD.169358.VC | PF06725;PF18348;COG3584;K11060;TIGR04211 | 3D (Asp-Asp-Asp) domain |
| NC\_008261.1\|provirus\_1084498\_1127691\_941 | 663 | NA | NA | NA |
| NC\_008261.1\|provirus\_1084498\_1127691\_942 | 1089 | GENOMAD.129207.VV | NA | NA |
| NC\_008261.1\|provirus\_1084498\_1127691\_943 | 342 | NA | NA | NA |
| NC\_008261.1\|provirus\_1084498\_1127691\_944 | 726 | GENOMAD.129093.VV | NA | NA |

###### 2.4.2.2 Phage ID: NC\_008261.1|provirus\_1784095\_1821197

**Phage-Host Genome Ideogram:**

**Genes Annotation (geNomad):**

| gene | length | marker | annotation\_accessions | annotation\_description |
| --- | --- | --- | --- | --- |
| NC\_008261.1\|provirus\_1784095\_1821197\_1520 | 486 | NA | NA | NA |
| NC\_008261.1\|provirus\_1784095\_1821197\_1521 | 303 | GENOMAD.099786.VV | NA | NA |
| NC\_008261.1\|provirus\_1784095\_1821197\_1522 | 1029 | GENOMAD.169358.VC | PF06725;PF18348;COG3584;K11060;TIGR04211 | 3D (Asp-Asp-Asp) domain |
| NC\_008261.1\|provirus\_1784095\_1821197\_1523 | 483 | GENOMAD.222318.VP | NA | NA |
| NC\_008261.1\|provirus\_1784095\_1821197\_1524 | 234 | GENOMAD.080405.VV | PF10779 | Haemolysin XhlA |
| NC\_008261.1\|provirus\_1784095\_1821197\_1525 | 186 | NA | NA | NA |
| NC\_008261.1\|provirus\_1784095\_1821197\_1526 | 2499 | GENOMAD.033703.VV | NA | NA |
| NC\_008261.1\|provirus\_1784095\_1821197\_1527 | 1881 | GENOMAD.082024.VV | PF10651;PF15425 | BppU N-terminal domain |
| NC\_008261.1\|provirus\_1784095\_1821197\_1528 | 3060 | GENOMAD.004842.VV | PF06605;PF18994;COG4926;TIGR01665 | Phage-related protein |
| NC\_008261.1\|provirus\_1784095\_1821197\_1529 | 711 | GENOMAD.004006.VV | PF20195;COG4722;TIGR01633 | Phage-related protein |
| NC\_008261.1\|provirus\_1784095\_1821197\_1530 | 3255 | GENOMAD.116638.VP | NA | NA |
| NC\_008261.1\|provirus\_1784095\_1821197\_1531 | 354 | GENOMAD.080513.VV | NA | NA |
| NC\_008261.1\|provirus\_1784095\_1821197\_1532 | 318 | GENOMAD.080540.VV | NA | NA |
| NC\_008261.1\|provirus\_1784095\_1821197\_1533 | 603 | GENOMAD.013991.VV | PF04630;TIGR01603 | phage major tail protein, phi13 family |
| NC\_008261.1\|provirus\_1784095\_1821197\_1534 | 348 | GENOMAD.056191.VV | PF05657 | NA |
| NC\_008261.1\|provirus\_1784095\_1821197\_1535 | 438 | GENOMAD.072939.VV | TIGR01725;COG5005 | phage protein, HK97 gp10 family |
| NC\_008261.1\|provirus\_1784095\_1821197\_1536 | 330 | GENOMAD.078312.VV | PF05521;TIGR01563;COG5614 | phage head-tail adaptor, putative, SPP1 family |
| NC\_008261.1\|provirus\_1784095\_1821197\_1537 | 279 | GENOMAD.034954.VV | PF05135;TIGR01560 | Phage gp6-like head-tail connector protein |
| NC\_008261.1\|provirus\_1784095\_1821197\_1538 | 1185 | GENOMAD.014413.VV | PF05065;TIGR01554;COG4653 | phage major capsid protein, HK97 family |
| NC\_008261.1\|provirus\_1784095\_1821197\_1539 | 606 | GENOMAD.126260.VV | PF04586;COG3740;K06904;TIGR01543 | Phage head maturation protease |
| NC\_008261.1\|provirus\_1784095\_1821197\_1540 | 1248 | GENOMAD.126260.VV | PF04586;COG3740;K06904;TIGR01543 | Phage head maturation protease |
| NC\_008261.1\|provirus\_1784095\_1821197\_1541 | 1740 | GENOMAD.192500.VP | PF18352;COG4626 | Phage terminase-like protein, large subunit, contains N-terminal HTH domain |
| NC\_008261.1\|provirus\_1784095\_1821197\_1542 | 513 | GENOMAD.136127.VP | PF05119;COG3747;TIGR01558 | Phage terminase, small subunit |
| NC\_008261.1\|provirus\_1784095\_1821197\_1543 | 822 | NA | NA | NA |
| NC\_008261.1\|provirus\_1784095\_1821197\_1544 | 549 | GENOMAD.027124.VV | PF07104 | NA |
| NC\_008261.1\|provirus\_1784095\_1821197\_1545 | 354 | NA | NA | NA |
| NC\_008261.1\|provirus\_1784095\_1821197\_1546 | 207 | GENOMAD.117627.VV | NA | NA |
| NC\_008261.1\|provirus\_1784095\_1821197\_1547 | 462 | GENOMAD.062141.VV | PF05263;TIGR01636 | phage transcriptional activator, RinA family |
| NC\_008261.1\|provirus\_1784095\_1821197\_1548 | 390 | NA | NA | NA |
| NC\_008261.1\|provirus\_1784095\_1821197\_1549 | 147 | NA | NA | NA |
| NC\_008261.1\|provirus\_1784095\_1821197\_1550 | 1284 | GENOMAD.068048.VV | PF20307;TIGR03600;K02314;COG0305 | phage replicative helicase, DnaB family, HK022 subfamily |
| NC\_008261.1\|provirus\_1784095\_1821197\_1551 | 741 | GENOMAD.096239.VV | NA | NA |
| NC\_008261.1\|provirus\_1784095\_1821197\_1552 | 159 | NA | NA | NA |
| NC\_008261.1\|provirus\_1784095\_1821197\_1553 | 180 | GENOMAD.148814.VV | NA | NA |
| NC\_008261.1\|provirus\_1784095\_1821197\_1554 | 231 | NA | NA | NA |
| NC\_008261.1\|provirus\_1784095\_1821197\_1555 | 498 | GENOMAD.142555.VV | PF06892;TIGR02612;K22299;COG3620 | Phage regulatory protein CII (CP76) |
| NC\_008261.1\|provirus\_1784095\_1821197\_1556 | 189 | NA | NA | NA |
| NC\_008261.1\|provirus\_1784095\_1821197\_1557 | 345 | NA | NA | NA |
| NC\_008261.1\|provirus\_1784095\_1821197\_1558 | 1590 | GENOMAD.219971.CV | NA | NA |
| NC\_008261.1\|provirus\_1784095\_1821197\_1559 | 855 | NA | NA | NA |
| NC\_008261.1\|provirus\_1784095\_1821197\_1560 | 498 | GENOMAD.123021.VV | PF06114;COG2856 | IrrE N-terminal-like domain |
| NC\_008261.1\|provirus\_1784095\_1821197\_1561 | 1509 | GENOMAD.169508.VP | PF04708;PF13262;COG1961;K14060 | Site-specific DNA recombinase related to the DNA invertase Pin |
| NC\_008261.1\|provirus\_1784095\_1821197\_1562 | 339 | NA | NA | NA |
| NC\_008261.1\|provirus\_1784095\_1821197\_1563 | 378 | NA | NA | NA |

##### 2.4.3 vOTUs

When more than 1 phage is detected in the host genome, we perform a
clustering step using the tool dRep.

A threshold of 0.95 was applied to the ANI similarity index to define
clusters of virus operational taxonomic units (vOTUs).

**Final cluster designations**

| genomad\_summary$seq\_name | genome | secondary\_cluster | threshold | cluster\_method | comparison\_algorithm | primary\_cluster |
| --- | --- | --- | --- | --- | --- | --- |
| NC\_008261.1\|provirus\_1084498\_1127691 | sequence\_000000.fasta.fasta | 1\_0 | 0.05 | average | ANImf | 1 |
| NC\_008261.1\|provirus\_1784095\_1821197 | sequence\_000001.fasta.fasta | 2\_0 | 0.05 | average | ANImf | 2 |

**Primary clustering plot**

##### 2.4.4 Prophage-Host Prediction

Prediction of potential hosts for the prophage contigs.

| virus | host\_genome | host\_taxonomy | main\_method | confidence\_score | additional\_methods |
| --- | --- | --- | --- | --- | --- |
| NC\_008261.1\|provirus\_1084498\_1127691 | RS\_GCF\_000013285.1 | d\_\_Bacteria;p\_\_Bacillota\_A;c\_\_Clostridia;o\_\_Clostridiales;f\_\_Clostridiaceae;g\_\_Sarcina;s\_\_Sarcina perfringens | blast | 98.2 | iPHoP-RF;75.00 |
| NC\_008261.1\|provirus\_1784095\_1821197 | RS\_GCF\_000013285.1 | d\_\_Bacteria;p\_\_Bacillota\_A;c\_\_Clostridia;o\_\_Clostridiales;f\_\_Clostridiaceae;g\_\_Sarcina;s\_\_Sarcina perfringens | blast | 96.9 | CRISPR;93.40 iPHoP-RF;85.40 |

## 2.5 NC\_009012

##### 2.5.1 Host Genome

**GTDB-Tk taxonomy**:

classification

d\_\_Bacteria;p\_\_Bacillota\_A;c\_\_Clostridia;o\_\_Acetivibrionales;f\_\_Acetivibrionaceae;g\_\_Hungateiclostridium;s\_\_Hungateiclostridium
thermocellum

**CheckM2 Summary**:

| total\_contigs | contig\_n50 | completeness | contamination |
| --- | --- | --- | --- |
| 1 | 3843301 | 100 | 1.09 |

**Defense-Finder Systems**:

| sys\_id | type | subtype | sys\_beg | sys\_end | protein\_in\_syst | genes\_count | name\_of\_profiles\_in\_sys |
| --- | --- | --- | --- | --- | --- | --- | --- |
| NC\_009012.1\_RM\_Type\_III\_8 | RM | RM\_Type\_III | NC\_009012.1\_532 | NC\_009012.1\_533 | NC\_009012.1\_532,NC\_009012.1\_533 | 2 | RM\_Type\_III\_\_Type\_III\_MTases\_FAM\_0,RM\_Type\_III\_\_Type\_III\_REases\_FAM\_0.einsi\_trimmed |
| NC\_009012.1\_SEFIR\_3 | SEFIR | SEFIR | NC\_009012.1\_1184 | NC\_009012.1\_1184 | NC\_009012.1\_1184 | 1 | SEFIR\_\_bSEFIR |
| NC\_009012.1\_RloC\_2 | RloC | RloC | NC\_009012.1\_1211 | NC\_009012.1\_1211 | NC\_009012.1\_1211 | 1 | RloC\_\_RloC |
| NC\_009012.1\_RM\_Type\_II\_4 | RM | RM\_Type\_II | NC\_009012.1\_1577 | NC\_009012.1\_1579 | NC\_009012.1\_1577,NC\_009012.1\_1578,NC\_009012.1\_1579 | 3 | RM\_Type\_II\_\_Type\_II\_MTases\_FAM\_38,RM\_Type\_II\_\_Type\_II\_MTases\_FAM\_4,RM\_Type\_II\_\_Type\_II\_REase32 |
| NC\_009012.1\_AbiU\_1 | AbiU | AbiU | NC\_009012.1\_1698 | NC\_009012.1\_1698 | NC\_009012.1\_1698 | 1 | AbiU\_\_AbiU |
| NC\_009012.1\_RM\_Type\_II\_5 | RM | RM\_Type\_II | NC\_009012.1\_1826 | NC\_009012.1\_1827 | NC\_009012.1\_1826,NC\_009012.1\_1827 | 2 | RM\_Type\_II\_\_Type\_II\_MTases\_FAM\_16,RM\_Type\_II\_\_Type\_II\_REase01 |
| NC\_009012.1\_CAS\_Class1-Subtype-III-D\_11 | Cas | CAS\_Class1-Subtype-III-D | NC\_009012.1\_2140 | NC\_009012.1\_2149 | NC\_009012.1\_2140,NC\_009012.1\_2141,NC\_009012.1\_2142,NC\_009012.1\_2143,NC\_009012.1\_2144,NC\_009012.1\_2145,NC\_009012.1\_2146,NC\_009012.1\_2147,NC\_009012.1\_2149 | 9 | cas10\_III-D\_3,csm2gr11\_III-D\_6,csm2gr11\_III-D\_7,csm3gr7\_III-A\_III-D\_2,csm3gr7\_III\_1,csm3gr7\_III\_1,csm3gr7\_III\_IV,csx10gr5\_III-D\_2,csx1\_III\_9 |
| NC\_009012.1\_CAS\_Class1-Subtype-I-B\_9 | Cas | CAS\_Class1-Subtype-I-B | NC\_009012.1\_2404 | NC\_009012.1\_2411 | NC\_009012.1\_2404,NC\_009012.1\_2405,NC\_009012.1\_2406,NC\_009012.1\_2407,NC\_009012.1\_2408,NC\_009012.1\_2409,NC\_009012.1\_2410,NC\_009012.1\_2411 | 8 | cas1\_I\_II\_III\_IV\_V\_VI\_7,cas2\_I\_II\_III\_IV\_V\_VI\_3,cas3\_I\_5,cas4\_I\_II\_III\_IV\_V\_VI\_6,cas5\_I-B\_1,cas6\_I\_II\_III\_IV\_V\_VI\_14,cas7\_I-B\_6,cas8b1\_I-B\_12 |
| NC\_009012.1\_RM\_Type\_II\_6 | RM | RM\_Type\_II | NC\_009012.1\_2429 | NC\_009012.1\_2430 | NC\_009012.1\_2429,NC\_009012.1\_2430 | 2 | RM\_Type\_II\_\_Type\_II\_MTases\_FAM\_2,RM\_Type\_II\_\_Type\_II\_REase30 |
| NC\_009012.1\_RM\_Type\_II\_7 | RM | RM\_Type\_II | NC\_009012.1\_2587 | NC\_009012.1\_2588 | NC\_009012.1\_2587,NC\_009012.1\_2588 | 2 | RM\_Type\_II\_\_Type\_II\_MTases\_FAM\_4,RM\_Type\_II\_\_Type\_II\_REase07 |
| NC\_009012.1\_CAS\_Class1-Subtype-I-B\_10 | Cas | CAS\_Class1-Subtype-I-B | NC\_009012.1\_3357 | NC\_009012.1\_3361 | NC\_009012.1\_3357,NC\_009012.1\_3358,NC\_009012.1\_3359,NC\_009012.1\_3360,NC\_009012.1\_3361 | 5 | cas3\_I\_5,cas5\_I-B\_17,cas6\_I\_II\_III\_IV\_V\_VI\_12,cas7\_I-B\_8,cas8b1\_I-B\_4 |
| NC\_009012.1\_CAS\_Class1-Subtype-III-D\_12 | Cas | CAS\_Class1-Subtype-III-D | NC\_009012.1\_3361 | NC\_009012.1\_3378 | NC\_009012.1\_3361,NC\_009012.1\_3365,NC\_009012.1\_3366,NC\_009012.1\_3367,NC\_009012.1\_3368,NC\_009012.1\_3369,NC\_009012.1\_3370,NC\_009012.1\_3372,NC\_009012.1\_3375,NC\_009012.1\_3376,NC\_009012.1\_3377,NC\_009012.1\_3378 | 12 | cas10\_III-D\_3,cas1\_I\_II\_III\_IV\_V\_VI\_10,cas2\_I\_II\_III\_IV\_V\_VI\_3,cas4\_I\_II\_III\_IV\_V\_VI\_1,cas6\_I\_II\_III\_IV\_V\_VI\_12,csm3gr7\_III-D\_2,csm3gr7\_III-D\_3,csm3gr7\_III\_1,csx10gr5\_III-D\_3,csx19\_III-D\_11,csx1\_III\_21,csx1\_III\_9 |
| NC\_009012.1\_CAS\_Class1-Subtype-III-D\_12 | Cas | CAS\_Class1-Subtype-III-D | NC\_009012.1\_3361 | NC\_009012.1\_3378 | NC\_009012.1\_3361,NC\_009012.1\_3365,NC\_009012.1\_3366,NC\_009012.1\_3367,NC\_009012.1\_3368,NC\_009012.1\_3369,NC\_009012.1\_3370,NC\_009012.1\_3372,NC\_009012.1\_3375,NC\_009012.1\_3376,NC\_009012.1\_3377,NC\_009012.1\_3378 | 12 | cas10\_III-D\_3,cas1\_I\_II\_III\_IV\_V\_VI\_10,cas2\_I\_II\_III\_IV\_V\_VI\_3,cas4\_I\_II\_III\_IV\_V\_VI\_1,cas6\_I\_II\_III\_IV\_V\_VI\_12,csm3gr7\_III-D\_2,csm3gr7\_III-D\_3,csm3gr7\_III\_1,csx10gr5\_III-D\_3,csx19\_III-D\_11,csx1\_III\_21,csx1\_III\_9 |

**Genomad and CheckV Summary**:

| seq\_name | length | gene\_count | viral\_genes | checkv\_quality | miuvig\_quality | taxonomy | topology | coordinates |
| --- | --- | --- | --- | --- | --- | --- | --- | --- |
| NC\_009012.1\|provirus\_1938476\_1983993 | 45518 | 52 | 23 | Medium-quality | Genome-fragment | Viruses;Duplodnaviria;Heunggongvirae;Uroviricota;Caudoviricetes;; | Provirus | 1938476-1983993 |
| NC\_009012.1\|provirus\_2921887\_2969530 | 47644 | 65 | 28 | Medium-quality | Genome-fragment | Viruses;Duplodnaviria;Heunggongvirae;Uroviricota;Caudoviricetes;; | Provirus | 2921887-2969530 |
| NC\_009012.1\|provirus\_2022140\_2067593 | 45454 | 54 | 21 | Medium-quality | Genome-fragment | Viruses;Duplodnaviria;Heunggongvirae;Uroviricota;Caudoviricetes;; | Provirus | 2022140-2067593 |
| NC\_009012.1\|provirus\_3339258\_3389798 | 50541 | 55 | 6 | Medium-quality | Genome-fragment | Viruses;Duplodnaviria;Heunggongvirae;Uroviricota;Caudoviricetes;; | Provirus | 3339258-3389798 |
| NC\_009012.1\|provirus\_2330336\_2378640 | 48305 | 46 | 3 | Low-quality | Genome-fragment | Viruses;Duplodnaviria;Heunggongvirae;Uroviricota;Caudoviricetes;; | Provirus | 2330336-2378640 |

**Host Genome Ideogram with Phages**:

In
this circular plot, **“c”** indicates the contig, and the
number that follows (e.g., **c1**) represents the contig
number. If multiple contigs are present in the genome, each will be
shown with a distinct label (e.g., **c1**,
**c2**, etc.).

##### 2.5.2 Prophages

**Select prophage to show:**

###### 2.5.2.1 Phage ID: NC\_009012.1|provirus\_1938476\_1983993

**Phage-Host Genome Ideogram:**

**Genes Annotation (geNomad):**

| gene | length | marker | annotation\_accessions | annotation\_description |
| --- | --- | --- | --- | --- |
| NC\_009012.1\|provirus\_1938476\_1983993\_1694 | 492 | NA | NA | NA |
| NC\_009012.1\|provirus\_1938476\_1983993\_1695 | 1569 | GENOMAD.097162.VV | PF20307;TIGR03600;COG0305;K18947 | phage replicative helicase, DnaB family, HK022 subfamily |
| NC\_009012.1\|provirus\_1938476\_1983993\_1696 | 219 | GENOMAD.180442.VC | PF20612 | SHOCT domain |
| NC\_009012.1\|provirus\_1938476\_1983993\_1697 | 1005 | GENOMAD.215835.VV | PF01510;COG5632;K11066 | N-acetylmuramoyl-L-alanine amidase CwlA |
| NC\_009012.1\|provirus\_1938476\_1983993\_1698 | 420 | GENOMAD.083633.VV | NA | NA |
| NC\_009012.1\|provirus\_1938476\_1983993\_1699 | 2469 | GENOMAD.116546.VV | PF06605;TIGR01665;COG4926 | phage minor structural protein, N-terminal region |
| NC\_009012.1\|provirus\_1938476\_1983993\_1700 | 576 | GENOMAD.066153.VV | PF06605;TIGR01665;COG4926 | phage minor structural protein, N-terminal region |
| NC\_009012.1\|provirus\_1938476\_1983993\_1701 | 195 | GENOMAD.089015.VV | NA | NA |
| NC\_009012.1\|provirus\_1938476\_1983993\_1702 | 2523 | GENOMAD.116546.VV | PF06605;TIGR01665;COG4926 | phage minor structural protein, N-terminal region |
| NC\_009012.1\|provirus\_1938476\_1983993\_1703 | 774 | GENOMAD.116614.VV | PF20195;COG4722;TIGR01633 | Phage-related protein |
| NC\_009012.1\|provirus\_1938476\_1983993\_1704 | 2289 | GENOMAD.036973.VV | K02334;COG4722 | Phage-related protein |
| NC\_009012.1\|provirus\_1938476\_1983993\_1705 | 273 | NA | NA | NA |
| NC\_009012.1\|provirus\_1938476\_1983993\_1706 | 405 | NA | NA | NA |
| NC\_009012.1\|provirus\_1938476\_1983993\_1707 | 192 | GENOMAD.223693.VP | PF09550;TIGR02216 | Phage tail assembly chaperone protein, TAC |
| NC\_009012.1\|provirus\_1938476\_1983993\_1708 | 384 | GENOMAD.166825.VP | PF11836 | Phage tail tube protein, GTA-gp10 |
| NC\_009012.1\|provirus\_1938476\_1983993\_1709 | 597 | GENOMAD.005050.VV | PF04630;TIGR01603 | phage major tail protein, phi13 family |
| NC\_009012.1\|provirus\_1938476\_1983993\_1710 | 345 | GENOMAD.077616.VV | PF05657 | NA |
| NC\_009012.1\|provirus\_1938476\_1983993\_1711 | 432 | GENOMAD.032585.VV | PF11114;COG5005;TIGR01725 | Mu-like prophage protein gpG |
| NC\_009012.1\|provirus\_1938476\_1983993\_1712 | 336 | GENOMAD.040271.VV | PF05521;COG5614;TIGR01563 | Bacteriophage head-tail adaptor |
| NC\_009012.1\|provirus\_1938476\_1983993\_1713 | 309 | GENOMAD.028909.VV | PF05135;TIGR01560 | Phage gp6-like head-tail connector protein |
| NC\_009012.1\|provirus\_1938476\_1983993\_1714 | 1203 | GENOMAD.168658.VV | PF05065;PF04586;PF12518;COG4653;TIGR01554;K06904 | Predicted phage phi-C31 gp36 major capsid-like protein |
| NC\_009012.1\|provirus\_1938476\_1983993\_1715 | 729 | GENOMAD.028909.VV | PF05135;TIGR01560 | Phage gp6-like head-tail connector protein |
| NC\_009012.1\|provirus\_1938476\_1983993\_1716 | 1323 | GENOMAD.179073.VP | NA | NA |
| NC\_009012.1\|provirus\_1938476\_1983993\_1717 | 1785 | NA | NA | NA |
| NC\_009012.1\|provirus\_1938476\_1983993\_1718 | 1230 | GENOMAD.181434.VP | PF20441 | Terminase large subunit, endonuclease domain |
| NC\_009012.1\|provirus\_1938476\_1983993\_1719 | 336 | NA | NA | NA |
| NC\_009012.1\|provirus\_1938476\_1983993\_1720 | 195 | GENOMAD.140940.VV | PF16468 | NA |
| NC\_009012.1\|provirus\_1938476\_1983993\_1721 | 468 | GENOMAD.191984.VP | PF07128 | NA |
| NC\_009012.1\|provirus\_1938476\_1983993\_1722 | 900 | GENOMAD.105515.VV | NA | NA |
| NC\_009012.1\|provirus\_1938476\_1983993\_1723 | 231 | NA | NA | NA |
| NC\_009012.1\|provirus\_1938476\_1983993\_1724 | 693 | GENOMAD.105515.VV | NA | NA |
| NC\_009012.1\|provirus\_1938476\_1983993\_1725 | 1254 | GENOMAD.208446.VP | PF01555;COG4725;K13581 | N6-adenosine-specific RNA methylase IME4 |
| NC\_009012.1\|provirus\_1938476\_1983993\_1726 | 1299 | GENOMAD.005053.VV | NA | NA |
| NC\_009012.1\|provirus\_1938476\_1983993\_1727 | 183 | GENOMAD.195270.VP | NA | NA |
| NC\_009012.1\|provirus\_1938476\_1983993\_1728 | 552 | GENOMAD.098194.VV | PF05119;COG3747;TIGR01558 | Phage terminase, small subunit |
| NC\_009012.1\|provirus\_1938476\_1983993\_1729 | 360 | GENOMAD.112725.VV | PF16786 | Recombination enhancement, RecA-dependent nuclease |
| NC\_009012.1\|provirus\_1938476\_1983993\_1730 | 243 | GENOMAD.212426.VP | NA | NA |
| NC\_009012.1\|provirus\_1938476\_1983993\_1731 | 999 | NA | NA | NA |
| NC\_009012.1\|provirus\_1938476\_1983993\_1732 | 411 | NA | NA | NA |
| NC\_009012.1\|provirus\_1938476\_1983993\_1733 | 456 | GENOMAD.056402.VV | PF05263;TIGR01636 | phage transcriptional activator, RinA family |
| NC\_009012.1\|provirus\_1938476\_1983993\_1734 | 303 | GENOMAD.138066.VP | PF03838;COG3331;TIGR00648 | Penicillin-binding protein-related factor A, putative recombinase |
| NC\_009012.1\|provirus\_1938476\_1983993\_1735 | 2556 | NA | NA | NA |
| NC\_009012.1\|provirus\_1938476\_1983993\_1736 | 426 | GENOMAD.179451.VC | PF24963 | NA |
| NC\_009012.1\|provirus\_1938476\_1983993\_1737 | 798 | GENOMAD.153366.VP | PF03374;COG3645 | Phage antirepressor protein YoqD, KilAC domain |
| NC\_009012.1\|provirus\_1938476\_1983993\_1738 | 1923 | GENOMAD.166868.VC | PF00476;COG0749 | DNA polymerase I - 3’-5’ exonuclease and polymerase domains |
| NC\_009012.1\|provirus\_1938476\_1983993\_1739 | 753 | GENOMAD.211539.VV | NA | NA |
| NC\_009012.1\|provirus\_1938476\_1983993\_1740 | 441 | GENOMAD.026103.VV | PF23984 | Pam3 gp33-like protein |
| NC\_009012.1\|provirus\_1938476\_1983993\_1741 | 1359 | GENOMAD.003266.VV | PF00176;K14440;COG1061;TIGR04095 | Superfamily II DNA or RNA helicase |
| NC\_009012.1\|provirus\_1938476\_1983993\_1742 | 183 | NA | NA | NA |
| NC\_009012.1\|provirus\_1938476\_1983993\_1743 | 1920 | GENOMAD.213960.VV | NA | NA |
| NC\_009012.1\|provirus\_1938476\_1983993\_1744 | 207 | NA | NA | NA |
| NC\_009012.1\|provirus\_1938476\_1983993\_1745 | 948 | NA | NA | NA |

###### 2.5.2.2 Phage ID: NC\_009012.1|provirus\_2921887\_2969530

**Phage-Host Genome Ideogram:**

**Genes Annotation (geNomad):**

| gene | length | marker | annotation\_accessions | annotation\_description |
| --- | --- | --- | --- | --- |
| NC\_009012.1\|provirus\_2921887\_2969530\_2577 | 3264 | GENOMAD.064018.VV | PF13872;K14440;COG1111;TIGR04095 | ERCC4-related helicase |
| NC\_009012.1\|provirus\_2921887\_2969530\_2578 | 1809 | NA | NA | NA |
| NC\_009012.1\|provirus\_2921887\_2969530\_2579 | 357 | NA | NA | NA |
| NC\_009012.1\|provirus\_2921887\_2969530\_2580 | 3129 | NA | NA | NA |
| NC\_009012.1\|provirus\_2921887\_2969530\_2581 | 1113 | GENOMAD.016861.VV | PF13671;COG2019;TIGR01359;K13829 | AAA domain |
| NC\_009012.1\|provirus\_2921887\_2969530\_2582 | 423 | NA | NA | NA |
| NC\_009012.1\|provirus\_2921887\_2969530\_2583 | 207 | GENOMAD.087773.VV | PF14083 | PGDYG protein |
| NC\_009012.1\|provirus\_2921887\_2969530\_2584 | 465 | GENOMAD.170160.VC | PF18184 | SMODS and SLOG-associating 2TM effector domain 3 |
| NC\_009012.1\|provirus\_2921887\_2969530\_2585 | 441 | NA | NA | NA |
| NC\_009012.1\|provirus\_2921887\_2969530\_2586 | 429 | NA | NA | NA |
| NC\_009012.1\|provirus\_2921887\_2969530\_2587 | 240 | NA | NA | NA |
| NC\_009012.1\|provirus\_2921887\_2969530\_2588 | 747 | GENOMAD.132419.VP | PF03374;COG3645 | Phage antirepressor protein YoqD, KilAC domain |
| NC\_009012.1\|provirus\_2921887\_2969530\_2589 | 219 | NA | NA | NA |
| NC\_009012.1\|provirus\_2921887\_2969530\_2590 | 213 | GENOMAD.209016.VC | NA | NA |
| NC\_009012.1\|provirus\_2921887\_2969530\_2591 | 174 | NA | NA | NA |
| NC\_009012.1\|provirus\_2921887\_2969530\_2592 | 219 | GENOMAD.088748.VV | NA | NA |
| NC\_009012.1\|provirus\_2921887\_2969530\_2593 | 552 | NA | NA | NA |
| NC\_009012.1\|provirus\_2921887\_2969530\_2594 | 906 | NA | NA | NA |
| NC\_009012.1\|provirus\_2921887\_2969530\_2595 | 288 | NA | NA | NA |
| NC\_009012.1\|provirus\_2921887\_2969530\_2596 | 189 | NA | NA | NA |
| NC\_009012.1\|provirus\_2921887\_2969530\_2597 | 1083 | GENOMAD.021533.VV | TIGR01714 | phage replisome organizer, putative, N-terminal region |
| NC\_009012.1\|provirus\_2921887\_2969530\_2598 | 837 | NA | NA | NA |
| NC\_009012.1\|provirus\_2921887\_2969530\_2599 | 375 | GENOMAD.151656.VP | PF17288;K06909;TIGR01547;COG1783 | phage terminase, large subunit, PBSX family |
| NC\_009012.1\|provirus\_2921887\_2969530\_2600 | 423 | GENOMAD.221128.VP | PF05263;TIGR01636;COG2739 | phage transcriptional activator, RinA family |
| NC\_009012.1\|provirus\_2921887\_2969530\_2601 | 243 | NA | NA | NA |
| NC\_009012.1\|provirus\_2921887\_2969530\_2602 | 843 | GENOMAD.178342.VP | PF02086;PF06576;PF06147;K21507;TIGR00571;COG0338 | DNA adenine methylase (dam) |
| NC\_009012.1\|provirus\_2921887\_2969530\_2603 | 1281 | NA | NA | NA |
| NC\_009012.1\|provirus\_2921887\_2969530\_2604 | 846 | GENOMAD.225783.VP | NA | NA |
| NC\_009012.1\|provirus\_2921887\_2969530\_2605 | 150 | NA | NA | NA |
| NC\_009012.1\|provirus\_2921887\_2969530\_2606 | 450 | GENOMAD.026460.VV | PF03592;COG3728;K07474 | Phage terminase, small subunit |
| NC\_009012.1\|provirus\_2921887\_2969530\_2607 | 1248 | GENOMAD.141197.VP | PF17288;K06909;TIGR01547;COG1783 | Terminase RNAseH like domain |
| NC\_009012.1\|provirus\_2921887\_2969530\_2608 | 1434 | GENOMAD.006049.VV | NA | NA |
| NC\_009012.1\|provirus\_2921887\_2969530\_2609 | 150 | NA | NA | NA |
| NC\_009012.1\|provirus\_2921887\_2969530\_2610 | 1410 | GENOMAD.003968.VV | PF06152;COG5585;TIGR01641 | NAD+–asparagine ADP-ribosyltransferase |
| NC\_009012.1\|provirus\_2921887\_2969530\_2611 | 207 | GENOMAD.066155.VV | PF06372 | NA |
| NC\_009012.1\|provirus\_2921887\_2969530\_2612 | 687 | GENOMAD.007545.VV | PF06810 | Phage minor structural protein GP20 |
| NC\_009012.1\|provirus\_2921887\_2969530\_2613 | 924 | GENOMAD.092843.VP | NA | NA |
| NC\_009012.1\|provirus\_2921887\_2969530\_2614 | 147 | NA | NA | NA |
| NC\_009012.1\|provirus\_2921887\_2969530\_2615 | 402 | GENOMAD.047671.VV | PF11436 | NA |
| NC\_009012.1\|provirus\_2921887\_2969530\_2616 | 336 | GENOMAD.074010.VV | PF12206 | NA |
| NC\_009012.1\|provirus\_2921887\_2969530\_2617 | 411 | GENOMAD.019155.VV | PF11114;TIGR01725 | phage protein, HK97 gp10 family |
| NC\_009012.1\|provirus\_2921887\_2969530\_2618 | 420 | GENOMAD.025660.VV | PF20765 | Phage tail terminator protein |
| NC\_009012.1\|provirus\_2921887\_2969530\_2619 | 1290 | NA | NA | NA |
| NC\_009012.1\|provirus\_2921887\_2969530\_2620 | 1047 | GENOMAD.118787.VV | PF04984;PF17482;COG3497 | Phage tail sheath protein subtilisin-like domain; Phage tail sheath C-terminal domain |
| NC\_009012.1\|provirus\_2921887\_2969530\_2621 | 477 | GENOMAD.018728.VV | PF09393 | Phage tail tube protein |
| NC\_009012.1\|provirus\_2921887\_2969530\_2622 | 414 | GENOMAD.009389.VV | PF08890;PF17482 | Phage XkdN-like tail assembly chaperone protein, TAC; Phage tail sheath C-terminal domain |
| NC\_009012.1\|provirus\_2921887\_2969530\_2623 | 192 | GENOMAD.059912.VV | NA | NA |
| NC\_009012.1\|provirus\_2921887\_2969530\_2624 | 1836 | GENOMAD.118787.VV | PF04984;PF17482;COG3497 | Phage tail sheath protein subtilisin-like domain; Phage tail sheath C-terminal domain |
| NC\_009012.1\|provirus\_2921887\_2969530\_2625 | 660 | GENOMAD.017357.VV | PF06995;COG1652 | Nucleoid-associated protein YgaU, contains BON and LysM domains |
| NC\_009012.1\|provirus\_2921887\_2969530\_2626 | 945 | GENOMAD.018966.VV | PF14594;COG4379;TIGR03361;K06905 | Mu-like prophage tail protein gpP |
| NC\_009012.1\|provirus\_2921887\_2969530\_2627 | 234 | GENOMAD.096038.VV | PF10844 | NA |
| NC\_009012.1\|provirus\_2921887\_2969530\_2628 | 396 | GENOMAD.016318.VV | PF10934;COG4381;TIGR03357 | Mu-like prophage protein gp46 |
| NC\_009012.1\|provirus\_2921887\_2969530\_2629 | 1059 | GENOMAD.004833.VV | PF04865;COG3299 | Baseplate J-like protein |
| NC\_009012.1\|provirus\_2921887\_2969530\_2630 | 603 | GENOMAD.015577.VV | PF10076;COG3778;TIGR02242 | Uncharacterized protein YmfQ in lambdoid prophage, DUF2313 family |
| NC\_009012.1\|provirus\_2921887\_2969530\_2631 | 285 | NA | NA | NA |
| NC\_009012.1\|provirus\_2921887\_2969530\_2632 | 1383 | GENOMAD.087125.VV | NA | NA |
| NC\_009012.1\|provirus\_2921887\_2969530\_2633 | 261 | GENOMAD.213587.VP | NA | NA |
| NC\_009012.1\|provirus\_2921887\_2969530\_2634 | 153 | GENOMAD.065816.VV | PF09693;TIGR01669 | phage uncharacterized protein, XkdX family |
| NC\_009012.1\|provirus\_2921887\_2969530\_2635 | 291 | GENOMAD.171716.VV | PF07439 | NA |
| NC\_009012.1\|provirus\_2921887\_2969530\_2636 | 657 | GENOMAD.220135.VV | COG5632 | N-acetylmuramoyl-L-alanine amidase CwlA |
| NC\_009012.1\|provirus\_2921887\_2969530\_2637 | 300 | GENOMAD.151236.VC | PF06946 | Bacteriophage A118-like holin, Hol118 |
| NC\_009012.1\|provirus\_2921887\_2969530\_2638 | 636 | NA | NA | NA |
| NC\_009012.1\|provirus\_2921887\_2969530\_2639 | 432 | GENOMAD.206152.VV | PF01845;COG2337;K07171 | mRNA-degrading endonuclease, toxin component of the MazEF toxin-antitoxin module |
| NC\_009012.1\|provirus\_2921887\_2969530\_2640 | 486 | NA | NA | NA |
| NC\_009012.1\|provirus\_2921887\_2969530\_2641 | 165 | NA | NA | NA |

###### 2.5.2.3 Phage ID: NC\_009012.1|provirus\_2022140\_2067593

**Phage-Host Genome Ideogram:**

**Genes Annotation (geNomad):**

| gene | length | marker | annotation\_accessions | annotation\_description |
| --- | --- | --- | --- | --- |
| NC\_009012.1\|provirus\_2022140\_2067593\_1786 | 1329 | NA | NA | NA |
| NC\_009012.1\|provirus\_2022140\_2067593\_1787 | 1290 | GENOMAD.022372.VV | PF04708;COG1961;K14060 | Site-specific DNA recombinase related to the DNA invertase Pin |
| NC\_009012.1\|provirus\_2022140\_2067593\_1788 | 672 | NA | NA | NA |
| NC\_009012.1\|provirus\_2022140\_2067593\_1789 | 486 | NA | NA | NA |
| NC\_009012.1\|provirus\_2022140\_2067593\_1790 | 411 | GENOMAD.083633.VV | NA | NA |
| NC\_009012.1\|provirus\_2022140\_2067593\_1791 | 1689 | GENOMAD.068767.VV | NA | NA |
| NC\_009012.1\|provirus\_2022140\_2067593\_1792 | 1158 | GENOMAD.222380.VP | NA | NA |
| NC\_009012.1\|provirus\_2022140\_2067593\_1793 | 1905 | GENOMAD.004340.VV | PF06605;PF18994;COG4926;TIGR01665 | Phage-related protein |
| NC\_009012.1\|provirus\_2022140\_2067593\_1794 | 705 | GENOMAD.004006.VV | PF20195;COG4722;TIGR01633 | Phage-related protein |
| NC\_009012.1\|provirus\_2022140\_2067593\_1795 | 2280 | GENOMAD.089120.VP | NA | NA |
| NC\_009012.1\|provirus\_2022140\_2067593\_1796 | 126 | GENOMAD.202970.VV | PF09550;TIGR02216 | NA |
| NC\_009012.1\|provirus\_2022140\_2067593\_1797 | 324 | GENOMAD.136836.VV | PF16478 | Phage tail tube protein, GTA-gp10 |
| NC\_009012.1\|provirus\_2022140\_2067593\_1798 | 441 | GENOMAD.053493.CC | PF14101 | NA |
| NC\_009012.1\|provirus\_2022140\_2067593\_1799 | 1044 | NA | NA | NA |
| NC\_009012.1\|provirus\_2022140\_2067593\_1800 | 108 | NA | NA | NA |
| NC\_009012.1\|provirus\_2022140\_2067593\_1801 | 573 | GENOMAD.013991.VV | PF04630;TIGR01603 | phage major tail protein, phi13 family |
| NC\_009012.1\|provirus\_2022140\_2067593\_1802 | 330 | GENOMAD.018077.VV | PF05657 | NA |
| NC\_009012.1\|provirus\_2022140\_2067593\_1803 | 393 | GENOMAD.066487.VV | TIGR01725;COG5005 | phage protein, HK97 gp10 family |
| NC\_009012.1\|provirus\_2022140\_2067593\_1804 | 327 | GENOMAD.124210.VP | PF05521;COG5614;TIGR01563 | Bacteriophage head-tail adaptor |
| NC\_009012.1\|provirus\_2022140\_2067593\_1805 | 600 | GENOMAD.158311.VV | PF05135;TIGR02215 | phage conserved hypothetical protein, phiE125 gp8 family |
| NC\_009012.1\|provirus\_2022140\_2067593\_1806 | 441 | NA | NA | NA |
| NC\_009012.1\|provirus\_2022140\_2067593\_1807 | 1290 | GENOMAD.093418.VV | PF05065;TIGR01554;COG4653 | phage major capsid protein, HK97 family |
| NC\_009012.1\|provirus\_2022140\_2067593\_1808 | 744 | GENOMAD.158277.VP | PF00574;PF19602;K01358;TIGR00493;COG3904 | ATP-dependent Clp endopeptidase, proteolytic subunit ClpP |
| NC\_009012.1\|provirus\_2022140\_2067593\_1809 | 1260 | GENOMAD.179073.VP | NA | NA |
| NC\_009012.1\|provirus\_2022140\_2067593\_1810 | 2550 | GENOMAD.181434.VP | PF20441 | Terminase large subunit, endonuclease domain |
| NC\_009012.1\|provirus\_2022140\_2067593\_1811 | 483 | GENOMAD.168120.VP | PF05119;COG3747;TIGR01558 | Phage terminase, small subunit |
| NC\_009012.1\|provirus\_2022140\_2067593\_1812 | 216 | GENOMAD.225559.VP | NA | NA |
| NC\_009012.1\|provirus\_2022140\_2067593\_1813 | 753 | GENOMAD.105515.VV | NA | NA |
| NC\_009012.1\|provirus\_2022140\_2067593\_1814 | 312 | GENOMAD.224340.VP | PF19854 | NA |
| NC\_009012.1\|provirus\_2022140\_2067593\_1815 | 480 | GENOMAD.191984.VP | PF07128 | NA |
| NC\_009012.1\|provirus\_2022140\_2067593\_1816 | 924 | GENOMAD.105515.VV | NA | NA |
| NC\_009012.1\|provirus\_2022140\_2067593\_1817 | 1452 | GENOMAD.136632.VV | PF00145;TIGR00675;K00558;COG0270 | DNA-methyltransferase (dcm) |
| NC\_009012.1\|provirus\_2022140\_2067593\_1818 | 1236 | GENOMAD.038338.VV | COG3392 | NA |
| NC\_009012.1\|provirus\_2022140\_2067593\_1819 | 426 | GENOMAD.133740.VV | PF07750;COG5352;K13583;TIGR00721 | GcrA cell cycle regulator |
| NC\_009012.1\|provirus\_2022140\_2067593\_1820 | 381 | GENOMAD.219652.VP | NA | NA |
| NC\_009012.1\|provirus\_2022140\_2067593\_1821 | 246 | NA | NA | NA |
| NC\_009012.1\|provirus\_2022140\_2067593\_1822 | 405 | GENOMAD.159035.VP | PF07374;TIGR01636;COG2739 | phage transcriptional activator, RinA family |
| NC\_009012.1\|provirus\_2022140\_2067593\_1823 | 303 | GENOMAD.212426.VP | NA | NA |
| NC\_009012.1\|provirus\_2022140\_2067593\_1824 | 1875 | GENOMAD.024099.VV | NA | NA |
| NC\_009012.1\|provirus\_2022140\_2067593\_1825 | 1692 | GENOMAD.102034.VP | TIGR01636 | NA |
| NC\_009012.1\|provirus\_2022140\_2067593\_1826 | 459 | GENOMAD.031678.VV | PF05037 | NA |
| NC\_009012.1\|provirus\_2022140\_2067593\_1827 | 1431 | GENOMAD.016341.VV | PF13479;PF12684;TIGR01618;K07465;COG1468 | phage nucleotide-binding protein |
| NC\_009012.1\|provirus\_2022140\_2067593\_1828 | 366 | GENOMAD.010330.VV | PF03838;COG3331;TIGR00648 | Recombination protein U |
| NC\_009012.1\|provirus\_2022140\_2067593\_1829 | 1179 | NA | NA | NA |
| NC\_009012.1\|provirus\_2022140\_2067593\_1830 | 90 | NA | NA | NA |
| NC\_009012.1\|provirus\_2022140\_2067593\_1831 | 765 | GENOMAD.107118.VP | NA | NA |
| NC\_009012.1\|provirus\_2022140\_2067593\_1832 | 252 | GENOMAD.222352.VV | NA | NA |
| NC\_009012.1\|provirus\_2022140\_2067593\_1833 | 324 | NA | NA | NA |
| NC\_009012.1\|provirus\_2022140\_2067593\_1834 | 228 | NA | NA | NA |
| NC\_009012.1\|provirus\_2022140\_2067593\_1835 | 711 | NA | NA | NA |
| NC\_009012.1\|provirus\_2022140\_2067593\_1836 | 369 | GENOMAD.213960.VV | NA | NA |
| NC\_009012.1\|provirus\_2022140\_2067593\_1837 | 381 | NA | NA | NA |
| NC\_009012.1\|provirus\_2022140\_2067593\_1838 | 789 | NA | NA | NA |
| NC\_009012.1\|provirus\_2022140\_2067593\_1839 | 417 | NA | NA | NA |
| NC\_009012.1\|provirus\_2022140\_2067593\_1840 | 408 | NA | NA | NA |

###### 2.5.2.4 Phage ID: NC\_009012.1|provirus\_3339258\_3389798

**Phage-Host Genome Ideogram:**

**Genes Annotation (geNomad):**

| gene | length | marker | annotation\_accessions | annotation\_description |
| --- | --- | --- | --- | --- |
| NC\_009012.1\|provirus\_3339258\_3389798\_2977 | 240 | NA | NA | NA |
| NC\_009012.1\|provirus\_3339258\_3389798\_2978 | 807 | GENOMAD.169139.VV | PF09250 | Bifunctional DNA primase/polymerase, N-terminal |
| NC\_009012.1\|provirus\_3339258\_3389798\_2979 | 1848 | NA | NA | NA |
| NC\_009012.1\|provirus\_3339258\_3389798\_2980 | 1239 | NA | NA | NA |
| NC\_009012.1\|provirus\_3339258\_3389798\_2981 | 138 | NA | NA | NA |
| NC\_009012.1\|provirus\_3339258\_3389798\_2982 | 330 | NA | NA | NA |
| NC\_009012.1\|provirus\_3339258\_3389798\_2983 | 471 | NA | NA | NA |
| NC\_009012.1\|provirus\_3339258\_3389798\_2984 | 405 | NA | NA | NA |
| NC\_009012.1\|provirus\_3339258\_3389798\_2985 | 759 | NA | NA | NA |
| NC\_009012.1\|provirus\_3339258\_3389798\_2986 | 1485 | NA | NA | NA |
| NC\_009012.1\|provirus\_3339258\_3389798\_2987 | 717 | NA | NA | NA |
| NC\_009012.1\|provirus\_3339258\_3389798\_2988 | 348 | NA | NA | NA |
| NC\_009012.1\|provirus\_3339258\_3389798\_2989 | 1686 | NA | NA | NA |
| NC\_009012.1\|provirus\_3339258\_3389798\_2990 | 579 | NA | NA | NA |
| NC\_009012.1\|provirus\_3339258\_3389798\_2991 | 390 | NA | NA | NA |
| NC\_009012.1\|provirus\_3339258\_3389798\_2992 | 138 | GENOMAD.223510.VP | PF10122;TIGR04104;COG4530 | Mu-like prophage protein Com |
| NC\_009012.1\|provirus\_3339258\_3389798\_2993 | 552 | NA | NA | NA |
| NC\_009012.1\|provirus\_3339258\_3389798\_2994 | 1842 | NA | NA | NA |
| NC\_009012.1\|provirus\_3339258\_3389798\_2995 | 1239 | NA | NA | NA |
| NC\_009012.1\|provirus\_3339258\_3389798\_2996 | 345 | NA | NA | NA |
| NC\_009012.1\|provirus\_3339258\_3389798\_2997 | 435 | NA | NA | NA |
| NC\_009012.1\|provirus\_3339258\_3389798\_2998 | 345 | NA | NA | NA |
| NC\_009012.1\|provirus\_3339258\_3389798\_2999 | 246 | NA | NA | NA |
| NC\_009012.1\|provirus\_3339258\_3389798\_3000 | 189 | NA | NA | NA |
| NC\_009012.1\|provirus\_3339258\_3389798\_3001 | 1206 | NA | NA | NA |
| NC\_009012.1\|provirus\_3339258\_3389798\_3002 | 288 | NA | NA | NA |
| NC\_009012.1\|provirus\_3339258\_3389798\_3003 | 390 | NA | NA | NA |
| NC\_009012.1\|provirus\_3339258\_3389798\_3004 | 141 | GENOMAD.223510.VP | PF10122;TIGR04104;COG4530 | Mu-like prophage protein Com |
| NC\_009012.1\|provirus\_3339258\_3389798\_3005 | 2160 | GENOMAD.197438.VP | PF09250 | Bifunctional DNA primase/polymerase, N-terminal |
| NC\_009012.1\|provirus\_3339258\_3389798\_3006 | 1314 | GENOMAD.093418.VV | PF05065;TIGR01554;COG4653 | phage major capsid protein, HK97 family |
| NC\_009012.1\|provirus\_3339258\_3389798\_3007 | 303 | NA | NA | NA |
| NC\_009012.1\|provirus\_3339258\_3389798\_3008 | 1245 | NA | NA | NA |
| NC\_009012.1\|provirus\_3339258\_3389798\_3009 | 288 | NA | NA | NA |
| NC\_009012.1\|provirus\_3339258\_3389798\_3010 | 297 | NA | NA | NA |
| NC\_009012.1\|provirus\_3339258\_3389798\_3011 | 807 | GENOMAD.169139.VV | PF09250 | Bifunctional DNA primase/polymerase, N-terminal |
| NC\_009012.1\|provirus\_3339258\_3389798\_3012 | 1848 | NA | NA | NA |
| NC\_009012.1\|provirus\_3339258\_3389798\_3013 | 1239 | NA | NA | NA |
| NC\_009012.1\|provirus\_3339258\_3389798\_3014 | 528 | NA | NA | NA |
| NC\_009012.1\|provirus\_3339258\_3389798\_3015 | 471 | NA | NA | NA |
| NC\_009012.1\|provirus\_3339258\_3389798\_3016 | 1137 | NA | NA | NA |
| NC\_009012.1\|provirus\_3339258\_3389798\_3017 | 348 | NA | NA | NA |
| NC\_009012.1\|provirus\_3339258\_3389798\_3018 | 780 | NA | NA | NA |
| NC\_009012.1\|provirus\_3339258\_3389798\_3019 | 930 | NA | NA | NA |
| NC\_009012.1\|provirus\_3339258\_3389798\_3020 | 243 | GENOMAD.227820.VP | PF15597 | Immunity protein 59 |
| NC\_009012.1\|provirus\_3339258\_3389798\_3021 | 603 | NA | NA | NA |
| NC\_009012.1\|provirus\_3339258\_3389798\_3022 | 1005 | NA | NA | NA |
| NC\_009012.1\|provirus\_3339258\_3389798\_3023 | 237 | GENOMAD.218640.VC | NA | NA |
| NC\_009012.1\|provirus\_3339258\_3389798\_3024 | 390 | NA | NA | NA |
| NC\_009012.1\|provirus\_3339258\_3389798\_3025 | 138 | GENOMAD.223510.VP | PF10122;TIGR04104;COG4530 | Mu-like prophage protein Com |
| NC\_009012.1\|provirus\_3339258\_3389798\_3026 | 2160 | GENOMAD.197438.VP | PF09250 | Bifunctional DNA primase/polymerase, N-terminal |
| NC\_009012.1\|provirus\_3339258\_3389798\_3027 | 1314 | GENOMAD.061958.VV | PF05065 | Phage capsid family |
| NC\_009012.1\|provirus\_3339258\_3389798\_3028 | 303 | NA | NA | NA |
| NC\_009012.1\|provirus\_3339258\_3389798\_3029 | 1245 | NA | NA | NA |
| NC\_009012.1\|provirus\_3339258\_3389798\_3030 | 186 | NA | NA | NA |
| NC\_009012.1\|provirus\_3339258\_3389798\_3031 | 294 | NA | NA | NA |

###### 2.5.2.5 Phage ID: NC\_009012.1|provirus\_2330336\_2378640

**Phage-Host Genome Ideogram:**

**Genes Annotation (geNomad):**

| gene | length | marker | annotation\_accessions | annotation\_description |
| --- | --- | --- | --- | --- |
| NC\_009012.1\|provirus\_2330336\_2378640\_2057 | 933 | NA | NA | NA |
| NC\_009012.1\|provirus\_2330336\_2378640\_2058 | 1341 | NA | NA | NA |
| NC\_009012.1\|provirus\_2330336\_2378640\_2059 | 198 | NA | NA | NA |
| NC\_009012.1\|provirus\_2330336\_2378640\_2060 | 966 | GENOMAD.221871.VP | K13281;TIGR00629;COG4294 | UV damage endonuclease UvdE |
| NC\_009012.1\|provirus\_2330336\_2378640\_2061 | 735 | NA | NA | NA |
| NC\_009012.1\|provirus\_2330336\_2378640\_2062 | 1083 | NA | NA | NA |
| NC\_009012.1\|provirus\_2330336\_2378640\_2063 | 2451 | NA | NA | NA |
| NC\_009012.1\|provirus\_2330336\_2378640\_2064 | 963 | NA | NA | NA |
| NC\_009012.1\|provirus\_2330336\_2378640\_2065 | 2514 | GENOMAD.208624.VC | NA | NA |
| NC\_009012.1\|provirus\_2330336\_2378640\_2066 | 1530 | NA | NA | NA |
| NC\_009012.1\|provirus\_2330336\_2378640\_2067 | 564 | NA | NA | NA |
| NC\_009012.1\|provirus\_2330336\_2378640\_2068 | 141 | NA | NA | NA |
| NC\_009012.1\|provirus\_2330336\_2378640\_2069 | 327 | NA | NA | NA |
| NC\_009012.1\|provirus\_2330336\_2378640\_2070 | 1737 | NA | NA | NA |
| NC\_009012.1\|provirus\_2330336\_2378640\_2071 | 3201 | NA | NA | NA |
| NC\_009012.1\|provirus\_2330336\_2378640\_2072 | 1380 | NA | NA | NA |
| NC\_009012.1\|provirus\_2330336\_2378640\_2073 | 306 | NA | NA | NA |
| NC\_009012.1\|provirus\_2330336\_2378640\_2074 | 330 | NA | NA | NA |
| NC\_009012.1\|provirus\_2330336\_2378640\_2075 | 321 | NA | NA | NA |
| NC\_009012.1\|provirus\_2330336\_2378640\_2076 | 1524 | NA | NA | NA |
| NC\_009012.1\|provirus\_2330336\_2378640\_2077 | 564 | NA | NA | NA |
| NC\_009012.1\|provirus\_2330336\_2378640\_2078 | 948 | NA | NA | NA |
| NC\_009012.1\|provirus\_2330336\_2378640\_2079 | 327 | NA | NA | NA |
| NC\_009012.1\|provirus\_2330336\_2378640\_2080 | 246 | NA | NA | NA |
| NC\_009012.1\|provirus\_2330336\_2378640\_2081 | 1353 | NA | NA | NA |
| NC\_009012.1\|provirus\_2330336\_2378640\_2082 | 807 | GENOMAD.197438.VP | PF09250 | Bifunctional DNA primase/polymerase, N-terminal |
| NC\_009012.1\|provirus\_2330336\_2378640\_2083 | 138 | GENOMAD.223510.VP | PF10122;TIGR04104;COG4530 | Mu-like prophage protein Com |
| NC\_009012.1\|provirus\_2330336\_2378640\_2084 | 390 | NA | NA | NA |
| NC\_009012.1\|provirus\_2330336\_2378640\_2085 | 243 | NA | NA | NA |
| NC\_009012.1\|provirus\_2330336\_2378640\_2086 | 1185 | NA | NA | NA |
| NC\_009012.1\|provirus\_2330336\_2378640\_2087 | 348 | NA | NA | NA |
| NC\_009012.1\|provirus\_2330336\_2378640\_2088 | 1272 | GENOMAD.061958.VV | PF05065 | Phage capsid family |
| NC\_009012.1\|provirus\_2330336\_2378640\_2089 | 2160 | GENOMAD.197438.VP | PF09250 | Bifunctional DNA primase/polymerase, N-terminal |
| NC\_009012.1\|provirus\_2330336\_2378640\_2090 | 138 | GENOMAD.223510.VP | PF10122;TIGR04104;COG4530 | Mu-like prophage protein Com |
| NC\_009012.1\|provirus\_2330336\_2378640\_2091 | 390 | NA | NA | NA |
| NC\_009012.1\|provirus\_2330336\_2378640\_2092 | 279 | GENOMAD.218057.VV | NA | NA |
| NC\_009012.1\|provirus\_2330336\_2378640\_2093 | 894 | NA | NA | NA |
| NC\_009012.1\|provirus\_2330336\_2378640\_2094 | 780 | NA | NA | NA |
| NC\_009012.1\|provirus\_2330336\_2378640\_2095 | 348 | NA | NA | NA |
| NC\_009012.1\|provirus\_2330336\_2378640\_2096 | 987 | NA | NA | NA |
| NC\_009012.1\|provirus\_2330336\_2378640\_2097 | 948 | NA | NA | NA |
| NC\_009012.1\|provirus\_2330336\_2378640\_2098 | 351 | NA | NA | NA |
| NC\_009012.1\|provirus\_2330336\_2378640\_2099 | 1524 | NA | NA | NA |
| NC\_009012.1\|provirus\_2330336\_2378640\_2100 | 726 | NA | NA | NA |
| NC\_009012.1\|provirus\_2330336\_2378640\_2101 | 315 | NA | NA | NA |
| NC\_009012.1\|provirus\_2330336\_2378640\_2102 | 771 | NA | NA | NA |

##### 2.5.3 vOTUs

When more than 1 phage is detected in the host genome, we perform a
clustering step using the tool dRep.

A threshold of 0.95 was applied to the ANI similarity index to define
clusters of virus operational taxonomic units (vOTUs).

**Final cluster designations**

| genomad\_summary$seq\_name | genome | secondary\_cluster | threshold | cluster\_method | comparison\_algorithm | primary\_cluster |
| --- | --- | --- | --- | --- | --- | --- |
| NC\_009012.1\|provirus\_1938476\_1983993 | sequence\_000002.fasta.fasta | 1\_1 | 0.05 | average | ANImf | 1 |
| NC\_009012.1\|provirus\_2921887\_2969530 | sequence\_000004.fasta.fasta | 1\_2 | 0.05 | average | ANImf | 1 |
| NC\_009012.1\|provirus\_2022140\_2067593 | sequence\_000000.fasta.fasta | 2\_0 | 0.05 | average | ANImf | 2 |
| NC\_009012.1\|provirus\_3339258\_3389798 | sequence\_000001.fasta.fasta | 3\_0 | 0.05 | average | ANImf | 3 |
| NC\_009012.1\|provirus\_2330336\_2378640 | sequence\_000003.fasta.fasta | 4\_0 | 0.05 | average | ANImf | 4 |

**Primary clustering plot**

##### 2.5.4 Prophage-Host Prediction

Prediction of potential hosts for the prophage contigs.

| virus | host\_genome | host\_taxonomy | main\_method | confidence\_score | additional\_methods |
| --- | --- | --- | --- | --- | --- |
| NC\_009012.1\|provirus\_1938476\_1983993 | RS\_GCF\_000015865.1 | d\_\_Bacteria;p\_\_Bacillota\_A;c\_\_Clostridia;o\_\_Acetivibrionales;f\_\_Acetivibrionaceae;g\_\_Hungateiclostridium;s\_\_Hungateiclostridium thermocellum | blast | 94.9 | iPHoP-RF;71.30 |
| NC\_009012.1\|provirus\_2022140\_2067593 | RS\_GCF\_000015865.1 | d\_\_Bacteria;p\_\_Bacillota\_A;c\_\_Clostridia;o\_\_Acetivibrionales;f\_\_Acetivibrionaceae;g\_\_Hungateiclostridium;s\_\_Hungateiclostridium thermocellum | blast | 93.9 | None |
| NC\_009012.1\|provirus\_2330336\_2378640 | RS\_GCF\_000015865.1 | d\_\_Bacteria;p\_\_Bacillota\_A;c\_\_Clostridia;o\_\_Acetivibrionales;f\_\_Acetivibrionaceae;g\_\_Hungateiclostridium;s\_\_Hungateiclostridium thermocellum | blast | 97.7 | iPHoP-RF;83.30 |
| NC\_009012.1\|provirus\_2330336\_2378640 | RS\_GCF\_004102745.1 | d\_\_Bacteria;p\_\_Bacillota\_A;c\_\_Clostridia;o\_\_Acetivibrionales;f\_\_Acetivibrionaceae;g\_\_Hungateiclostridium;s\_\_Hungateiclostridium mesophilum | blast | 90.4 | None |
| NC\_009012.1\|provirus\_2330336\_2378640 | 3300025605\_6 | d\_\_Bacteria;p\_\_Bacillota\_A;c\_\_Clostridia;o\_\_Acetivibrionales;f\_\_Acetivibrionaceae;g**Hungateiclostridium;s** | blast | 90.0 | iPHoP-RF;50.70 |
| NC\_009012.1\|provirus\_2921887\_2969530 | RS\_GCF\_000015865.1 | d\_\_Bacteria;p\_\_Bacillota\_A;c\_\_Clostridia;o\_\_Acetivibrionales;f\_\_Acetivibrionaceae;g\_\_Hungateiclostridium;s\_\_Hungateiclostridium thermocellum | blast | 93.1 | iPHoP-RF;51.50 |
| NC\_009012.1\|provirus\_2921887\_2969530 | 3300025605\_6 | d\_\_Bacteria;p\_\_Bacillota\_A;c\_\_Clostridia;o\_\_Acetivibrionales;f\_\_Acetivibrionaceae;g**Hungateiclostridium;s** | blast | 91.8 | None |
| NC\_009012.1\|provirus\_3339258\_3389798 | RS\_GCF\_000015865.1 | d\_\_Bacteria;p\_\_Bacillota\_A;c\_\_Clostridia;o\_\_Acetivibrionales;f\_\_Acetivibrionaceae;g\_\_Hungateiclostridium;s\_\_Hungateiclostridium thermocellum | blast | 98.1 | iPHoP-RF;76.80 |

##### 2.5.5 AMG Predictions

Prediction of auxiliary metabolic genes in the prophage contigs.

| protein | scaffold | amg\_ko | amg\_ko\_name | pfam | pfam\_name |
| --- | --- | --- | --- | --- | --- |
| NC\_009012.1\|provirus\_2022140\_2067593\_32 | NC\_009012.1\|provirus\_2022140\_2067593 | K00558 | DNMT1, dcm; DNA (cytosine-5)-methyltransferase 1 [EC:2.1.1.37] | PF00145.17 | C-5 cytosine-specific DNA methylase |

## 2.6 NC\_012982

##### 2.6.1 Host Genome

**GTDB-Tk taxonomy**:

classification

d\_\_Bacteria;p\_\_Pseudomonadota;c\_\_Alphaproteobacteria;o\_\_Caulobacterales;f\_\_Hyphomonadaceae;g\_\_Hirschia;s\_\_Hirschia
baltica

**CheckM2 Summary**:

| total\_contigs | contig\_n50 | completeness | contamination |
| --- | --- | --- | --- |
| 1 | 3455622 | 100 | 0.05 |

**Defense-Finder Systems**:

| sys\_id | type | subtype | sys\_beg | sys\_end | protein\_in\_syst | genes\_count | name\_of\_profiles\_in\_sys |
| --- | --- | --- | --- | --- | --- | --- | --- |
| NC\_012982.1\_AbiC\_1 | AbiC | AbiC | NC\_012982.1\_420 | NC\_012982.1\_420 | NC\_012982.1\_420 | 1 | AbiC\_\_AbiC |
| NC\_012982.1\_RM\_Type\_IIG\_5 | RM | RM\_Type\_IIG | NC\_012982.1\_623 | NC\_012982.1\_623 | NC\_012982.1\_623 | 1 | RM\_Type\_IIG\_\_Type\_IIG\_FAM\_1.einsi\_trimmed |
| NC\_012982.1\_RM\_Type\_III\_6 | RM | RM\_Type\_III | NC\_012982.1\_1629 | NC\_012982.1\_1630 | NC\_012982.1\_1629,NC\_012982.1\_1630 | 2 | RM\_Type\_III\_\_Type\_III\_MTases\_FAM\_0,RM\_Type\_III\_\_Type\_III\_REases\_FAM\_1.einsi\_trimmed |
| NC\_012982.1\_CBASS\_I\_3 | CBASS | CBASS\_I | NC\_012982.1\_1825 | NC\_012982.1\_1826 | NC\_012982.1\_1825,NC\_012982.1\_1826 | 2 | CBASS\_\_Cyclase\_II,CBASS\_\_Phospholipase |
| NC\_012982.1\_Rst\_DUF4238\_4 | Rst\_DUF4238 | Rst\_DUF4238 | NC\_012982.1\_2416 | NC\_012982.1\_2416 | NC\_012982.1\_2416 | 1 | Rst\_DUF4238\_\_DUF4238\_Pers |

## 2.7 NC\_014008

##### 2.7.1 Host Genome

**GTDB-Tk taxonomy**:

classification

d\_\_Bacteria;p\_\_Verrucomicrobiota;c\_\_Verrucomicrobiae;o\_\_Opitutales;f\_\_Coraliomargaritaceae;g\_\_Coraliomargarita;s\_\_Coraliomargarita
akajimensis

**CheckM2 Summary**:

| total\_contigs | contig\_n50 | completeness | contamination |
| --- | --- | --- | --- |
| 1 | 3750771 | 100 | 0.02 |

**Defense-Finder Systems**:

| sys\_id | type | subtype | sys\_beg | sys\_end | protein\_in\_syst | genes\_count | name\_of\_profiles\_in\_sys |
| --- | --- | --- | --- | --- | --- | --- | --- |
| NC\_014008.1\_RM\_Type\_I\_6 | RM | RM\_Type\_I | NC\_014008.1\_1177 | NC\_014008.1\_1181 | NC\_014008.1\_1177,NC\_014008.1\_1180,NC\_014008.1\_1181 | 3 | RM\_\_Type\_I\_MTases\_FAM\_3,RM\_\_Type\_I\_REases\_FAM\_1.einsi\_trimmed,RM\_\_Type\_I\_S\_52 |
| NC\_014008.1\_Viperin\_3 | Viperin | Viperin | NC\_014008.1\_1182 | NC\_014008.1\_1182 | NC\_014008.1\_1182 | 1 | Viperin\_\_pVip |
| NC\_014008.1\_dGTPase\_4 | dGTPase | dGTPase | NC\_014008.1\_1769 | NC\_014008.1\_1769 | NC\_014008.1\_1769 | 1 | dGTPase\_\_Sp\_dGTPase |
| NC\_014008.1\_Ceres\_1 | Ceres | Ceres | NC\_014008.1\_2641 | NC\_014008.1\_2641 | NC\_014008.1\_2641 | 1 | Ceres\_\_CrsA1 |
| NC\_014008.1\_Gao\_Mza\_2 | Gao\_Mza | Gao\_Mza | NC\_014008.1\_2741 | NC\_014008.1\_2744 | NC\_014008.1\_2741,NC\_014008.1\_2743,NC\_014008.1\_2744 | 3 | Gao\_Mza\_\_MzaB,Gao\_Mza\_\_MzaC,Gao\_Mza\_\_MzaE |

## 2.8 NC\_014168

##### 2.8.1 Host Genome

**GTDB-Tk taxonomy**:

classification

d\_\_Bacteria;p\_\_Actinomycetota;c\_\_Actinomycetes;o\_\_Mycobacteriales;f\_\_Mycobacteriaceae;g\_\_Segniliparus;s\_\_Segniliparus
rotundus

**CheckM2 Summary**:

| total\_contigs | contig\_n50 | completeness | contamination |
| --- | --- | --- | --- |
| 1 | 3157527 | 99.99 | 0.03 |

**Defense-Finder Systems**:

| sys\_id | type | subtype | sys\_beg | sys\_end | protein\_in\_syst | genes\_count | name\_of\_profiles\_in\_sys |
| --- | --- | --- | --- | --- | --- | --- | --- |
| NC\_014168.1\_RM\_Type\_II\_2 | RM | RM\_Type\_II | NC\_014168.1\_1355 | NC\_014168.1\_1356 | NC\_014168.1\_1355,NC\_014168.1\_1356 | 2 | RM\_Type\_II\_\_Type\_II\_MTases\_FAM\_41,RM\_Type\_II\_\_Type\_II\_REase25 |
| NC\_014168.1\_RM\_Type\_IIG\_3 | RM | RM\_Type\_IIG | NC\_014168.1\_1738 | NC\_014168.1\_1738 | NC\_014168.1\_1738 | 1 | RM\_Type\_IIG\_\_Type\_IIG\_FAM\_2.einsi\_trimmed |
| NC\_014168.1\_DS-20\_1 | DS-20 | DS-20 | NC\_014168.1\_2356 | NC\_014168.1\_2356 | NC\_014168.1\_2356 | 1 | DS-20\_\_DS-20 |

**Genomad and CheckV Summary**:

| seq\_name | length | gene\_count | viral\_genes | checkv\_quality | miuvig\_quality | taxonomy | topology | coordinates |
| --- | --- | --- | --- | --- | --- | --- | --- | --- |
| NC\_014168.1\|provirus\_27479\_67296 | 39818 | 60 | 29 | Medium-quality | Genome-fragment | Viruses;Duplodnaviria;Heunggongvirae;Uroviricota;Caudoviricetes;; | Provirus | 27479-67296 |
| NC\_014168.1\|provirus\_1141631\_1162952 | 21322 | 36 | 19 | Medium-quality | Genome-fragment | Viruses;Duplodnaviria;Heunggongvirae;Uroviricota;Caudoviricetes;; | Provirus | 1141631-1162952 |

**Host Genome Ideogram with Phages**:

In
this circular plot, **“c”** indicates the contig, and the
number that follows (e.g., **c1**) represents the contig
number. If multiple contigs are present in the genome, each will be
shown with a distinct label (e.g., **c1**,
**c2**, etc.).

##### 2.8.2 Prophages

**Select prophage to show:**

###### 2.8.2.1 Phage ID: NC\_014168.1|provirus\_27479\_67296

**Phage-Host Genome Ideogram:**

**Genes Annotation (geNomad):**

| gene | length | marker | annotation\_accessions | annotation\_description |
| --- | --- | --- | --- | --- |
| NC\_014168.1\|provirus\_27479\_67296\_28 | 897 | NA | NA | NA |
| NC\_014168.1\|provirus\_27479\_67296\_29 | 549 | NA | NA | NA |
| NC\_014168.1\|provirus\_27479\_67296\_30 | 198 | NA | NA | NA |
| NC\_014168.1\|provirus\_27479\_67296\_31 | 753 | GENOMAD.032144.VV | PF12684 | NA |
| NC\_014168.1\|provirus\_27479\_67296\_32 | 657 | GENOMAD.124252.VP | PF03837;TIGR01913 | phage recombination protein Bet |
| NC\_014168.1\|provirus\_27479\_67296\_33 | 402 | NA | NA | NA |
| NC\_014168.1\|provirus\_27479\_67296\_34 | 354 | NA | NA | NA |
| NC\_014168.1\|provirus\_27479\_67296\_35 | 153 | NA | NA | NA |
| NC\_014168.1\|provirus\_27479\_67296\_36 | 420 | NA | NA | NA |
| NC\_014168.1\|provirus\_27479\_67296\_37 | 516 | GENOMAD.033304.VV | PF13395 | NA |
| NC\_014168.1\|provirus\_27479\_67296\_38 | 936 | GENOMAD.028141.VV | PF09681;TIGR01714 | N-terminal phage replisome organiser (Phage\_rep\_org\_N) |
| NC\_014168.1\|provirus\_27479\_67296\_39 | 366 | NA | NA | NA |
| NC\_014168.1\|provirus\_27479\_67296\_40 | 387 | GENOMAD.018151.VV | PF01870;COG1591 | Holliday junction resolvase, archaeal type |
| NC\_014168.1\|provirus\_27479\_67296\_41 | 201 | NA | NA | NA |
| NC\_014168.1\|provirus\_27479\_67296\_42 | 312 | NA | NA | NA |
| NC\_014168.1\|provirus\_27479\_67296\_43 | 213 | NA | NA | NA |
| NC\_014168.1\|provirus\_27479\_67296\_44 | 237 | NA | NA | NA |
| NC\_014168.1\|provirus\_27479\_67296\_45 | 321 | NA | NA | NA |
| NC\_014168.1\|provirus\_27479\_67296\_46 | 471 | NA | NA | NA |
| NC\_014168.1\|provirus\_27479\_67296\_47 | 429 | NA | NA | NA |
| NC\_014168.1\|provirus\_27479\_67296\_48 | 579 | NA | NA | NA |
| NC\_014168.1\|provirus\_27479\_67296\_49 | 312 | NA | NA | NA |
| NC\_014168.1\|provirus\_27479\_67296\_50 | 780 | NA | NA | NA |
| NC\_014168.1\|provirus\_27479\_67296\_51 | 222 | NA | NA | NA |
| NC\_014168.1\|provirus\_27479\_67296\_52 | 1761 | NA | NA | NA |
| NC\_014168.1\|provirus\_27479\_67296\_53 | 657 | NA | NA | NA |
| NC\_014168.1\|provirus\_27479\_67296\_54 | 321 | NA | NA | NA |
| NC\_014168.1\|provirus\_27479\_67296\_55 | 255 | GENOMAD.083226.VV | PF23850 | NA |
| NC\_014168.1\|provirus\_27479\_67296\_56 | 384 | NA | NA | NA |
| NC\_014168.1\|provirus\_27479\_67296\_57 | 480 | NA | NA | NA |
| NC\_014168.1\|provirus\_27479\_67296\_58 | 369 | GENOMAD.215374.PC | PF13827 | NA |
| NC\_014168.1\|provirus\_27479\_67296\_59 | 279 | GENOMAD.014408.VV | PF13395;TIGR02646 | HNH endonuclease |
| NC\_014168.1\|provirus\_27479\_67296\_60 | 186 | NA | NA | NA |
| NC\_014168.1\|provirus\_27479\_67296\_61 | 264 | GENOMAD.047538.VV | NA | NA |
| NC\_014168.1\|provirus\_27479\_67296\_62 | 1416 | GENOMAD.002060.VV | PF20441;PF04466;COG4626;TIGR01547;K06909 | Phage terminase-like protein, large subunit, contains N-terminal HTH domain |
| NC\_014168.1\|provirus\_27479\_67296\_63 | 1377 | GENOMAD.080886.VV | PF05133;TIGR01538 | phage portal protein, SPP1 family |
| NC\_014168.1\|provirus\_27479\_67296\_64 | 684 | GENOMAD.097054.VV | PF04233;TIGR01641 | phage putative head morphogenesis protein, SPP1 gp7 family |
| NC\_014168.1\|provirus\_27479\_67296\_65 | 477 | GENOMAD.040462.VV | NA | NA |
| NC\_014168.1\|provirus\_27479\_67296\_66 | 909 | GENOMAD.013023.VV | PF05065;TIGR01554;COG4653 | phage major capsid protein, HK97 family |
| NC\_014168.1\|provirus\_27479\_67296\_67 | 174 | NA | NA | NA |
| NC\_014168.1\|provirus\_27479\_67296\_68 | 468 | GENOMAD.062537.VV | PF09355 | Phage protein Gp19/Gp15/Gp42 |
| NC\_014168.1\|provirus\_27479\_67296\_69 | 342 | GENOMAD.038727.VV | PF12206;COG5614;TIGR01563 | Minor capsid protein |
| NC\_014168.1\|provirus\_27479\_67296\_70 | 309 | GENOMAD.087122.VV | NA | NA |
| NC\_014168.1\|provirus\_27479\_67296\_71 | 399 | NA | NA | NA |
| NC\_014168.1\|provirus\_27479\_67296\_72 | 648 | GENOMAD.004252.VV | NA | NA |
| NC\_014168.1\|provirus\_27479\_67296\_73 | 333 | NA | NA | NA |
| NC\_014168.1\|provirus\_27479\_67296\_74 | 384 | GENOMAD.056037.VV | PF17318;PF08765;TIGR02417 | Mor transcription activator family |
| NC\_014168.1\|provirus\_27479\_67296\_75 | 3723 | GENOMAD.016295.VV | COG3953 | SLT domain protein |
| NC\_014168.1\|provirus\_27479\_67296\_76 | 1263 | GENOMAD.053786.VV | COG4722 | Phage-related protein |
| NC\_014168.1\|provirus\_27479\_67296\_77 | 1773 | GENOMAD.019112.VV | NA | NA |
| NC\_014168.1\|provirus\_27479\_67296\_78 | 252 | NA | NA | NA |
| NC\_014168.1\|provirus\_27479\_67296\_79 | 1062 | GENOMAD.059914.VV | NA | NA |
| NC\_014168.1\|provirus\_27479\_67296\_80 | 1266 | GENOMAD.079434.VV | NA | NA |
| NC\_014168.1\|provirus\_27479\_67296\_81 | 1260 | GENOMAD.047467.VV | COG5632 | N-acetylmuramoyl-L-alanine amidase CwlA |
| NC\_014168.1\|provirus\_27479\_67296\_82 | 357 | NA | NA | NA |
| NC\_014168.1\|provirus\_27479\_67296\_83 | 276 | NA | NA | NA |
| NC\_014168.1\|provirus\_27479\_67296\_84 | 207 | GENOMAD.158505.VV | PF24596 | NA |
| NC\_014168.1\|provirus\_27479\_67296\_85 | 360 | GENOMAD.073706.VV | PF23778 | Putative phage holin family 2 |
| NC\_014168.1\|provirus\_27479\_67296\_86 | 372 | NA | NA | NA |
| NC\_014168.1\|provirus\_27479\_67296\_87 | 336 | NA | NA | NA |
| NC\_014168.1\|provirus\_27479\_67296\_88 | 273 | NA | NA | NA |
| NC\_014168.1\|provirus\_27479\_67296\_89 | 1263 | NA | NA | NA |
| NC\_014168.1\|provirus\_27479\_67296\_90 | 1050 | NA | NA | NA |

###### 2.8.2.2 Phage ID: NC\_014168.1|provirus\_1141631\_1162952

**Phage-Host Genome Ideogram:**

**Genes Annotation (geNomad):**

| gene | length | marker | annotation\_accessions | annotation\_description |
| --- | --- | --- | --- | --- |
| NC\_014168.1\|provirus\_1141631\_1162952\_1152 | 318 | NA | NA | NA |
| NC\_014168.1\|provirus\_1141631\_1162952\_1153 | 360 | GENOMAD.082050.VV | NA | NA |
| NC\_014168.1\|provirus\_1141631\_1162952\_1154 | 333 | GENOMAD.076859.VV | NA | NA |
| NC\_014168.1\|provirus\_1141631\_1162952\_1155 | 312 | GENOMAD.103239.VV | PF10665;TIGR01563;COG5614 | Minor capsid protein |
| NC\_014168.1\|provirus\_1141631\_1162952\_1156 | 237 | NA | NA | NA |
| NC\_014168.1\|provirus\_1141631\_1162952\_1157 | 510 | GENOMAD.028700.VV | TIGR02215 | phage conserved hypothetical protein, phiE125 gp8 family |
| NC\_014168.1\|provirus\_1141631\_1162952\_1158 | 192 | NA | NA | NA |
| NC\_014168.1\|provirus\_1141631\_1162952\_1159 | 1281 | GENOMAD.098207.VP | PF06673;TIGR01554;COG1659 | Lactococcus lactis bacteriophage major capsid protein |
| NC\_014168.1\|provirus\_1141631\_1162952\_1160 | 693 | GENOMAD.207278.VV | PF04586;COG3740;K06904;TIGR01543 | Phage head maturation protease |
| NC\_014168.1\|provirus\_1141631\_1162952\_1161 | 1302 | GENOMAD.138175.VV | PF04860;TIGR01537;COG4695 | phage portal protein, HK97 family |
| NC\_014168.1\|provirus\_1141631\_1162952\_1162 | 1497 | GENOMAD.037086.VV | PF20441;PF03354;COG4626;TIGR01547 | Phage terminase-like protein, large subunit, contains N-terminal HTH domain |
| NC\_014168.1\|provirus\_1141631\_1162952\_1163 | 291 | GENOMAD.064187.VV | TIGR01558 | NA |
| NC\_014168.1\|provirus\_1141631\_1162952\_1164 | 507 | NA | NA | NA |
| NC\_014168.1\|provirus\_1141631\_1162952\_1165 | 552 | NA | NA | NA |
| NC\_014168.1\|provirus\_1141631\_1162952\_1166 | 282 | NA | NA | NA |
| NC\_014168.1\|provirus\_1141631\_1162952\_1167 | 276 | GENOMAD.041582.VV | PF13395;TIGR02646 | TIGR02646 family protein |
| NC\_014168.1\|provirus\_1141631\_1162952\_1168 | 1119 | GENOMAD.064801.VV | PF00145;TIGR00675;COG0270;K15336 | DNA-methyltransferase (dcm) |
| NC\_014168.1\|provirus\_1141631\_1162952\_1169 | 657 | GENOMAD.066690.VV | PF10122;COG5349;TIGR01206 | Uncharacterized conserved protein, DUF983 family |
| NC\_014168.1\|provirus\_1141631\_1162952\_1170 | 249 | NA | NA | NA |
| NC\_014168.1\|provirus\_1141631\_1162952\_1171 | 258 | NA | NA | NA |
| NC\_014168.1\|provirus\_1141631\_1162952\_1172 | 378 | GENOMAD.063447.VV | PF11417 | NA |
| NC\_014168.1\|provirus\_1141631\_1162952\_1173 | 795 | NA | NA | NA |
| NC\_014168.1\|provirus\_1141631\_1162952\_1174 | 426 | NA | NA | NA |
| NC\_014168.1\|provirus\_1141631\_1162952\_1175 | 387 | NA | NA | NA |
| NC\_014168.1\|provirus\_1141631\_1162952\_1176 | 783 | GENOMAD.021092.VV | PF12684;TIGR00372;COG1468;K07465 | CRISPR-associated protein Cas4 |
| NC\_014168.1\|provirus\_1141631\_1162952\_1177 | 291 | NA | NA | NA |
| NC\_014168.1\|provirus\_1141631\_1162952\_1178 | 228 | NA | NA | NA |
| NC\_014168.1\|provirus\_1141631\_1162952\_1179 | 375 | GENOMAD.067886.VV | NA | NA |
| NC\_014168.1\|provirus\_1141631\_1162952\_1180 | 843 | GENOMAD.161686.VP | PF20188 | NA |
| NC\_014168.1\|provirus\_1141631\_1162952\_1181 | 921 | NA | NA | NA |
| NC\_014168.1\|provirus\_1141631\_1162952\_1182 | 255 | GENOMAD.222641.VP | PF08667;COG5606;TIGR02612;K07727 | Predicted DNA-binding protein, XRE-type HTH domain |
| NC\_014168.1\|provirus\_1141631\_1162952\_1183 | 429 | NA | NA | NA |
| NC\_014168.1\|provirus\_1141631\_1162952\_1184 | 861 | NA | NA | NA |
| NC\_014168.1\|provirus\_1141631\_1162952\_1185 | 264 | NA | NA | NA |
| NC\_014168.1\|provirus\_1141631\_1162952\_1186 | 1356 | NA | NA | NA |

##### 2.8.3 vOTUs

When more than 1 phage is detected in the host genome, we perform a
clustering step using the tool dRep.

A threshold of 0.95 was applied to the ANI similarity index to define
clusters of virus operational taxonomic units (vOTUs).

**Final cluster designations**

| genomad\_summary$seq\_name | genome | secondary\_cluster | threshold | cluster\_method | comparison\_algorithm | primary\_cluster |
| --- | --- | --- | --- | --- | --- | --- |
| NC\_014168.1\|provirus\_27479\_67296 | sequence\_000000.fasta.fasta | 1\_0 | 0.05 | average | ANImf | 1 |
| NC\_014168.1\|provirus\_1141631\_1162952 | sequence\_000001.fasta.fasta | 2\_0 | 0.05 | average | ANImf | 2 |

**Primary clustering plot**

##### 2.8.4 Prophage-Host Prediction

Prediction of potential hosts for the prophage contigs.

| virus | host\_genome | host\_taxonomy | main\_method | confidence\_score | additional\_methods |
| --- | --- | --- | --- | --- | --- |
| NC\_014168.1\|provirus\_1141631\_1162952 | RS\_GCF\_000092825.1 | d\_\_Bacteria;p\_\_Actinomycetota;c\_\_Actinomycetia;o\_\_Mycobacteriales;f\_\_Mycobacteriaceae;g\_\_Segniliparus;s\_\_Segniliparus rotundus | blast | 95.7 | iPHoP-RF;68.00 |
| NC\_014168.1\|provirus\_27479\_67296 | RS\_GCF\_000092825.1 | d\_\_Bacteria;p\_\_Actinomycetota;c\_\_Actinomycetia;o\_\_Mycobacteriales;f\_\_Mycobacteriaceae;g\_\_Segniliparus;s\_\_Segniliparus rotundus | blast | 96.0 | iPHoP-RF;79.30 |

##### 2.8.5 AMG Predictions

Prediction of auxiliary metabolic genes in the prophage contigs.

| protein | scaffold | amg\_ko | amg\_ko\_name | pfam | pfam\_name |
| --- | --- | --- | --- | --- | --- |
| NC\_014168.1\|provirus\_1141631\_1162952\_17 | NC\_014168.1\|provirus\_1141631\_1162952 | K00558 | DNMT1, dcm; DNA (cytosine-5)-methyltransferase 1 [EC:2.1.1.37] | PF00145.17 | C-5 cytosine-specific DNA methylase |

## 2.9 NC\_014211

##### 2.9.1 Host Genome

**GTDB-Tk taxonomy**:

classification

d\_\_Bacteria;p\_\_Actinomycetota;c\_\_Actinomycetes;o\_\_Streptosporangiales;f\_\_Streptosporangiaceae;g\_\_Nocardiopsis;s\_\_Nocardiopsis
dassonvillei

**CheckM2 Summary**:

| total\_contigs | contig\_n50 | completeness | contamination |
| --- | --- | --- | --- |
| 1 | 775354 | 14.07 | 0.01 |

No Defense-Finder systems detected.

## 2.10 NC\_014212

##### 2.10.1 Host Genome

**GTDB-Tk taxonomy**:

classification

d\_\_Bacteria;p\_\_Deinococcota;c\_\_Deinococci;o\_\_Deinococcales;f\_\_Thermaceae;g\_\_Allomeiothermus;s\_\_Allomeiothermus
silvanus

**CheckM2 Summary**:

| total\_contigs | contig\_n50 | completeness | contamination |
| --- | --- | --- | --- |
| 1 | 3249394 | 99.99 | 0.16 |

**Defense-Finder Systems**:

| sys\_id | type | subtype | sys\_beg | sys\_end | protein\_in\_syst | genes\_count | name\_of\_profiles\_in\_sys |
| --- | --- | --- | --- | --- | --- | --- | --- |
| NC\_014212.1\_Ceres\_1 | Ceres | Ceres | NC\_014212.1\_142 | NC\_014212.1\_142 | NC\_014212.1\_142 | 1 | Ceres\_\_CrsA1 |
| NC\_014212.1\_RM\_Type\_III\_2 | RM | RM\_Type\_III | NC\_014212.1\_571 | NC\_014212.1\_572 | NC\_014212.1\_571,NC\_014212.1\_572 | 2 | RM\_Type\_III\_\_Type\_III\_MTases\_FAM\_0,RM\_Type\_III\_\_Type\_III\_REases\_FAM\_0.einsi\_trimmed |
| NC\_014212.1\_CAS\_Class1-Subtype-III-A\_4 | Cas | CAS\_Class1-Subtype-III-A | NC\_014212.1\_712 | NC\_014212.1\_723 | NC\_014212.1\_712,NC\_014212.1\_713,NC\_014212.1\_714,NC\_014212.1\_715,NC\_014212.1\_716,NC\_014212.1\_719,NC\_014212.1\_720,NC\_014212.1\_721,NC\_014212.1\_722,NC\_014212.1\_723 | 10 | cas10\_III-A\_1,cas1\_I\_II\_III\_IV\_V\_VI\_1,cas2\_I\_II\_III\_IV\_V\_VI\_8,cas6\_I\_II\_III\_IV\_V\_VI\_15,csm2gr11\_III-A\_1,csm3gr7\_III-A\_1,csm4gr5\_III-A\_2,csm5gr7\_III-A\_2,csx1\_III\_21,csx1\_III\_21 |
| NC\_014212.1\_CAS\_Class1-Subtype-I-E\_3 | Cas | CAS\_Class1-Subtype-I-E | NC\_014212.1\_1246 | NC\_014212.1\_1255 | NC\_014212.1\_1246,NC\_014212.1\_1248,NC\_014212.1\_1249,NC\_014212.1\_1251,NC\_014212.1\_1252,NC\_014212.1\_1253,NC\_014212.1\_1254,NC\_014212.1\_1255 | 8 | cas1\_I-E\_1,cas2\_I-E\_1,cas3\_I\_2,cas5\_I-E\_2,cas6e\_I\_II\_III\_IV\_V\_VI\_2,cas7\_I-E\_16,cas8e\_I-E\_5,cse2gr11\_I-E\_8 |

**Genomad and CheckV Summary**:

| seq\_name | length | gene\_count | viral\_genes | checkv\_quality | miuvig\_quality | taxonomy | topology | coordinates |
| --- | --- | --- | --- | --- | --- | --- | --- | --- |
| NC\_014212.1\|provirus\_1746722\_1764354 | 17633 | 38 | 3 | Medium-quality | Genome-fragment | Viruses;Varidnaviria;Helvetiavirae;Dividoviricota;Laserviricetes;Halopanivirales;Matsushitaviridae | Provirus | 1746722-1764354 |
| NC\_014212.1\|provirus\_1892278\_1921074 | 28797 | 40 | 6 | Medium-quality | Genome-fragment | Viruses;Duplodnaviria;Heunggongvirae;Uroviricota;Caudoviricetes;; | Provirus | 1892278-1921074 |
| NC\_014212.1\|provirus\_1170297\_1211081 | 40785 | 50 | 10 | Medium-quality | Genome-fragment | Viruses;Duplodnaviria;Heunggongvirae;Uroviricota;Caudoviricetes;; | Provirus | 1170297-1211081 |

**Host Genome Ideogram with Phages**:

In
this circular plot, **“c”** indicates the contig, and the
number that follows (e.g., **c1**) represents the contig
number. If multiple contigs are present in the genome, each will be
shown with a distinct label (e.g., **c1**,
**c2**, etc.).

##### 2.10.2 Prophages

**Select prophage to show:**

###### 2.10.2.1 Phage ID: NC\_014212.1|provirus\_1746722\_1764354

**Phage-Host Genome Ideogram:**

**Genes Annotation (geNomad):**

| gene | length | marker | annotation\_accessions | annotation\_description |
| --- | --- | --- | --- | --- |
| NC\_014212.1\|provirus\_1746722\_1764354\_1776 | 981 | NA | NA | NA |
| NC\_014212.1\|provirus\_1746722\_1764354\_1777 | 570 | NA | NA | NA |
| NC\_014212.1\|provirus\_1746722\_1764354\_1778 | 708 | NA | NA | NA |
| NC\_014212.1\|provirus\_1746722\_1764354\_1779 | 189 | NA | NA | NA |
| NC\_014212.1\|provirus\_1746722\_1764354\_1780 | 207 | NA | NA | NA |
| NC\_014212.1\|provirus\_1746722\_1764354\_1781 | 171 | NA | NA | NA |
| NC\_014212.1\|provirus\_1746722\_1764354\_1782 | 279 | NA | NA | NA |
| NC\_014212.1\|provirus\_1746722\_1764354\_1783 | 123 | NA | NA | NA |
| NC\_014212.1\|provirus\_1746722\_1764354\_1784 | 93 | NA | NA | NA |
| NC\_014212.1\|provirus\_1746722\_1764354\_1785 | 300 | NA | NA | NA |
| NC\_014212.1\|provirus\_1746722\_1764354\_1786 | 345 | GENOMAD.138277.VP | NA | NA |
| NC\_014212.1\|provirus\_1746722\_1764354\_1787 | 333 | NA | NA | NA |
| NC\_014212.1\|provirus\_1746722\_1764354\_1788 | 1467 | NA | NA | NA |
| NC\_014212.1\|provirus\_1746722\_1764354\_1789 | 210 | GENOMAD.118491.VV | NA | NA |
| NC\_014212.1\|provirus\_1746722\_1764354\_1790 | 333 | NA | NA | NA |
| NC\_014212.1\|provirus\_1746722\_1764354\_1791 | 243 | NA | NA | NA |
| NC\_014212.1\|provirus\_1746722\_1764354\_1792 | 270 | NA | NA | NA |
| NC\_014212.1\|provirus\_1746722\_1764354\_1793 | 204 | NA | NA | NA |
| NC\_014212.1\|provirus\_1746722\_1764354\_1794 | 471 | NA | NA | NA |
| NC\_014212.1\|provirus\_1746722\_1764354\_1795 | 366 | NA | NA | NA |
| NC\_014212.1\|provirus\_1746722\_1764354\_1796 | 678 | NA | NA | NA |
| NC\_014212.1\|provirus\_1746722\_1764354\_1797 | 201 | GENOMAD.187456.VV | NA | NA |
| NC\_014212.1\|provirus\_1746722\_1764354\_1798 | 738 | NA | NA | NA |
| NC\_014212.1\|provirus\_1746722\_1764354\_1799 | 432 | GENOMAD.226936.VP | NA | NA |
| NC\_014212.1\|provirus\_1746722\_1764354\_1800 | 462 | NA | NA | NA |
| NC\_014212.1\|provirus\_1746722\_1764354\_1801 | 885 | GENOMAD.175220.VV | PF22210 | Large MCP VP17 central beta-barrel |
| NC\_014212.1\|provirus\_1746722\_1764354\_1802 | 243 | NA | NA | NA |
| NC\_014212.1\|provirus\_1746722\_1764354\_1803 | 636 | NA | NA | NA |
| NC\_014212.1\|provirus\_1746722\_1764354\_1804 | 300 | NA | NA | NA |
| NC\_014212.1\|provirus\_1746722\_1764354\_1805 | 177 | NA | NA | NA |
| NC\_014212.1\|provirus\_1746722\_1764354\_1806 | 507 | GENOMAD.201673.VV | NA | NA |
| NC\_014212.1\|provirus\_1746722\_1764354\_1807 | 189 | GENOMAD.131894.VV | PF04531;TIGR01598;COG5546 | NA |
| NC\_014212.1\|provirus\_1746722\_1764354\_1808 | 174 | NA | NA | NA |
| NC\_014212.1\|provirus\_1746722\_1764354\_1809 | 456 | GENOMAD.199413.VV | NA | NA |
| NC\_014212.1\|provirus\_1746722\_1764354\_1810 | 1050 | GENOMAD.107709.VV | NA | NA |
| NC\_014212.1\|provirus\_1746722\_1764354\_1811 | 1014 | GENOMAD.109373.VV | NA | NA |
| NC\_014212.1\|provirus\_1746722\_1764354\_1812 | 264 | NA | NA | NA |
| NC\_014212.1\|provirus\_1746722\_1764354\_1813 | 750 | NA | NA | NA |

###### 2.10.2.2 Phage ID: NC\_014212.1|provirus\_1892278\_1921074

**Phage-Host Genome Ideogram:**

**Genes Annotation (geNomad):**

| gene | length | marker | annotation\_accessions | annotation\_description |
| --- | --- | --- | --- | --- |
| NC\_014212.1\|provirus\_1892278\_1921074\_1934 | 438 | NA | NA | NA |
| NC\_014212.1\|provirus\_1892278\_1921074\_1935 | 312 | NA | NA | NA |
| NC\_014212.1\|provirus\_1892278\_1921074\_1936 | 240 | NA | NA | NA |
| NC\_014212.1\|provirus\_1892278\_1921074\_1937 | 231 | NA | NA | NA |
| NC\_014212.1\|provirus\_1892278\_1921074\_1938 | 276 | NA | NA | NA |
| NC\_014212.1\|provirus\_1892278\_1921074\_1939 | 219 | NA | NA | NA |
| NC\_014212.1\|provirus\_1892278\_1921074\_1940 | 396 | NA | NA | NA |
| NC\_014212.1\|provirus\_1892278\_1921074\_1941 | 480 | NA | NA | NA |
| NC\_014212.1\|provirus\_1892278\_1921074\_1942 | 702 | NA | NA | NA |
| NC\_014212.1\|provirus\_1892278\_1921074\_1943 | 441 | NA | NA | NA |
| NC\_014212.1\|provirus\_1892278\_1921074\_1944 | 249 | NA | NA | NA |
| NC\_014212.1\|provirus\_1892278\_1921074\_1945 | 342 | GENOMAD.188989.VV | PF10711 | Hypothetical protein (DUF2513) |
| NC\_014212.1\|provirus\_1892278\_1921074\_1946 | 321 | NA | NA | NA |
| NC\_014212.1\|provirus\_1892278\_1921074\_1947 | 462 | GENOMAD.102149.VV | PF11985 | Bacteriophage Mu, Gp27 |
| NC\_014212.1\|provirus\_1892278\_1921074\_1948 | 1284 | GENOMAD.025283.VV | NA | NA |
| NC\_014212.1\|provirus\_1892278\_1921074\_1949 | 1536 | GENOMAD.116653.VV | PF06074;COG4383 | Mu-like prophage protein gp29 |
| NC\_014212.1\|provirus\_1892278\_1921074\_1950 | 1173 | GENOMAD.169564.VP | COG2369 | Uncharacterized conserved protein, contains phage Mu gpF-like domain |
| NC\_014212.1\|provirus\_1892278\_1921074\_1951 | 1200 | NA | NA | NA |
| NC\_014212.1\|provirus\_1892278\_1921074\_1952 | 453 | NA | NA | NA |
| NC\_014212.1\|provirus\_1892278\_1921074\_1953 | 540 | NA | NA | NA |
| NC\_014212.1\|provirus\_1892278\_1921074\_1954 | 447 | GENOMAD.191448.VP | PF05069;TIGR01635;COG5005 | phage virion morphogenesis (putative tail completion) protein |
| NC\_014212.1\|provirus\_1892278\_1921074\_1955 | 501 | NA | NA | NA |
| NC\_014212.1\|provirus\_1892278\_1921074\_1956 | 213 | NA | NA | NA |
| NC\_014212.1\|provirus\_1892278\_1921074\_1957 | 1422 | GENOMAD.041181.VV | PF10758;COG3497 | Phage tail sheath protein FI |
| NC\_014212.1\|provirus\_1892278\_1921074\_1958 | 432 | GENOMAD.115837.VV | PF10772 | Bacteriophage HP1, Orf24 |
| NC\_014212.1\|provirus\_1892278\_1921074\_1959 | 375 | NA | NA | NA |
| NC\_014212.1\|provirus\_1892278\_1921074\_1960 | 132 | NA | NA | NA |
| NC\_014212.1\|provirus\_1892278\_1921074\_1961 | 2082 | GENOMAD.106657.VP | COG5280 | Phage-related minor tail protein |
| NC\_014212.1\|provirus\_1892278\_1921074\_1962 | 579 | GENOMAD.163406.VV | NA | NA |
| NC\_014212.1\|provirus\_1892278\_1921074\_1963 | 666 | GENOMAD.068883.VV | TIGR03361;COG3500 | type VI secretion system Vgr family protein |
| NC\_014212.1\|provirus\_1892278\_1921074\_1964 | 477 | GENOMAD.068883.VV | TIGR03361;COG3500 | type VI secretion system Vgr family protein |
| NC\_014212.1\|provirus\_1892278\_1921074\_1965 | 375 | NA | NA | NA |
| NC\_014212.1\|provirus\_1892278\_1921074\_1966 | 1083 | GENOMAD.106861.VV | PF04865;COG3299 | Baseplate J-like protein |
| NC\_014212.1\|provirus\_1892278\_1921074\_1967 | 612 | GENOMAD.067612.VV | PF10076;COG4385;TIGR01634 | Bacteriophage P2-related tail formation protein |
| NC\_014212.1\|provirus\_1892278\_1921074\_1968 | 1155 | NA | NA | NA |
| NC\_014212.1\|provirus\_1892278\_1921074\_1969 | 177 | NA | NA | NA |
| NC\_014212.1\|provirus\_1892278\_1921074\_1970 | 501 | NA | NA | NA |
| NC\_014212.1\|provirus\_1892278\_1921074\_1971 | 1002 | NA | NA | NA |
| NC\_014212.1\|provirus\_1892278\_1921074\_1972 | 282 | NA | NA | NA |
| NC\_014212.1\|provirus\_1892278\_1921074\_1973 | 1392 | NA | NA | NA |
| NC\_014212.1\|provirus\_1892278\_1921074\_1974 | 2181 | GENOMAD.090865.VV | PF13086;TIGR02785;COG1074;K19781 | helicase-exonuclease AddAB, AddA subunit, Firmicutes type |

###### 2.10.2.3 Phage ID: NC\_014212.1|provirus\_1170297\_1211081

**Phage-Host Genome Ideogram:**

**Genes Annotation (geNomad):**

| gene | length | marker | annotation\_accessions | annotation\_description |
| --- | --- | --- | --- | --- |
| NC\_014212.1\|provirus\_1170297\_1211081\_1203 | 1164 | NA | NA | NA |
| NC\_014212.1\|provirus\_1170297\_1211081\_1204 | 261 | NA | NA | NA |
| NC\_014212.1\|provirus\_1170297\_1211081\_1205 | 1200 | NA | NA | NA |
| NC\_014212.1\|provirus\_1170297\_1211081\_1206 | 453 | NA | NA | NA |
| NC\_014212.1\|provirus\_1170297\_1211081\_1207 | 654 | GENOMAD.212789.VC | PF08667;TIGR00673;COG1974;K22299 | cyanase |
| NC\_014212.1\|provirus\_1170297\_1211081\_1208 | 180 | NA | NA | NA |
| NC\_014212.1\|provirus\_1170297\_1211081\_1209 | 231 | NA | NA | NA |
| NC\_014212.1\|provirus\_1170297\_1211081\_1210 | 189 | NA | NA | NA |
| NC\_014212.1\|provirus\_1170297\_1211081\_1211 | 273 | NA | NA | NA |
| NC\_014212.1\|provirus\_1170297\_1211081\_1212 | 123 | NA | NA | NA |
| NC\_014212.1\|provirus\_1170297\_1211081\_1213 | 219 | NA | NA | NA |
| NC\_014212.1\|provirus\_1170297\_1211081\_1214 | 618 | NA | NA | NA |
| NC\_014212.1\|provirus\_1170297\_1211081\_1215 | 1734 | NA | NA | NA |
| NC\_014212.1\|provirus\_1170297\_1211081\_1216 | 336 | GENOMAD.004598.VV | PF18743;COG1591 | REase\_AHJR-like |
| NC\_014212.1\|provirus\_1170297\_1211081\_1217 | 2694 | NA | NA | NA |
| NC\_014212.1\|provirus\_1170297\_1211081\_1218 | 243 | NA | NA | NA |
| NC\_014212.1\|provirus\_1170297\_1211081\_1219 | 837 | NA | NA | NA |
| NC\_014212.1\|provirus\_1170297\_1211081\_1220 | 468 | NA | NA | NA |
| NC\_014212.1\|provirus\_1170297\_1211081\_1221 | 456 | GENOMAD.172137.VP | PF07141;COG3728;TIGR02036 | Putative bacteriophage terminase small subunit |
| NC\_014212.1\|provirus\_1170297\_1211081\_1222 | 1305 | GENOMAD.190694.VP | PF04466;PF17288;TIGR01547;COG1783;K06909 | phage terminase, large subunit, PBSX family |
| NC\_014212.1\|provirus\_1170297\_1211081\_1223 | 1428 | NA | NA | NA |
| NC\_014212.1\|provirus\_1170297\_1211081\_1224 | 873 | GENOMAD.017886.VV | PF04233;TIGR01641 | phage putative head morphogenesis protein, SPP1 gp7 family |
| NC\_014212.1\|provirus\_1170297\_1211081\_1225 | 180 | NA | NA | NA |
| NC\_014212.1\|provirus\_1170297\_1211081\_1226 | 1644 | NA | NA | NA |
| NC\_014212.1\|provirus\_1170297\_1211081\_1227 | 201 | NA | NA | NA |
| NC\_014212.1\|provirus\_1170297\_1211081\_1228 | 873 | NA | NA | NA |
| NC\_014212.1\|provirus\_1170297\_1211081\_1229 | 660 | NA | NA | NA |
| NC\_014212.1\|provirus\_1170297\_1211081\_1230 | 1653 | GENOMAD.196129.VP | PF05125;TIGR01551;COG4653 | phage major capsid protein, P2 family |
| NC\_014212.1\|provirus\_1170297\_1211081\_1231 | 234 | NA | NA | NA |
| NC\_014212.1\|provirus\_1170297\_1211081\_1232 | 555 | NA | NA | NA |
| NC\_014212.1\|provirus\_1170297\_1211081\_1233 | 327 | NA | NA | NA |
| NC\_014212.1\|provirus\_1170297\_1211081\_1234 | 540 | NA | NA | NA |
| NC\_014212.1\|provirus\_1170297\_1211081\_1235 | 435 | NA | NA | NA |
| NC\_014212.1\|provirus\_1170297\_1211081\_1236 | 789 | NA | NA | NA |
| NC\_014212.1\|provirus\_1170297\_1211081\_1237 | 486 | NA | NA | NA |
| NC\_014212.1\|provirus\_1170297\_1211081\_1238 | 3738 | GENOMAD.183233.VP | NA | NA |
| NC\_014212.1\|provirus\_1170297\_1211081\_1239 | 441 | NA | NA | NA |
| NC\_014212.1\|provirus\_1170297\_1211081\_1240 | 1077 | NA | NA | NA |
| NC\_014212.1\|provirus\_1170297\_1211081\_1241 | 1905 | NA | NA | NA |
| NC\_014212.1\|provirus\_1170297\_1211081\_1242 | 543 | NA | NA | NA |
| NC\_014212.1\|provirus\_1170297\_1211081\_1243 | 540 | NA | NA | NA |
| NC\_014212.1\|provirus\_1170297\_1211081\_1244 | 312 | NA | NA | NA |
| NC\_014212.1\|provirus\_1170297\_1211081\_1245 | 1839 | NA | NA | NA |
| NC\_014212.1\|provirus\_1170297\_1211081\_1246 | 561 | NA | NA | NA |
| NC\_014212.1\|provirus\_1170297\_1211081\_1247 | 312 | GENOMAD.103334.VV | NA | NA |
| NC\_014212.1\|provirus\_1170297\_1211081\_1248 | 792 | GENOMAD.024392.VV | NA | NA |
| NC\_014212.1\|provirus\_1170297\_1211081\_1249 | 549 | GENOMAD.052331.VV | NA | NA |
| NC\_014212.1\|provirus\_1170297\_1211081\_1250 | 216 | GENOMAD.199026.VV | NA | NA |
| NC\_014212.1\|provirus\_1170297\_1211081\_1251 | 300 | NA | NA | NA |
| NC\_014212.1\|provirus\_1170297\_1211081\_1252 | 1002 | NA | NA | NA |

##### 2.10.3 vOTUs

When more than 1 phage is detected in the host genome, we perform a
clustering step using the tool dRep.

A threshold of 0.95 was applied to the ANI similarity index to define
clusters of virus operational taxonomic units (vOTUs).

**Final cluster designations**

| genomad\_summary$seq\_name | genome | secondary\_cluster | threshold | cluster\_method | comparison\_algorithm | primary\_cluster |
| --- | --- | --- | --- | --- | --- | --- |
| NC\_014212.1\|provirus\_1746722\_1764354 | sequence\_000000.fasta.fasta | 1\_0 | 0.05 | average | ANImf | 1 |
| NC\_014212.1\|provirus\_1892278\_1921074 | sequence\_000002.fasta.fasta | 2\_0 | 0.05 | average | ANImf | 2 |
| NC\_014212.1\|provirus\_1170297\_1211081 | sequence\_000001.fasta.fasta | 3\_0 | 0.05 | average | ANImf | 3 |

**Primary clustering plot**

##### 2.10.4 Prophage-Host Prediction

Prediction of potential hosts for the prophage contigs.

| virus | host\_genome | host\_taxonomy | main\_method | confidence\_score | additional\_methods |
| --- | --- | --- | --- | --- | --- |
| NC\_014212.1\|provirus\_1170297\_1211081 | RS\_GCF\_000092125.1 | d\_\_Bacteria;p\_\_Deinococcota;c\_\_Deinococci;o\_\_Deinococcales;f\_\_Thermaceae;g\_\_Meiothermus\_B;s\_\_Meiothermus\_B silvanus | blast | 95.9 | iPHoP-RF;76.80 |
| NC\_014212.1\|provirus\_1170297\_1211081 | RS\_GCF\_003226535.1 | d\_\_Bacteria;p\_\_Deinococcota;c\_\_Deinococci;o\_\_Deinococcales;f\_\_Thermaceae;g\_\_Meiothermus\_B;s\_\_Meiothermus\_B sp003226535 | blast | 92.4 | iPHoP-RF;74.60 |
| NC\_014212.1\|provirus\_1170297\_1211081 | RS\_GCF\_000430045.1 | d\_\_Bacteria;p\_\_Deinococcota;c\_\_Deinococci;o\_\_Deinococcales;f\_\_Thermaceae;g\_\_Calidithermus;s\_\_Calidithermus chliarophilus | blast | 90.7 | CRISPR;76.20 iPHoP-RF;71.10 |
| NC\_014212.1\|provirus\_1746722\_1764354 | RS\_GCF\_000092125.1 | d\_\_Bacteria;p\_\_Deinococcota;c\_\_Deinococci;o\_\_Deinococcales;f\_\_Thermaceae;g\_\_Meiothermus\_B;s\_\_Meiothermus\_B silvanus | blast | 95.2 | iPHoP-RF;86.00 |
| NC\_014212.1\|provirus\_1892278\_1921074 | RS\_GCF\_000092125.1 | d\_\_Bacteria;p\_\_Deinococcota;c\_\_Deinococci;o\_\_Deinococcales;f\_\_Thermaceae;g\_\_Meiothermus\_B;s\_\_Meiothermus\_B silvanus | blast | 94.5 | iPHoP-RF;74.80 |
| NC\_014212.1\|provirus\_1892278\_1921074 | RS\_GCF\_000430045.1 | d\_\_Bacteria;p\_\_Deinococcota;c\_\_Deinococci;o\_\_Deinococcales;f\_\_Thermaceae;g\_\_Calidithermus;s\_\_Calidithermus chliarophilus | blast | 92.9 | iPHoP-RF;63.50 |
| NC\_014212.1\|provirus\_1892278\_1921074 | RS\_GCF\_000373205.1 | d\_\_Bacteria;p\_\_Deinococcota;c\_\_Deinococci;o\_\_Deinococcales;f\_\_Thermaceae;g\_\_Calidithermus;s\_\_Calidithermus timidus | blast | 90.9 | None |
| NC\_014212.1\|provirus\_1892278\_1921074 | RS\_GCF\_003574095.1 | d\_\_Bacteria;p\_\_Deinococcota;c\_\_Deinococci;o\_\_Deinococcales;f\_\_Thermaceae;g\_\_Calidithermus;s\_\_Calidithermus roseus | blast | 90.6 | None |
| NC\_014212.1\|provirus\_1892278\_1921074 | RS\_GCF\_003574035.1 | d\_\_Bacteria;p\_\_Deinococcota;c\_\_Deinococci;o\_\_Deinococcales;f\_\_Thermaceae;g\_\_Meiothermus;s\_\_Meiothermus hypogaeus | blast | 90.5 | iPHoP-RF;59.80 |

##### 2.10.5 AMG Predictions

Prediction of auxiliary metabolic genes in the prophage contigs.

| protein | scaffold | amg\_ko | amg\_ko\_name | pfam | pfam\_name |
| --- | --- | --- | --- | --- | --- |
| NC\_014212.1\|provirus\_1892278\_1921074\_39 | NC\_014212.1\|provirus\_1892278\_1921074 | K01840 | manB; phosphomannomutase [EC:5.4.2.8] | PF02878.16 | Phosphoglucomutase/phosphomannomutase, alpha/beta/alpha domain I |

## 2.11 NC\_014363

##### 2.11.1 Host Genome

**GTDB-Tk taxonomy**:

classification

d\_\_Bacteria;p\_\_Actinomycetota;c\_\_Coriobacteriia;o\_\_Coriobacteriales;f\_\_Atopobiaceae;g\_\_Olsenella;s\_\_Olsenella
uli

**CheckM2 Summary**:

| total\_contigs | contig\_n50 | completeness | contamination |
| --- | --- | --- | --- |
| 1 | 2051896 | 99.7 | 0.35 |

**Defense-Finder Systems**:

| sys\_id | type | subtype | sys\_beg | sys\_end | protein\_in\_syst | genes\_count | name\_of\_profiles\_in\_sys |
| --- | --- | --- | --- | --- | --- | --- | --- |
| NC\_014363.1\_AbiE\_1 | AbiE | AbiE | NC\_014363.1\_207 | NC\_014363.1\_208 | NC\_014363.1\_207,NC\_014363.1\_208 | 2 | AbiEii\_\_AbiEi\_4,AbiEii\_\_AbiEii |
| NC\_014363.1\_AbiE\_2 | AbiE | AbiE | NC\_014363.1\_473 | NC\_014363.1\_474 | NC\_014363.1\_473,NC\_014363.1\_474 | 2 | AbiEii\_\_AbiEi\_4,AbiEii\_\_AbiEii |
| NC\_014363.1\_VP1839\_3 | VP1839 | VP1839 | NC\_014363.1\_1184 | NC\_014363.1\_1184 | NC\_014363.1\_1184 | 1 | VP1839\_\_VP1839 |
| NC\_014363.1\_RM\_Type\_II\_6 | RM | RM\_Type\_II | NC\_014363.1\_1186 | NC\_014363.1\_1187 | NC\_014363.1\_1186,NC\_014363.1\_1187 | 2 | RM\_Type\_II\_\_Type\_II\_MTases\_FAM\_0,RM\_Type\_II\_\_Type\_II\_REase15 |
| NC\_014363.1\_CAS\_Class2-Subtype-II-A\_7 | Cas | CAS\_Class2-Subtype-II-A | NC\_014363.1\_1233 | NC\_014363.1\_1236 | NC\_014363.1\_1233,NC\_014363.1\_1234,NC\_014363.1\_1235,NC\_014363.1\_1236 | 4 | cas1\_I\_II\_III\_IV\_V\_VI\_5,cas2\_I\_II\_III\_IV\_V\_VI\_6,cas9\_II-A\_1,csn2\_II-A\_3 |
| NC\_014363.1\_RM\_Type\_I\_5 | RM | RM\_Type\_I | NC\_014363.1\_1659 | NC\_014363.1\_1664 | NC\_014363.1\_1659,NC\_014363.1\_1662,NC\_014363.1\_1663,NC\_014363.1\_1664 | 4 | RM\_\_Type\_I\_MTases\_FAM\_2,RM\_\_Type\_I\_REases\_FAM\_2.einsi\_trimmed,RM\_\_Type\_I\_S\_04,RM\_\_Type\_I\_S\_04 |

## 2.12 NC\_014364

##### 2.12.1 Host Genome

**GTDB-Tk taxonomy**:

classification

d\_\_Bacteria;p\_\_Spirochaetota;c\_\_Spirochaetia;o\_\_DSM-16054;f\_\_Sediminispirochaetaceae;g\_\_Sediminispirochaeta;s\_\_Sediminispirochaeta
smaragdinae

**CheckM2 Summary**:

| total\_contigs | contig\_n50 | completeness | contamination |
| --- | --- | --- | --- |
| 1 | 4653970 | 99.98 | 1.88 |

**Defense-Finder Systems**:

| sys\_id | type | subtype | sys\_beg | sys\_end | protein\_in\_syst | genes\_count | name\_of\_profiles\_in\_sys |
| --- | --- | --- | --- | --- | --- | --- | --- |
| NC\_014364.1\_TIR-IV\_4 | TIR-IV | TIR-IV | NC\_014364.1\_392 | NC\_014364.1\_393 | NC\_014364.1\_392,NC\_014364.1\_393 | 2 | TIR-IV\_\_TIR-IV\_A,TIR-IV\_\_TIR-IV\_B |
| NC\_014364.1\_SoFic\_3 | SoFIC | SoFic | NC\_014364.1\_418 | NC\_014364.1\_418 | NC\_014364.1\_418 | 1 | SoFic\_\_SoFic |
| NC\_014364.1\_MazEF\_2 | MazEF | MazEF | NC\_014364.1\_697 | NC\_014364.1\_698 | NC\_014364.1\_697,NC\_014364.1\_698 | 2 | MazEF\_\_MazE,MazEF\_\_MazF |
| NC\_014364.1\_CAS\_Class1-Subtype-I-C\_5 | Cas | CAS\_Class1-Subtype-I-C | NC\_014364.1\_1062 | NC\_014364.1\_1069 | NC\_014364.1\_1062,NC\_014364.1\_1063,NC\_014364.1\_1064,NC\_014364.1\_1065,NC\_014364.1\_1066,NC\_014364.1\_1067,NC\_014364.1\_1068,NC\_014364.1\_1069 | 8 | WYL\_I\_II\_III\_IV\_V\_VI\_4,cas1\_I\_II\_III\_IV\_V\_VI\_6,cas2\_I\_II\_III\_IV\_V\_VI\_3,cas3\_I\_5,cas4\_I\_II\_III\_IV\_V\_VI\_6,cas5\_I-C\_11,cas7\_I-C\_7,cas8c\_I-C\_4 |
| NC\_014364.1\_CAS\_Class1-Subtype-IV-B\_6 | Cas | CAS\_Class1-Subtype-IV-B | NC\_014364.1\_2494 | NC\_014364.1\_2499 | NC\_014364.1\_2494,NC\_014364.1\_2496,NC\_014364.1\_2497,NC\_014364.1\_2498,NC\_014364.1\_2499 | 5 | csf1gr8\_IV-A\_5,csf2gr7\_IV\_1,csf3gr5\_IV-B\_1,csf4gr11\_IV-B\_4,cysH\_IV-B\_1 |
| NC\_014364.1\_AbiE\_1 | AbiE | AbiE | NC\_014364.1\_2619 | NC\_014364.1\_2620 | NC\_014364.1\_2619,NC\_014364.1\_2620 | 2 | AbiEii\_\_AbiEi\_4,AbiEii\_\_AbiEii |

**Genomad and CheckV Summary**:

| seq\_name | length | gene\_count | viral\_genes | checkv\_quality | miuvig\_quality | taxonomy | topology | coordinates |
| --- | --- | --- | --- | --- | --- | --- | --- | --- |
| NC\_014364.1\|provirus\_2666253\_2703772 | 37520 | 56 | 15 | Medium-quality | Genome-fragment | Viruses;Duplodnaviria;Heunggongvirae;Uroviricota;Caudoviricetes;; | Provirus | 2666253-2703772 |
| NC\_014364.1\|provirus\_2976748\_3014759 | 38012 | 44 | 12 | Medium-quality | Genome-fragment | Viruses;Duplodnaviria;Heunggongvirae;Uroviricota;Caudoviricetes;; | Provirus | 2976748-3014759 |
| NC\_014364.1\|provirus\_2119034\_2157334 | 38301 | 47 | 12 | Medium-quality | Genome-fragment | Viruses;Duplodnaviria;Heunggongvirae;Uroviricota;Caudoviricetes;; | Provirus | 2119034-2157334 |
| NC\_014364.1\|provirus\_2329685\_2350621 | 20937 | 29 | 6 | Low-quality | Genome-fragment | Viruses;Duplodnaviria;Heunggongvirae;Uroviricota;Caudoviricetes;; | Provirus | 2329685-2350621 |

**Host Genome Ideogram with Phages**:

In
this circular plot, **“c”** indicates the contig, and the
number that follows (e.g., **c1**) represents the contig
number. If multiple contigs are present in the genome, each will be
shown with a distinct label (e.g., **c1**,
**c2**, etc.).

##### 2.12.2 Prophages

**Select prophage to show:**

###### 2.12.2.1 Phage ID: NC\_014364.1|provirus\_2666253\_2703772

**Phage-Host Genome Ideogram:**

**Genes Annotation (geNomad):**

| gene | length | marker | annotation\_accessions | annotation\_description |
| --- | --- | --- | --- | --- |
| NC\_014364.1\|provirus\_2666253\_2703772\_2490 | 1263 | NA | NA | NA |
| NC\_014364.1\|provirus\_2666253\_2703772\_2491 | 441 | GENOMAD.167054.PV | PF06892;TIGR02612;COG3655 | mobile mystery protein A |
| NC\_014364.1\|provirus\_2666253\_2703772\_2492 | 384 | NA | NA | NA |
| NC\_014364.1\|provirus\_2666253\_2703772\_2493 | 249 | NA | NA | NA |
| NC\_014364.1\|provirus\_2666253\_2703772\_2494 | 417 | NA | NA | NA |
| NC\_014364.1\|provirus\_2666253\_2703772\_2495 | 561 | NA | NA | NA |
| NC\_014364.1\|provirus\_2666253\_2703772\_2496 | 426 | NA | NA | NA |
| NC\_014364.1\|provirus\_2666253\_2703772\_2497 | 249 | NA | NA | NA |
| NC\_014364.1\|provirus\_2666253\_2703772\_2498 | 102 | NA | NA | NA |
| NC\_014364.1\|provirus\_2666253\_2703772\_2499 | 372 | NA | NA | NA |
| NC\_014364.1\|provirus\_2666253\_2703772\_2500 | 234 | NA | NA | NA |
| NC\_014364.1\|provirus\_2666253\_2703772\_2501 | 612 | GENOMAD.008561.VV | PF16778 | Phage tail assembly chaperone protein |
| NC\_014364.1\|provirus\_2666253\_2703772\_2502 | 3315 | GENOMAD.109300.VV | NA | NA |
| NC\_014364.1\|provirus\_2666253\_2703772\_2503 | 2178 | GENOMAD.040051.VV | NA | NA |
| NC\_014364.1\|provirus\_2666253\_2703772\_2504 | 276 | NA | NA | NA |
| NC\_014364.1\|provirus\_2666253\_2703772\_2505 | 402 | NA | NA | NA |
| NC\_014364.1\|provirus\_2666253\_2703772\_2506 | 756 | NA | NA | NA |
| NC\_014364.1\|provirus\_2666253\_2703772\_2507 | 399 | NA | NA | NA |
| NC\_014364.1\|provirus\_2666253\_2703772\_2508 | 426 | GENOMAD.060383.VV | PF04883;TIGR01725;COG5005 | phage protein, HK97 gp10 family |
| NC\_014364.1\|provirus\_2666253\_2703772\_2509 | 558 | GENOMAD.080176.VV | PF11436;TIGR02215 | phage conserved hypothetical protein, phiE125 gp8 family |
| NC\_014364.1\|provirus\_2666253\_2703772\_2510 | 414 | GENOMAD.210649.VV | NA | NA |
| NC\_014364.1\|provirus\_2666253\_2703772\_2511 | 1251 | GENOMAD.090434.VV | PF05065;TIGR01554;COG4653 | phage major capsid protein, HK97 family |
| NC\_014364.1\|provirus\_2666253\_2703772\_2512 | 708 | GENOMAD.126169.VV | PF04586;K06904;COG3740;TIGR01543 | Phage head maturation protease |
| NC\_014364.1\|provirus\_2666253\_2703772\_2513 | 1617 | GENOMAD.179073.VP | NA | NA |
| NC\_014364.1\|provirus\_2666253\_2703772\_2514 | 1725 | GENOMAD.194580.VP | PF03354;PF05521;PF20441;COG4626;TIGR01563 | Phage terminase-like protein, large subunit, contains N-terminal HTH domain |
| NC\_014364.1\|provirus\_2666253\_2703772\_2515 | 510 | GENOMAD.170837.VV | PF05119;COG3747;TIGR01558 | Phage terminase, small subunit |
| NC\_014364.1\|provirus\_2666253\_2703772\_2516 | 399 | NA | NA | NA |
| NC\_014364.1\|provirus\_2666253\_2703772\_2517 | 708 | NA | NA | NA |
| NC\_014364.1\|provirus\_2666253\_2703772\_2518 | 237 | NA | NA | NA |
| NC\_014364.1\|provirus\_2666253\_2703772\_2519 | 678 | GENOMAD.222821.VP | TIGR03116 | CRISPR type IV/AFERR-associated protein Csf3 |
| NC\_014364.1\|provirus\_2666253\_2703772\_2520 | 942 | NA | NA | NA |
| NC\_014364.1\|provirus\_2666253\_2703772\_2521 | 354 | NA | NA | NA |
| NC\_014364.1\|provirus\_2666253\_2703772\_2522 | 741 | GENOMAD.222446.VP | TIGR03114 | CRISPR type AFERR-associated protein Csf1 |
| NC\_014364.1\|provirus\_2666253\_2703772\_2523 | 186 | NA | NA | NA |
| NC\_014364.1\|provirus\_2666253\_2703772\_2524 | 192 | NA | NA | NA |
| NC\_014364.1\|provirus\_2666253\_2703772\_2525 | 219 | NA | NA | NA |
| NC\_014364.1\|provirus\_2666253\_2703772\_2526 | 402 | NA | NA | NA |
| NC\_014364.1\|provirus\_2666253\_2703772\_2527 | 1089 | GENOMAD.018355.VV | PF09681;TIGR01714 | phage replisome organizer, putative, N-terminal region |
| NC\_014364.1\|provirus\_2666253\_2703772\_2528 | 597 | NA | NA | NA |
| NC\_014364.1\|provirus\_2666253\_2703772\_2529 | 414 | GENOMAD.002027.VV | PF05766 | Bacteriophage Lambda NinG protein |
| NC\_014364.1\|provirus\_2666253\_2703772\_2530 | 2514 | GENOMAD.126436.VP | PF13872 | NA |
| NC\_014364.1\|provirus\_2666253\_2703772\_2531 | 477 | GENOMAD.111167.VV | PF16784 | Putative HNHc nuclease |
| NC\_014364.1\|provirus\_2666253\_2703772\_2532 | 606 | GENOMAD.176983.VP | NA | NA |
| NC\_014364.1\|provirus\_2666253\_2703772\_2533 | 813 | GENOMAD.177891.VP | PF03837;TIGR01913;COG3723 | phage recombination protein Bet |
| NC\_014364.1\|provirus\_2666253\_2703772\_2534 | 756 | NA | NA | NA |
| NC\_014364.1\|provirus\_2666253\_2703772\_2535 | 204 | NA | NA | NA |
| NC\_014364.1\|provirus\_2666253\_2703772\_2536 | 735 | GENOMAD.014438.VV | NA | NA |
| NC\_014364.1\|provirus\_2666253\_2703772\_2537 | 348 | GENOMAD.100208.VV | NA | NA |
| NC\_014364.1\|provirus\_2666253\_2703772\_2538 | 264 | NA | NA | NA |
| NC\_014364.1\|provirus\_2666253\_2703772\_2539 | 342 | NA | NA | NA |
| NC\_014364.1\|provirus\_2666253\_2703772\_2540 | 423 | NA | NA | NA |
| NC\_014364.1\|provirus\_2666253\_2703772\_2541 | 219 | NA | NA | NA |
| NC\_014364.1\|provirus\_2666253\_2703772\_2542 | 348 | NA | NA | NA |
| NC\_014364.1\|provirus\_2666253\_2703772\_2543 | 504 | NA | NA | NA |
| NC\_014364.1\|provirus\_2666253\_2703772\_2544 | 291 | GENOMAD.219421.VC | NA | NA |
| NC\_014364.1\|provirus\_2666253\_2703772\_2545 | 363 | GENOMAD.227798.VP | NA | NA |

###### 2.12.2.2 Phage ID: NC\_014364.1|provirus\_2976748\_3014759

**Phage-Host Genome Ideogram:**

**Genes Annotation (geNomad):**

| gene | length | marker | annotation\_accessions | annotation\_description |
| --- | --- | --- | --- | --- |
| NC\_014364.1\|provirus\_2976748\_3014759\_2796 | 1251 | NA | NA | NA |
| NC\_014364.1\|provirus\_2976748\_3014759\_2797 | 474 | NA | NA | NA |
| NC\_014364.1\|provirus\_2976748\_3014759\_2798 | 186 | NA | NA | NA |
| NC\_014364.1\|provirus\_2976748\_3014759\_2799 | 315 | NA | NA | NA |
| NC\_014364.1\|provirus\_2976748\_3014759\_2800 | 456 | NA | NA | NA |
| NC\_014364.1\|provirus\_2976748\_3014759\_2801 | 381 | NA | NA | NA |
| NC\_014364.1\|provirus\_2976748\_3014759\_2802 | 228 | NA | NA | NA |
| NC\_014364.1\|provirus\_2976748\_3014759\_2803 | 543 | GENOMAD.112664.VP | PF11195 | NA |
| NC\_014364.1\|provirus\_2976748\_3014759\_2804 | 255 | NA | NA | NA |
| NC\_014364.1\|provirus\_2976748\_3014759\_2805 | 1092 | GENOMAD.032961.VV | PF13479;TIGR01618;COG2087;K04484 | phage nucleotide-binding protein |
| NC\_014364.1\|provirus\_2976748\_3014759\_2806 | 651 | GENOMAD.025041.VV | NA | NA |
| NC\_014364.1\|provirus\_2976748\_3014759\_2807 | 639 | GENOMAD.067346.VV | PF06023;TIGR00372;COG4343 | CRISPR-associated protein Cas4 |
| NC\_014364.1\|provirus\_2976748\_3014759\_2808 | 474 | GENOMAD.111167.VV | PF16784 | Putative HNHc nuclease |
| NC\_014364.1\|provirus\_2976748\_3014759\_2809 | 447 | GENOMAD.002027.VV | PF05766 | Bacteriophage Lambda NinG protein |
| NC\_014364.1\|provirus\_2976748\_3014759\_2810 | 543 | NA | NA | NA |
| NC\_014364.1\|provirus\_2976748\_3014759\_2811 | 1098 | NA | NA | NA |
| NC\_014364.1\|provirus\_2976748\_3014759\_2812 | 402 | NA | NA | NA |
| NC\_014364.1\|provirus\_2976748\_3014759\_2813 | 219 | NA | NA | NA |
| NC\_014364.1\|provirus\_2976748\_3014759\_2814 | 207 | NA | NA | NA |
| NC\_014364.1\|provirus\_2976748\_3014759\_2815 | 720 | NA | NA | NA |
| NC\_014364.1\|provirus\_2976748\_3014759\_2816 | 870 | NA | NA | NA |
| NC\_014364.1\|provirus\_2976748\_3014759\_2817 | 678 | NA | NA | NA |
| NC\_014364.1\|provirus\_2976748\_3014759\_2818 | 177 | NA | NA | NA |
| NC\_014364.1\|provirus\_2976748\_3014759\_2819 | 543 | NA | NA | NA |
| NC\_014364.1\|provirus\_2976748\_3014759\_2820 | 330 | NA | NA | NA |
| NC\_014364.1\|provirus\_2976748\_3014759\_2821 | 1683 | GENOMAD.060970.VV | PF12236 | Bacteriophage head to tail connecting protein |
| NC\_014364.1\|provirus\_2976748\_3014759\_2822 | 303 | GENOMAD.118268.VV | NA | NA |
| NC\_014364.1\|provirus\_2976748\_3014759\_2823 | 807 | GENOMAD.072294.VV | NA | NA |
| NC\_014364.1\|provirus\_2976748\_3014759\_2824 | 987 | GENOMAD.004790.VV | PF19307 | Phage capsid-like protein |
| NC\_014364.1\|provirus\_2976748\_3014759\_2825 | 375 | GENOMAD.013948.VV | PF21190 | Bbp16 |
| NC\_014364.1\|provirus\_2976748\_3014759\_2826 | 315 | NA | NA | NA |
| NC\_014364.1\|provirus\_2976748\_3014759\_2827 | 594 | GENOMAD.061091.VV | PF17212 | Tail tubular protein |
| NC\_014364.1\|provirus\_2976748\_3014759\_2828 | 1713 | NA | NA | NA |
| NC\_014364.1\|provirus\_2976748\_3014759\_2829 | 888 | NA | NA | NA |
| NC\_014364.1\|provirus\_2976748\_3014759\_2830 | 1947 | NA | NA | NA |
| NC\_014364.1\|provirus\_2976748\_3014759\_2831 | 6246 | NA | NA | NA |
| NC\_014364.1\|provirus\_2976748\_3014759\_2832 | 3657 | GENOMAD.089557.VV | K03987 | NA |
| NC\_014364.1\|provirus\_2976748\_3014759\_2833 | 612 | GENOMAD.007262.VV | PF16778 | Phage tail assembly chaperone protein |
| NC\_014364.1\|provirus\_2976748\_3014759\_2834 | 234 | NA | NA | NA |
| NC\_014364.1\|provirus\_2976748\_3014759\_2835 | 480 | NA | NA | NA |
| NC\_014364.1\|provirus\_2976748\_3014759\_2836 | 222 | NA | NA | NA |
| NC\_014364.1\|provirus\_2976748\_3014759\_2837 | 417 | NA | NA | NA |
| NC\_014364.1\|provirus\_2976748\_3014759\_2838 | 1110 | NA | NA | NA |

###### 2.12.2.3 Phage ID: NC\_014364.1|provirus\_2119034\_2157334

**Phage-Host Genome Ideogram:**

**Genes Annotation (geNomad):**

| gene | length | marker | annotation\_accessions | annotation\_description |
| --- | --- | --- | --- | --- |
| NC\_014364.1\|provirus\_2119034\_2157334\_1977 | 765 | GENOMAD.168064.VC | TIGR02869 | spore cortex-lytic enzyme |
| NC\_014364.1\|provirus\_2119034\_2157334\_1978 | 1188 | NA | NA | NA |
| NC\_014364.1\|provirus\_2119034\_2157334\_1979 | 282 | NA | NA | NA |
| NC\_014364.1\|provirus\_2119034\_2157334\_1980 | 1875 | NA | NA | NA |
| NC\_014364.1\|provirus\_2119034\_2157334\_1981 | 1542 | NA | NA | NA |
| NC\_014364.1\|provirus\_2119034\_2157334\_1982 | 1671 | NA | NA | NA |
| NC\_014364.1\|provirus\_2119034\_2157334\_1983 | 1191 | NA | NA | NA |
| NC\_014364.1\|provirus\_2119034\_2157334\_1984 | 219 | NA | NA | NA |
| NC\_014364.1\|provirus\_2119034\_2157334\_1985 | 369 | NA | NA | NA |
| NC\_014364.1\|provirus\_2119034\_2157334\_1986 | 189 | NA | NA | NA |
| NC\_014364.1\|provirus\_2119034\_2157334\_1987 | 489 | GENOMAD.053051.VV | PF10772;PF02086;TIGR00571;COG0338 | Bacteriophage HP1, Orf24; D12 class N6 adenine-specific DNA methyltransferase |
| NC\_014364.1\|provirus\_2119034\_2157334\_1988 | 348 | NA | NA | NA |
| NC\_014364.1\|provirus\_2119034\_2157334\_1989 | 819 | GENOMAD.177891.VP | PF03837;TIGR01913;COG3723 | phage recombination protein Bet |
| NC\_014364.1\|provirus\_2119034\_2157334\_1990 | 690 | GENOMAD.183319.VP | NA | NA |
| NC\_014364.1\|provirus\_2119034\_2157334\_1991 | 477 | GENOMAD.111167.VV | PF16784 | Putative HNHc nuclease |
| NC\_014364.1\|provirus\_2119034\_2157334\_1992 | 393 | GENOMAD.002027.VV | PF05766 | Bacteriophage Lambda NinG protein |
| NC\_014364.1\|provirus\_2119034\_2157334\_1993 | 603 | NA | NA | NA |
| NC\_014364.1\|provirus\_2119034\_2157334\_1994 | 1089 | GENOMAD.018355.VV | PF09681;TIGR01714 | phage replisome organizer, putative, N-terminal region |
| NC\_014364.1\|provirus\_2119034\_2157334\_1995 | 402 | NA | NA | NA |
| NC\_014364.1\|provirus\_2119034\_2157334\_1996 | 219 | NA | NA | NA |
| NC\_014364.1\|provirus\_2119034\_2157334\_1997 | 435 | NA | NA | NA |
| NC\_014364.1\|provirus\_2119034\_2157334\_1998 | 729 | NA | NA | NA |
| NC\_014364.1\|provirus\_2119034\_2157334\_1999 | 510 | GENOMAD.170837.VV | PF05119;COG3747;TIGR01558 | Phage terminase, small subunit |
| NC\_014364.1\|provirus\_2119034\_2157334\_2000 | 1722 | GENOMAD.194580.VP | PF03354;PF05521;PF20441;COG4626;TIGR01563 | Phage terminase-like protein, large subunit, contains N-terminal HTH domain |
| NC\_014364.1\|provirus\_2119034\_2157334\_2001 | 1242 | GENOMAD.179073.VP | NA | NA |
| NC\_014364.1\|provirus\_2119034\_2157334\_2002 | 834 | GENOMAD.096083.VP | PF00574;K01358;TIGR00493;COG0740 | ATP-dependent Clp endopeptidase, proteolytic subunit ClpP |
| NC\_014364.1\|provirus\_2119034\_2157334\_2003 | 1200 | GENOMAD.080602.VV | PF05135;TIGR02215 | phage conserved hypothetical protein, phiE125 gp8 family |
| NC\_014364.1\|provirus\_2119034\_2157334\_2004 | 423 | GENOMAD.129880.VV | NA | NA |
| NC\_014364.1\|provirus\_2119034\_2157334\_2005 | 558 | GENOMAD.080176.VV | PF11436;TIGR02215 | phage conserved hypothetical protein, phiE125 gp8 family |
| NC\_014364.1\|provirus\_2119034\_2157334\_2006 | 453 | GENOMAD.060383.VV | PF04883;TIGR01725;COG5005 | phage protein, HK97 gp10 family |
| NC\_014364.1\|provirus\_2119034\_2157334\_2007 | 399 | NA | NA | NA |
| NC\_014364.1\|provirus\_2119034\_2157334\_2008 | 696 | NA | NA | NA |
| NC\_014364.1\|provirus\_2119034\_2157334\_2009 | 402 | NA | NA | NA |
| NC\_014364.1\|provirus\_2119034\_2157334\_2010 | 276 | NA | NA | NA |
| NC\_014364.1\|provirus\_2119034\_2157334\_2011 | 2274 | GENOMAD.009561.VV | NA | NA |
| NC\_014364.1\|provirus\_2119034\_2157334\_2012 | 3279 | GENOMAD.109300.VV | NA | NA |
| NC\_014364.1\|provirus\_2119034\_2157334\_2013 | 612 | GENOMAD.008561.VV | PF16778 | Phage tail assembly chaperone protein |
| NC\_014364.1\|provirus\_2119034\_2157334\_2014 | 234 | NA | NA | NA |
| NC\_014364.1\|provirus\_2119034\_2157334\_2015 | 471 | NA | NA | NA |
| NC\_014364.1\|provirus\_2119034\_2157334\_2016 | 249 | NA | NA | NA |
| NC\_014364.1\|provirus\_2119034\_2157334\_2017 | 390 | NA | NA | NA |
| NC\_014364.1\|provirus\_2119034\_2157334\_2018 | 369 | NA | NA | NA |
| NC\_014364.1\|provirus\_2119034\_2157334\_2019 | 324 | NA | NA | NA |
| NC\_014364.1\|provirus\_2119034\_2157334\_2020 | 786 | NA | NA | NA |
| NC\_014364.1\|provirus\_2119034\_2157334\_2021 | 1242 | NA | NA | NA |

###### 2.12.2.4 Phage ID: NC\_014364.1|provirus\_2329685\_2350621

**Phage-Host Genome Ideogram:**

**Genes Annotation (geNomad):**

| gene | length | marker | annotation\_accessions | annotation\_description |
| --- | --- | --- | --- | --- |
| NC\_014364.1\|provirus\_2329685\_2350621\_2174 | 1266 | NA | NA | NA |
| NC\_014364.1\|provirus\_2329685\_2350621\_2175 | 516 | NA | NA | NA |
| NC\_014364.1\|provirus\_2329685\_2350621\_2176 | 384 | NA | NA | NA |
| NC\_014364.1\|provirus\_2329685\_2350621\_2177 | 1224 | NA | NA | NA |
| NC\_014364.1\|provirus\_2329685\_2350621\_2178 | 618 | NA | NA | NA |
| NC\_014364.1\|provirus\_2329685\_2350621\_2179 | 1035 | NA | NA | NA |
| NC\_014364.1\|provirus\_2329685\_2350621\_2180 | 1068 | NA | NA | NA |
| NC\_014364.1\|provirus\_2329685\_2350621\_2181 | 342 | NA | NA | NA |
| NC\_014364.1\|provirus\_2329685\_2350621\_2182 | 231 | NA | NA | NA |
| NC\_014364.1\|provirus\_2329685\_2350621\_2183 | 246 | NA | NA | NA |
| NC\_014364.1\|provirus\_2329685\_2350621\_2184 | 204 | NA | NA | NA |
| NC\_014364.1\|provirus\_2329685\_2350621\_2185 | 543 | GENOMAD.112664.VP | PF11195 | NA |
| NC\_014364.1\|provirus\_2329685\_2350621\_2186 | 255 | NA | NA | NA |
| NC\_014364.1\|provirus\_2329685\_2350621\_2187 | 1116 | GENOMAD.090362.VV | PF13479;TIGR01618;COG2087 | phage nucleotide-binding protein |
| NC\_014364.1\|provirus\_2329685\_2350621\_2188 | 651 | GENOMAD.025041.VV | NA | NA |
| NC\_014364.1\|provirus\_2329685\_2350621\_2189 | 639 | GENOMAD.067346.VV | PF06023;TIGR00372;COG4343 | CRISPR-associated protein Cas4 |
| NC\_014364.1\|provirus\_2329685\_2350621\_2190 | 474 | GENOMAD.111167.VV | PF16784 | Putative HNHc nuclease |
| NC\_014364.1\|provirus\_2329685\_2350621\_2191 | 234 | NA | NA | NA |
| NC\_014364.1\|provirus\_2329685\_2350621\_2192 | 540 | NA | NA | NA |
| NC\_014364.1\|provirus\_2329685\_2350621\_2193 | 1056 | NA | NA | NA |
| NC\_014364.1\|provirus\_2329685\_2350621\_2194 | 414 | NA | NA | NA |
| NC\_014364.1\|provirus\_2329685\_2350621\_2195 | 219 | NA | NA | NA |
| NC\_014364.1\|provirus\_2329685\_2350621\_2196 | 192 | NA | NA | NA |
| NC\_014364.1\|provirus\_2329685\_2350621\_2197 | 201 | NA | NA | NA |
| NC\_014364.1\|provirus\_2329685\_2350621\_2198 | 1287 | NA | NA | NA |
| NC\_014364.1\|provirus\_2329685\_2350621\_2199 | 981 | NA | NA | NA |
| NC\_014364.1\|provirus\_2329685\_2350621\_2200 | 507 | GENOMAD.170837.VV | PF05119;COG3747;TIGR01558 | Phage terminase, small subunit |
| NC\_014364.1\|provirus\_2329685\_2350621\_2201 | 1323 | GENOMAD.194580.VP | PF03354;PF05521;PF20441;COG4626;TIGR01563 | Phage terminase-like protein, large subunit, contains N-terminal HTH domain |

##### 2.12.3 vOTUs

When more than 1 phage is detected in the host genome, we perform a
clustering step using the tool dRep.

A threshold of 0.95 was applied to the ANI similarity index to define
clusters of virus operational taxonomic units (vOTUs).

**Final cluster designations**

| genomad\_summary$seq\_name | genome | secondary\_cluster | threshold | cluster\_method | comparison\_algorithm | primary\_cluster |
| --- | --- | --- | --- | --- | --- | --- |
| NC\_014364.1\|provirus\_2666253\_2703772 | sequence\_000000.fasta.fasta | 1\_1 | 0.05 | average | ANImf | 1 |
| NC\_014364.1\|provirus\_2976748\_3014759 | sequence\_000002.fasta.fasta | 1\_2 | 0.05 | average | ANImf | 1 |
| NC\_014364.1\|provirus\_2119034\_2157334 | sequence\_000001.fasta.fasta | 2\_0 | 0.05 | average | ANImf | 2 |
| NC\_014364.1\|provirus\_2329685\_2350621 | sequence\_000003.fasta.fasta | 3\_0 | 0.05 | average | ANImf | 3 |

**Primary clustering plot**

##### 2.12.4 Prophage-Host Prediction

Prediction of potential hosts for the prophage contigs.

| virus | host\_genome | host\_taxonomy | main\_method | confidence\_score | additional\_methods |
| --- | --- | --- | --- | --- | --- |
| NC\_014364.1\|provirus\_2119034\_2157334 | RS\_GCF\_000143985.1 | d\_\_Bacteria;p\_\_Spirochaetota;c\_\_Spirochaetia;o\_\_DSM-16054;f\_\_Sediminispirochaetaceae;g\_\_Sediminispirochaeta;s\_\_Sediminispirochaeta smaragdinae | CRISPR | 95.4 | blast;94.80 iPHoP-RF;70.40 |
| NC\_014364.1\|provirus\_2119034\_2157334 | RS\_GCF\_000378205.1 | d\_\_Bacteria;p\_\_Spirochaetota;c\_\_Spirochaetia;o\_\_DSM-16054;f\_\_Sediminispirochaetaceae;g\_\_Sediminispirochaeta;s\_\_Sediminispirochaeta bajacaliforniensis | CRISPR | 93.4 | blast;92.00 |
| NC\_014364.1\|provirus\_2329685\_2350621 | RS\_GCF\_000143985.1 | d\_\_Bacteria;p\_\_Spirochaetota;c\_\_Spirochaetia;o\_\_DSM-16054;f\_\_Sediminispirochaetaceae;g\_\_Sediminispirochaeta;s\_\_Sediminispirochaeta smaragdinae | blast | 95.7 | iPHoP-RF;65.10 |
| NC\_014364.1\|provirus\_2329685\_2350621 | RS\_GCF\_000378205.1 | d\_\_Bacteria;p\_\_Spirochaetota;c\_\_Spirochaetia;o\_\_DSM-16054;f\_\_Sediminispirochaetaceae;g\_\_Sediminispirochaeta;s\_\_Sediminispirochaeta bajacaliforniensis | blast | 92.1 | iPHoP-RF;52.80 |
| NC\_014364.1\|provirus\_2666253\_2703772 | RS\_GCF\_000143985.1 | d\_\_Bacteria;p\_\_Spirochaetota;c\_\_Spirochaetia;o\_\_DSM-16054;f\_\_Sediminispirochaetaceae;g\_\_Sediminispirochaeta;s\_\_Sediminispirochaeta smaragdinae | blast | 96.2 | CRISPR;74.20 iPHoP-RF;72.60 |
| NC\_014364.1\|provirus\_2976748\_3014759 | RS\_GCF\_000143985.1 | d\_\_Bacteria;p\_\_Spirochaetota;c\_\_Spirochaetia;o\_\_DSM-16054;f\_\_Sediminispirochaetaceae;g\_\_Sediminispirochaeta;s\_\_Sediminispirochaeta smaragdinae | blast | 96.2 | iPHoP-RF;69.80 |

##### 2.12.5 AMG Predictions

Prediction of auxiliary metabolic genes in the prophage contigs.

| protein | scaffold | amg\_ko | amg\_ko\_name | pfam | pfam\_name |
| --- | --- | --- | --- | --- | --- |
| NC\_014364.1\|provirus\_2666253\_2703772\_28 | NC\_014364.1\|provirus\_2666253\_2703772 | K00390 | cysH; phosphoadenosine phosphosulfate reductase [EC:1.8.4.8 1.8.4.10] | PF01507.19 | Phosphoadenosine phosphosulfate reductase family |

## 2.13 NC\_015761

##### 2.13.1 Host Genome

**GTDB-Tk taxonomy**:

classification

d\_\_Bacteria;p\_\_Pseudomonadota;c\_\_Gammaproteobacteria;o\_\_Enterobacterales;f\_\_Enterobacteriaceae;g\_\_Salmonella;s\_\_Salmonella
bongori

**CheckM2 Summary**:

| total\_contigs | contig\_n50 | completeness | contamination |
| --- | --- | --- | --- |
| 1 | 4460105 | 100 | 0.14 |

**Defense-Finder Systems**:

| sys\_id | type | subtype | sys\_beg | sys\_end | protein\_in\_syst | genes\_count | name\_of\_profiles\_in\_sys |
| --- | --- | --- | --- | --- | --- | --- | --- |
| NC\_015761.1\_PrrC\_6 | PrrC | PrrC | NC\_015761.1\_263 | NC\_015761.1\_265 | NC\_015761.1\_263,NC\_015761.1\_264,NC\_015761.1\_265 | 3 | PrrC\_\_EcoprrI,PrrC\_\_EcoprrI,RM\_\_Type\_I\_REases\_FAM\_0.einsi\_trimmed |
| NC\_015761.1\_DarTG\_2 | DarTG | DarTG | NC\_015761.1\_279 | NC\_015761.1\_280 | NC\_015761.1\_279,NC\_015761.1\_280 | 2 | DarTG\_\_DarG,DarTG\_\_DarT |
| NC\_015761.1\_RM\_Type\_III\_7 | RM | RM\_Type\_III | NC\_015761.1\_310 | NC\_015761.1\_311 | NC\_015761.1\_310,NC\_015761.1\_311 | 2 | RM\_Type\_III\_\_Type\_III\_MTases\_FAM\_0,RM\_Type\_III\_\_Type\_III\_REases\_FAM\_0.einsi\_trimmed |
| NC\_015761.1\_PfiAT\_4 | PfiAT | PfiAT | NC\_015761.1\_932 | NC\_015761.1\_933 | NC\_015761.1\_932,NC\_015761.1\_933 | 2 | PfiAT\_\_PfiA,PfiAT\_\_PfiT |
| NC\_015761.1\_CAS\_Class1-Subtype-I-E\_8 | Cas | CAS\_Class1-Subtype-I-E | NC\_015761.1\_2549 | NC\_015761.1\_2556 | NC\_015761.1\_2549,NC\_015761.1\_2550,NC\_015761.1\_2551,NC\_015761.1\_2552,NC\_015761.1\_2553,NC\_015761.1\_2554,NC\_015761.1\_2555,NC\_015761.1\_2556 | 8 | cas1\_I-E\_1,cas2\_I-E\_2,cas3\_I\_5,cas5\_I-E\_3,cas6e\_I\_II\_III\_IV\_V\_VI\_1,cas7\_I-E\_2,cas8e\_I-E\_1,cse2gr11\_I-E\_2 |
| NC\_015761.1\_dCTPdeaminase\_5 | dCTPdeaminase | dCTPdeaminase | NC\_015761.1\_3620 | NC\_015761.1\_3620 | NC\_015761.1\_3620 | 1 | dCTPdeaminase\_\_dCTPdeaminase |
| NC\_015761.1\_Mokosh\_TypeII\_3 | Mokosh | Mokosh\_TypeII | NC\_015761.1\_3957 | NC\_015761.1\_3957 | NC\_015761.1\_3957 | 1 | Mokosh\_TypeII\_\_MkoC |
| NC\_015761.1\_DS-17\_1 | DS-17 | DS-17 | NC\_015761.1\_3965 | NC\_015761.1\_3965 | NC\_015761.1\_3965 | 1 | DS-17\_\_DS-17 |

**Genomad and CheckV Summary**:

| seq\_name | length | gene\_count | viral\_genes | checkv\_quality | miuvig\_quality | taxonomy | topology | coordinates |
| --- | --- | --- | --- | --- | --- | --- | --- | --- |
| NC\_015761.1\|provirus\_3084741\_3115735 | 30995 | 41 | 32 | High-quality | High-quality | Viruses;Duplodnaviria;Heunggongvirae;Uroviricota;Caudoviricetes;; | Provirus | 3084741-3115735 |
| NC\_015761.1\|provirus\_847397\_862760 | 15364 | 19 | 11 | Low-quality | Genome-fragment | Viruses;Duplodnaviria;Heunggongvirae;Uroviricota;Caudoviricetes;; | Provirus | 847397-862760 |
| NC\_015761.1\|provirus\_1007223\_1043740 | 36518 | 51 | 17 | Medium-quality | Genome-fragment | Viruses;Duplodnaviria;Heunggongvirae;Uroviricota;Caudoviricetes;; | Provirus | 1007223-1043740 |

**Host Genome Ideogram with Phages**:

In
this circular plot, **“c”** indicates the contig, and the
number that follows (e.g., **c1**) represents the contig
number. If multiple contigs are present in the genome, each will be
shown with a distinct label (e.g., **c1**,
**c2**, etc.).

##### 2.13.2 Prophages

**Select prophage to show:**

###### 2.13.2.1 Phage ID: NC\_015761.1|provirus\_3084741\_3115735

**Phage-Host Genome Ideogram:**

**Genes Annotation (geNomad):**

| gene | length | marker | annotation\_accessions | annotation\_description |
| --- | --- | --- | --- | --- |
| NC\_015761.1\|provirus\_3084741\_3115735\_2808 | 219 | GENOMAD.223866.VP | PF04606;TIGR02098 | Ogr/Delta-like zinc finger |
| NC\_015761.1\|provirus\_3084741\_3115735\_2809 | 1170 | GENOMAD.209614.VP | PF05954;K06905;COG3500;TIGR03361 | Phage protein D |
| NC\_015761.1\|provirus\_3084741\_3115735\_2810 | 486 | GENOMAD.125908.VV | PF06995;K06906;COG3499 | Phage P2 GpU |
| NC\_015761.1\|provirus\_3084741\_3115735\_2811 | 2442 | GENOMAD.130201.VP | COG5283 | Phage-related tail protein |
| NC\_015761.1\|provirus\_3084741\_3115735\_2812 | 336 | GENOMAD.135969.VP | PF10109 | Phage tail assembly chaperone proteins, E, or 41 or 14 |
| NC\_015761.1\|provirus\_3084741\_3115735\_2813 | 519 | GENOMAD.222086.VV | PF04985;K06908;TIGR01611;COG3498 | phage contractile tail tube protein, P2 family |
| NC\_015761.1\|provirus\_3084741\_3115735\_2814 | 1188 | GENOMAD.102019.VV | PF10758;COG3497 | Phage tail sheath protein FI |
| NC\_015761.1\|provirus\_3084741\_3115735\_2815 | 549 | GENOMAD.209102.VP | NA | NA |
| NC\_015761.1\|provirus\_3084741\_3115735\_2816 | 1977 | GENOMAD.151930.VP | PF12571;COG5301 | Phage-related tail fibre protein |
| NC\_015761.1\|provirus\_3084741\_3115735\_2817 | 531 | GENOMAD.114503.VV | PF09684;COG4385;TIGR01634 | Bacteriophage P2-related tail formation protein |
| NC\_015761.1\|provirus\_3084741\_3115735\_2818 | 909 | GENOMAD.105501.VV | PF03434;COG3948 | Phage-related baseplate assembly protein |
| NC\_015761.1\|provirus\_3084741\_3115735\_2819 | 348 | GENOMAD.123275.VP | PF04965;K06903;COG3628;TIGR03357 | Phage baseplate assembly protein W |
| NC\_015761.1\|provirus\_3084741\_3115735\_2820 | 642 | GENOMAD.166314.VP | K22111;TIGR01644;COG4540 | phage baseplate assembly protein V |
| NC\_015761.1\|provirus\_3084741\_3115735\_2821 | 450 | GENOMAD.140989.VP | PF05069;TIGR01635;COG5005 | phage virion morphogenesis (putative tail completion) protein |
| NC\_015761.1\|provirus\_3084741\_3115735\_2822 | 468 | GENOMAD.125799.VP | PF06891 | P2 phage tail completion protein R (GpR) |
| NC\_015761.1\|provirus\_3084741\_3115735\_2823 | 159 | GENOMAD.116277.VV | NA | NA |
| NC\_015761.1\|provirus\_3084741\_3115735\_2824 | 414 | GENOMAD.191084.VP | PF10828;TIGR03495 | phage lysis regulatory protein, LysB family |
| NC\_015761.1\|provirus\_3084741\_3115735\_2825 | 498 | NA | NA | NA |
| NC\_015761.1\|provirus\_3084741\_3115735\_2826 | 297 | GENOMAD.177230.VP | PF04550;TIGR01594 | Phage holin family 2 |
| NC\_015761.1\|provirus\_3084741\_3115735\_2827 | 189 | GENOMAD.159179.VP | COG5004;K06370 | P2-like prophage tail protein X |
| NC\_015761.1\|provirus\_3084741\_3115735\_2828 | 507 | GENOMAD.119083.VV | PF05926 | Phage head completion protein (GPL) |
| NC\_015761.1\|provirus\_3084741\_3115735\_2829 | 750 | GENOMAD.034571.VV | PF05944 | Phage small terminase subunit |
| NC\_015761.1\|provirus\_3084741\_3115735\_2830 | 1068 | GENOMAD.125799.VP | PF06891 | P2 phage tail completion protein R (GpR) |
| NC\_015761.1\|provirus\_3084741\_3115735\_2831 | 855 | GENOMAD.042716.VV | COG4388 | Mu-like prophage I protein |
| NC\_015761.1\|provirus\_3084741\_3115735\_2832 | 1770 | GENOMAD.151636.VP | COG5484 | Uncharacterized protein YjcR, contains N-terminal HTH domain |
| NC\_015761.1\|provirus\_3084741\_3115735\_2833 | 1047 | NA | NA | NA |
| NC\_015761.1\|provirus\_3084741\_3115735\_2834 | 1005 | NA | NA | NA |
| NC\_015761.1\|provirus\_3084741\_3115735\_2835 | 549 | NA | NA | NA |
| NC\_015761.1\|provirus\_3084741\_3115735\_2836 | 732 | GENOMAD.226160.VP | NA | NA |
| NC\_015761.1\|provirus\_3084741\_3115735\_2837 | 441 | GENOMAD.137104.VV | NA | NA |
| NC\_015761.1\|provirus\_3084741\_3115735\_2838 | 2220 | GENOMAD.182861.VP | PF05840 | Bacteriophage replication gene A protein (GPA) |
| NC\_015761.1\|provirus\_3084741\_3115735\_2839 | 522 | GENOMAD.170058.VV | NA | NA |
| NC\_015761.1\|provirus\_3084741\_3115735\_2840 | 225 | NA | NA | NA |
| NC\_015761.1\|provirus\_3084741\_3115735\_2841 | 228 | GENOMAD.071646.VV | PF10809 | NA |
| NC\_015761.1\|provirus\_3084741\_3115735\_2842 | 201 | GENOMAD.167874.VV | NA | NA |
| NC\_015761.1\|provirus\_3084741\_3115735\_2843 | 228 | GENOMAD.087231.VV | PF10893 | Bacteriophage 186, Fil |
| NC\_015761.1\|provirus\_3084741\_3115735\_2844 | 510 | GENOMAD.170830.VP | PF06892 | Phage regulatory protein CII (CP76) |
| NC\_015761.1\|provirus\_3084741\_3115735\_2845 | 264 | GENOMAD.120542.VV | PF07618;COG3311;TIGR02405 | Putative transcription regulator (DUF1323) |
| NC\_015761.1\|provirus\_3084741\_3115735\_2846 | 579 | GENOMAD.069724.VV | PF16452 | Bacteriophage CI repressor C-terminal domain |
| NC\_015761.1\|provirus\_3084741\_3115735\_2847 | 1038 | GENOMAD.212346.VP | PF16452;PF06892;COG1974 | Bacteriophage CI repressor C-terminal domain; Phage regulatory protein CII (CP76) |
| NC\_015761.1\|provirus\_3084741\_3115735\_2848 | 336 | NA | NA | NA |

###### 2.13.2.2 Phage ID: NC\_015761.1|provirus\_847397\_862760

**Phage-Host Genome Ideogram:**

**Genes Annotation (geNomad):**

| gene | length | marker | annotation\_accessions | annotation\_description |
| --- | --- | --- | --- | --- |
| NC\_015761.1\|provirus\_847397\_862760\_745 | 240 | NA | NA | NA |
| NC\_015761.1\|provirus\_847397\_862760\_746 | 414 | GENOMAD.138177.VP | PF04717;PF18715;COG4540;TIGR01644 | Phage P2 baseplate assembly protein gpV |
| NC\_015761.1\|provirus\_847397\_862760\_747 | 360 | GENOMAD.121581.VP | PF04965;COG3628;K06903;TIGR03357 | Phage baseplate assembly protein W |
| NC\_015761.1\|provirus\_847397\_862760\_748 | 909 | GENOMAD.105501.VV | PF03434;COG3948 | Phage-related baseplate assembly protein |
| NC\_015761.1\|provirus\_847397\_862760\_749 | 606 | GENOMAD.114503.VV | PF09684;COG4385;TIGR01634 | Bacteriophage P2-related tail formation protein |
| NC\_015761.1\|provirus\_847397\_862760\_750 | 918 | GENOMAD.118528.VP | PF12571;COG5301 | Phage-related tail fibre protein |
| NC\_015761.1\|provirus\_847397\_862760\_751 | 1158 | NA | NA | NA |
| NC\_015761.1\|provirus\_847397\_862760\_752 | 465 | GENOMAD.168001.VP | PF16778 | Phage tail assembly chaperone protein |
| NC\_015761.1\|provirus\_847397\_862760\_753 | 255 | NA | NA | NA |
| NC\_015761.1\|provirus\_847397\_862760\_754 | 258 | NA | NA | NA |
| NC\_015761.1\|provirus\_847397\_862760\_755 | 186 | GENOMAD.120968.VV | COG3497 | Phage tail sheath protein FI |
| NC\_015761.1\|provirus\_847397\_862760\_756 | 303 | GENOMAD.135969.VP | PF10109 | Phage tail assembly chaperone proteins, E, or 41 or 14 |
| NC\_015761.1\|provirus\_847397\_862760\_757 | 1149 | GENOMAD.130201.VP | COG5283 | Phage-related tail protein |
| NC\_015761.1\|provirus\_847397\_862760\_758 | 2025 | GENOMAD.096277.VV | NA | NA |
| NC\_015761.1\|provirus\_847397\_862760\_759 | 387 | GENOMAD.116424.VV | PF06995;K06906;COG3499 | Phage protein U |
| NC\_015761.1\|provirus\_847397\_862760\_760 | 1053 | GENOMAD.129122.VP | PF05954;K06905;COG3500;TIGR03361 | Phage protein D |
| NC\_015761.1\|provirus\_847397\_862760\_761 | 219 | GENOMAD.222803.VP | PF04606;TIGR04165;COG1326 | Ogr/Delta-like zinc finger |
| NC\_015761.1\|provirus\_847397\_862760\_762 | 1686 | NA | NA | NA |
| NC\_015761.1\|provirus\_847397\_862760\_763 | 378 | NA | NA | NA |

###### 2.13.2.3 Phage ID: NC\_015761.1|provirus\_1007223\_1043740

**Phage-Host Genome Ideogram:**

**Genes Annotation (geNomad):**

| gene | length | marker | annotation\_accessions | annotation\_description |
| --- | --- | --- | --- | --- |
| NC\_015761.1\|provirus\_1007223\_1043740\_886 | 1020 | GENOMAD.212993.VP | PF14659;K21039 | Phage integrase, N-terminal SAM-like domain |
| NC\_015761.1\|provirus\_1007223\_1043740\_887 | 225 | GENOMAD.151670.VV | PF13986 | NA |
| NC\_015761.1\|provirus\_1007223\_1043740\_888 | 141 | GENOMAD.224144.VP | PF10798;K21975 | Biofilm development protein YmgB/AriR |
| NC\_015761.1\|provirus\_1007223\_1043740\_889 | 807 | NA | NA | NA |
| NC\_015761.1\|provirus\_1007223\_1043740\_890 | 849 | GENOMAD.212816.VV | PF20613;TIGR03843 | HipA-like kinase |
| NC\_015761.1\|provirus\_1007223\_1043740\_891 | 468 | GENOMAD.091484.VV | PF13411;COG3415;K22302;TIGR00721 | Transposase |
| NC\_015761.1\|provirus\_1007223\_1043740\_892 | 228 | GENOMAD.151341.VV | PF11242;TIGR00673;COG5606;K18830 | cyanase |
| NC\_015761.1\|provirus\_1007223\_1043740\_893 | 321 | GENOMAD.069430.VV | PF18010;TIGR00721;COG4220 | Cry35Ab1 HTH C-terminal domain |
| NC\_015761.1\|provirus\_1007223\_1043740\_894 | 906 | GENOMAD.208204.VP | PF04492;TIGR01610 | phage replication protein O, N-terminal domain |
| NC\_015761.1\|provirus\_1007223\_1043740\_895 | 693 | GENOMAD.194512.VV | PF06992 | NA |
| NC\_015761.1\|provirus\_1007223\_1043740\_896 | 258 | GENOMAD.072808.VV | PF15944 | NA |
| NC\_015761.1\|provirus\_1007223\_1043740\_897 | 912 | GENOMAD.068933.VV | NA | NA |
| NC\_015761.1\|provirus\_1007223\_1043740\_898 | 327 | NA | NA | NA |
| NC\_015761.1\|provirus\_1007223\_1043740\_899 | 108 | GENOMAD.226031.VP | NA | NA |
| NC\_015761.1\|provirus\_1007223\_1043740\_900 | 432 | GENOMAD.213484.VP | PF05509;COG4877 | Plasmid stability protein |
| NC\_015761.1\|provirus\_1007223\_1043740\_901 | 168 | GENOMAD.168832.VV | NA | NA |
| NC\_015761.1\|provirus\_1007223\_1043740\_902 | 915 | GENOMAD.222649.VP | PF10548;PF10547;COG3617 | Prophage antirepressor |
| NC\_015761.1\|provirus\_1007223\_1043740\_903 | 747 | GENOMAD.167364.VP | NA | NA |
| NC\_015761.1\|provirus\_1007223\_1043740\_904 | 234 | NA | NA | NA |
| NC\_015761.1\|provirus\_1007223\_1043740\_905 | 603 | GENOMAD.040195.VV | PF07105 | NA |
| NC\_015761.1\|provirus\_1007223\_1043740\_906 | 672 | GENOMAD.058019.VV | PF06323 | Phage antitermination protein Q |
| NC\_015761.1\|provirus\_1007223\_1043740\_907 | 573 | GENOMAD.209828.VP | PF10543;COG3646 | ORF6N domain |
| NC\_015761.1\|provirus\_1007223\_1043740\_908 | 501 | NA | NA | NA |
| NC\_015761.1\|provirus\_1007223\_1043740\_909 | 180 | GENOMAD.175300.VV | NA | NA |
| NC\_015761.1\|provirus\_1007223\_1043740\_910 | 426 | NA | NA | NA |
| NC\_015761.1\|provirus\_1007223\_1043740\_911 | 744 | NA | NA | NA |
| NC\_015761.1\|provirus\_1007223\_1043740\_912 | 390 | GENOMAD.222896.VP | PF16931 | Putative phage holin |
| NC\_015761.1\|provirus\_1007223\_1043740\_913 | 282 | GENOMAD.214895.VP | PF05449 | Putative 3TM holin, Phage\_holin\_3 |
| NC\_015761.1\|provirus\_1007223\_1043740\_914 | 615 | GENOMAD.021660.VV | COG3179;K18950 | Predicted chitinase |
| NC\_015761.1\|provirus\_1007223\_1043740\_915 | 543 | GENOMAD.108518.VV | NA | NA |
| NC\_015761.1\|provirus\_1007223\_1043740\_916 | 531 | GENOMAD.068419.VV | PF10549 | ORF11CD3 domain |
| NC\_015761.1\|provirus\_1007223\_1043740\_917 | 360 | GENOMAD.062288.VV | PF10721 | NA |
| NC\_015761.1\|provirus\_1007223\_1043740\_918 | 411 | NA | NA | NA |
| NC\_015761.1\|provirus\_1007223\_1043740\_919 | 306 | GENOMAD.210890.VV | NA | NA |
| NC\_015761.1\|provirus\_1007223\_1043740\_920 | 351 | GENOMAD.179073.VP | NA | NA |
| NC\_015761.1\|provirus\_1007223\_1043740\_921 | 258 | NA | NA | NA |
| NC\_015761.1\|provirus\_1007223\_1043740\_922 | 1113 | NA | NA | NA |
| NC\_015761.1\|provirus\_1007223\_1043740\_923 | 117 | NA | NA | NA |
| NC\_015761.1\|provirus\_1007223\_1043740\_924 | 579 | GENOMAD.222575.VP | PF06416 | Effector protein NleG |
| NC\_015761.1\|provirus\_1007223\_1043740\_925 | 2349 | NA | NA | NA |
| NC\_015761.1\|provirus\_1007223\_1043740\_926 | 1131 | NA | NA | NA |
| NC\_015761.1\|provirus\_1007223\_1043740\_927 | 198 | NA | NA | NA |
| NC\_015761.1\|provirus\_1007223\_1043740\_928 | 330 | GENOMAD.182552.PC | PF15781;COG3668;TIGR00053;K06218 | Plasmid stabilization system protein ParE |
| NC\_015761.1\|provirus\_1007223\_1043740\_929 | 231 | NA | NA | NA |
| NC\_015761.1\|provirus\_1007223\_1043740\_930 | 294 | GENOMAD.221919.VP | PF00589 | Phage integrase family |
| NC\_015761.1\|provirus\_1007223\_1043740\_931 | 579 | GENOMAD.222575.VP | PF06416 | Effector protein NleG |
| NC\_015761.1\|provirus\_1007223\_1043740\_932 | 348 | NA | NA | NA |
| NC\_015761.1\|provirus\_1007223\_1043740\_933 | 855 | GENOMAD.120636.VV | PF09612;TIGR02192 | protein YibB |
| NC\_015761.1\|provirus\_1007223\_1043740\_934 | 231 | NA | NA | NA |
| NC\_015761.1\|provirus\_1007223\_1043740\_935 | 2808 | NA | NA | NA |

##### 2.13.3 vOTUs

When more than 1 phage is detected in the host genome, we perform a
clustering step using the tool dRep.

A threshold of 0.95 was applied to the ANI similarity index to define
clusters of virus operational taxonomic units (vOTUs).

**Final cluster designations**

| genomad\_summary$seq\_name | genome | secondary\_cluster | threshold | cluster\_method | comparison\_algorithm | primary\_cluster |
| --- | --- | --- | --- | --- | --- | --- |
| NC\_015761.1\|provirus\_3084741\_3115735 | sequence\_000000.fasta.fasta | 1\_0 | 0.05 | average | ANImf | 1 |
| NC\_015761.1\|provirus\_847397\_862760 | sequence\_000001.fasta.fasta | 2\_0 | 0.05 | average | ANImf | 2 |
| NC\_015761.1\|provirus\_1007223\_1043740 | sequence\_000002.fasta.fasta | 3\_0 | 0.05 | average | ANImf | 3 |

**Primary clustering plot**

##### 2.13.4 Virulence Genes

Screening of virulence genes present in the prophage contigs.

| sequence | start | end | strand | gene | coverage | coverage\_map | gaps | percent\_coverage | percent\_identity | database | accession | product | resistance |
| --- | --- | --- | --- | --- | --- | --- | --- | --- | --- | --- | --- | --- | --- |
| NC\_015761.1\|provirus\_1007223\_1043740 | 24347 | 26694 |  | sopA | 1-2348/2349 | =============== | 0/0 | 99.96 | 89.91 | vfdb | NP\_461011 | (sopA) type III secretion system effector SopA E3 ubiquitin ligase [TTSS(SPI-1 encode) (VF0116)] [Salmonella enterica subsp. enterica serovar Typhimurium str. LT2] | NA |

##### 2.13.5 Prophage-Host Prediction

Prediction of potential hosts for the prophage contigs.

| virus | host\_genome | host\_taxonomy | main\_method | confidence\_score | additional\_methods |
| --- | --- | --- | --- | --- | --- |
| NC\_015761.1\|provirus\_1007223\_1043740 | GB\_GCA\_900446925.1 | d\_\_Bacteria;p\_\_Pseudomonadota;c\_\_Gammaproteobacteria;o\_\_Enterobacterales;f\_\_Enterobacteriaceae;g\_\_Citrobacter\_B;s\_\_Citrobacter\_B koseri | blast | 96.9 | iPHoP-RF;82.80 |
| NC\_015761.1\|provirus\_1007223\_1043740 | GB\_GCA\_900478215.1 | d\_\_Bacteria;p\_\_Pseudomonadota;c\_\_Gammaproteobacteria;o\_\_Enterobacterales;f\_\_Enterobacteriaceae;g\_\_Salmonella;s\_\_Salmonella houtenae | blast | 96.9 | iPHoP-RF;83.30 |
| NC\_015761.1\|provirus\_1007223\_1043740 | RS\_GCF\_000006945.2 | d\_\_Bacteria;p\_\_Pseudomonadota;c\_\_Gammaproteobacteria;o\_\_Enterobacterales;f\_\_Enterobacteriaceae;g\_\_Salmonella;s\_\_Salmonella enterica | blast | 96.9 | iPHoP-RF;86.70 |
| NC\_015761.1\|provirus\_1007223\_1043740 | RS\_GCF\_000252995.1 | d\_\_Bacteria;p\_\_Pseudomonadota;c\_\_Gammaproteobacteria;o\_\_Enterobacterales;f\_\_Enterobacteriaceae;g\_\_Salmonella;s\_\_Salmonella bongori | blast | 96.9 | iPHoP-RF;84.20 |
| NC\_015761.1\|provirus\_1007223\_1043740 | RS\_GCF\_008692785.1 | d\_\_Bacteria;p\_\_Pseudomonadota;c\_\_Gammaproteobacteria;o\_\_Enterobacterales;f\_\_Enterobacteriaceae;g\_\_Salmonella;s\_\_Salmonella diarizonae | blast | 96.9 | iPHoP-RF;87.60 |
| NC\_015761.1\|provirus\_1007223\_1043740 | RS\_GCF\_008692845.1 | d\_\_Bacteria;p\_\_Pseudomonadota;c\_\_Gammaproteobacteria;o\_\_Enterobacterales;f\_\_Enterobacteriaceae;g\_\_Salmonella;s\_\_Salmonella arizonae | blast | 96.9 | iPHoP-RF;87.60 |
| NC\_015761.1\|provirus\_1007223\_1043740 | RS\_GCF\_006874705.1 | d\_\_Bacteria;p\_\_Pseudomonadota;c\_\_Gammaproteobacteria;o\_\_Enterobacterales;f\_\_Enterobacteriaceae;g\_\_Leclercia;s\_\_Leclercia adecarboxylata\_C | blast | 92.7 | None |
| NC\_015761.1\|provirus\_3084741\_3115735 | RS\_GCF\_000006945.2 | d\_\_Bacteria;p\_\_Pseudomonadota;c\_\_Gammaproteobacteria;o\_\_Enterobacterales;f\_\_Enterobacteriaceae;g\_\_Salmonella;s\_\_Salmonella enterica | CRISPR | 98.5 | blast;96.80 iPHoP-RF;89.80 |
| NC\_015761.1\|provirus\_3084741\_3115735 | RS\_GCF\_008692785.1 | d\_\_Bacteria;p\_\_Pseudomonadota;c\_\_Gammaproteobacteria;o\_\_Enterobacterales;f\_\_Enterobacteriaceae;g\_\_Salmonella;s\_\_Salmonella diarizonae | CRISPR | 98.1 | iPHoP-RF;90.80 |
| NC\_015761.1\|provirus\_3084741\_3115735 | RS\_GCF\_000252995.1 | d\_\_Bacteria;p\_\_Pseudomonadota;c\_\_Gammaproteobacteria;o\_\_Enterobacterales;f\_\_Enterobacteriaceae;g\_\_Salmonella;s\_\_Salmonella bongori | blast | 96.9 | iPHoP-RF;90.10 |
| NC\_015761.1\|provirus\_3084741\_3115735 | RS\_GCF\_002918555.1 | d\_\_Bacteria;p\_\_Pseudomonadota;c\_\_Gammaproteobacteria;o\_\_Enterobacterales;f\_\_Enterobacteriaceae;g\_\_Citrobacter\_C;s\_\_Citrobacter\_C amalonaticus\_A | blast | 93.2 | None |
| NC\_015761.1\|provirus\_3084741\_3115735 | GB\_GCA\_900446925.1 | d\_\_Bacteria;p\_\_Pseudomonadota;c\_\_Gammaproteobacteria;o\_\_Enterobacterales;f\_\_Enterobacteriaceae;g\_\_Citrobacter\_B;s\_\_Citrobacter\_B koseri | iPHoP-RF | 92.1 | CRISPR;67.50 |
| NC\_015761.1\|provirus\_847397\_862760 | RS\_GCF\_003697165.2 | d\_\_Bacteria;p\_\_Pseudomonadota;c\_\_Gammaproteobacteria;o\_\_Enterobacterales;f\_\_Enterobacteriaceae;g\_\_Escherichia;s\_\_Escherichia coli | blast | 96.9 | iPHoP-RF;88.90 |
| NC\_015761.1\|provirus\_847397\_862760 | RS\_GCF\_000759775.1 | d\_\_Bacteria;p\_\_Pseudomonadota;c\_\_Gammaproteobacteria;o\_\_Enterobacterales;f\_\_Enterobacteriaceae;g\_\_Escherichia;s\_\_Escherichia albertii | blast | 96.3 | iPHoP-RF;86.70 |
| NC\_015761.1\|provirus\_847397\_862760 | RS\_GCF\_001729745.1 | d\_\_Bacteria;p\_\_Pseudomonadota;c\_\_Gammaproteobacteria;o\_\_Enterobacterales;f\_\_Enterobacteriaceae;g\_\_Enterobacter;s\_\_Enterobacter hormaechei\_A | blast | 96.2 | iPHoP-RF;73.00 |
| NC\_015761.1\|provirus\_847397\_862760 | RS\_GCF\_000252995.1 | d\_\_Bacteria;p\_\_Pseudomonadota;c\_\_Gammaproteobacteria;o\_\_Enterobacterales;f\_\_Enterobacteriaceae;g\_\_Salmonella;s\_\_Salmonella bongori | blast | 93.9 | iPHoP-RF;71.50 |
| NC\_015761.1\|provirus\_847397\_862760 | GB\_GCA\_900478215.1 | d\_\_Bacteria;p\_\_Pseudomonadota;c\_\_Gammaproteobacteria;o\_\_Enterobacterales;f\_\_Enterobacteriaceae;g\_\_Salmonella;s\_\_Salmonella houtenae | blast | 91.9 | iPHoP-RF;73.70 |
| NC\_015761.1\|provirus\_847397\_862760 | RS\_GCF\_000006945.2 | d\_\_Bacteria;p\_\_Pseudomonadota;c\_\_Gammaproteobacteria;o\_\_Enterobacterales;f\_\_Enterobacteriaceae;g\_\_Salmonella;s\_\_Salmonella enterica | blast | 91.7 | iPHoP-RF;88.50 |
| NC\_015761.1\|provirus\_847397\_862760 | RS\_GCF\_008692845.1 | d\_\_Bacteria;p\_\_Pseudomonadota;c\_\_Gammaproteobacteria;o\_\_Enterobacterales;f\_\_Enterobacteriaceae;g\_\_Salmonella;s\_\_Salmonella arizonae | blast | 91.7 | iPHoP-RF;84.50 |
| NC\_015761.1\|provirus\_847397\_862760 | RS\_GCF\_002900365.1 | d\_\_Bacteria;p\_\_Pseudomonadota;c\_\_Gammaproteobacteria;o\_\_Enterobacterales;f\_\_Enterobacteriaceae;g\_\_Escherichia;s\_\_Escherichia marmotae | blast | 90.0 | iPHoP-RF;87.00 |

## 2.14 NC\_017033

##### 2.14.1 Host Genome

**GTDB-Tk taxonomy**:

classification

d\_\_Bacteria;p\_\_Pseudomonadota;c\_\_Gammaproteobacteria;o\_\_Xanthomonadales;f\_\_Rhodanobacteraceae;g\_\_Frateuria;s\_\_Frateuria
aurantia

**CheckM2 Summary**:

| total\_contigs | contig\_n50 | completeness | contamination |
| --- | --- | --- | --- |
| 1 | 3603458 | 99.99 | 0.04 |

**Defense-Finder Systems**:

| sys\_id | type | subtype | sys\_beg | sys\_end | protein\_in\_syst | genes\_count | name\_of\_profiles\_in\_sys |
| --- | --- | --- | --- | --- | --- | --- | --- |
| NC\_017033.1\_PrrC\_3 | PrrC | PrrC | NC\_017033.1\_586 | NC\_017033.1\_589 | NC\_017033.1\_586,NC\_017033.1\_587,NC\_017033.1\_588,NC\_017033.1\_589 | 4 | PrrC\_\_EcoprrI,PrrC\_\_PrrC,RM\_\_Type\_I\_REases\_FAM\_0.einsi\_trimmed,RM\_\_Type\_I\_S\_02 |
| NC\_017033.1\_DS-6\_1 | DS-6 | DS-6 | NC\_017033.1\_1067 | NC\_017033.1\_1068 | NC\_017033.1\_1067,NC\_017033.1\_1068 | 2 | DS-6\_\_DS-6A,DS-6\_\_DS-6B |
| NC\_017033.1\_Mokosh\_TypeII\_2 | Mokosh | Mokosh\_TypeII | NC\_017033.1\_1863 | NC\_017033.1\_1863 | NC\_017033.1\_1863 | 1 | Mokosh\_TypeII\_\_MkoC |

**Genomad and CheckV Summary**:

| seq\_name | length | gene\_count | viral\_genes | checkv\_quality | miuvig\_quality | taxonomy | topology | coordinates |
| --- | --- | --- | --- | --- | --- | --- | --- | --- |
| NC\_017033.1\|provirus\_1557694\_1600481 | 42788 | 71 | 28 | Medium-quality | Genome-fragment | Viruses;Duplodnaviria;Heunggongvirae;Uroviricota;Caudoviricetes;; | Provirus | 1557694-1600481 |
| NC\_017033.1\|provirus\_1050333\_1094910 | 44578 | 68 | 28 | High-quality | High-quality | Viruses;Duplodnaviria;Heunggongvirae;Uroviricota;Caudoviricetes;; | Provirus | 1050333-1094910 |
| NC\_017033.1\|provirus\_2089294\_2107522 | 18229 | 25 | 3 | Low-quality | Genome-fragment | Viruses;Duplodnaviria;Heunggongvirae;Uroviricota;Caudoviricetes;; | Provirus | 2089294-2107522 |

**Host Genome Ideogram with Phages**:

In
this circular plot, **“c”** indicates the contig, and the
number that follows (e.g., **c1**) represents the contig
number. If multiple contigs are present in the genome, each will be
shown with a distinct label (e.g., **c1**,
**c2**, etc.).

##### 2.14.2 Prophages

**Select prophage to show:**

###### 2.14.2.1 Phage ID: NC\_017033.1|provirus\_1557694\_1600481

**Phage-Host Genome Ideogram:**

**Genes Annotation (geNomad):**

| gene | length | marker | annotation\_accessions | annotation\_description |
| --- | --- | --- | --- | --- |
| NC\_017033.1\|provirus\_1557694\_1600481\_1419 | 288 | NA | NA | NA |
| NC\_017033.1\|provirus\_1557694\_1600481\_1420 | 189 | NA | NA | NA |
| NC\_017033.1\|provirus\_1557694\_1600481\_1421 | 309 | GENOMAD.060822.VV | NA | NA |
| NC\_017033.1\|provirus\_1557694\_1600481\_1422 | 369 | NA | NA | NA |
| NC\_017033.1\|provirus\_1557694\_1600481\_1423 | 279 | NA | NA | NA |
| NC\_017033.1\|provirus\_1557694\_1600481\_1424 | 330 | NA | NA | NA |
| NC\_017033.1\|provirus\_1557694\_1600481\_1425 | 1656 | NA | NA | NA |
| NC\_017033.1\|provirus\_1557694\_1600481\_1426 | 333 | NA | NA | NA |
| NC\_017033.1\|provirus\_1557694\_1600481\_1427 | 228 | NA | NA | NA |
| NC\_017033.1\|provirus\_1557694\_1600481\_1428 | 1755 | GENOMAD.151126.VP | NA | NA |
| NC\_017033.1\|provirus\_1557694\_1600481\_1429 | 741 | NA | NA | NA |
| NC\_017033.1\|provirus\_1557694\_1600481\_1430 | 285 | NA | NA | NA |
| NC\_017033.1\|provirus\_1557694\_1600481\_1431 | 195 | NA | NA | NA |
| NC\_017033.1\|provirus\_1557694\_1600481\_1432 | 216 | NA | NA | NA |
| NC\_017033.1\|provirus\_1557694\_1600481\_1433 | 126 | NA | NA | NA |
| NC\_017033.1\|provirus\_1557694\_1600481\_1434 | 834 | GENOMAD.115107.VV | NA | NA |
| NC\_017033.1\|provirus\_1557694\_1600481\_1435 | 387 | GENOMAD.094188.VV | NA | NA |
| NC\_017033.1\|provirus\_1557694\_1600481\_1436 | 237 | GENOMAD.189851.VV | NA | NA |
| NC\_017033.1\|provirus\_1557694\_1600481\_1437 | 393 | NA | NA | NA |
| NC\_017033.1\|provirus\_1557694\_1600481\_1438 | 753 | NA | NA | NA |
| NC\_017033.1\|provirus\_1557694\_1600481\_1439 | 603 | GENOMAD.205774.CP | PF08000;PF20612 | Bacterial PH domain; SHOCT domain |
| NC\_017033.1\|provirus\_1557694\_1600481\_1440 | 327 | NA | NA | NA |
| NC\_017033.1\|provirus\_1557694\_1600481\_1441 | 243 | NA | NA | NA |
| NC\_017033.1\|provirus\_1557694\_1600481\_1442 | 294 | NA | NA | NA |
| NC\_017033.1\|provirus\_1557694\_1600481\_1443 | 210 | NA | NA | NA |
| NC\_017033.1\|provirus\_1557694\_1600481\_1444 | 303 | GENOMAD.045996.VV | PF09012;TIGR02702;COG1777 | FeoC like transcriptional regulator |
| NC\_017033.1\|provirus\_1557694\_1600481\_1445 | 876 | NA | NA | NA |
| NC\_017033.1\|provirus\_1557694\_1600481\_1446 | 492 | GENOMAD.166457.VP | PF06992 | Replication protein P |
| NC\_017033.1\|provirus\_1557694\_1600481\_1447 | 207 | GENOMAD.063929.VV | PF05810;COG4068 | Predicted nucleic acid-binding protein, contains Zn-ribbon domain |
| NC\_017033.1\|provirus\_1557694\_1600481\_1448 | 198 | NA | NA | NA |
| NC\_017033.1\|provirus\_1557694\_1600481\_1449 | 195 | NA | NA | NA |
| NC\_017033.1\|provirus\_1557694\_1600481\_1450 | 450 | GENOMAD.061349.VV | PF05772 | NinB protein |
| NC\_017033.1\|provirus\_1557694\_1600481\_1451 | 348 | GENOMAD.036842.VV | PF07102 | Putative nuclease YbcO |
| NC\_017033.1\|provirus\_1557694\_1600481\_1452 | 174 | NA | NA | NA |
| NC\_017033.1\|provirus\_1557694\_1600481\_1453 | 336 | GENOMAD.040593.VV | NA | NA |
| NC\_017033.1\|provirus\_1557694\_1600481\_1454 | 618 | GENOMAD.069895.VV | PF17302;TIGR02642 | Tryptophan RNA-binding attenuator protein inhibitory protein |
| NC\_017033.1\|provirus\_1557694\_1600481\_1455 | 264 | NA | NA | NA |
| NC\_017033.1\|provirus\_1557694\_1600481\_1456 | 504 | GENOMAD.044304.VV | NA | NA |
| NC\_017033.1\|provirus\_1557694\_1600481\_1457 | 300 | NA | NA | NA |
| NC\_017033.1\|provirus\_1557694\_1600481\_1458 | 267 | NA | NA | NA |
| NC\_017033.1\|provirus\_1557694\_1600481\_1459 | 342 | NA | NA | NA |
| NC\_017033.1\|provirus\_1557694\_1600481\_1460 | 336 | NA | NA | NA |
| NC\_017033.1\|provirus\_1557694\_1600481\_1461 | 264 | GENOMAD.077615.VV | NA | NA |
| NC\_017033.1\|provirus\_1557694\_1600481\_1462 | 672 | GENOMAD.076022.VV | NA | NA |
| NC\_017033.1\|provirus\_1557694\_1600481\_1463 | 1557 | GENOMAD.013124.VV | PF13262;COG5410 | NA |
| NC\_017033.1\|provirus\_1557694\_1600481\_1464 | 1539 | GENOMAD.003432.VV | PF06381;K09961;TIGR01555;COG3567 | phage-related protein, HI1409 family |
| NC\_017033.1\|provirus\_1557694\_1600481\_1465 | 654 | GENOMAD.083949.VV | PF04233;COG2369;TIGR01641 | Uncharacterized conserved protein, contains phage Mu gpF-like domain |
| NC\_017033.1\|provirus\_1557694\_1600481\_1466 | 180 | NA | NA | NA |
| NC\_017033.1\|provirus\_1557694\_1600481\_1467 | 1311 | GENOMAD.099067.VP | PF09979;K09960;COG3566 | NA |
| NC\_017033.1\|provirus\_1557694\_1600481\_1468 | 495 | GENOMAD.033203.VV | PF23982 | Putative phage cement protein |
| NC\_017033.1\|provirus\_1557694\_1600481\_1469 | 1008 | GENOMAD.085797.VV | PF09950;COG4834 | Encapsulating protein for peroxidase |
| NC\_017033.1\|provirus\_1557694\_1600481\_1470 | 225 | GENOMAD.170487.VV | NA | NA |
| NC\_017033.1\|provirus\_1557694\_1600481\_1471 | 396 | GENOMAD.005453.VV | PF11863;COG4386 | Mu-like prophage tail sheath protein gpL |
| NC\_017033.1\|provirus\_1557694\_1600481\_1472 | 543 | GENOMAD.008778.VV | PF05069;COG5005;TIGR01635 | NA |
| NC\_017033.1\|provirus\_1557694\_1600481\_1473 | 384 | GENOMAD.040078.VV | NA | NA |
| NC\_017033.1\|provirus\_1557694\_1600481\_1474 | 555 | GENOMAD.023692.VV | PF23961 | Phage neck terminator protein |
| NC\_017033.1\|provirus\_1557694\_1600481\_1475 | 1485 | GENOMAD.005453.VV | PF11863;COG4386 | Mu-like prophage tail sheath protein gpL |
| NC\_017033.1\|provirus\_1557694\_1600481\_1476 | 441 | GENOMAD.017010.VV | PF11681 | Bacteriophage KPP10, Structural protein ORF10 |
| NC\_017033.1\|provirus\_1557694\_1600481\_1477 | 390 | GENOMAD.055746.VV | PF10876 | Phage tail assembly chaperone protein, TAC |
| NC\_017033.1\|provirus\_1557694\_1600481\_1478 | 2253 | GENOMAD.015777.VV | NA | NA |
| NC\_017033.1\|provirus\_1557694\_1600481\_1479 | 621 | GENOMAD.017318.VV | COG3499 | NA |
| NC\_017033.1\|provirus\_1557694\_1600481\_1480 | 315 | GENOMAD.105312.VV | PF22479 | Cyanophage baseplate Pam3 plug gp18 |
| NC\_017033.1\|provirus\_1557694\_1600481\_1481 | 948 | GENOMAD.123309.VP | PF21829 | NA |
| NC\_017033.1\|provirus\_1557694\_1600481\_1482 | 660 | GENOMAD.006053.VV | PF18352;COG4540;TIGR01644 | Phage protein Gp138 N-terminal domain |
| NC\_017033.1\|provirus\_1557694\_1600481\_1483 | 348 | GENOMAD.009002.VV | PF10934;COG3628 | Phage baseplate assembly protein W |
| NC\_017033.1\|provirus\_1557694\_1600481\_1484 | 1179 | GENOMAD.005285.VV | PF03434;COG3299 | Uncharacterized phage protein gp47/JayE |
| NC\_017033.1\|provirus\_1557694\_1600481\_1485 | 585 | GENOMAD.008482.VV | PF11041;TIGR02242 | Bacteriophage Mu-like, Gp48 |
| NC\_017033.1\|provirus\_1557694\_1600481\_1486 | 1809 | GENOMAD.051520.VV | NA | NA |
| NC\_017033.1\|provirus\_1557694\_1600481\_1487 | 486 | NA | NA | NA |
| NC\_017033.1\|provirus\_1557694\_1600481\_1488 | 177 | NA | NA | NA |
| NC\_017033.1\|provirus\_1557694\_1600481\_1489 | 972 | NA | NA | NA |

###### 2.14.2.2 Phage ID: NC\_017033.1|provirus\_1050333\_1094910

**Phage-Host Genome Ideogram:**

**Genes Annotation (geNomad):**

| gene | length | marker | annotation\_accessions | annotation\_description |
| --- | --- | --- | --- | --- |
| NC\_017033.1\|provirus\_1050333\_1094910\_928 | 1320 | NA | NA | NA |
| NC\_017033.1\|provirus\_1050333\_1094910\_929 | 546 | NA | NA | NA |
| NC\_017033.1\|provirus\_1050333\_1094910\_930 | 204 | NA | NA | NA |
| NC\_017033.1\|provirus\_1050333\_1094910\_931 | 1020 | NA | NA | NA |
| NC\_017033.1\|provirus\_1050333\_1094910\_932 | 219 | GENOMAD.180717.VC | PF13986 | NA |
| NC\_017033.1\|provirus\_1050333\_1094910\_933 | 249 | GENOMAD.104567.VV | NA | NA |
| NC\_017033.1\|provirus\_1050333\_1094910\_934 | 228 | NA | NA | NA |
| NC\_017033.1\|provirus\_1050333\_1094910\_935 | 282 | NA | NA | NA |
| NC\_017033.1\|provirus\_1050333\_1094910\_936 | 258 | GENOMAD.178404.VP | NA | NA |
| NC\_017033.1\|provirus\_1050333\_1094910\_937 | 684 | GENOMAD.171154.VP | PF13986;PF05551 | NA |
| NC\_017033.1\|provirus\_1050333\_1094910\_938 | 351 | NA | NA | NA |
| NC\_017033.1\|provirus\_1050333\_1094910\_939 | 252 | NA | NA | NA |
| NC\_017033.1\|provirus\_1050333\_1094910\_940 | 480 | NA | NA | NA |
| NC\_017033.1\|provirus\_1050333\_1094910\_941 | 303 | NA | NA | NA |
| NC\_017033.1\|provirus\_1050333\_1094910\_942 | 276 | NA | NA | NA |
| NC\_017033.1\|provirus\_1050333\_1094910\_943 | 129 | NA | NA | NA |
| NC\_017033.1\|provirus\_1050333\_1094910\_944 | 690 | NA | NA | NA |
| NC\_017033.1\|provirus\_1050333\_1094910\_945 | 279 | NA | NA | NA |
| NC\_017033.1\|provirus\_1050333\_1094910\_946 | 432 | NA | NA | NA |
| NC\_017033.1\|provirus\_1050333\_1094910\_947 | 192 | NA | NA | NA |
| NC\_017033.1\|provirus\_1050333\_1094910\_948 | 294 | NA | NA | NA |
| NC\_017033.1\|provirus\_1050333\_1094910\_949 | 168 | NA | NA | NA |
| NC\_017033.1\|provirus\_1050333\_1094910\_950 | 258 | GENOMAD.209492.VC | NA | NA |
| NC\_017033.1\|provirus\_1050333\_1094910\_951 | 564 | GENOMAD.193207.VC | NA | NA |
| NC\_017033.1\|provirus\_1050333\_1094910\_952 | 357 | NA | NA | NA |
| NC\_017033.1\|provirus\_1050333\_1094910\_953 | 696 | NA | NA | NA |
| NC\_017033.1\|provirus\_1050333\_1094910\_954 | 213 | NA | NA | NA |
| NC\_017033.1\|provirus\_1050333\_1094910\_955 | 510 | GENOMAD.177794.VC | PF09639 | YjcQ protein |
| NC\_017033.1\|provirus\_1050333\_1094910\_956 | 627 | GENOMAD.120202.VV | PF06892 | NA |
| NC\_017033.1\|provirus\_1050333\_1094910\_957 | 990 | GENOMAD.105893.VV | PF07120;COG3756 | Uncharacterized conserved protein YdaU, DUF1376 family |
| NC\_017033.1\|provirus\_1050333\_1094910\_958 | 663 | NA | NA | NA |
| NC\_017033.1\|provirus\_1050333\_1094910\_959 | 201 | NA | NA | NA |
| NC\_017033.1\|provirus\_1050333\_1094910\_960 | 507 | NA | NA | NA |
| NC\_017033.1\|provirus\_1050333\_1094910\_961 | 390 | GENOMAD.022482.VV | PF16786 | Recombination enhancement, RecA-dependent nuclease |
| NC\_017033.1\|provirus\_1050333\_1094910\_962 | 498 | GENOMAD.159849.VP | PF07102;PF13264 | Putative nuclease YbcO |
| NC\_017033.1\|provirus\_1050333\_1094910\_963 | 384 | GENOMAD.099176.VV | NA | NA |
| NC\_017033.1\|provirus\_1050333\_1094910\_964 | 684 | GENOMAD.069895.VV | PF17302;TIGR02642 | Tryptophan RNA-binding attenuator protein inhibitory protein |
| NC\_017033.1\|provirus\_1050333\_1094910\_965 | 306 | NA | NA | NA |
| NC\_017033.1\|provirus\_1050333\_1094910\_966 | 315 | NA | NA | NA |
| NC\_017033.1\|provirus\_1050333\_1094910\_967 | 507 | GENOMAD.219399.VP | PF10549;COG3646;TIGR02681 | Phage regulatory protein Rha |
| NC\_017033.1\|provirus\_1050333\_1094910\_968 | 219 | NA | NA | NA |
| NC\_017033.1\|provirus\_1050333\_1094910\_969 | 477 | GENOMAD.083189.VV | PF00959;COG3772;K01185 | Phage-related lysozyme (muramidase), GH24 family |
| NC\_017033.1\|provirus\_1050333\_1094910\_970 | 483 | GENOMAD.077452.VV | PF03245;K14744 | Bacteriophage Rz lysis protein |
| NC\_017033.1\|provirus\_1050333\_1094910\_971 | 342 | NA | NA | NA |
| NC\_017033.1\|provirus\_1050333\_1094910\_972 | 531 | GENOMAD.161251.VP | PF07471;K22014;COG4220 | Phage DNA packaging protein, Nu1 subunit of terminase |
| NC\_017033.1\|provirus\_1050333\_1094910\_973 | 1905 | GENOMAD.129396.VP | PF20454 | Terminase large subunit gpA, endonuclease domain |
| NC\_017033.1\|provirus\_1050333\_1094910\_974 | 237 | NA | NA | NA |
| NC\_017033.1\|provirus\_1050333\_1094910\_975 | 1602 | GENOMAD.019804.VV | PF05136;TIGR01539;COG5511 | phage portal protein, lambda family |
| NC\_017033.1\|provirus\_1050333\_1094910\_976 | 1230 | GENOMAD.160855.VP | PF00574;TIGR00706;COG0740 | signal peptide peptidase SppA, 36K type |
| NC\_017033.1\|provirus\_1050333\_1094910\_977 | 417 | GENOMAD.191487.VP | PF02924 | Bacteriophage lambda head decoration protein D |
| NC\_017033.1\|provirus\_1050333\_1094910\_978 | 1002 | GENOMAD.124233.VP | PF03864 | Phage major capsid protein E |
| NC\_017033.1\|provirus\_1050333\_1094910\_979 | 309 | GENOMAD.092349.VV | PF13856 | ATP-binding sugar transporter from pro-phage |
| NC\_017033.1\|provirus\_1050333\_1094910\_980 | 597 | GENOMAD.066423.VV | PF06763 | Prophage minor tail protein Z (GPZ) |
| NC\_017033.1\|provirus\_1050333\_1094910\_981 | 582 | GENOMAD.065270.VV | PF09646 | Gp37 protein |
| NC\_017033.1\|provirus\_1050333\_1094910\_982 | 588 | GENOMAD.162066.VP | PF04717;COG4540;TIGR01644 | Phage P2 baseplate assembly protein gpV |
| NC\_017033.1\|provirus\_1050333\_1094910\_983 | 357 | GENOMAD.099345.VV | NA | NA |
| NC\_017033.1\|provirus\_1050333\_1094910\_984 | 330 | GENOMAD.136378.VP | PF05136;PF04965;TIGR01539;K06903;COG5511 | phage portal protein, lambda family |
| NC\_017033.1\|provirus\_1050333\_1094910\_985 | 885 | GENOMAD.105501.VV | PF03434;COG3948 | Phage-related baseplate assembly protein |
| NC\_017033.1\|provirus\_1050333\_1094910\_986 | 546 | GENOMAD.114503.VV | PF09684;COG4385;TIGR01634 | Bacteriophage P2-related tail formation protein |
| NC\_017033.1\|provirus\_1050333\_1094910\_987 | 1968 | GENOMAD.208390.VP | COG5301 | Phage-related tail fibre protein |
| NC\_017033.1\|provirus\_1050333\_1094910\_988 | 486 | NA | NA | NA |
| NC\_017033.1\|provirus\_1050333\_1094910\_989 | 1221 | GENOMAD.102019.VV | PF10758;COG3497 | Phage tail sheath protein FI |
| NC\_017033.1\|provirus\_1050333\_1094910\_990 | 504 | GENOMAD.099825.VV | PF04985;K06908;COG3498;TIGR01611 | Phage tail tube protein FII |
| NC\_017033.1\|provirus\_1050333\_1094910\_991 | 324 | GENOMAD.073828.VV | PF10109 | Phage tail assembly chaperone proteins, E, or 41 or 14 |
| NC\_017033.1\|provirus\_1050333\_1094910\_992 | 2319 | GENOMAD.141796.VV | COG3941 | Phage tail tape-measure protein, controls tail length |
| NC\_017033.1\|provirus\_1050333\_1094910\_993 | 843 | GENOMAD.124222.VP | K06906;COG3499 | Phage protein U |
| NC\_017033.1\|provirus\_1050333\_1094910\_994 | 207 | GENOMAD.159179.VP | COG5004;K06370 | P2-like prophage tail protein X |
| NC\_017033.1\|provirus\_1050333\_1094910\_995 | 1053 | GENOMAD.158351.VP | PF05954;K06905;COG3500;TIGR03361 | Phage protein D |

###### 2.14.2.3 Phage ID: NC\_017033.1|provirus\_2089294\_2107522

**Phage-Host Genome Ideogram:**

**Genes Annotation (geNomad):**

| gene | length | marker | annotation\_accessions | annotation\_description |
| --- | --- | --- | --- | --- |
| NC\_017033.1\|provirus\_2089294\_2107522\_1926 | 987 | GENOMAD.147153.VV | PF07120;COG3756 | Uncharacterized conserved protein YdaU, DUF1376 family |
| NC\_017033.1\|provirus\_2089294\_2107522\_1927 | 447 | GENOMAD.220574.VP | PF06892 | Phage regulatory protein CII (CP76) |
| NC\_017033.1\|provirus\_2089294\_2107522\_1928 | 504 | GENOMAD.226021.VP | PF14205 | NA |
| NC\_017033.1\|provirus\_2089294\_2107522\_1929 | 750 | NA | NA | NA |
| NC\_017033.1\|provirus\_2089294\_2107522\_1930 | 900 | NA | NA | NA |
| NC\_017033.1\|provirus\_2089294\_2107522\_1931 | 465 | NA | NA | NA |
| NC\_017033.1\|provirus\_2089294\_2107522\_1932 | 264 | NA | NA | NA |
| NC\_017033.1\|provirus\_2089294\_2107522\_1933 | 888 | NA | NA | NA |
| NC\_017033.1\|provirus\_2089294\_2107522\_1934 | 246 | NA | NA | NA |
| NC\_017033.1\|provirus\_2089294\_2107522\_1935 | 132 | NA | NA | NA |
| NC\_017033.1\|provirus\_2089294\_2107522\_1936 | 321 | NA | NA | NA |
| NC\_017033.1\|provirus\_2089294\_2107522\_1937 | 339 | NA | NA | NA |
| NC\_017033.1\|provirus\_2089294\_2107522\_1938 | 282 | NA | NA | NA |
| NC\_017033.1\|provirus\_2089294\_2107522\_1939 | 822 | GENOMAD.192631.VV | PF03837;TIGR01913 | RecT family |
| NC\_017033.1\|provirus\_2089294\_2107522\_1940 | 1752 | GENOMAD.151126.VP | NA | NA |
| NC\_017033.1\|provirus\_2089294\_2107522\_1941 | 228 | NA | NA | NA |
| NC\_017033.1\|provirus\_2089294\_2107522\_1942 | 333 | NA | NA | NA |
| NC\_017033.1\|provirus\_2089294\_2107522\_1943 | 222 | GENOMAD.224342.VP | PF09035;COG3311 | Predicted DNA-binding transcriptional regulator AlpA |
| NC\_017033.1\|provirus\_2089294\_2107522\_1944 | 927 | NA | NA | NA |
| NC\_017033.1\|provirus\_2089294\_2107522\_1945 | 1797 | NA | NA | NA |
| NC\_017033.1\|provirus\_2089294\_2107522\_1946 | 933 | GENOMAD.166800.VP | COG4643 | Uncharacterized domain associated with phage/plasmid primase |
| NC\_017033.1\|provirus\_2089294\_2107522\_1947 | 369 | NA | NA | NA |
| NC\_017033.1\|provirus\_2089294\_2107522\_1948 | 348 | NA | NA | NA |
| NC\_017033.1\|provirus\_2089294\_2107522\_1949 | 456 | NA | NA | NA |
| NC\_017033.1\|provirus\_2089294\_2107522\_1950 | 972 | NA | NA | NA |

##### 2.14.3 vOTUs

When more than 1 phage is detected in the host genome, we perform a
clustering step using the tool dRep.

A threshold of 0.95 was applied to the ANI similarity index to define
clusters of virus operational taxonomic units (vOTUs).

**Final cluster designations**

| genomad\_summary$seq\_name | genome | secondary\_cluster | threshold | cluster\_method | comparison\_algorithm | primary\_cluster |
| --- | --- | --- | --- | --- | --- | --- |
| NC\_017033.1\|provirus\_1557694\_1600481 | sequence\_000001.fasta.fasta | 1\_0 | 0.05 | average | ANImf | 1 |
| NC\_017033.1\|provirus\_1050333\_1094910 | sequence\_000002.fasta.fasta | 2\_0 | 0.05 | average | ANImf | 2 |
| NC\_017033.1\|provirus\_2089294\_2107522 | sequence\_000000.fasta.fasta | 3\_0 | 0.05 | average | ANImf | 3 |

**Primary clustering plot**

##### 2.14.4 Prophage-Host Prediction

Prediction of potential hosts for the prophage contigs.

| virus | host\_genome | host\_taxonomy | main\_method | confidence\_score | additional\_methods |
| --- | --- | --- | --- | --- | --- |
| NC\_017033.1\|provirus\_1050333\_1094910 | RS\_GCF\_000242255.2 | d\_\_Bacteria;p\_\_Pseudomonadota;c\_\_Gammaproteobacteria;o\_\_Xanthomonadales;f\_\_Rhodanobacteraceae;g\_\_Frateuria;s\_\_Frateuria aurantia | blast | 98.2 | None |
| NC\_017033.1\|provirus\_1557694\_1600481 | RS\_GCF\_000242255.2 | d\_\_Bacteria;p\_\_Pseudomonadota;c\_\_Gammaproteobacteria;o\_\_Xanthomonadales;f\_\_Rhodanobacteraceae;g\_\_Frateuria;s\_\_Frateuria aurantia | blast | 98.4 | iPHoP-RF;64.30 |
| NC\_017033.1\|provirus\_2089294\_2107522 | RS\_GCF\_000242255.2 | d\_\_Bacteria;p\_\_Pseudomonadota;c\_\_Gammaproteobacteria;o\_\_Xanthomonadales;f\_\_Rhodanobacteraceae;g\_\_Frateuria;s\_\_Frateuria aurantia | blast | 95.2 | None |

##### 2.14.5 AMG Predictions

Prediction of auxiliary metabolic genes in the prophage contigs.

| protein | scaffold | amg\_ko | amg\_ko\_name | pfam | pfam\_name |
| --- | --- | --- | --- | --- | --- |
| NC\_017033.1\|provirus\_1557694\_1600481\_7 | NC\_017033.1\|provirus\_1557694\_1600481 | K00558 | DNMT1, dcm; DNA (cytosine-5)-methyltransferase 1 [EC:2.1.1.37] | PF00145.17 | C-5 cytosine-specific DNA methylase |

## 2.15 NC\_017095

##### 2.15.1 Host Genome

**GTDB-Tk taxonomy**:

classification

d\_\_Bacteria;p\_\_Thermotogota;c\_\_Thermotogae;o\_\_Thermotogales;f\_\_Fervidobacteriaceae;g\_\_Fervidobacterium;s\_\_Fervidobacterium
pennivorans

**CheckM2 Summary**:

| total\_contigs | contig\_n50 | completeness | contamination |
| --- | --- | --- | --- |
| 1 | 2166381 | 99.95 | 2.67 |

**Defense-Finder Systems**:

| sys\_id | type | subtype | sys\_beg | sys\_end | protein\_in\_syst | genes\_count | name\_of\_profiles\_in\_sys |
| --- | --- | --- | --- | --- | --- | --- | --- |
| NC\_017095.1\_RM\_Type\_I\_4 | RM | RM\_Type\_I | NC\_017095.1\_540 | NC\_017095.1\_543 | NC\_017095.1\_540,NC\_017095.1\_541,NC\_017095.1\_542,NC\_017095.1\_543 | 4 | RM\_\_Type\_I\_MTases\_FAM\_0,RM\_\_Type\_I\_MTases\_FAM\_0,RM\_\_Type\_I\_REases\_FAM\_0.einsi\_trimmed,RM\_\_Type\_I\_S\_51 |
| NC\_017095.1\_CAS\_Class1-Subtype-III-B\_7 | Cas | CAS\_Class1-Subtype-III-B | NC\_017095.1\_1486 | NC\_017095.1\_1502 | NC\_017095.1\_1486,NC\_017095.1\_1487,NC\_017095.1\_1488,NC\_017095.1\_1489,NC\_017095.1\_1490,NC\_017095.1\_1491,NC\_017095.1\_1495,NC\_017095.1\_1497,NC\_017095.1\_1499,NC\_017095.1\_1500,NC\_017095.1\_1502 | 11 | HTH\_III\_1,cas10\_III\_6,cas2\_I\_II\_III\_IV\_V\_VI\_3,cas2\_I\_II\_III\_IV\_V\_VI\_3,cas6\_I\_II\_III\_IV\_V\_VI\_14,cmr1gr7\_III-B\_1,cmr3gr5\_III-B\_III-C\_6,cmr4gr7\_III-B\_III-C\_1,cmr5gr11\_III-B\_4,cmr6gr7\_III-B\_3,csx1\_III\_9 |
| NC\_017095.1\_CAS\_Class1-Subtype-III-A\_6 | Cas | CAS\_Class1-Subtype-III-A | NC\_017095.1\_1495 | NC\_017095.1\_1512 | NC\_017095.1\_1495,NC\_017095.1\_1497,NC\_017095.1\_1499,NC\_017095.1\_1500,NC\_017095.1\_1502,NC\_017095.1\_1503,NC\_017095.1\_1504,NC\_017095.1\_1505,NC\_017095.1\_1506,NC\_017095.1\_1507,NC\_017095.1\_1508,NC\_017095.1\_1509,NC\_017095.1\_1512 | 13 | HTH\_III\_1,cas10\_III-A\_1,cas1\_I\_II\_III\_IV\_V\_VI\_7,cas2\_I\_II\_III\_IV\_V\_VI\_3,cas2\_I\_II\_III\_IV\_V\_VI\_3,cas6\_I\_II\_III\_IV\_V\_VI\_14,cas6\_I\_II\_III\_IV\_V\_VI\_19,casR\_III\_1,csm2gr11\_III-A\_15,csm3gr7\_III-A\_1,csm4gr5\_III-A\_1,csm5gr7\_III-A\_2,csx1\_III\_9 |
| NC\_017095.1\_CAS\_Class1-Subtype-III-A\_6 | Cas | CAS\_Class1-Subtype-III-A | NC\_017095.1\_1495 | NC\_017095.1\_1512 | NC\_017095.1\_1495,NC\_017095.1\_1497,NC\_017095.1\_1499,NC\_017095.1\_1500,NC\_017095.1\_1502,NC\_017095.1\_1503,NC\_017095.1\_1504,NC\_017095.1\_1505,NC\_017095.1\_1506,NC\_017095.1\_1507,NC\_017095.1\_1508,NC\_017095.1\_1509,NC\_017095.1\_1512 | 13 | HTH\_III\_1,cas10\_III-A\_1,cas1\_I\_II\_III\_IV\_V\_VI\_7,cas2\_I\_II\_III\_IV\_V\_VI\_3,cas2\_I\_II\_III\_IV\_V\_VI\_3,cas6\_I\_II\_III\_IV\_V\_VI\_14,cas6\_I\_II\_III\_IV\_V\_VI\_19,casR\_III\_1,csm2gr11\_III-A\_15,csm3gr7\_III-A\_1,csm4gr5\_III-A\_1,csm5gr7\_III-A\_2,csx1\_III\_9 |
| NC\_017095.1\_Esos\_1 | Esos | Esos | NC\_017095.1\_1541 | NC\_017095.1\_1541 | NC\_017095.1\_1541 | 1 | Esos\_\_VCA0450 |
| NC\_017095.1\_VP1840\_2 | VP1840 | VP1840 | NC\_017095.1\_1651 | NC\_017095.1\_1651 | NC\_017095.1\_1651 | 1 | VP1840\_\_VP1840 |
| NC\_017095.1\_RM\_Type\_II\_5 | RM | RM\_Type\_II | NC\_017095.1\_1662 | NC\_017095.1\_1664 | NC\_017095.1\_1662,NC\_017095.1\_1663,NC\_017095.1\_1664 | 3 | RM\_Type\_II\_\_Type\_II\_MTases\_FAM\_38,RM\_Type\_II\_\_Type\_II\_MTases\_FAM\_4,RM\_Type\_II\_\_Type\_II\_REase32 |

## 2.16 NC\_018014

##### 2.16.1 Host Genome

**GTDB-Tk taxonomy**:

classification

d\_\_Bacteria;p\_\_Acidobacteriota;c\_\_Terriglobia;o\_\_Terriglobales;f\_\_Acidobacteriaceae;g\_\_Terriglobus;s\_\_Terriglobus
roseus

**CheckM2 Summary**:

| total\_contigs | contig\_n50 | completeness | contamination |
| --- | --- | --- | --- |
| 1 | 5227858 | 99.99 | 9.38 |

**Defense-Finder Systems**:

| sys\_id | type | subtype | sys\_beg | sys\_end | protein\_in\_syst | genes\_count | name\_of\_profiles\_in\_sys |
| --- | --- | --- | --- | --- | --- | --- | --- |
| NC\_018014.1\_DS-6\_5 | DS-6 | DS-6 | NC\_018014.1\_132 | NC\_018014.1\_133 | NC\_018014.1\_132,NC\_018014.1\_133 | 2 | DS-6\_\_DS-6A,DS-6\_\_DS-6B |
| NC\_018014.1\_Pycsar\_6 | Pycsar | Pycsar | NC\_018014.1\_571 | NC\_018014.1\_573 | NC\_018014.1\_571,NC\_018014.1\_572,NC\_018014.1\_573 | 3 | CBASS\_\_2TM\_5,Pycsar\_\_AG\_cyclase,Pycsar\_\_AG\_cyclase |
| NC\_018014.1\_DISARM\_1\_4 | DISARM | DISARM\_1 | NC\_018014.1\_3050 | NC\_018014.1\_3053 | NC\_018014.1\_3050,NC\_018014.1\_3051,NC\_018014.1\_3052,NC\_018014.1\_3053 | 4 | DISARM\_1\_\_drmMI,DISARM\_\_drmA,DISARM\_\_drmB,DISARM\_\_drmC |
| NC\_018014.1\_CBASS\_III\_3 | CBASS | CBASS\_III | NC\_018014.1\_3068 | NC\_018014.1\_3071 | NC\_018014.1\_3068,NC\_018014.1\_3069,NC\_018014.1\_3070,NC\_018014.1\_3071 | 4 | CBASS\_\_Cyclase\_II,CBASS\_\_Endonuc\_big,CBASS\_\_TRIP13,CBASS\_\_bacHORMA\_1 |

## 2.17 NC\_018068

##### 2.17.1 Host Genome

**GTDB-Tk taxonomy**:

classification

d\_\_Bacteria;p\_\_Bacillota\_B;c\_\_Desulfitobacteriia;o\_\_Desulfitobacteriales;f\_\_Desulfitobacteriaceae;g\_\_Desulfosporosinus;s\_\_Desulfosporosinus
acidiphilus

**CheckM2 Summary**:

| total\_contigs | contig\_n50 | completeness | contamination |
| --- | --- | --- | --- |
| 1 | 4926837 | 99.99 | 0.42 |

**Defense-Finder Systems**:

| sys\_id | type | subtype | sys\_beg | sys\_end | protein\_in\_syst | genes\_count | name\_of\_profiles\_in\_sys |
| --- | --- | --- | --- | --- | --- | --- | --- |
| NC\_018068.1\_Hachiman\_2 | Hachiman | Hachiman | NC\_018068.1\_1171 | NC\_018068.1\_1172 | NC\_018068.1\_1171,NC\_018068.1\_1172 | 2 | Hachiman\_\_HamA\_1,Hachiman\_\_HamB |
| NC\_018068.1\_RM\_Type\_I\_10 | RM | RM\_Type\_I | NC\_018068.1\_1178 | NC\_018068.1\_1180 | NC\_018068.1\_1178,NC\_018068.1\_1179,NC\_018068.1\_1180 | 3 | RM\_\_Type\_I\_MTases\_FAM\_1,RM\_\_Type\_I\_REases\_FAM\_2.einsi\_trimmed,RM\_\_Type\_I\_S\_03 |
| NC\_018068.1\_RM\_Type\_IV\_14 | RM | RM\_Type\_IV | NC\_018068.1\_1186 | NC\_018068.1\_1186 | NC\_018068.1\_1186 | 1 | RM\_Type\_IV\_\_Type\_IV\_03 |
| NC\_018068.1\_RM\_Type\_II\_11 | RM | RM\_Type\_II | NC\_018068.1\_1192 | NC\_018068.1\_1193 | NC\_018068.1\_1192,NC\_018068.1\_1193 | 2 | RM\_Type\_II\_\_Type\_II\_MTases\_FAM\_7,RM\_Type\_II\_\_Type\_II\_REase10 |
| NC\_018068.1\_CAS\_Class1-Subtype-I-C\_15 | Cas | CAS\_Class1-Subtype-I-C | NC\_018068.1\_1411 | NC\_018068.1\_1418 | NC\_018068.1\_1411,NC\_018068.1\_1412,NC\_018068.1\_1413,NC\_018068.1\_1414,NC\_018068.1\_1415,NC\_018068.1\_1417,NC\_018068.1\_1418 | 7 | cas1\_I\_II\_III\_IV\_V\_VI\_6,cas2\_I\_II\_III\_IV\_V\_VI\_3,cas3\_I\_5,cas4\_I\_II\_III\_IV\_V\_VI\_6,cas5\_I-C\_11,cas7\_I-C\_13,cas8c\_I-C\_4 |
| NC\_018068.1\_RM\_Type\_III\_13 | RM | RM\_Type\_III | NC\_018068.1\_1541 | NC\_018068.1\_1542 | NC\_018068.1\_1541,NC\_018068.1\_1542 | 2 | RM\_Type\_III\_\_Type\_III\_MTases\_FAM\_0,RM\_Type\_III\_\_Type\_III\_REases\_FAM\_0.einsi\_trimmed |
| NC\_018068.1\_Ceres\_1 | Ceres | Ceres | NC\_018068.1\_3152 | NC\_018068.1\_3152 | NC\_018068.1\_3152 | 1 | Ceres\_\_CrsA1 |
| NC\_018068.1\_MazEF\_6 | MazEF | MazEF | NC\_018068.1\_3456 | NC\_018068.1\_3457 | NC\_018068.1\_3456,NC\_018068.1\_3457 | 2 | MazEF\_\_MazE,MazEF\_\_MazF |
| NC\_018068.1\_Rst\_HelicaseDUF2290\_7 | Rst\_HelicaseDUF2290 | Rst\_HelicaseDUF2290 | NC\_018068.1\_3757 | NC\_018068.1\_3758 | NC\_018068.1\_3757,NC\_018068.1\_3758 | 2 | Rst\_HelicaseDUF2290\_\_DUF2290,Rst\_HelicaseDUF2290\_\_Helicase |
| NC\_018068.1\_Hachiman\_3 | Hachiman | Hachiman | NC\_018068.1\_3759 | NC\_018068.1\_3760 | NC\_018068.1\_3759,NC\_018068.1\_3760 | 2 | Hachiman\_\_HamA\_1,Hachiman\_\_HamB |
| NC\_018068.1\_Septu\_8 | Septu | Septu | NC\_018068.1\_4412 | NC\_018068.1\_4413 | NC\_018068.1\_4412,NC\_018068.1\_4413 | 2 | Septu\_\_PtuA,Septu\_\_PtuB |
| NC\_018068.1\_RM\_Type\_II\_12 | RM | RM\_Type\_II | NC\_018068.1\_4429 | NC\_018068.1\_4430 | NC\_018068.1\_4429,NC\_018068.1\_4430 | 2 | RM\_Type\_II\_\_Type\_II\_MTases\_FAM\_25,RM\_Type\_II\_\_Type\_II\_REase09 |

**Genomad and CheckV Summary**:

| seq\_name | length | gene\_count | viral\_genes | checkv\_quality | miuvig\_quality | taxonomy | topology | coordinates |
| --- | --- | --- | --- | --- | --- | --- | --- | --- |
| NC\_018068.1\|provirus\_1361108\_1399802 | 38695 | 52 | 24 | Medium-quality | Genome-fragment | Viruses;Duplodnaviria;Heunggongvirae;Uroviricota;Caudoviricetes;; | Provirus | 1361108-1399802 |
| NC\_018068.1\|provirus\_2072558\_2098541 | 25984 | 36 | 4 | High-quality | High-quality | Viruses;Duplodnaviria;Heunggongvirae;Uroviricota;Caudoviricetes;; | Provirus | 2072558-2098541 |

**Host Genome Ideogram with Phages**:

In
this circular plot, **“c”** indicates the contig, and the
number that follows (e.g., **c1**) represents the contig
number. If multiple contigs are present in the genome, each will be
shown with a distinct label (e.g., **c1**,
**c2**, etc.).

##### 2.17.2 Prophages

**Select prophage to show:**

###### 2.17.2.1 Phage ID: NC\_018068.1|provirus\_1361108\_1399802

**Phage-Host Genome Ideogram:**

**Genes Annotation (geNomad):**

| gene | length | marker | annotation\_accessions | annotation\_description |
| --- | --- | --- | --- | --- |
| NC\_018068.1\|provirus\_1361108\_1399802\_1216 | 417 | NA | NA | NA |
| NC\_018068.1\|provirus\_1361108\_1399802\_1217 | 285 | NA | NA | NA |
| NC\_018068.1\|provirus\_1361108\_1399802\_1218 | 213 | NA | NA | NA |
| NC\_018068.1\|provirus\_1361108\_1399802\_1219 | 825 | GENOMAD.133073.VP | PF03374;COG3645 | Phage antirepressor protein YoqD, KilAC domain |
| NC\_018068.1\|provirus\_1361108\_1399802\_1220 | 1245 | GENOMAD.014802.VV | PF00176;K20093;COG1061;TIGR04095 | Superfamily II DNA or RNA helicase |
| NC\_018068.1\|provirus\_1361108\_1399802\_1221 | 285 | GENOMAD.062524.VV | PF03838;COG3331;TIGR00648;K03552 | Penicillin-binding protein-related factor A, putative recombinase |
| NC\_018068.1\|provirus\_1361108\_1399802\_1222 | 1668 | GENOMAD.016341.VV | PF13479;PF12684;TIGR01618;K07465;COG1468 | phage nucleotide-binding protein |
| NC\_018068.1\|provirus\_1361108\_1399802\_1223 | 450 | GENOMAD.031678.VV | PF05037 | NA |
| NC\_018068.1\|provirus\_1361108\_1399802\_1224 | 1695 | GENOMAD.102034.VP | TIGR01636 | NA |
| NC\_018068.1\|provirus\_1361108\_1399802\_1225 | 1875 | GENOMAD.024099.VV | NA | NA |
| NC\_018068.1\|provirus\_1361108\_1399802\_1226 | 381 | GENOMAD.192072.VP | NA | NA |
| NC\_018068.1\|provirus\_1361108\_1399802\_1227 | 453 | GENOMAD.076519.VV | PF05263;TIGR01636;COG2739;K01994 | phage transcriptional activator, RinA family |
| NC\_018068.1\|provirus\_1361108\_1399802\_1228 | 189 | NA | NA | NA |
| NC\_018068.1\|provirus\_1361108\_1399802\_1229 | 207 | NA | NA | NA |
| NC\_018068.1\|provirus\_1361108\_1399802\_1230 | 357 | GENOMAD.221055.VP | TIGR02646 | NA |
| NC\_018068.1\|provirus\_1361108\_1399802\_1231 | 528 | GENOMAD.098194.VV | PF05119;COG3747;TIGR01558 | Phage terminase, small subunit |
| NC\_018068.1\|provirus\_1361108\_1399802\_1232 | 1254 | GENOMAD.038338.VV | COG3392 | NA |
| NC\_018068.1\|provirus\_1361108\_1399802\_1233 | 525 | GENOMAD.133508.VP | PF13392 | HNH endonuclease |
| NC\_018068.1\|provirus\_1361108\_1399802\_1234 | 186 | GENOMAD.209016.VC | NA | NA |
| NC\_018068.1\|provirus\_1361108\_1399802\_1235 | 816 | GENOMAD.105515.VV | NA | NA |
| NC\_018068.1\|provirus\_1361108\_1399802\_1236 | 240 | GENOMAD.225559.VP | NA | NA |
| NC\_018068.1\|provirus\_1361108\_1399802\_1237 | 252 | NA | NA | NA |
| NC\_018068.1\|provirus\_1361108\_1399802\_1238 | 210 | NA | NA | NA |
| NC\_018068.1\|provirus\_1361108\_1399802\_1239 | 1548 | GENOMAD.190509.VP | PF03354;COG4626 | Phage terminase-like protein, large subunit, contains N-terminal HTH domain |
| NC\_018068.1\|provirus\_1361108\_1399802\_1240 | 1164 | GENOMAD.179073.VP | NA | NA |
| NC\_018068.1\|provirus\_1361108\_1399802\_1241 | 1068 | GENOMAD.158277.VP | PF00574;PF19602;K01358;TIGR00493;COG3904 | ATP-dependent Clp endopeptidase, proteolytic subunit ClpP |
| NC\_018068.1\|provirus\_1361108\_1399802\_1242 | 1275 | GENOMAD.092606.VV | PF05065;COG4653;TIGR01554 | Predicted phage phi-C31 gp36 major capsid-like protein |
| NC\_018068.1\|provirus\_1361108\_1399802\_1243 | 477 | GENOMAD.077304.VV | NA | NA |
| NC\_018068.1\|provirus\_1361108\_1399802\_1244 | 594 | GENOMAD.067701.VV | PF05135;TIGR02215 | phage conserved hypothetical protein, phiE125 gp8 family |
| NC\_018068.1\|provirus\_1361108\_1399802\_1245 | 330 | GENOMAD.027939.VV | PF05521;COG5614;TIGR01563 | Bacteriophage head-tail adaptor |
| NC\_018068.1\|provirus\_1361108\_1399802\_1246 | 414 | GENOMAD.018908.VV | PF04883;TIGR01725;COG5005 | Bacteriophage HK97-gp10, putative tail-component |
| NC\_018068.1\|provirus\_1361108\_1399802\_1247 | 435 | GENOMAD.029331.VV | PF20765 | Phage tail terminator protein |
| NC\_018068.1\|provirus\_1361108\_1399802\_1248 | 1068 | GENOMAD.014203.VV | PF04984;PF17482;COG4386 | Mu-like prophage tail sheath protein gpL |
| NC\_018068.1\|provirus\_1361108\_1399802\_1249 | 429 | GENOMAD.008035.VV | PF09393 | Phage tail tube protein |
| NC\_018068.1\|provirus\_1361108\_1399802\_1250 | 402 | GENOMAD.020710.VV | PF08890 | Phage XkdN-like tail assembly chaperone protein, TAC |
| NC\_018068.1\|provirus\_1361108\_1399802\_1251 | 2172 | GENOMAD.117212.VP | NA | NA |
| NC\_018068.1\|provirus\_1361108\_1399802\_1252 | 405 | GENOMAD.025221.VV | PF06995;COG1652 | Nucleoid-associated protein YgaU, contains BON and LysM domains |
| NC\_018068.1\|provirus\_1361108\_1399802\_1253 | 993 | GENOMAD.020668.VV | PF05954;COG4379;K06905;TIGR03361 | Mu-like prophage tail protein gpP |
| NC\_018068.1\|provirus\_1361108\_1399802\_1254 | 369 | GENOMAD.046601.VV | PF10844 | NA |
| NC\_018068.1\|provirus\_1361108\_1399802\_1255 | 432 | GENOMAD.015505.VV | PF10934;COG4381 | Mu-like prophage protein gp46 |
| NC\_018068.1\|provirus\_1361108\_1399802\_1256 | 1074 | GENOMAD.110264.VV | PF04865;COG3948 | Phage-related baseplate assembly protein |
| NC\_018068.1\|provirus\_1361108\_1399802\_1257 | 522 | GENOMAD.018612.VV | PF10076;COG3778;TIGR02242 | Uncharacterized protein YmfQ in lambdoid prophage, DUF2313 family |
| NC\_018068.1\|provirus\_1361108\_1399802\_1258 | 567 | GENOMAD.069954.VV | NA | NA |
| NC\_018068.1\|provirus\_1361108\_1399802\_1259 | 336 | NA | NA | NA |
| NC\_018068.1\|provirus\_1361108\_1399802\_1260 | 1644 | GENOMAD.014429.VV | NA | NA |
| NC\_018068.1\|provirus\_1361108\_1399802\_1261 | 339 | GENOMAD.213587.VP | NA | NA |
| NC\_018068.1\|provirus\_1361108\_1399802\_1262 | 138 | GENOMAD.041649.VV | PF09693;TIGR01669 | phage uncharacterized protein, XkdX family |
| NC\_018068.1\|provirus\_1361108\_1399802\_1263 | 240 | NA | NA | NA |
| NC\_018068.1\|provirus\_1361108\_1399802\_1264 | 321 | GENOMAD.130222.VV | COG3105 | NA |
| NC\_018068.1\|provirus\_1361108\_1399802\_1265 | 486 | GENOMAD.151608.VV | PF19988 | NA |
| NC\_018068.1\|provirus\_1361108\_1399802\_1266 | 774 | NA | NA | NA |
| NC\_018068.1\|provirus\_1361108\_1399802\_1267 | 414 | NA | NA | NA |

###### 2.17.2.2 Phage ID: NC\_018068.1|provirus\_2072558\_2098541

**Phage-Host Genome Ideogram:**

**Genes Annotation (geNomad):**

| gene | length | marker | annotation\_accessions | annotation\_description |
| --- | --- | --- | --- | --- |
| NC\_018068.1\|provirus\_2072558\_2098541\_1928 | 177 | GENOMAD.171344.VV | PF07278;K22014;COG4220;TIGR01764 | Phage DNA packaging protein, Nu1 subunit of terminase |
| NC\_018068.1\|provirus\_2072558\_2098541\_1929 | 189 | NA | NA | NA |
| NC\_018068.1\|provirus\_2072558\_2098541\_1930 | 204 | NA | NA | NA |
| NC\_018068.1\|provirus\_2072558\_2098541\_1931 | 147 | NA | NA | NA |
| NC\_018068.1\|provirus\_2072558\_2098541\_1932 | 867 | NA | NA | NA |
| NC\_018068.1\|provirus\_2072558\_2098541\_1933 | 2490 | NA | NA | NA |
| NC\_018068.1\|provirus\_2072558\_2098541\_1934 | 264 | NA | NA | NA |
| NC\_018068.1\|provirus\_2072558\_2098541\_1935 | 213 | NA | NA | NA |
| NC\_018068.1\|provirus\_2072558\_2098541\_1936 | 189 | NA | NA | NA |
| NC\_018068.1\|provirus\_2072558\_2098541\_1937 | 309 | GENOMAD.037442.VV | NA | NA |
| NC\_018068.1\|provirus\_2072558\_2098541\_1938 | 135 | NA | NA | NA |
| NC\_018068.1\|provirus\_2072558\_2098541\_1939 | 144 | NA | NA | NA |
| NC\_018068.1\|provirus\_2072558\_2098541\_1940 | 204 | GENOMAD.220824.VP | PF18903 | NA |
| NC\_018068.1\|provirus\_2072558\_2098541\_1941 | 129 | NA | NA | NA |
| NC\_018068.1\|provirus\_2072558\_2098541\_1942 | 207 | GENOMAD.207723.VP | PF10960 | BhlA holin family |
| NC\_018068.1\|provirus\_2072558\_2098541\_1943 | 417 | GENOMAD.126365.VV | NA | NA |
| NC\_018068.1\|provirus\_2072558\_2098541\_1944 | 888 | GENOMAD.136497.VV | PF17236 | Phage capsid-like protein |
| NC\_018068.1\|provirus\_2072558\_2098541\_1945 | 192 | NA | NA | NA |
| NC\_018068.1\|provirus\_2072558\_2098541\_1946 | 813 | GENOMAD.225339.VP | NA | NA |
| NC\_018068.1\|provirus\_2072558\_2098541\_1947 | 1497 | GENOMAD.225126.VP | PF06862;TIGR01587;COG4098;K17677 | CRISPR-associated helicase Cas3 |
| NC\_018068.1\|provirus\_2072558\_2098541\_1948 | 393 | GENOMAD.142024.VV | NA | NA |
| NC\_018068.1\|provirus\_2072558\_2098541\_1949 | 1572 | GENOMAD.180947.VP | PF04466;COG5323;TIGR01547;K21523 | Large terminase phage packaging protein |
| NC\_018068.1\|provirus\_2072558\_2098541\_1950 | 1461 | GENOMAD.003260.VV | PF05133;TIGR01538;COG3567 | phage portal protein, SPP1 family |
| NC\_018068.1\|provirus\_2072558\_2098541\_1951 | 294 | NA | NA | NA |
| NC\_018068.1\|provirus\_2072558\_2098541\_1952 | 351 | GENOMAD.223401.VP | PF09851 | NA |
| NC\_018068.1\|provirus\_2072558\_2098541\_1953 | 1242 | NA | NA | NA |
| NC\_018068.1\|provirus\_2072558\_2098541\_1954 | 867 | NA | NA | NA |
| NC\_018068.1\|provirus\_2072558\_2098541\_1955 | 219 | NA | NA | NA |
| NC\_018068.1\|provirus\_2072558\_2098541\_1956 | 261 | NA | NA | NA |
| NC\_018068.1\|provirus\_2072558\_2098541\_1957 | 348 | GENOMAD.220167.VP | PF20449 | NA |
| NC\_018068.1\|provirus\_2072558\_2098541\_1958 | 1119 | NA | NA | NA |
| NC\_018068.1\|provirus\_2072558\_2098541\_1959 | 213 | NA | NA | NA |
| NC\_018068.1\|provirus\_2072558\_2098541\_1960 | 987 | NA | NA | NA |

##### 2.17.3 vOTUs

When more than 1 phage is detected in the host genome, we perform a
clustering step using the tool dRep.

A threshold of 0.95 was applied to the ANI similarity index to define
clusters of virus operational taxonomic units (vOTUs).

**Final cluster designations**

| genomad\_summary$seq\_name | genome | secondary\_cluster | threshold | cluster\_method | comparison\_algorithm | primary\_cluster |
| --- | --- | --- | --- | --- | --- | --- |
| NC\_018068.1\|provirus\_1361108\_1399802 | sequence\_000000.fasta.fasta | 1\_0 | 0.05 | average | ANImf | 1 |
| NC\_018068.1\|provirus\_2072558\_2098541 | sequence\_000001.fasta.fasta | 2\_0 | 0.05 | average | ANImf | 2 |

**Primary clustering plot**

##### 2.17.4 Prophage-Host Prediction

Prediction of potential hosts for the prophage contigs.

| virus | host\_genome | host\_taxonomy | main\_method | confidence\_score | additional\_methods |
| --- | --- | --- | --- | --- | --- |
| NC\_018068.1\|provirus\_1361108\_1399802 | RS\_GCF\_000255115.2 | d\_\_Bacteria;p\_\_Bacillota\_B;c\_\_Desulfitobacteriia;o\_\_Desulfitobacteriales;f\_\_Desulfitobacteriaceae;g\_\_Desulfosporosinus;s\_\_Desulfosporosinus acidiphilus | blast | 96.4 | iPHoP-RF;53.60 |
| NC\_018068.1\|provirus\_1361108\_1399802 | RS\_GCF\_001707885.1 | d\_\_Bacteria;p\_\_Bacillota\_B;c\_\_Desulfitobacteriia;o\_\_Desulfitobacteriales;f\_\_Desulfitobacteriaceae;g\_\_Desulfosporosinus;s\_\_Desulfosporosinus sp001707885 | iPHoP-RF | 95.4 | None |
| NC\_018068.1\|provirus\_1361108\_1399802 | RS\_GCF\_001936615.1 | d\_\_Bacteria;p\_\_Bacillota\_B;c\_\_Desulfitobacteriia;o\_\_Desulfitobacteriales;f\_\_Desulfitobacteriaceae;g\_\_Desulfosporosinus;s\_\_Desulfosporosinus metallidurans | iPHoP-RF | 95.4 | None |
| NC\_018068.1\|provirus\_1361108\_1399802 | RS\_GCF\_000224515.1 | d\_\_Bacteria;p\_\_Bacillota\_B;c\_\_Desulfitobacteriia;o\_\_Desulfitobacteriales;f\_\_Desulfitobacteriaceae;g\_\_Desulfosporosinus;s\_\_Desulfosporosinus sp000224515 | iPHoP-RF | 95.1 | None |
| NC\_018068.1\|provirus\_1361108\_1399802 | RS\_GCF\_000960765.1 | d\_\_Bacteria;p\_\_Bacillota\_B;c\_\_Desulfitobacteriia;o\_\_Desulfitobacteriales;f\_\_Desulfitobacteriaceae;g\_\_Desulfosporosinus;s\_\_Desulfosporosinus sp000960765 | iPHoP-RF | 93.4 | None |
| NC\_018068.1\|provirus\_1361108\_1399802 | RS\_GCF\_004766055.1 | d\_\_Bacteria;p\_\_Bacillota\_B;c\_\_Desulfitobacteriia;o\_\_Desulfitobacteriales;f\_\_Desulfitobacteriaceae;g\_\_Desulfosporosinus;s\_\_Desulfosporosinus sp004766055 | iPHoP-RF | 93.1 | None |
| NC\_018068.1\|provirus\_1361108\_1399802 | 3300011997\_21 | d\_\_Bacteria;p\_\_Bacillota\_B;c\_\_Desulfitobacteriia;o\_\_Desulfitobacteriales;f\_\_Desulfitobacteriaceae;g**Desulfosporosinus;s** | iPHoP-RF | 92.1 | None |
| NC\_018068.1\|provirus\_1361108\_1399802 | GB\_GCA\_002404215.1 | d\_\_Bacteria;p\_\_Bacillota\_B;c\_\_Desulfitobacteriia;o\_\_Desulfitobacteriales;f\_\_Desulfitobacteriaceae;g\_\_Desulfosporosinus;s\_\_Desulfosporosinus sp002404215 | iPHoP-RF | 91.8 | None |
| NC\_018068.1\|provirus\_1361108\_1399802 | RS\_GCF\_001029285.1 | d\_\_Bacteria;p\_\_Bacillota\_B;c\_\_Desulfitobacteriia;o\_\_Desulfitobacteriales;f\_\_Desulfitobacteriaceae;g\_\_Desulfosporosinus;s\_\_Desulfosporosinus acididurans | blast | 91.8 | None |
| NC\_018068.1\|provirus\_1361108\_1399802 | RS\_GCF\_002196705.1 | d\_\_Bacteria;p\_\_Bacillota\_B;c\_\_Desulfitobacteriia;o\_\_Desulfitobacteriales;f\_\_Desulfitobacteriaceae;g\_\_Desulfosporosinus;s\_\_Desulfosporosinus sp002196705 | blast | 91.2 | iPHoP-RF;78.30 |
| NC\_018068.1\|provirus\_1361108\_1399802 | GB\_GCA\_900290375.1 | d\_\_Bacteria;p\_\_Bacillota\_B;c\_\_Desulfitobacteriia;o\_\_Desulfitobacteriales;f\_\_Desulfitobacteriaceae;g\_\_Desulfosporosinus;s\_\_Desulfosporosinus infrequens | blast | 90.3 | iPHoP-RF;73.50 |
| NC\_018068.1\|provirus\_2072558\_2098541 | RS\_GCF\_000255115.2 | d\_\_Bacteria;p\_\_Bacillota\_B;c\_\_Desulfitobacteriia;o\_\_Desulfitobacteriales;f\_\_Desulfitobacteriaceae;g\_\_Desulfosporosinus;s\_\_Desulfosporosinus acidiphilus | blast | 93.6 | iPHoP-RF;69.50 |

## 2.18 NC\_018515

##### 2.18.1 Host Genome

**GTDB-Tk taxonomy**:

classification

d\_\_Bacteria;p\_\_Bacillota\_B;c\_\_Desulfitobacteriia;o\_\_Desulfitobacteriales;f\_\_Desulfitobacteriaceae;g\_\_Desulfosporosinus;s\_\_Desulfosporosinus
meridiei

**CheckM2 Summary**:

| total\_contigs | contig\_n50 | completeness | contamination |
| --- | --- | --- | --- |
| 1 | 4873567 | 100 | 1.83 |

**Defense-Finder Systems**:

| sys\_id | type | subtype | sys\_beg | sys\_end | protein\_in\_syst | genes\_count | name\_of\_profiles\_in\_sys |
| --- | --- | --- | --- | --- | --- | --- | --- |
| NC\_018515.1\_AbiAlpha\_1 | AbiAlpha | AbiAlpha | NC\_018515.1\_121 | NC\_018515.1\_121 | NC\_018515.1\_121 | 1 | AbiAlpha\_\_AbiAlpha |
| NC\_018515.1\_VP1839\_11 | VP1839 | VP1839 | NC\_018515.1\_122 | NC\_018515.1\_122 | NC\_018515.1\_122 | 1 | VP1839\_\_VP1839 |
| NC\_018515.1\_CBASS\_IV\_7 | CBASS | CBASS\_IV | NC\_018515.1\_324 | NC\_018515.1\_328 | NC\_018515.1\_324,NC\_018515.1\_325,NC\_018515.1\_326,NC\_018515.1\_327,NC\_018515.1\_328 | 5 | CBASS\_\_2TM\_type\_IV,CBASS\_\_Cyclase\_SMODS,CBASS\_\_OGG,CBASS\_\_QueC,CBASS\_\_TGT |
| NC\_018515.1\_BREX\_I\_4 | BREX | BREX\_I | NC\_018515.1\_329 | NC\_018515.1\_335 | NC\_018515.1\_329,NC\_018515.1\_330,NC\_018515.1\_331,NC\_018515.1\_332,NC\_018515.1\_334,NC\_018515.1\_335 | 6 | BREX\_\_brxA\_DUF1819,BREX\_\_brxB\_DUF1788,BREX\_\_brxC,BREX\_\_brxL,BREX\_\_pglX1,BREX\_\_pglZA |
| NC\_018515.1\_Prometheus\_9 | Prometheus | Prometheus | NC\_018515.1\_337 | NC\_018515.1\_337 | NC\_018515.1\_337 | 1 | Prometheus\_\_ProA |
| NC\_018515.1\_Azaca\_2 | Azaca | Azaca | NC\_018515.1\_349 | NC\_018515.1\_351 | NC\_018515.1\_349,NC\_018515.1\_350,NC\_018515.1\_351 | 3 | Azaca\_\_ZacA,Azaca\_\_ZacB,Azaca\_\_ZacC |
| NC\_018515.1\_Wadjet\_III\_13 | Wadjet | Wadjet\_III | NC\_018515.1\_1811 | NC\_018515.1\_1814 | NC\_018515.1\_1811,NC\_018515.1\_1812,NC\_018515.1\_1813,NC\_018515.1\_1814 | 4 | Wadjet\_\_JetA\_III,Wadjet\_\_JetB\_III,Wadjet\_\_JetC\_III,Wadjet\_\_JetD\_III |
| NC\_018515.1\_SpbK\_10 | SpbK | SpbK | NC\_018515.1\_2272 | NC\_018515.1\_2272 | NC\_018515.1\_2272 | 1 | SpbK\_\_SpbK |
| NC\_018515.1\_Kiwa\_8 | Kiwa | Kiwa | NC\_018515.1\_3261 | NC\_018515.1\_3262 | NC\_018515.1\_3261,NC\_018515.1\_3262 | 2 | Kiwa\_\_KwaA,Kiwa\_\_KwaB\_2 |

**Genomad and CheckV Summary**:

| seq\_name | length | gene\_count | viral\_genes | checkv\_quality | miuvig\_quality | taxonomy | topology | coordinates |
| --- | --- | --- | --- | --- | --- | --- | --- | --- |
| NC\_018515.1\|provirus\_3418112\_3436097 | 17986 | 25 | 18 | Low-quality | Genome-fragment | Viruses;Duplodnaviria;Heunggongvirae;Uroviricota;Caudoviricetes;; | Provirus | 3418112-3436097 |
| NC\_018515.1\|provirus\_4659544\_4693604 | 34061 | 46 | 20 | Medium-quality | Genome-fragment | Viruses;Duplodnaviria;Heunggongvirae;Uroviricota;Caudoviricetes;; | Provirus | 4659544-4693604 |

**Host Genome Ideogram with Phages**:

In
this circular plot, **“c”** indicates the contig, and the
number that follows (e.g., **c1**) represents the contig
number. If multiple contigs are present in the genome, each will be
shown with a distinct label (e.g., **c1**,
**c2**, etc.).

##### 2.18.2 Prophages

**Select prophage to show:**

###### 2.18.2.1 Phage ID: NC\_018515.1|provirus\_3418112\_3436097

**Phage-Host Genome Ideogram:**

**Genes Annotation (geNomad):**

| gene | length | marker | annotation\_accessions | annotation\_description |
| --- | --- | --- | --- | --- |
| NC\_018515.1\|provirus\_3418112\_3436097\_3127 | 576 | NA | NA | NA |
| NC\_018515.1\|provirus\_3418112\_3436097\_3128 | 345 | GENOMAD.178485.VV | NA | NA |
| NC\_018515.1\|provirus\_3418112\_3436097\_3129 | 756 | NA | NA | NA |
| NC\_018515.1\|provirus\_3418112\_3436097\_3130 | 249 | GENOMAD.042597.VV | PF10779 | Haemolysin XhlA |
| NC\_018515.1\|provirus\_3418112\_3436097\_3131 | 171 | GENOMAD.019319.VV | PF09693;TIGR01669 | phage uncharacterized protein, XkdX family |
| NC\_018515.1\|provirus\_3418112\_3436097\_3132 | 300 | NA | NA | NA |
| NC\_018515.1\|provirus\_3418112\_3436097\_3133 | 1080 | GENOMAD.010477.VV | NA | NA |
| NC\_018515.1\|provirus\_3418112\_3436097\_3134 | 612 | GENOMAD.018612.VV | PF10076;COG3778;TIGR02242 | Uncharacterized protein YmfQ in lambdoid prophage, DUF2313 family |
| NC\_018515.1\|provirus\_3418112\_3436097\_3135 | 717 | GENOMAD.072355.VV | PF18454 | Major tropism determinant N-terminal domain |
| NC\_018515.1\|provirus\_3418112\_3436097\_3136 | 1023 | GENOMAD.037496.VV | PF18454 | Major tropism determinant N-terminal domain |
| NC\_018515.1\|provirus\_3418112\_3436097\_3137 | 414 | GENOMAD.105081.VV | NA | NA |
| NC\_018515.1\|provirus\_3418112\_3436097\_3138 | 1038 | GENOMAD.004833.VV | PF04865;COG3299 | Baseplate J-like protein |
| NC\_018515.1\|provirus\_3418112\_3436097\_3139 | 402 | GENOMAD.016318.VV | PF10934;COG4381;TIGR03357 | Mu-like prophage protein gp46 |
| NC\_018515.1\|provirus\_3418112\_3436097\_3140 | 357 | GENOMAD.020599.VV | PF10844 | NA |
| NC\_018515.1\|provirus\_3418112\_3436097\_3141 | 987 | GENOMAD.018966.VV | PF14594;COG4379;TIGR03361;K06905 | Mu-like prophage tail protein gpP |
| NC\_018515.1\|provirus\_3418112\_3436097\_3142 | 663 | GENOMAD.015578.VV | PF06995;COG1652 | Nucleoid-associated protein YgaU, contains BON and LysM domains |
| NC\_018515.1\|provirus\_3418112\_3436097\_3143 | 1689 | GENOMAD.032671.VV | NA | NA |
| NC\_018515.1\|provirus\_3418112\_3436097\_3144 | 423 | GENOMAD.001212.VV | PF08890 | Phage XkdN-like tail assembly chaperone protein, TAC |
| NC\_018515.1\|provirus\_3418112\_3436097\_3145 | 468 | GENOMAD.011307.VV | PF09393 | Phage tail tube protein |
| NC\_018515.1\|provirus\_3418112\_3436097\_3146 | 1317 | GENOMAD.013578.VV | PF17481;PF04984;PF17482;COG4386 | Mu-like prophage tail sheath protein gpL |
| NC\_018515.1\|provirus\_3418112\_3436097\_3147 | 183 | NA | NA | NA |
| NC\_018515.1\|provirus\_3418112\_3436097\_3148 | 420 | GENOMAD.023771.VV | PF20765 | Phage tail terminator protein |
| NC\_018515.1\|provirus\_3418112\_3436097\_3149 | 471 | NA | NA | NA |
| NC\_018515.1\|provirus\_3418112\_3436097\_3150 | 597 | GENOMAD.220722.VV | COG3655 | DNA-binding transcriptional regulator, XRE family |
| NC\_018515.1\|provirus\_3418112\_3436097\_3151 | 480 | GENOMAD.123021.VV | PF06114;COG2856 | IrrE N-terminal-like domain |

###### 2.18.2.2 Phage ID: NC\_018515.1|provirus\_4659544\_4693604

**Phage-Host Genome Ideogram:**

**Genes Annotation (geNomad):**

| gene | length | marker | annotation\_accessions | annotation\_description |
| --- | --- | --- | --- | --- |
| NC\_018515.1\|provirus\_4659544\_4693604\_4266 | 246 | NA | NA | NA |
| NC\_018515.1\|provirus\_4659544\_4693604\_4267 | 783 | GENOMAD.053051.VV | PF10772;PF02086;TIGR00571;COG0338 | Bacteriophage HP1, Orf24; D12 class N6 adenine-specific DNA methyltransferase |
| NC\_018515.1\|provirus\_4659544\_4693604\_4268 | 726 | NA | NA | NA |
| NC\_018515.1\|provirus\_4659544\_4693604\_4269 | 1290 | NA | NA | NA |
| NC\_018515.1\|provirus\_4659544\_4693604\_4270 | 831 | NA | NA | NA |
| NC\_018515.1\|provirus\_4659544\_4693604\_4271 | 501 | GENOMAD.220913.VP | PF09682;TIGR01673 | Bacteriophage holin of superfamily 6 (Holin\_LLH) |
| NC\_018515.1\|provirus\_4659544\_4693604\_4272 | 846 | NA | NA | NA |
| NC\_018515.1\|provirus\_4659544\_4693604\_4273 | 366 | NA | NA | NA |
| NC\_018515.1\|provirus\_4659544\_4693604\_4274 | 255 | NA | NA | NA |
| NC\_018515.1\|provirus\_4659544\_4693604\_4275 | 177 | GENOMAD.202651.VV | NA | NA |
| NC\_018515.1\|provirus\_4659544\_4693604\_4276 | 279 | NA | NA | NA |
| NC\_018515.1\|provirus\_4659544\_4693604\_4277 | 1095 | GENOMAD.135924.VV | NA | NA |
| NC\_018515.1\|provirus\_4659544\_4693604\_4278 | 1377 | GENOMAD.007293.VV | NA | NA |
| NC\_018515.1\|provirus\_4659544\_4693604\_4279 | 1092 | GENOMAD.006687.VV | PF14594;COG4926;TIGR01665 | Siphovirus ReqiPepy6 Gp37-like protein |
| NC\_018515.1\|provirus\_4659544\_4693604\_4280 | 864 | GENOMAD.009232.VV | PF16774;COG4722;TIGR01633 | Phage-related protein |
| NC\_018515.1\|provirus\_4659544\_4693604\_4281 | 2739 | GENOMAD.110013.VP | COG5280 | Phage-related minor tail protein |
| NC\_018515.1\|provirus\_4659544\_4693604\_4282 | 228 | GENOMAD.095592.VV | NA | NA |
| NC\_018515.1\|provirus\_4659544\_4693604\_4283 | 327 | GENOMAD.044337.VV | NA | NA |
| NC\_018515.1\|provirus\_4659544\_4693604\_4284 | 564 | GENOMAD.009652.VV | TIGR01537 | NA |
| NC\_018515.1\|provirus\_4659544\_4693604\_4285 | 360 | GENOMAD.058243.VV | PF11367 | NA |
| NC\_018515.1\|provirus\_4659544\_4693604\_4286 | 426 | GENOMAD.072075.VV | PF11114;TIGR01725;COG5005 | phage protein, HK97 gp10 family |
| NC\_018515.1\|provirus\_4659544\_4693604\_4287 | 312 | GENOMAD.008635.VV | PF05521;TIGR01563;COG5614 | phage head-tail adaptor, putative, SPP1 family |
| NC\_018515.1\|provirus\_4659544\_4693604\_4288 | 270 | GENOMAD.053459.VV | NA | NA |
| NC\_018515.1\|provirus\_4659544\_4693604\_4289 | 198 | GENOMAD.188189.VV | NA | NA |
| NC\_018515.1\|provirus\_4659544\_4693604\_4290 | 1224 | GENOMAD.088359.VV | PF05065;PF18316;PF17078;TIGR01554;COG4653 | phage major capsid protein, HK97 family |
| NC\_018515.1\|provirus\_4659544\_4693604\_4291 | 621 | GENOMAD.116539.VV | PF04586;PF05065;K06904;COG3740;TIGR01543 | Phage head maturation protease |
| NC\_018515.1\|provirus\_4659544\_4693604\_4292 | 1230 | NA | NA | NA |
| NC\_018515.1\|provirus\_4659544\_4693604\_4293 | 1755 | GENOMAD.194580.VP | PF03354;PF05521;PF20441;COG4626;TIGR01563 | Phage terminase-like protein, large subunit, contains N-terminal HTH domain |
| NC\_018515.1\|provirus\_4659544\_4693604\_4294 | 384 | GENOMAD.092852.VV | TIGR01558 | NA |
| NC\_018515.1\|provirus\_4659544\_4693604\_4295 | 363 | GENOMAD.080571.VV | NA | NA |
| NC\_018515.1\|provirus\_4659544\_4693604\_4296 | 786 | NA | NA | NA |
| NC\_018515.1\|provirus\_4659544\_4693604\_4297 | 849 | NA | NA | NA |
| NC\_018515.1\|provirus\_4659544\_4693604\_4298 | 513 | NA | NA | NA |
| NC\_018515.1\|provirus\_4659544\_4693604\_4299 | 363 | GENOMAD.151656.VP | PF17288;K06909;TIGR01547;COG1783 | phage terminase, large subunit, PBSX family |
| NC\_018515.1\|provirus\_4659544\_4693604\_4300 | 522 | NA | NA | NA |
| NC\_018515.1\|provirus\_4659544\_4693604\_4301 | 975 | GENOMAD.017981.VV | NA | NA |
| NC\_018515.1\|provirus\_4659544\_4693604\_4302 | 168 | NA | NA | NA |
| NC\_018515.1\|provirus\_4659544\_4693604\_4303 | 744 | GENOMAD.129273.VC | PF12706;TIGR02651;COG1234;K06167 | ribonuclease Z |
| NC\_018515.1\|provirus\_4659544\_4693604\_4304 | 828 | GENOMAD.121864.VP | PF03837;COG3723;K07455;TIGR00616 | Recombinational DNA repair protein RecT |
| NC\_018515.1\|provirus\_4659544\_4693604\_4305 | 108 | NA | NA | NA |
| NC\_018515.1\|provirus\_4659544\_4693604\_4306 | 1965 | GENOMAD.062081.VV | PF04423;COG1579 | Predicted nucleic acid-binding protein, contains Zn-ribbon domain |
| NC\_018515.1\|provirus\_4659544\_4693604\_4307 | 387 | NA | NA | NA |
| NC\_018515.1\|provirus\_4659544\_4693604\_4308 | 273 | GENOMAD.199250.VV | NA | NA |
| NC\_018515.1\|provirus\_4659544\_4693604\_4309 | 264 | NA | NA | NA |
| NC\_018515.1\|provirus\_4659544\_4693604\_4310 | 267 | NA | NA | NA |
| NC\_018515.1\|provirus\_4659544\_4693604\_4311 | 114 | NA | NA | NA |

##### 2.18.3 vOTUs

When more than 1 phage is detected in the host genome, we perform a
clustering step using the tool dRep.

A threshold of 0.95 was applied to the ANI similarity index to define
clusters of virus operational taxonomic units (vOTUs).

**Final cluster designations**

| genomad\_summary$seq\_name | genome | secondary\_cluster | threshold | cluster\_method | comparison\_algorithm | primary\_cluster |
| --- | --- | --- | --- | --- | --- | --- |
| NC\_018515.1\|provirus\_3418112\_3436097 | sequence\_000000.fasta.fasta | 1\_0 | 0.05 | average | ANImf | 1 |
| NC\_018515.1\|provirus\_4659544\_4693604 | sequence\_000001.fasta.fasta | 2\_0 | 0.05 | average | ANImf | 2 |

**Primary clustering plot**

##### 2.18.4 Prophage-Host Prediction

Prediction of potential hosts for the prophage contigs.

| virus | host\_genome | host\_taxonomy | main\_method | confidence\_score | additional\_methods |
| --- | --- | --- | --- | --- | --- |
| NC\_018515.1\|provirus\_3418112\_3436097 | RS\_GCF\_000231385.2 | d\_\_Bacteria;p\_\_Bacillota\_B;c\_\_Desulfitobacteriia;o\_\_Desulfitobacteriales;f\_\_Desulfitobacteriaceae;g\_\_Desulfosporosinus;s\_\_Desulfosporosinus meridiei | blast | 95.9 | iPHoP-RF;79.80 |
| NC\_018515.1\|provirus\_3418112\_3436097 | RS\_GCF\_004766055.1 | d\_\_Bacteria;p\_\_Bacillota\_B;c\_\_Desulfitobacteriia;o\_\_Desulfitobacteriales;f\_\_Desulfitobacteriaceae;g\_\_Desulfosporosinus;s\_\_Desulfosporosinus sp004766055 | iPHoP-RF | 95.7 | None |
| NC\_018515.1\|provirus\_3418112\_3436097 | GB\_GCA\_002404215.1 | d\_\_Bacteria;p\_\_Bacillota\_B;c\_\_Desulfitobacteriia;o\_\_Desulfitobacteriales;f\_\_Desulfitobacteriaceae;g\_\_Desulfosporosinus;s\_\_Desulfosporosinus sp002404215 | iPHoP-RF | 95.1 | None |
| NC\_018515.1\|provirus\_3418112\_3436097 | 3300011997\_21 | d\_\_Bacteria;p\_\_Bacillota\_B;c\_\_Desulfitobacteriia;o\_\_Desulfitobacteriales;f\_\_Desulfitobacteriaceae;g**Desulfosporosinus;s** | iPHoP-RF | 94.7 | None |
| NC\_018515.1\|provirus\_3418112\_3436097 | GB\_GCA\_021779415.1 | d\_\_Bacteria;p\_\_Bacillota\_B;c\_\_Desulfitobacteriia;o\_\_Desulfitobacteriales;f\_\_Desulfitobacteriaceae;g\_\_Desulfosporosinus;s\_\_Desulfosporosinus sp021779415 | iPHoP-RF | 92.8 | None |
| NC\_018515.1\|provirus\_3418112\_3436097 | RS\_GCF\_023897015.1 | d\_\_Bacteria;p\_\_Bacillota\_B;c\_\_Desulfitobacteriia;o\_\_Desulfitobacteriales;f\_\_Desulfitobacteriaceae;g\_\_Desulfosporosinus;s\_\_Desulfosporosinus nitroreducens | blast | 91.8 | iPHoP-RF;73.20 |
| NC\_018515.1\|provirus\_3418112\_3436097 | GB\_GCA\_003132105.1 | d\_\_Bacteria;p\_\_Bacillota\_B;c\_\_Desulfitobacteriia;o\_\_Desulfitobacteriales;f\_\_Desulfitobacteriaceae;g\_\_Desulfosporosinus;s\_\_Desulfosporosinus sp003132105 | iPHoP-RF | 91.4 | None |
| NC\_018515.1\|provirus\_3418112\_3436097 | RS\_GCF\_020595055.1 | d\_\_Bacteria;p\_\_Bacillota\_B;c\_\_Desulfitobacteriia;o\_\_Desulfitobacteriales;f\_\_Desulfitobacteriaceae;g\_\_Desulfosporosinus;s\_\_Desulfosporosinus sp020595055 | blast | 91.4 | iPHoP-RF;74.10 |
| NC\_018515.1\|provirus\_3418112\_3436097 | RS\_GCF\_000235605.1 | d\_\_Bacteria;p\_\_Bacillota\_B;c\_\_Desulfitobacteriia;o\_\_Desulfitobacteriales;f\_\_Desulfitobacteriaceae;g\_\_Desulfosporosinus;s\_\_Desulfosporosinus orientis | blast | 90.7 | iPHoP-RF;73.00 |
| NC\_018515.1\|provirus\_3418112\_3436097 | RS\_GCF\_900100785.1 | d\_\_Bacteria;p\_\_Bacillota\_B;c\_\_Desulfitobacteriia;o\_\_Desulfitobacteriales;f\_\_Desulfitobacteriaceae;g\_\_Desulfosporosinus;s\_\_Desulfosporosinus hippei | blast | 90.3 | iPHoP-RF;71.70 |
| NC\_018515.1\|provirus\_4659544\_4693604 | RS\_GCF\_000231385.2 | d\_\_Bacteria;p\_\_Bacillota\_B;c\_\_Desulfitobacteriia;o\_\_Desulfitobacteriales;f\_\_Desulfitobacteriaceae;g\_\_Desulfosporosinus;s\_\_Desulfosporosinus meridiei | blast | 96.5 | iPHoP-RF;74.60 |
| NC\_018515.1\|provirus\_4659544\_4693604 | RS\_GCF\_023897015.1 | d\_\_Bacteria;p\_\_Bacillota\_B;c\_\_Desulfitobacteriia;o\_\_Desulfitobacteriales;f\_\_Desulfitobacteriaceae;g\_\_Desulfosporosinus;s\_\_Desulfosporosinus nitroreducens | blast | 95.6 | iPHoP-RF;78.00 |
| NC\_018515.1\|provirus\_4659544\_4693604 | RS\_GCF\_900100785.1 | d\_\_Bacteria;p\_\_Bacillota\_B;c\_\_Desulfitobacteriia;o\_\_Desulfitobacteriales;f\_\_Desulfitobacteriaceae;g\_\_Desulfosporosinus;s\_\_Desulfosporosinus hippei | blast | 92.3 | iPHoP-RF;69.50 |
| NC\_018515.1\|provirus\_4659544\_4693604 | 3300011997\_21 | d\_\_Bacteria;p\_\_Bacillota\_B;c\_\_Desulfitobacteriia;o\_\_Desulfitobacteriales;f\_\_Desulfitobacteriaceae;g**Desulfosporosinus;s** | iPHoP-RF | 91.4 | None |
| NC\_018515.1\|provirus\_4659544\_4693604 | GB\_GCA\_002404215.1 | d\_\_Bacteria;p\_\_Bacillota\_B;c\_\_Desulfitobacteriia;o\_\_Desulfitobacteriales;f\_\_Desulfitobacteriaceae;g\_\_Desulfosporosinus;s\_\_Desulfosporosinus sp002404215 | iPHoP-RF | 91.1 | None |
| NC\_018515.1\|provirus\_4659544\_4693604 | RS\_GCF\_000244895.1 | d\_\_Bacteria;p\_\_Bacillota\_B;c\_\_Desulfitobacteriia;o\_\_Desulfitobacteriales;f\_\_Desulfitobacteriaceae;g\_\_Desulfosporosinus;s\_\_Desulfosporosinus youngiae | blast | 90.6 | iPHoP-RF;82.50 |
| NC\_018515.1\|provirus\_4659544\_4693604 | GB\_GCA\_021779415.1 | d\_\_Bacteria;p\_\_Bacillota\_B;c\_\_Desulfitobacteriia;o\_\_Desulfitobacteriales;f\_\_Desulfitobacteriaceae;g\_\_Desulfosporosinus;s\_\_Desulfosporosinus sp021779415 | iPHoP-RF | 90.1 | None |
| NC\_018515.1\|provirus\_4659544\_4693604 | GB\_GCA\_900290375.1 | d\_\_Bacteria;p\_\_Bacillota\_B;c\_\_Desulfitobacteriia;o\_\_Desulfitobacteriales;f\_\_Desulfitobacteriaceae;g\_\_Desulfosporosinus;s\_\_Desulfosporosinus infrequens | iPHoP-RF | 90.1 | None |

##### 2.18.5 AMG Predictions

Prediction of auxiliary metabolic genes in the prophage contigs.

| protein | scaffold | amg\_ko | amg\_ko\_name | pfam | pfam\_name |
| --- | --- | --- | --- | --- | --- |
| NC\_018515.1\|provirus\_3418112\_3436097\_1 | NC\_018515.1\|provirus\_3418112\_3436097 | K00106 | XDH; xanthine dehydrogenase/oxidase [EC:1.17.1.4 1.17.3.2] | PF02738.18 | Molybdopterin-binding domain of aldehyde dehydrogenase |
| NC\_018515.1\|provirus\_4659544\_4693604\_38 | NC\_018515.1\|provirus\_4659544\_4693604 | K06167 | phnP; phosphoribosyl 1,2-cyclic phosphate phosphodiesterase [EC:3.1.4.55] | PF00753.27 | Metallo-beta-lactamase superfamily |

## 2.19 NC\_019897

##### 2.19.1 Host Genome

**GTDB-Tk taxonomy**:

classification

d\_\_Bacteria;p\_\_Bacillota;c\_\_Bacilli;o\_\_Paenibacillales;f\_\_Paenibacillaceae;g\_\_Thermobacillus;s\_\_Thermobacillus
xylanilyticus

**CheckM2 Summary**:

| total\_contigs | contig\_n50 | completeness | contamination |
| --- | --- | --- | --- |
| 1 | 4206343 | 99.95 | 0.34 |

**Defense-Finder Systems**:

| sys\_id | type | subtype | sys\_beg | sys\_end | protein\_in\_syst | genes\_count | name\_of\_profiles\_in\_sys |
| --- | --- | --- | --- | --- | --- | --- | --- |
| NC\_019897.1\_RM\_Type\_III\_11 | RM | RM\_Type\_III | NC\_019897.1\_86 | NC\_019897.1\_87 | NC\_019897.1\_86,NC\_019897.1\_87 | 2 | RM\_Type\_III\_\_Type\_III\_MTases\_FAM\_0,RM\_Type\_III\_\_Type\_III\_REases\_FAM\_0.einsi\_trimmed |
| NC\_019897.1\_MazEF\_3 | MazEF | MazEF | NC\_019897.1\_128 | NC\_019897.1\_129 | NC\_019897.1\_128,NC\_019897.1\_129 | 2 | MazEF\_\_MazE,MazEF\_\_MazF |
| NC\_019897.1\_RM\_Type\_II\_9 | RM | RM\_Type\_II | NC\_019897.1\_789 | NC\_019897.1\_790 | NC\_019897.1\_789,NC\_019897.1\_790 | 2 | RM\_Type\_II\_\_Type\_II\_MTases\_FAM\_2,RM\_Type\_II\_\_Type\_II\_REase27 |
| NC\_019897.1\_Mokosh\_TypeII\_4 | Mokosh | Mokosh\_TypeII | NC\_019897.1\_913 | NC\_019897.1\_913 | NC\_019897.1\_913 | 1 | Mokosh\_TypeII\_\_MkoC |
| NC\_019897.1\_Druantia\_I\_2 | Druantia | Druantia\_I | NC\_019897.1\_2741 | NC\_019897.1\_2744 | NC\_019897.1\_2741,NC\_019897.1\_2742,NC\_019897.1\_2743,NC\_019897.1\_2744 | 4 | Druantia\_I\_\_DruB,Druantia\_I\_\_DruC,Druantia\_I\_\_DruD,Druantia\_\_DruE\_1 |
| NC\_019897.1\_CAS\_Class1-Subtype-III-D\_14 | Cas | CAS\_Class1-Subtype-III-D | NC\_019897.1\_3030 | NC\_019897.1\_3035 | NC\_019897.1\_3030,NC\_019897.1\_3031,NC\_019897.1\_3032,NC\_019897.1\_3033,NC\_019897.1\_3034,NC\_019897.1\_3035 | 6 | cas10\_III-D\_3,csm3gr7\_III-D\_2,csm3gr7\_III-D\_3,csm3gr7\_III\_1,csx19\_III-D\_17,csx1\_III\_11 |
| NC\_019897.1\_CAS\_Class1-Subtype-I-B\_12 | Cas | CAS\_Class1-Subtype-I-B | NC\_019897.1\_3046 | NC\_019897.1\_3053 | NC\_019897.1\_3046,NC\_019897.1\_3047,NC\_019897.1\_3048,NC\_019897.1\_3049,NC\_019897.1\_3050,NC\_019897.1\_3051,NC\_019897.1\_3052,NC\_019897.1\_3053 | 8 | cas1\_I\_II\_III\_IV\_V\_VI\_7,cas2\_I\_II\_III\_IV\_V\_VI\_3,cas3\_I\_2,cas4\_I\_II\_III\_IV\_V\_VI\_6,cas5\_I-B\_17,cas6\_I\_II\_III\_IV\_V\_VI\_20,cas7b\_I-B\_I-C\_2,cas8b1\_I-B\_8 |
| NC\_019897.1\_CAS\_Class1-Subtype-I-C\_13 | Cas | CAS\_Class1-Subtype-I-C | NC\_019897.1\_3155 | NC\_019897.1\_3158 | NC\_019897.1\_3155,NC\_019897.1\_3156,NC\_019897.1\_3157,NC\_019897.1\_3158 | 4 | cas3\_I\_5,cas5\_I-C\_5,cas7\_I-C\_7,cas8c\_I-C\_4 |
| NC\_019897.1\_Wadjet\_II\_6 | Wadjet | Wadjet\_II | NC\_019897.1\_3286 | NC\_019897.1\_3289 | NC\_019897.1\_3286,NC\_019897.1\_3287,NC\_019897.1\_3288,NC\_019897.1\_3289 | 4 | Wadjet\_\_JetA\_II,Wadjet\_\_JetB\_II,Wadjet\_\_JetC\_II,Wadjet\_\_JetD\_II |
| NC\_019897.1\_RM\_Type\_I\_8 | RM | RM\_Type\_I | NC\_019897.1\_3326 | NC\_019897.1\_3328 | NC\_019897.1\_3326,NC\_019897.1\_3327,NC\_019897.1\_3328 | 3 | RM\_\_Type\_I\_MTases\_FAM\_1,RM\_\_Type\_I\_REases\_FAM\_2.einsi\_trimmed,RM\_\_Type\_I\_S\_51 |
| NC\_019897.1\_RM\_Type\_II\_10 | RM | RM\_Type\_II | NC\_019897.1\_3527 | NC\_019897.1\_3528 | NC\_019897.1\_3527,NC\_019897.1\_3528 | 2 | RM\_Type\_II\_\_Type\_II\_MTases\_FAM\_16,RM\_Type\_II\_\_Type\_II\_REase01 |
| NC\_019897.1\_Ceres\_1 | Ceres | Ceres | NC\_019897.1\_3704 | NC\_019897.1\_3704 | NC\_019897.1\_3704 | 1 | Ceres\_\_CrsA1 |

## 2.20 NC\_019904

##### 2.20.1 Host Genome

**GTDB-Tk taxonomy**:

classification

d\_\_Bacteria;p\_\_Bacteroidota;c\_\_Bacteroidia;o\_\_Cytophagales;f\_\_Cyclobacteriaceae;g\_\_Echinicola;s\_\_Echinicola
vietnamensis

**CheckM2 Summary**:

| total\_contigs | contig\_n50 | completeness | contamination |
| --- | --- | --- | --- |
| 1 | 5608040 | 100 | 0.23 |

**Defense-Finder Systems**:

| sys\_id | type | subtype | sys\_beg | sys\_end | protein\_in\_syst | genes\_count | name\_of\_profiles\_in\_sys |
| --- | --- | --- | --- | --- | --- | --- | --- |
| NC\_019904.1\_HEC-05\_12 | HEC-05 | HEC-05 | NC\_019904.1\_318 | NC\_019904.1\_318 | NC\_019904.1\_318 | 1 | HEC-05\_\_HEC-05 |
| NC\_019904.1\_RM\_Type\_I\_19 | RM | RM\_Type\_I | NC\_019904.1\_371 | NC\_019904.1\_374 | NC\_019904.1\_371,NC\_019904.1\_372,NC\_019904.1\_374 | 3 | RM\_\_Type\_I\_MTases\_FAM\_1,RM\_\_Type\_I\_REases\_FAM\_2.einsi\_trimmed,RM\_\_Type\_I\_S\_01 |
| NC\_019904.1\_dCTPdeaminase\_15 | dCTPdeaminase | dCTPdeaminase | NC\_019904.1\_408 | NC\_019904.1\_408 | NC\_019904.1\_408 | 1 | dCTPdeaminase\_\_dCTPdeaminase |
| NC\_019904.1\_RM\_Type\_I\_20 | RM | RM\_Type\_I | NC\_019904.1\_411 | NC\_019904.1\_413 | NC\_019904.1\_411,NC\_019904.1\_412,NC\_019904.1\_413 | 3 | RM\_\_Type\_I\_MTases\_FAM\_1,RM\_\_Type\_I\_REases\_FAM\_2.einsi\_trimmed,RM\_\_Type\_I\_S\_03 |
| NC\_019904.1\_Gabija\_8 | Gabija | Gabija | NC\_019904.1\_430 | NC\_019904.1\_431 | NC\_019904.1\_430,NC\_019904.1\_431 | 2 | Gabija\_\_GajA,Gabija\_\_GajB\_2 |
| NC\_019904.1\_Gao\_Qat\_11 | Gao\_Qat | Gao\_Qat | NC\_019904.1\_1323 | NC\_019904.1\_1326 | NC\_019904.1\_1323,NC\_019904.1\_1324,NC\_019904.1\_1325,NC\_019904.1\_1326 | 4 | Gao\_Qat\_\_QatA,Gao\_Qat\_\_QatB,Gao\_Qat\_\_QatC,Gao\_Qat\_\_QatD |
| NC\_019904.1\_SoFic\_14 | SoFIC | SoFic | NC\_019904.1\_1676 | NC\_019904.1\_1676 | NC\_019904.1\_1676 | 1 | SoFic\_\_SoFic |
| NC\_019904.1\_Gabija\_9 | Gabija | Gabija | NC\_019904.1\_1900 | NC\_019904.1\_1901 | NC\_019904.1\_1900,NC\_019904.1\_1901 | 2 | Gabija\_\_GajA,Gabija\_\_GajB\_3 |
| NC\_019904.1\_RM\_Type\_II\_22 | RM | RM\_Type\_II | NC\_019904.1\_1947 | NC\_019904.1\_1949 | NC\_019904.1\_1947,NC\_019904.1\_1948,NC\_019904.1\_1949 | 3 | RM\_Type\_II\_\_Type\_II\_MTases\_FAM\_2,RM\_Type\_II\_\_Type\_II\_REase06,RM\_Type\_II\_\_Type\_II\_REase38 |
| NC\_019904.1\_Sirona\_13 | Sirona | Sirona | NC\_019904.1\_1950 | NC\_019904.1\_1950 | NC\_019904.1\_1950 | 1 | Sirona\_\_VCA0356 |
| NC\_019904.1\_CBASS\_II\_5 | CBASS | CBASS\_II | NC\_019904.1\_3143 | NC\_019904.1\_3145 | NC\_019904.1\_3143,NC\_019904.1\_3144,NC\_019904.1\_3145 | 3 | CBASS\_\_Cyclase\_II,CBASS\_\_E2,CBASS\_\_Effector\_2TM\_Sa\_NUDIX |
| NC\_019904.1\_AbiE\_2 | AbiE | AbiE | NC\_019904.1\_3195 | NC\_019904.1\_3196 | NC\_019904.1\_3195,NC\_019904.1\_3196 | 2 | AbiEii\_\_AbiEi\_3,AbiEii\_\_AbiEii |
| NC\_019904.1\_RM\_Type\_I\_21 | RM | RM\_Type\_I | NC\_019904.1\_4383 | NC\_019904.1\_4389 | NC\_019904.1\_4383,NC\_019904.1\_4387,NC\_019904.1\_4388,NC\_019904.1\_4389 | 4 | RM\_\_Type\_I\_MTases\_FAM\_0,RM\_\_Type\_I\_REases\_FAM\_0.einsi\_trimmed,RM\_\_Type\_I\_S\_01,RM\_\_Type\_I\_S\_51 |
| NC\_019904.1\_AbiD\_1 | AbiD | AbiD | NC\_019904.1\_4385 | NC\_019904.1\_4385 | NC\_019904.1\_4385 | 1 | AbiD\_\_AbiD |
| NC\_019904.1\_Cernunnos\_7 | Cernunnos | Cernunnos | NC\_019904.1\_4390 | NC\_019904.1\_4390 | NC\_019904.1\_4390 | 1 | Cernunnos\_\_VCA0410 |
| NC\_019904.1\_Gabija\_10 | Gabija | Gabija | NC\_019904.1\_4391 | NC\_019904.1\_4392 | NC\_019904.1\_4391,NC\_019904.1\_4392 | 2 | Gabija\_\_GajA,Gabija\_\_GajB\_1 |
| NC\_019904.1\_Ceres\_6 | Ceres | Ceres | NC\_019904.1\_4414 | NC\_019904.1\_4414 | NC\_019904.1\_4414 | 1 | Ceres\_\_CrsA1 |

## 2.21 NC\_019936

##### 2.21.1 Host Genome

**GTDB-Tk taxonomy**:

classification

d\_\_Bacteria;p\_\_Pseudomonadota;c\_\_Gammaproteobacteria;o\_\_Pseudomonadales;f\_\_Pseudomonadaceae;g\_\_Stutzerimonas;s\_\_Stutzerimonas
stutzeri\_AE

**CheckM2 Summary**:

| total\_contigs | contig\_n50 | completeness | contamination |
| --- | --- | --- | --- |
| 1 | 4575057 | 100 | 0.13 |

**Defense-Finder Systems**:

| sys\_id | type | subtype | sys\_beg | sys\_end | protein\_in\_syst | genes\_count | name\_of\_profiles\_in\_sys |
| --- | --- | --- | --- | --- | --- | --- | --- |
| NC\_019936.1\_Gabija\_5 | Gabija | Gabija | NC\_019936.1\_617 | NC\_019936.1\_618 | NC\_019936.1\_617,NC\_019936.1\_618 | 2 | Gabija\_\_GajA,Gabija\_\_GajB\_3 |
| NC\_019936.1\_RM\_Type\_I\_12 | RM | RM\_Type\_I | NC\_019936.1\_860 | NC\_019936.1\_862 | NC\_019936.1\_860,NC\_019936.1\_861,NC\_019936.1\_862 | 3 | RM\_\_Type\_I\_MTases\_FAM\_3,RM\_\_Type\_I\_REases\_FAM\_1.einsi\_trimmed,RM\_\_Type\_I\_S\_52 |
| NC\_019936.1\_Shango\_6 | Shango | Shango | NC\_019936.1\_896 | NC\_019936.1\_898 | NC\_019936.1\_896,NC\_019936.1\_897,NC\_019936.1\_898 | 3 | Shango\_\_SngA,Shango\_\_SngB,Shango\_\_SngC |
| NC\_019936.1\_RM\_Type\_I\_13 | RM | RM\_Type\_I | NC\_019936.1\_1070 | NC\_019936.1\_1072 | NC\_019936.1\_1070,NC\_019936.1\_1071,NC\_019936.1\_1072 | 3 | RM\_\_Type\_I\_MTases\_FAM\_1,RM\_\_Type\_I\_REases\_FAM\_2.einsi\_trimmed,RM\_\_Type\_I\_S\_03 |
| NC\_019936.1\_RM\_Type\_IV\_16 | RM | RM\_Type\_IV | NC\_019936.1\_1377 | NC\_019936.1\_1377 | NC\_019936.1\_1377 | 1 | RM\_Type\_IV\_\_FAM\_0 |
| NC\_019936.1\_AbiC\_1 | AbiC | AbiC | NC\_019936.1\_2817 | NC\_019936.1\_2817 | NC\_019936.1\_2817 | 1 | AbiC\_\_AbiC |
| NC\_019936.1\_RM\_Type\_I\_14 | RM | RM\_Type\_I | NC\_019936.1\_3312 | NC\_019936.1\_3317 | NC\_019936.1\_3312,NC\_019936.1\_3313,NC\_019936.1\_3314,NC\_019936.1\_3317 | 4 | RM\_\_Type\_I\_MTases\_FAM\_3,RM\_\_Type\_I\_REases\_FAM\_1.einsi\_trimmed,RM\_\_Type\_I\_REases\_FAM\_2.einsi\_trimmed,RM\_\_Type\_I\_S\_52 |
| NC\_019936.1\_CBASS\_III\_4 | CBASS | CBASS\_III | NC\_019936.1\_3322 | NC\_019936.1\_3325 | NC\_019936.1\_3322,NC\_019936.1\_3323,NC\_019936.1\_3324,NC\_019936.1\_3325 | 4 | CBASS\_\_Cyclase\_II,CBASS\_\_Endonuc\_big,CBASS\_\_TRIP13,CBASS\_\_bacHORMA\_1 |
| NC\_019936.1\_RM\_Type\_I\_15 | RM | RM\_Type\_I | NC\_019936.1\_3936 | NC\_019936.1\_3940 | NC\_019936.1\_3936,NC\_019936.1\_3939,NC\_019936.1\_3940 | 3 | RM\_\_Type\_I\_MTases\_FAM\_0,RM\_\_Type\_I\_REases\_FAM\_0.einsi\_trimmed,RM\_\_Type\_I\_S\_06 |
| NC\_019936.1\_dCTPdeaminase\_7 | dCTPdeaminase | dCTPdeaminase | NC\_019936.1\_4004 | NC\_019936.1\_4004 | NC\_019936.1\_4004 | 1 | dCTPdeaminase\_\_dCTPdeaminase |

**Genomad and CheckV Summary**:

| seq\_name | length | gene\_count | viral\_genes | checkv\_quality | miuvig\_quality | taxonomy | topology | coordinates |
| --- | --- | --- | --- | --- | --- | --- | --- | --- |
| NC\_019936.1\|provirus\_4296137\_4304306 | 8170 | 12 | 2 | High-quality | High-quality | Viruses;Monodnaviria;Loebvirae;Hofneiviricota;Faserviricetes;Tubulavirales;Inoviridae | Provirus | 4296137-4304306 |
| NC\_019936.1\|provirus\_3233292\_3241441 | 8150 | 14 | 3 | Medium-quality | Genome-fragment | Viruses;Monodnaviria;Loebvirae;Hofneiviricota;Faserviricetes;Tubulavirales;Inoviridae | Provirus | 3233292-3241441 |
| NC\_019936.1\|provirus\_2346826\_2361929 | 15104 | 25 | 2 | High-quality | High-quality | Viruses;Monodnaviria;Loebvirae;Hofneiviricota;Faserviricetes;Tubulavirales;Inoviridae | Provirus | 2346826-2361929 |

**Host Genome Ideogram with Phages**:

In
this circular plot, **“c”** indicates the contig, and the
number that follows (e.g., **c1**) represents the contig
number. If multiple contigs are present in the genome, each will be
shown with a distinct label (e.g., **c1**,
**c2**, etc.).

##### 2.21.2 Prophages

**Select prophage to show:**

###### 2.21.2.1 Phage ID: NC\_019936.1|provirus\_4296137\_4304306

**Phage-Host Genome Ideogram:**

**Genes Annotation (geNomad):**

| gene | length | marker | annotation\_accessions | annotation\_description |
| --- | --- | --- | --- | --- |
| NC\_019936.1\|provirus\_4296137\_4304306\_3987 | 216 | NA | NA | NA |
| NC\_019936.1\|provirus\_4296137\_4304306\_3988 | 582 | NA | NA | NA |
| NC\_019936.1\|provirus\_4296137\_4304306\_3989 | 372 | GENOMAD.225482.VP | NA | NA |
| NC\_019936.1\|provirus\_4296137\_4304306\_3990 | 180 | GENOMAD.226094.VP | NA | NA |
| NC\_019936.1\|provirus\_4296137\_4304306\_3991 | 216 | GENOMAD.225528.VP | NA | NA |
| NC\_019936.1\|provirus\_4296137\_4304306\_3992 | 210 | GENOMAD.224287.VP | PF05356 | Inovirus Coat protein B |
| NC\_019936.1\|provirus\_4296137\_4304306\_3993 | 1494 | GENOMAD.222416.VP | NA | NA |
| NC\_019936.1\|provirus\_4296137\_4304306\_3994 | 327 | GENOMAD.224290.VP | PF10734 | NA |
| NC\_019936.1\|provirus\_4296137\_4304306\_3995 | 1185 | GENOMAD.090645.VV | NA | NA |
| NC\_019936.1\|provirus\_4296137\_4304306\_3996 | 1281 | GENOMAD.226808.VP | PF02486 | Replication initiation factor |
| NC\_019936.1\|provirus\_4296137\_4304306\_3997 | 1059 | GENOMAD.212346.VP | PF16452;PF06892;COG1974 | Bacteriophage CI repressor C-terminal domain; Phage regulatory protein CII (CP76) |
| NC\_019936.1\|provirus\_4296137\_4304306\_3998 | 318 | GENOMAD.200462.VV | NA | NA |

###### 2.21.2.2 Phage ID: NC\_019936.1|provirus\_3233292\_3241441

**Phage-Host Genome Ideogram:**

**Genes Annotation (geNomad):**

| gene | length | marker | annotation\_accessions | annotation\_description |
| --- | --- | --- | --- | --- |
| NC\_019936.1\|provirus\_3233292\_3241441\_2952 | 1008 | GENOMAD.212346.VP | PF16452;PF06892;COG1974 | Bacteriophage CI repressor C-terminal domain; Phage regulatory protein CII (CP76) |
| NC\_019936.1\|provirus\_3233292\_3241441\_2953 | 1278 | GENOMAD.226808.VP | PF02486 | Replication initiation factor |
| NC\_019936.1\|provirus\_3233292\_3241441\_2954 | 1185 | GENOMAD.090645.VV | NA | NA |
| NC\_019936.1\|provirus\_3233292\_3241441\_2955 | 327 | GENOMAD.224290.VP | PF10734 | NA |
| NC\_019936.1\|provirus\_3233292\_3241441\_2956 | 1494 | GENOMAD.222416.VP | NA | NA |
| NC\_019936.1\|provirus\_3233292\_3241441\_2957 | 129 | NA | NA | NA |
| NC\_019936.1\|provirus\_3233292\_3241441\_2958 | 210 | GENOMAD.224287.VP | PF05356 | Inovirus Coat protein B |
| NC\_019936.1\|provirus\_3233292\_3241441\_2959 | 216 | GENOMAD.225528.VP | NA | NA |
| NC\_019936.1\|provirus\_3233292\_3241441\_2960 | 180 | GENOMAD.226094.VP | NA | NA |
| NC\_019936.1\|provirus\_3233292\_3241441\_2961 | 360 | GENOMAD.225482.VP | NA | NA |
| NC\_019936.1\|provirus\_3233292\_3241441\_2962 | 771 | NA | NA | NA |
| NC\_019936.1\|provirus\_3233292\_3241441\_2963 | 144 | GENOMAD.213484.VP | PF05509;COG4877 | Plasmid stability protein |
| NC\_019936.1\|provirus\_3233292\_3241441\_2964 | 144 | NA | NA | NA |
| NC\_019936.1\|provirus\_3233292\_3241441\_2965 | 216 | NA | NA | NA |

###### 2.21.2.3 Phage ID: NC\_019936.1|provirus\_2346826\_2361929

**Phage-Host Genome Ideogram:**

**Genes Annotation (geNomad):**

| gene | length | marker | annotation\_accessions | annotation\_description |
| --- | --- | --- | --- | --- |
| NC\_019936.1\|provirus\_2346826\_2361929\_2121 | 288 | NA | NA | NA |
| NC\_019936.1\|provirus\_2346826\_2361929\_2122 | 213 | GENOMAD.221461.VP | NA | NA |
| NC\_019936.1\|provirus\_2346826\_2361929\_2123 | 1263 | GENOMAD.223897.VP | PF05155 | Phage X family |
| NC\_019936.1\|provirus\_2346826\_2361929\_2124 | 147 | NA | NA | NA |
| NC\_019936.1\|provirus\_2346826\_2361929\_2125 | 345 | GENOMAD.228002.VV | PF17426 | Putative Gamma DNA binding protein G5P |
| NC\_019936.1\|provirus\_2346826\_2361929\_2126 | 147 | NA | NA | NA |
| NC\_019936.1\|provirus\_2346826\_2361929\_2127 | 1512 | GENOMAD.222416.VP | NA | NA |
| NC\_019936.1\|provirus\_2346826\_2361929\_2128 | 267 | GENOMAD.197598.VP | NA | NA |
| NC\_019936.1\|provirus\_2346826\_2361929\_2129 | 1194 | NA | NA | NA |
| NC\_019936.1\|provirus\_2346826\_2361929\_2130 | 288 | NA | NA | NA |
| NC\_019936.1\|provirus\_2346826\_2361929\_2131 | 213 | GENOMAD.221461.VP | NA | NA |
| NC\_019936.1\|provirus\_2346826\_2361929\_2132 | 1263 | GENOMAD.223897.VP | PF05155 | Phage X family |
| NC\_019936.1\|provirus\_2346826\_2361929\_2133 | 147 | NA | NA | NA |
| NC\_019936.1\|provirus\_2346826\_2361929\_2134 | 345 | GENOMAD.228002.VV | PF17426 | Putative Gamma DNA binding protein G5P |
| NC\_019936.1\|provirus\_2346826\_2361929\_2135 | 147 | NA | NA | NA |
| NC\_019936.1\|provirus\_2346826\_2361929\_2136 | 1512 | GENOMAD.222416.VP | NA | NA |
| NC\_019936.1\|provirus\_2346826\_2361929\_2137 | 267 | GENOMAD.197598.VP | NA | NA |
| NC\_019936.1\|provirus\_2346826\_2361929\_2138 | 1194 | NA | NA | NA |
| NC\_019936.1\|provirus\_2346826\_2361929\_2139 | 288 | NA | NA | NA |
| NC\_019936.1\|provirus\_2346826\_2361929\_2140 | 360 | NA | NA | NA |
| NC\_019936.1\|provirus\_2346826\_2361929\_2141 | 285 | NA | NA | NA |
| NC\_019936.1\|provirus\_2346826\_2361929\_2142 | 288 | NA | NA | NA |

##### 2.21.3 vOTUs

When more than 1 phage is detected in the host genome, we perform a
clustering step using the tool dRep.

A threshold of 0.95 was applied to the ANI similarity index to define
clusters of virus operational taxonomic units (vOTUs).

**Final cluster designations**

| genomad\_summary$seq\_name | genome | secondary\_cluster | threshold | cluster\_method | comparison\_algorithm | primary\_cluster |
| --- | --- | --- | --- | --- | --- | --- |
| NC\_019936.1\|provirus\_4296137\_4304306 | sequence\_000001.fasta.fasta | 1\_1 | 0.05 | average | ANImf | 1 |
| NC\_019936.1\|provirus\_3233292\_3241441 | sequence\_000002.fasta.fasta | 1\_2 | 0.05 | average | ANImf | 1 |
| NC\_019936.1\|provirus\_2346826\_2361929 | sequence\_000000.fasta.fasta | 2\_0 | 0.05 | average | ANImf | 2 |

**Primary clustering plot**

##### 2.21.4 Prophage-Host Prediction

Prediction of potential hosts for the prophage contigs.

| virus | host\_genome | host\_taxonomy | main\_method | confidence\_score | additional\_methods |
| --- | --- | --- | --- | --- | --- |
| NC\_019936.1\|provirus\_2346826\_2361929 | RS\_GCF\_015291885.1 | d\_\_Bacteria;p\_\_Pseudomonadota;c\_\_Gammaproteobacteria;o\_\_Pseudomonadales;f\_\_Pseudomonadaceae;g\_\_Stutzerimonas;s\_\_Stutzerimonas stutzeri\_AC | CRISPR | 97.4 | blast;95.20 iPHoP-RF;85.40 |
| NC\_019936.1\|provirus\_2346826\_2361929 | RS\_GCF\_003640395.1 | d\_\_Bacteria;p\_\_Pseudomonadota;c\_\_Gammaproteobacteria;o\_\_Pseudomonadales;f\_\_Pseudomonadaceae;g\_\_Stutzerimonas;s\_\_Stutzerimonas urumqiensis | CRISPR | 96.9 | iPHoP-RF;52.20 |
| NC\_019936.1\|provirus\_2346826\_2361929 | RS\_GCF\_000219605.1 | d\_\_Bacteria;p\_\_Pseudomonadota;c\_\_Gammaproteobacteria;o\_\_Pseudomonadales;f\_\_Pseudomonadaceae;g\_\_Stutzerimonas;s\_\_Stutzerimonas stutzeri | blast | 96.2 | CRISPR;90.50 iPHoP-RF;82.80 |
| NC\_019936.1\|provirus\_2346826\_2361929 | RS\_GCF\_000327065.1 | d\_\_Bacteria;p\_\_Pseudomonadota;c\_\_Gammaproteobacteria;o\_\_Pseudomonadales;f\_\_Pseudomonadaceae;g\_\_Stutzerimonas;s\_\_Stutzerimonas stutzeri\_AE | blast | 95.9 | iPHoP-RF;64.00 |
| NC\_019936.1\|provirus\_2346826\_2361929 | RS\_GCF\_002929225.1 | d\_\_Bacteria;p\_\_Pseudomonadota;c\_\_Gammaproteobacteria;o\_\_Pseudomonadales;f\_\_Pseudomonadaceae;g\_\_Stutzerimonas;s\_\_Stutzerimonas stutzeri\_U | blast | 94.1 | iPHoP-RF;87.60 |
| NC\_019936.1\|provirus\_2346826\_2361929 | RS\_GCF\_003935375.1 | d\_\_Bacteria;p\_\_Pseudomonadota;c\_\_Gammaproteobacteria;o\_\_Pseudomonadales;f\_\_Pseudomonadaceae;g\_\_Stutzerimonas;s\_\_Stutzerimonas xanthomarina\_A | blast | 94.0 | iPHoP-RF;87.90 CRISPR;82.70 |
| NC\_019936.1\|provirus\_2346826\_2361929 | GB\_GCA\_007713455.1 | d\_\_Bacteria;p\_\_Pseudomonadota;c\_\_Gammaproteobacteria;o\_\_Pseudomonadales;f\_\_Pseudomonadaceae;g\_\_Stutzerimonas;s\_\_Stutzerimonas sp007713455 | blast | 93.1 | iPHoP-RF;76.60 |
| NC\_019936.1\|provirus\_2346826\_2361929 | RS\_GCF\_021432085.1 | d\_\_Bacteria;p\_\_Pseudomonadota;c\_\_Gammaproteobacteria;o\_\_Pseudomonadales;f\_\_Pseudomonadaceae;g\_\_Stutzerimonas;s\_\_Stutzerimonas kunmingensis\_A | blast | 93.1 | iPHoP-RF;89.80 |
| NC\_019936.1\|provirus\_2346826\_2361929 | RS\_GCF\_000341615.1 | d\_\_Bacteria;p\_\_Pseudomonadota;c\_\_Gammaproteobacteria;o\_\_Pseudomonadales;f\_\_Pseudomonadaceae;g\_\_Stutzerimonas;s\_\_Stutzerimonas stutzeri\_G | blast | 91.8 | iPHoP-RF;74.80 |
| NC\_019936.1\|provirus\_2346826\_2361929 | RS\_GCF\_003696285.1 | d\_\_Bacteria;p\_\_Pseudomonadota;c\_\_Gammaproteobacteria;o\_\_Pseudomonadales;f\_\_Pseudomonadaceae;g\_\_Stutzerimonas;s\_\_Stutzerimonas nitrititolerans | blast | 90.6 | iPHoP-RF;52.20 |
| NC\_019936.1\|provirus\_2346826\_2361929 | RS\_GCF\_015070855.1 | d\_\_Bacteria;p\_\_Pseudomonadota;c\_\_Gammaproteobacteria;o\_\_Pseudomonadales;f\_\_Pseudomonadaceae;g\_\_Stutzerimonas;s\_\_Stutzerimonas lopnurensis | blast | 90.6 | iPHoP-RF;79.80 |
| NC\_019936.1\|provirus\_3233292\_3241441 | RS\_GCF\_000219605.1 | d\_\_Bacteria;p\_\_Pseudomonadota;c\_\_Gammaproteobacteria;o\_\_Pseudomonadales;f\_\_Pseudomonadaceae;g\_\_Stutzerimonas;s\_\_Stutzerimonas stutzeri | blast | 96.9 | iPHoP-RF;86.70 |
| NC\_019936.1\|provirus\_3233292\_3241441 | RS\_GCF\_002929225.1 | d\_\_Bacteria;p\_\_Pseudomonadota;c\_\_Gammaproteobacteria;o\_\_Pseudomonadales;f\_\_Pseudomonadaceae;g\_\_Stutzerimonas;s\_\_Stutzerimonas stutzeri\_U | blast | 96.9 | iPHoP-RF;86.30 |
| NC\_019936.1\|provirus\_3233292\_3241441 | RS\_GCF\_003696285.1 | d\_\_Bacteria;p\_\_Pseudomonadota;c\_\_Gammaproteobacteria;o\_\_Pseudomonadales;f\_\_Pseudomonadaceae;g\_\_Stutzerimonas;s\_\_Stutzerimonas nitrititolerans | blast | 96.9 | None |
| NC\_019936.1\|provirus\_3233292\_3241441 | RS\_GCF\_000495915.1 | d\_\_Bacteria;p\_\_Pseudomonadota;c\_\_Gammaproteobacteria;o\_\_Pseudomonadales;f\_\_Pseudomonadaceae;g\_\_Stutzerimonas;s\_\_Stutzerimonas chloritidismutans | blast | 96.8 | iPHoP-RF;88.90 |
| NC\_019936.1\|provirus\_3233292\_3241441 | RS\_GCF\_003935375.1 | d\_\_Bacteria;p\_\_Pseudomonadota;c\_\_Gammaproteobacteria;o\_\_Pseudomonadales;f\_\_Pseudomonadaceae;g\_\_Stutzerimonas;s\_\_Stutzerimonas xanthomarina\_A | blast | 96.8 | iPHoP-RF;81.10 |
| NC\_019936.1\|provirus\_3233292\_3241441 | RS\_GCF\_015291885.1 | d\_\_Bacteria;p\_\_Pseudomonadota;c\_\_Gammaproteobacteria;o\_\_Pseudomonadales;f\_\_Pseudomonadaceae;g\_\_Stutzerimonas;s\_\_Stutzerimonas stutzeri\_AC | blast | 96.8 | iPHoP-RF;83.10 |
| NC\_019936.1\|provirus\_3233292\_3241441 | RS\_GCF\_021432085.1 | d\_\_Bacteria;p\_\_Pseudomonadota;c\_\_Gammaproteobacteria;o\_\_Pseudomonadales;f\_\_Pseudomonadaceae;g\_\_Stutzerimonas;s\_\_Stutzerimonas kunmingensis\_A | blast | 96.8 | iPHoP-RF;82.50 |
| NC\_019936.1\|provirus\_3233292\_3241441 | RS\_GCF\_900114065.1 | d\_\_Bacteria;p\_\_Pseudomonadota;c\_\_Gammaproteobacteria;o\_\_Pseudomonadales;f\_\_Pseudomonadaceae;g\_\_Stutzerimonas;s\_\_Stutzerimonas kunmingensis | blast | 96.8 | iPHoP-RF;88.20 |
| NC\_019936.1\|provirus\_3233292\_3241441 | RS\_GCF\_000818015.1 | d\_\_Bacteria;p\_\_Pseudomonadota;c\_\_Gammaproteobacteria;o\_\_Pseudomonadales;f\_\_Pseudomonadaceae;g\_\_Stutzerimonas;s\_\_Stutzerimonas balearica | blast | 96.7 | None |
| NC\_019936.1\|provirus\_3233292\_3241441 | RS\_GCF\_003696315.1 | d\_\_Bacteria;p\_\_Pseudomonadota;c\_\_Gammaproteobacteria;o\_\_Pseudomonadales;f\_\_Pseudomonadaceae;g\_\_Stutzerimonas;s\_\_Stutzerimonas songnenensis | blast | 96.5 | iPHoP-RF;93.40 |
| NC\_019936.1\|provirus\_3233292\_3241441 | RS\_GCF\_000327065.1 | d\_\_Bacteria;p\_\_Pseudomonadota;c\_\_Gammaproteobacteria;o\_\_Pseudomonadales;f\_\_Pseudomonadaceae;g\_\_Stutzerimonas;s\_\_Stutzerimonas stutzeri\_AE | blast | 96.4 | iPHoP-RF;92.40 |
| NC\_019936.1\|provirus\_3233292\_3241441 | RS\_GCF\_002909485.1 | d\_\_Bacteria;p\_\_Pseudomonadota;c\_\_Gammaproteobacteria;o\_\_Pseudomonadales;f\_\_Pseudomonadaceae;g\_\_Stutzerimonas;s\_\_Stutzerimonas stutzeri\_AH | blast | 96.3 | iPHoP-RF;75.90 |
| NC\_019936.1\|provirus\_3233292\_3241441 | RS\_GCF\_000661915.1 | d\_\_Bacteria;p\_\_Pseudomonadota;c\_\_Gammaproteobacteria;o\_\_Pseudomonadales;f\_\_Pseudomonadaceae;g\_\_Stutzerimonas;s\_\_Stutzerimonas stutzeri\_A | blast | 94.0 | iPHoP-RF;91.40 |
| NC\_019936.1\|provirus\_3233292\_3241441 | RS\_GCF\_002890795.1 | d\_\_Bacteria;p\_\_Pseudomonadota;c\_\_Gammaproteobacteria;o\_\_Pseudomonadales;f\_\_Pseudomonadaceae;g\_\_Stutzerimonas;s\_\_Stutzerimonas stutzeri\_AA | blast | 92.7 | iPHoP-RF;77.10 |
| NC\_019936.1\|provirus\_3233292\_3241441 | RS\_GCF\_000341615.1 | d\_\_Bacteria;p\_\_Pseudomonadota;c\_\_Gammaproteobacteria;o\_\_Pseudomonadales;f\_\_Pseudomonadaceae;g\_\_Stutzerimonas;s\_\_Stutzerimonas stutzeri\_G | iPHoP-RF | 90.8 | None |
| NC\_019936.1\|provirus\_3233292\_3241441 | RS\_GCF\_024448505.1 | d\_\_Bacteria;p\_\_Pseudomonadota;c\_\_Gammaproteobacteria;o\_\_Pseudomonadales;f\_\_Pseudomonadaceae;g\_\_Stutzerimonas;s\_\_Stutzerimonas degradans | blast | 90.6 | iPHoP-RF;75.20 |
| NC\_019936.1\|provirus\_4296137\_4304306 | RS\_GCF\_003696285.1 | d\_\_Bacteria;p\_\_Pseudomonadota;c\_\_Gammaproteobacteria;o\_\_Pseudomonadales;f\_\_Pseudomonadaceae;g\_\_Stutzerimonas;s\_\_Stutzerimonas nitrititolerans | blast | 96.9 | iPHoP-RF;77.80 |
| NC\_019936.1\|provirus\_4296137\_4304306 | RS\_GCF\_000219605.1 | d\_\_Bacteria;p\_\_Pseudomonadota;c\_\_Gammaproteobacteria;o\_\_Pseudomonadales;f\_\_Pseudomonadaceae;g\_\_Stutzerimonas;s\_\_Stutzerimonas stutzeri | blast | 96.8 | iPHoP-RF;90.50 |
| NC\_019936.1\|provirus\_4296137\_4304306 | RS\_GCF\_015291885.1 | d\_\_Bacteria;p\_\_Pseudomonadota;c\_\_Gammaproteobacteria;o\_\_Pseudomonadales;f\_\_Pseudomonadaceae;g\_\_Stutzerimonas;s\_\_Stutzerimonas stutzeri\_AC | blast | 96.7 | iPHoP-RF;88.20 |
| NC\_019936.1\|provirus\_4296137\_4304306 | RS\_GCF\_000818015.1 | d\_\_Bacteria;p\_\_Pseudomonadota;c\_\_Gammaproteobacteria;o\_\_Pseudomonadales;f\_\_Pseudomonadaceae;g\_\_Stutzerimonas;s\_\_Stutzerimonas balearica | blast | 96.2 | iPHoP-RF;70.40 |
| NC\_019936.1\|provirus\_4296137\_4304306 | RS\_GCF\_003696315.1 | d\_\_Bacteria;p\_\_Pseudomonadota;c\_\_Gammaproteobacteria;o\_\_Pseudomonadales;f\_\_Pseudomonadaceae;g\_\_Stutzerimonas;s\_\_Stutzerimonas songnenensis | blast | 93.9 | iPHoP-RF;80.30 |
| NC\_019936.1\|provirus\_4296137\_4304306 | RS\_GCF\_000327065.1 | d\_\_Bacteria;p\_\_Pseudomonadota;c\_\_Gammaproteobacteria;o\_\_Pseudomonadales;f\_\_Pseudomonadaceae;g\_\_Stutzerimonas;s\_\_Stutzerimonas stutzeri\_AE | blast | 93.6 | iPHoP-RF;74.10 |
| NC\_019936.1\|provirus\_4296137\_4304306 | RS\_GCF\_002909485.1 | d\_\_Bacteria;p\_\_Pseudomonadota;c\_\_Gammaproteobacteria;o\_\_Pseudomonadales;f\_\_Pseudomonadaceae;g\_\_Stutzerimonas;s\_\_Stutzerimonas stutzeri\_AH | blast | 92.5 | iPHoP-RF;85.40 |
| NC\_019936.1\|provirus\_4296137\_4304306 | RS\_GCF\_000661915.1 | d\_\_Bacteria;p\_\_Pseudomonadota;c\_\_Gammaproteobacteria;o\_\_Pseudomonadales;f\_\_Pseudomonadaceae;g\_\_Stutzerimonas;s\_\_Stutzerimonas stutzeri\_A | blast | 91.7 | iPHoP-RF;68.90 |
| NC\_019936.1\|provirus\_4296137\_4304306 | RS\_GCF\_003935375.1 | d\_\_Bacteria;p\_\_Pseudomonadota;c\_\_Gammaproteobacteria;o\_\_Pseudomonadales;f\_\_Pseudomonadaceae;g\_\_Stutzerimonas;s\_\_Stutzerimonas xanthomarina\_A | iPHoP-RF | 91.4 | blast;90.20 |
| NC\_019936.1\|provirus\_4296137\_4304306 | RS\_GCF\_002929225.1 | d\_\_Bacteria;p\_\_Pseudomonadota;c\_\_Gammaproteobacteria;o\_\_Pseudomonadales;f\_\_Pseudomonadaceae;g\_\_Stutzerimonas;s\_\_Stutzerimonas stutzeri\_U | blast | 91.0 | iPHoP-RF;90.10 |
| NC\_019936.1\|provirus\_4296137\_4304306 | GB\_GCA\_003488145.1 | d\_\_Bacteria;p\_\_Pseudomonadota;c\_\_Gammaproteobacteria;o\_\_Pseudomonadales;f\_\_Pseudomonadaceae;g\_\_Stutzerimonas;s\_\_Stutzerimonas sp003488145 | iPHoP-RF | 90.8 | None |
| NC\_019936.1\|provirus\_4296137\_4304306 | RS\_GCF\_000495915.1 | d\_\_Bacteria;p\_\_Pseudomonadota;c\_\_Gammaproteobacteria;o\_\_Pseudomonadales;f\_\_Pseudomonadaceae;g\_\_Stutzerimonas;s\_\_Stutzerimonas chloritidismutans | blast | 90.7 | iPHoP-RF;89.80 |
| NC\_019936.1\|provirus\_4296137\_4304306 | RS\_GCF\_000935215.1 | d\_\_Bacteria;p\_\_Pseudomonadota;c\_\_Gammaproteobacteria;o\_\_Pseudomonadales;f\_\_Pseudomonadaceae;g\_\_Stutzerimonas;s\_\_Stutzerimonas stutzeri\_AD | blast | 90.2 | iPHoP-RF;87.30 |
| NC\_019936.1\|provirus\_4296137\_4304306 | RS\_GCF\_021432085.1 | d\_\_Bacteria;p\_\_Pseudomonadota;c\_\_Gammaproteobacteria;o\_\_Pseudomonadales;f\_\_Pseudomonadaceae;g\_\_Stutzerimonas;s\_\_Stutzerimonas kunmingensis\_A | blast | 90.2 | iPHoP-RF;86.00 |
| NC\_019936.1\|provirus\_4296137\_4304306 | RS\_GCF\_900114065.1 | d\_\_Bacteria;p\_\_Pseudomonadota;c\_\_Gammaproteobacteria;o\_\_Pseudomonadales;f\_\_Pseudomonadaceae;g\_\_Stutzerimonas;s\_\_Stutzerimonas kunmingensis | blast | 90.2 | iPHoP-RF;90.10 |
| NC\_019936.1\|provirus\_4296137\_4304306 | RS\_GCF\_014764705.1 | d\_\_Bacteria;p\_\_Pseudomonadota;c\_\_Gammaproteobacteria;o\_\_Pseudomonadales;f\_\_Pseudomonadaceae;g\_\_Stutzerimonas;s\_\_Stutzerimonas sp002692525 | iPHoP-RF | 90.1 | None |

## 2.22 NC\_021184

##### 2.22.1 Host Genome

**GTDB-Tk taxonomy**:

classification

d\_\_Bacteria;p\_\_Bacillota\_B;c\_\_Desulfotomaculia;o\_\_Desulfotomaculales;f\_\_Desulfallaceae;g\_\_Sporotomaculum;s\_\_Sporotomaculum
gibsoniae

**CheckM2 Summary**:

| total\_contigs | contig\_n50 | completeness | contamination |
| --- | --- | --- | --- |
| 1 | 4855529 | 100 | 3.04 |

**Defense-Finder Systems**:

| sys\_id | type | subtype | sys\_beg | sys\_end | protein\_in\_syst | genes\_count | name\_of\_profiles\_in\_sys |
| --- | --- | --- | --- | --- | --- | --- | --- |
| NC\_021184.1\_SoFic\_11 | SoFIC | SoFic | NC\_021184.1\_113 | NC\_021184.1\_113 | NC\_021184.1\_113 | 1 | SoFic\_\_SoFic |
| NC\_021184.1\_RM\_Type\_I\_21 | RM | RM\_Type\_I | NC\_021184.1\_221 | NC\_021184.1\_224 | NC\_021184.1\_221,NC\_021184.1\_223,NC\_021184.1\_224 | 3 | RM\_\_Type\_I\_MTases\_FAM\_0,RM\_\_Type\_I\_REases\_FAM\_0.einsi\_trimmed,RM\_\_Type\_I\_S\_51 |
| NC\_021184.1\_RM\_Type\_I\_22 | RM | RM\_Type\_I | NC\_021184.1\_242 | NC\_021184.1\_244 | NC\_021184.1\_242,NC\_021184.1\_243,NC\_021184.1\_244 | 3 | RM\_\_Type\_I\_MTases\_FAM\_2,RM\_\_Type\_I\_REases\_FAM\_2.einsi\_trimmed,RM\_\_Type\_I\_S\_52 |
| NC\_021184.1\_MazEF\_8 | MazEF | MazEF | NC\_021184.1\_395 | NC\_021184.1\_396 | NC\_021184.1\_395,NC\_021184.1\_396 | 2 | MazEF\_\_MazE,MazEF\_\_MazF |
| NC\_021184.1\_CAS\_Class1-Subtype-I-B\_29 | Cas | CAS\_Class1-Subtype-I-B | NC\_021184.1\_509 | NC\_021184.1\_516 | NC\_021184.1\_509,NC\_021184.1\_510,NC\_021184.1\_511,NC\_021184.1\_512,NC\_021184.1\_513,NC\_021184.1\_514,NC\_021184.1\_515,NC\_021184.1\_516 | 8 | cas1\_I\_II\_III\_IV\_V\_VI\_7,cas2\_I\_II\_III\_IV\_V\_VI\_3,cas3\_I\_2,cas4\_I\_II\_III\_IV\_V\_VI\_6,cas5\_I-B\_17,cas6\_I-B\_III\_1,cas7b\_I-B\_I-C\_2,cas8b1\_I-B\_9 |
| NC\_021184.1\_CAS\_Class1-Subtype-I-B\_30 | Cas | CAS\_Class1-Subtype-I-B | NC\_021184.1\_770 | NC\_021184.1\_779 | NC\_021184.1\_770,NC\_021184.1\_773,NC\_021184.1\_774,NC\_021184.1\_775,NC\_021184.1\_776,NC\_021184.1\_777,NC\_021184.1\_778,NC\_021184.1\_779 | 8 | cas1\_I\_II\_III\_IV\_V\_VI\_7,cas2\_I\_II\_III\_IV\_V\_VI\_3,cas3\_I\_5,cas4\_I\_II\_III\_IV\_V\_VI\_6,cas5\_I-B\_17,cas6\_I\_II\_III\_IV\_V\_VI\_20,cas7\_I-B\_8,cas8b1\_I-B\_14 |
| NC\_021184.1\_Wadjet\_II\_17 | Wadjet | Wadjet\_II | NC\_021184.1\_925 | NC\_021184.1\_928 | NC\_021184.1\_925,NC\_021184.1\_926,NC\_021184.1\_927,NC\_021184.1\_928 | 4 | Wadjet\_\_JetA\_II,Wadjet\_\_JetB\_II,Wadjet\_\_JetC\_II,Wadjet\_\_JetD\_II |
| NC\_021184.1\_RM\_Type\_I\_23 | RM | RM\_Type\_I | NC\_021184.1\_963 | NC\_021184.1\_967 | NC\_021184.1\_963,NC\_021184.1\_965,NC\_021184.1\_967 | 3 | RM\_\_Type\_I\_MTases\_FAM\_0,RM\_\_Type\_I\_REases\_FAM\_0.einsi\_trimmed,RM\_\_Type\_I\_S\_06 |
| NC\_021184.1\_Kiwa\_6 | Kiwa | Kiwa | NC\_021184.1\_1016 | NC\_021184.1\_1017 | NC\_021184.1\_1016,NC\_021184.1\_1017 | 2 | Kiwa\_\_KwaA,Kiwa\_\_KwaB |
| NC\_021184.1\_PfiAT\_10 | PfiAT | PfiAT | NC\_021184.1\_1033 | NC\_021184.1\_1034 | NC\_021184.1\_1033,NC\_021184.1\_1034 | 2 | PfiAT\_\_PfiA,PfiAT\_\_PfiT |
| NC\_021184.1\_BREX\_I\_4 | BREX | BREX\_I | NC\_021184.1\_1159 | NC\_021184.1\_1165 | NC\_021184.1\_1159,NC\_021184.1\_1160,NC\_021184.1\_1161,NC\_021184.1\_1162,NC\_021184.1\_1165 | 5 | BREX\_\_brxA\_DUF1819,BREX\_\_brxB\_DUF1788,BREX\_\_brxC,BREX\_\_pglX1,BREX\_\_pglZA |
| NC\_021184.1\_AbiE\_1 | AbiE | AbiE | NC\_021184.1\_1208 | NC\_021184.1\_1209 | NC\_021184.1\_1208,NC\_021184.1\_1209 | 2 | AbiEii\_\_AbiEi\_4,AbiEii\_\_AbiEii |
| NC\_021184.1\_RM\_Type\_III\_26 | RM | RM\_Type\_III | NC\_021184.1\_1419 | NC\_021184.1\_1420 | NC\_021184.1\_1419,NC\_021184.1\_1420 | 2 | RM\_Type\_III\_\_Type\_III\_MTases\_FAM\_0,RM\_Type\_III\_\_Type\_III\_REases\_FAM\_0.einsi\_trimmed |
| NC\_021184.1\_Gabija\_5 | Gabija | Gabija | NC\_021184.1\_1422 | NC\_021184.1\_1423 | NC\_021184.1\_1422,NC\_021184.1\_1423 | 2 | Gabija\_\_GajA,Gabija\_\_GajB\_2 |
| NC\_021184.1\_PD-T7-2\_9 | PD-T7-2 | PD-T7-2 | NC\_021184.1\_1424 | NC\_021184.1\_1428 | NC\_021184.1\_1424,NC\_021184.1\_1428 | 2 | PD-T7-2\_\_PD-T7-2\_A,PD-T7-2\_\_PD-T7-2\_B |
| NC\_021184.1\_RM\_Type\_IV\_28 | RM | RM\_Type\_IV | NC\_021184.1\_1656 | NC\_021184.1\_1656 | NC\_021184.1\_1656 | 1 | RM\_Type\_IV\_\_Type\_IV\_05 |
| NC\_021184.1\_RM\_Type\_II\_24 | RM | RM\_Type\_II | NC\_021184.1\_3394 | NC\_021184.1\_3396 | NC\_021184.1\_3394,NC\_021184.1\_3395,NC\_021184.1\_3396 | 3 | RM\_Type\_II\_\_Type\_II\_MTases\_FAM\_16,RM\_Type\_II\_\_Type\_II\_REase01,RM\_Type\_II\_\_Type\_II\_REase17 |
| NC\_021184.1\_Wadjet\_I\_15 | Wadjet | Wadjet\_I | NC\_021184.1\_3691 | NC\_021184.1\_3693 | NC\_021184.1\_3691,NC\_021184.1\_3692,NC\_021184.1\_3693 | 3 | Wadjet\_\_JetA\_I,Wadjet\_\_JetB\_I,Wadjet\_\_JetC\_I |
| NC\_021184.1\_Wadjet\_I\_16 | Wadjet | Wadjet\_I | NC\_021184.1\_3803 | NC\_021184.1\_3806 | NC\_021184.1\_3803,NC\_021184.1\_3804,NC\_021184.1\_3805,NC\_021184.1\_3806 | 4 | Wadjet\_\_JetA\_I,Wadjet\_\_JetB\_I,Wadjet\_\_JetC\_I,Wadjet\_\_JetD\_I |
| NC\_021184.1\_CAS\_Class1-Subtype-I-C\_31 | Cas | CAS\_Class1-Subtype-I-C | NC\_021184.1\_4025 | NC\_021184.1\_4031 | NC\_021184.1\_4025,NC\_021184.1\_4026,NC\_021184.1\_4027,NC\_021184.1\_4028,NC\_021184.1\_4029,NC\_021184.1\_4030,NC\_021184.1\_4031 | 7 | cas1\_I\_II\_III\_IV\_V\_VI\_6,cas2\_I\_II\_III\_IV\_V\_VI\_3,cas3\_I\_5,cas4\_I\_II\_III\_IV\_V\_VI\_6,cas5\_I-C\_11,cas7\_I-C\_7,cas8c\_I-C\_2 |
| NC\_021184.1\_RM\_Type\_III\_27 | RM | RM\_Type\_III | NC\_021184.1\_4091 | NC\_021184.1\_4092 | NC\_021184.1\_4091,NC\_021184.1\_4092 | 2 | RM\_Type\_III\_\_Type\_III\_MTases\_FAM\_0,RM\_Type\_III\_\_Type\_III\_REases\_FAM\_0.einsi\_trimmed |
| NC\_021184.1\_Lamassu-Hypothetical\_7 | Lamassu-Fam | Lamassu-Hypothetical | NC\_021184.1\_4235 | NC\_021184.1\_4237 | NC\_021184.1\_4235,NC\_021184.1\_4236,NC\_021184.1\_4237 | 3 | Lamassu-Fam\_\_LmuA\_effector\_hypothetical,Lamassu-Fam\_\_LmuB\_SMC\_hypothetical,Lamassu-Fam\_\_LmuC\_acc\_hypothetical |
| NC\_021184.1\_AbiH\_2 | AbiH | AbiH | NC\_021184.1\_4301 | NC\_021184.1\_4301 | NC\_021184.1\_4301 | 1 | AbiH\_\_AbiH |
| NC\_021184.1\_RM\_Type\_II\_25 | RM | RM\_Type\_II | NC\_021184.1\_4531 | NC\_021184.1\_4532 | NC\_021184.1\_4531,NC\_021184.1\_4532 | 2 | RM\_Type\_II\_\_Type\_II\_MTases\_FAM\_27,RM\_Type\_II\_\_Type\_II\_REase01 |

**Genomad and CheckV Summary**:

| seq\_name | length | gene\_count | viral\_genes | checkv\_quality | miuvig\_quality | taxonomy | topology | coordinates |
| --- | --- | --- | --- | --- | --- | --- | --- | --- |
| NC\_021184.1\|provirus\_4565036\_4615358 | 50323 | 58 | 23 | Medium-quality | Genome-fragment | Viruses;Duplodnaviria;Heunggongvirae;Uroviricota;Caudoviricetes;; | Provirus | 4565036-4615358 |
| NC\_021184.1\|provirus\_4471243\_4518329 | 47087 | 58 | 22 | High-quality | High-quality | Viruses;Duplodnaviria;Heunggongvirae;Uroviricota;Caudoviricetes;; | Provirus | 4471243-4518329 |
| NC\_021184.1\|provirus\_19778\_35564 | 15787 | 22 | 5 | Low-quality | Genome-fragment | Viruses;Duplodnaviria;Heunggongvirae;Uroviricota;Caudoviricetes;; | Provirus | 19778-35564 |

**Host Genome Ideogram with Phages**:

In
this circular plot, **“c”** indicates the contig, and the
number that follows (e.g., **c1**) represents the contig
number. If multiple contigs are present in the genome, each will be
shown with a distinct label (e.g., **c1**,
**c2**, etc.).

##### 2.22.2 Prophages

**Select prophage to show:**

###### 2.22.2.1 Phage ID: NC\_021184.1|provirus\_4565036\_4615358

**Phage-Host Genome Ideogram:**

**Genes Annotation (geNomad):**

| gene | length | marker | annotation\_accessions | annotation\_description |
| --- | --- | --- | --- | --- |
| NC\_021184.1\|provirus\_4565036\_4615358\_4320 | 594 | NA | NA | NA |
| NC\_021184.1\|provirus\_4565036\_4615358\_4321 | 2580 | NA | NA | NA |
| NC\_021184.1\|provirus\_4565036\_4615358\_4322 | 240 | NA | NA | NA |
| NC\_021184.1\|provirus\_4565036\_4615358\_4323 | 267 | NA | NA | NA |
| NC\_021184.1\|provirus\_4565036\_4615358\_4324 | 759 | NA | NA | NA |
| NC\_021184.1\|provirus\_4565036\_4615358\_4325 | 1011 | NA | NA | NA |
| NC\_021184.1\|provirus\_4565036\_4615358\_4326 | 657 | GENOMAD.055867.VV | COG5632 | N-acetylmuramoyl-L-alanine amidase CwlA |
| NC\_021184.1\|provirus\_4565036\_4615358\_4327 | 423 | GENOMAD.083633.VV | NA | NA |
| NC\_021184.1\|provirus\_4565036\_4615358\_4328 | 1098 | GENOMAD.006687.VV | PF14594;COG4926;TIGR01665 | Siphovirus ReqiPepy6 Gp37-like protein |
| NC\_021184.1\|provirus\_4565036\_4615358\_4329 | 141 | GENOMAD.166984.VP | NA | NA |
| NC\_021184.1\|provirus\_4565036\_4615358\_4330 | 279 | GENOMAD.169028.VV | NA | NA |
| NC\_021184.1\|provirus\_4565036\_4615358\_4331 | 1863 | NA | NA | NA |
| NC\_021184.1\|provirus\_4565036\_4615358\_4332 | 798 | GENOMAD.078804.VV | NA | NA |
| NC\_021184.1\|provirus\_4565036\_4615358\_4333 | 1077 | GENOMAD.007754.VV | PF08931 | Receptor-binding protein of phage tail base-plate Siphoviridae, head |
| NC\_021184.1\|provirus\_4565036\_4615358\_4334 | 852 | GENOMAD.008827.VV | PF20195;COG4722;TIGR01633 | Phage-related protein |
| NC\_021184.1\|provirus\_4565036\_4615358\_4335 | 3420 | GENOMAD.207179.VP | PF05521;COG5283;TIGR01563 | Phage-related tail protein |
| NC\_021184.1\|provirus\_4565036\_4615358\_4336 | 156 | GENOMAD.108553.VV | NA | NA |
| NC\_021184.1\|provirus\_4565036\_4615358\_4337 | 294 | GENOMAD.063000.VV | NA | NA |
| NC\_021184.1\|provirus\_4565036\_4615358\_4338 | 807 | GENOMAD.003250.VV | PF04630;TIGR01603 | phage major tail protein, phi13 family |
| NC\_021184.1\|provirus\_4565036\_4615358\_4339 | 327 | GENOMAD.018077.VV | PF05657 | NA |
| NC\_021184.1\|provirus\_4565036\_4615358\_4340 | 372 | GENOMAD.137411.VP | PF11114;TIGR01725;COG5005 | phage protein, HK97 gp10 family |
| NC\_021184.1\|provirus\_4565036\_4615358\_4341 | 333 | GENOMAD.142742.VP | PF05521;COG5614;TIGR01563 | Bacteriophage head-tail adaptor |
| NC\_021184.1\|provirus\_4565036\_4615358\_4342 | 303 | GENOMAD.041843.VV | PF05135;TIGR01560 | Phage gp6-like head-tail connector protein |
| NC\_021184.1\|provirus\_4565036\_4615358\_4343 | 687 | GENOMAD.111513.VV | NA | NA |
| NC\_021184.1\|provirus\_4565036\_4615358\_4344 | 1185 | GENOMAD.168658.VV | PF05065;PF04586;PF12518;COG4653;TIGR01554;K06904 | Predicted phage phi-C31 gp36 major capsid-like protein |
| NC\_021184.1\|provirus\_4565036\_4615358\_4345 | 699 | GENOMAD.028909.VV | PF05135;TIGR01560 | Phage gp6-like head-tail connector protein |
| NC\_021184.1\|provirus\_4565036\_4615358\_4346 | 1293 | GENOMAD.179073.VP | NA | NA |
| NC\_021184.1\|provirus\_4565036\_4615358\_4347 | 327 | NA | NA | NA |
| NC\_021184.1\|provirus\_4565036\_4615358\_4348 | 312 | NA | NA | NA |
| NC\_021184.1\|provirus\_4565036\_4615358\_4349 | 1557 | GENOMAD.181434.VP | PF20441 | Terminase large subunit, endonuclease domain |
| NC\_021184.1\|provirus\_4565036\_4615358\_4350 | 474 | GENOMAD.168120.VP | PF05119;COG3747;TIGR01558 | Phage terminase, small subunit |
| NC\_021184.1\|provirus\_4565036\_4615358\_4351 | 207 | GENOMAD.225559.VP | NA | NA |
| NC\_021184.1\|provirus\_4565036\_4615358\_4352 | 453 | GENOMAD.191984.VP | PF07128 | NA |
| NC\_021184.1\|provirus\_4565036\_4615358\_4353 | 339 | NA | NA | NA |
| NC\_021184.1\|provirus\_4565036\_4615358\_4354 | 909 | GENOMAD.105515.VV | NA | NA |
| NC\_021184.1\|provirus\_4565036\_4615358\_4355 | 189 | NA | NA | NA |
| NC\_021184.1\|provirus\_4565036\_4615358\_4356 | 765 | GENOMAD.105515.VV | NA | NA |
| NC\_021184.1\|provirus\_4565036\_4615358\_4357 | 1224 | GENOMAD.038338.VV | COG3392 | NA |
| NC\_021184.1\|provirus\_4565036\_4615358\_4358 | 1395 | GENOMAD.005053.VV | NA | NA |
| NC\_021184.1\|provirus\_4565036\_4615358\_4359 | 321 | NA | NA | NA |
| NC\_021184.1\|provirus\_4565036\_4615358\_4360 | 957 | NA | NA | NA |
| NC\_021184.1\|provirus\_4565036\_4615358\_4361 | 759 | NA | NA | NA |
| NC\_021184.1\|provirus\_4565036\_4615358\_4362 | 432 | GENOMAD.159035.VP | PF07374;TIGR01636;COG2739 | phage transcriptional activator, RinA family |
| NC\_021184.1\|provirus\_4565036\_4615358\_4363 | 198 | GENOMAD.212426.VP | NA | NA |
| NC\_021184.1\|provirus\_4565036\_4615358\_4364 | 1353 | GENOMAD.116250.VP | PF00270;COG1111;TIGR04095;K17677 | ERCC4-related helicase |
| NC\_021184.1\|provirus\_4565036\_4615358\_4365 | 282 | GENOMAD.004000.VV | COG1591 | NA |
| NC\_021184.1\|provirus\_4565036\_4615358\_4366 | 2247 | GENOMAD.016441.VV | TIGR01613 | phage/plasmid primase, P4 family, C-terminal domain |
| NC\_021184.1\|provirus\_4565036\_4615358\_4367 | 648 | NA | NA | NA |
| NC\_021184.1\|provirus\_4565036\_4615358\_4368 | 408 | NA | NA | NA |
| NC\_021184.1\|provirus\_4565036\_4615358\_4369 | 258 | GENOMAD.159187.VV | PF14205;TIGR02098;COG1996 | MJ0042 family finger-like domain |
| NC\_021184.1\|provirus\_4565036\_4615358\_4370 | 2301 | GENOMAD.038590.VV | PF00476;PF13482;K02334;COG0749;TIGR01388 | DNA polymerase I - 3’-5’ exonuclease and polymerase domains |
| NC\_021184.1\|provirus\_4565036\_4615358\_4371 | 558 | GENOMAD.039702.VV | PF10991 | NA |
| NC\_021184.1\|provirus\_4565036\_4615358\_4372 | 1122 | GENOMAD.136786.VV | PF10926;TIGR01896;COG2887;K07465 | CRISPR-associated exonuclease Csa1 |
| NC\_021184.1\|provirus\_4565036\_4615358\_4373 | 318 | GENOMAD.222352.VV | NA | NA |
| NC\_021184.1\|provirus\_4565036\_4615358\_4374 | 162 | GENOMAD.116519.VV | NA | NA |
| NC\_021184.1\|provirus\_4565036\_4615358\_4375 | 498 | GENOMAD.204246.VV | PF03333;K01994;TIGR03879 | Adhesin biosynthesis transcription regulatory protein |
| NC\_021184.1\|provirus\_4565036\_4615358\_4376 | 1863 | NA | NA | NA |
| NC\_021184.1\|provirus\_4565036\_4615358\_4377 | 204 | NA | NA | NA |

###### 2.22.2.2 Phage ID: NC\_021184.1|provirus\_4471243\_4518329

**Phage-Host Genome Ideogram:**

**Genes Annotation (geNomad):**

| gene | length | marker | annotation\_accessions | annotation\_description |
| --- | --- | --- | --- | --- |
| NC\_021184.1\|provirus\_4471243\_4518329\_4220 | 1776 | NA | NA | NA |
| NC\_021184.1\|provirus\_4471243\_4518329\_4221 | 456 | NA | NA | NA |
| NC\_021184.1\|provirus\_4471243\_4518329\_4222 | 1011 | NA | NA | NA |
| NC\_021184.1\|provirus\_4471243\_4518329\_4223 | 267 | NA | NA | NA |
| NC\_021184.1\|provirus\_4471243\_4518329\_4224 | 546 | NA | NA | NA |
| NC\_021184.1\|provirus\_4471243\_4518329\_4225 | 276 | GENOMAD.222700.VP | NA | NA |
| NC\_021184.1\|provirus\_4471243\_4518329\_4226 | 1197 | GENOMAD.014802.VV | PF00176;K20093;COG1061;TIGR04095 | Superfamily II DNA or RNA helicase |
| NC\_021184.1\|provirus\_4471243\_4518329\_4227 | 1701 | GENOMAD.016341.VV | PF13479;PF12684;TIGR01618;K07465;COG1468 | phage nucleotide-binding protein |
| NC\_021184.1\|provirus\_4471243\_4518329\_4228 | 453 | GENOMAD.031678.VV | PF05037 | NA |
| NC\_021184.1\|provirus\_4471243\_4518329\_4229 | 1695 | GENOMAD.102034.VP | TIGR01636 | NA |
| NC\_021184.1\|provirus\_4471243\_4518329\_4230 | 654 | GENOMAD.171549.VP | COG3617 | Prophage antirepressor |
| NC\_021184.1\|provirus\_4471243\_4518329\_4231 | 1911 | GENOMAD.024099.VV | NA | NA |
| NC\_021184.1\|provirus\_4471243\_4518329\_4232 | 345 | NA | NA | NA |
| NC\_021184.1\|provirus\_4471243\_4518329\_4233 | 225 | GENOMAD.212426.VP | NA | NA |
| NC\_021184.1\|provirus\_4471243\_4518329\_4234 | 369 | GENOMAD.159035.VP | PF07374;TIGR01636;COG2739 | phage transcriptional activator, RinA family |
| NC\_021184.1\|provirus\_4471243\_4518329\_4235 | 207 | NA | NA | NA |
| NC\_021184.1\|provirus\_4471243\_4518329\_4236 | 399 | NA | NA | NA |
| NC\_021184.1\|provirus\_4471243\_4518329\_4237 | 273 | NA | NA | NA |
| NC\_021184.1\|provirus\_4471243\_4518329\_4238 | 276 | NA | NA | NA |
| NC\_021184.1\|provirus\_4471243\_4518329\_4239 | 1395 | GENOMAD.038338.VV | COG3392 | NA |
| NC\_021184.1\|provirus\_4471243\_4518329\_4240 | 1224 | GENOMAD.038338.VV | COG3392 | NA |
| NC\_021184.1\|provirus\_4471243\_4518329\_4241 | 765 | GENOMAD.105515.VV | NA | NA |
| NC\_021184.1\|provirus\_4471243\_4518329\_4242 | 189 | NA | NA | NA |
| NC\_021184.1\|provirus\_4471243\_4518329\_4243 | 909 | GENOMAD.105515.VV | NA | NA |
| NC\_021184.1\|provirus\_4471243\_4518329\_4244 | 339 | NA | NA | NA |
| NC\_021184.1\|provirus\_4471243\_4518329\_4245 | 432 | GENOMAD.191984.VP | PF07128 | NA |
| NC\_021184.1\|provirus\_4471243\_4518329\_4246 | 207 | GENOMAD.225559.VP | NA | NA |
| NC\_021184.1\|provirus\_4471243\_4518329\_4247 | 474 | GENOMAD.168120.VP | PF05119;COG3747;TIGR01558 | Phage terminase, small subunit |
| NC\_021184.1\|provirus\_4471243\_4518329\_4248 | 1557 | GENOMAD.181434.VP | PF20441 | Terminase large subunit, endonuclease domain |
| NC\_021184.1\|provirus\_4471243\_4518329\_4249 | 312 | NA | NA | NA |
| NC\_021184.1\|provirus\_4471243\_4518329\_4250 | 330 | NA | NA | NA |
| NC\_021184.1\|provirus\_4471243\_4518329\_4251 | 1236 | GENOMAD.179073.VP | NA | NA |
| NC\_021184.1\|provirus\_4471243\_4518329\_4252 | 714 | GENOMAD.028909.VV | PF05135;TIGR01560 | Phage gp6-like head-tail connector protein |
| NC\_021184.1\|provirus\_4471243\_4518329\_4253 | 936 | GENOMAD.168658.VV | PF05065;PF04586;PF12518;COG4653;TIGR01554;K06904 | Predicted phage phi-C31 gp36 major capsid-like protein |
| NC\_021184.1\|provirus\_4471243\_4518329\_4254 | 1821 | NA | NA | NA |
| NC\_021184.1\|provirus\_4471243\_4518329\_4255 | 342 | NA | NA | NA |
| NC\_021184.1\|provirus\_4471243\_4518329\_4256 | 687 | GENOMAD.111513.VV | NA | NA |
| NC\_021184.1\|provirus\_4471243\_4518329\_4257 | 303 | GENOMAD.041843.VV | PF05135;TIGR01560 | Phage gp6-like head-tail connector protein |
| NC\_021184.1\|provirus\_4471243\_4518329\_4258 | 333 | GENOMAD.142742.VP | PF05521;COG5614;TIGR01563 | Bacteriophage head-tail adaptor |
| NC\_021184.1\|provirus\_4471243\_4518329\_4259 | 372 | GENOMAD.137411.VP | PF11114;TIGR01725;COG5005 | phage protein, HK97 gp10 family |
| NC\_021184.1\|provirus\_4471243\_4518329\_4260 | 327 | GENOMAD.018077.VV | PF05657 | NA |
| NC\_021184.1\|provirus\_4471243\_4518329\_4261 | 567 | GENOMAD.003250.VV | PF04630;TIGR01603 | phage major tail protein, phi13 family |
| NC\_021184.1\|provirus\_4471243\_4518329\_4262 | 300 | GENOMAD.063000.VV | NA | NA |
| NC\_021184.1\|provirus\_4471243\_4518329\_4263 | 156 | GENOMAD.108553.VV | NA | NA |
| NC\_021184.1\|provirus\_4471243\_4518329\_4264 | 267 | NA | NA | NA |
| NC\_021184.1\|provirus\_4471243\_4518329\_4265 | 321 | NA | NA | NA |
| NC\_021184.1\|provirus\_4471243\_4518329\_4266 | 3429 | GENOMAD.181727.VP | COG5280 | Phage-related minor tail protein |
| NC\_021184.1\|provirus\_4471243\_4518329\_4267 | 852 | GENOMAD.008827.VV | PF20195;COG4722;TIGR01633 | Phage-related protein |
| NC\_021184.1\|provirus\_4471243\_4518329\_4268 | 1071 | GENOMAD.007754.VV | PF08931 | Receptor-binding protein of phage tail base-plate Siphoviridae, head |
| NC\_021184.1\|provirus\_4471243\_4518329\_4269 | 798 | GENOMAD.078804.VV | NA | NA |
| NC\_021184.1\|provirus\_4471243\_4518329\_4270 | 1863 | NA | NA | NA |
| NC\_021184.1\|provirus\_4471243\_4518329\_4271 | 279 | GENOMAD.169028.VV | NA | NA |
| NC\_021184.1\|provirus\_4471243\_4518329\_4272 | 141 | GENOMAD.166984.VP | NA | NA |
| NC\_021184.1\|provirus\_4471243\_4518329\_4273 | 711 | NA | NA | NA |
| NC\_021184.1\|provirus\_4471243\_4518329\_4274 | 1098 | GENOMAD.006687.VV | PF14594;COG4926;TIGR01665 | Siphovirus ReqiPepy6 Gp37-like protein |
| NC\_021184.1\|provirus\_4471243\_4518329\_4275 | 411 | GENOMAD.083633.VV | NA | NA |
| NC\_021184.1\|provirus\_4471243\_4518329\_4276 | 687 | NA | NA | NA |
| NC\_021184.1\|provirus\_4471243\_4518329\_4277 | 858 | NA | NA | NA |

###### 2.22.2.3 Phage ID: NC\_021184.1|provirus\_19778\_35564

**Phage-Host Genome Ideogram:**

**Genes Annotation (geNomad):**

| gene | length | marker | annotation\_accessions | annotation\_description |
| --- | --- | --- | --- | --- |
| NC\_021184.1\|provirus\_19778\_35564\_13 | 1269 | NA | NA | NA |
| NC\_021184.1\|provirus\_19778\_35564\_14 | 1185 | GENOMAD.016861.VV | PF13671;COG2019;TIGR01359;K13829 | AAA domain |
| NC\_021184.1\|provirus\_19778\_35564\_15 | 456 | NA | NA | NA |
| NC\_021184.1\|provirus\_19778\_35564\_16 | 264 | GENOMAD.221988.VV | PF06806;COG3311;TIGR01764 | Predicted DNA-binding transcriptional regulator AlpA |
| NC\_021184.1\|provirus\_19778\_35564\_17 | 621 | NA | NA | NA |
| NC\_021184.1\|provirus\_19778\_35564\_18 | 1278 | NA | NA | NA |
| NC\_021184.1\|provirus\_19778\_35564\_19 | 240 | NA | NA | NA |
| NC\_021184.1\|provirus\_19778\_35564\_20 | 171 | NA | NA | NA |
| NC\_021184.1\|provirus\_19778\_35564\_21 | 207 | NA | NA | NA |
| NC\_021184.1\|provirus\_19778\_35564\_22 | 357 | GENOMAD.096257.VV | NA | NA |
| NC\_021184.1\|provirus\_19778\_35564\_23 | 189 | NA | NA | NA |
| NC\_021184.1\|provirus\_19778\_35564\_24 | 573 | GENOMAD.103011.VV | PF04586;COG3740;K06904;TIGR01543 | Phage head maturation protease |
| NC\_021184.1\|provirus\_19778\_35564\_25 | 1245 | GENOMAD.113164.VP | PF05521;COG5614;TIGR01563 | Bacteriophage head-tail adaptor |
| NC\_021184.1\|provirus\_19778\_35564\_26 | 2025 | GENOMAD.110013.VP | COG5280 | Phage-related minor tail protein |
| NC\_021184.1\|provirus\_19778\_35564\_27 | 1161 | NA | NA | NA |
| NC\_021184.1\|provirus\_19778\_35564\_28 | 465 | NA | NA | NA |
| NC\_021184.1\|provirus\_19778\_35564\_29 | 291 | NA | NA | NA |
| NC\_021184.1\|provirus\_19778\_35564\_30 | 282 | NA | NA | NA |
| NC\_021184.1\|provirus\_19778\_35564\_31 | 231 | NA | NA | NA |
| NC\_021184.1\|provirus\_19778\_35564\_32 | 141 | NA | NA | NA |
| NC\_021184.1\|provirus\_19778\_35564\_33 | 123 | NA | NA | NA |
| NC\_021184.1\|provirus\_19778\_35564\_34 | 171 | NA | NA | NA |

##### 2.22.3 vOTUs

When more than 1 phage is detected in the host genome, we perform a
clustering step using the tool dRep.

A threshold of 0.95 was applied to the ANI similarity index to define
clusters of virus operational taxonomic units (vOTUs).

**Final cluster designations**

| genomad\_summary$seq\_name | genome | secondary\_cluster | threshold | cluster\_method | comparison\_algorithm | primary\_cluster |
| --- | --- | --- | --- | --- | --- | --- |
| NC\_021184.1\|provirus\_4565036\_4615358 | sequence\_000001.fasta.fasta | 1\_1 | 0.05 | average | ANImf | 1 |
| NC\_021184.1\|provirus\_4471243\_4518329 | sequence\_000002.fasta.fasta | 1\_2 | 0.05 | average | ANImf | 1 |
| NC\_021184.1\|provirus\_19778\_35564 | sequence\_000000.fasta.fasta | 2\_0 | 0.05 | average | ANImf | 2 |

**Primary clustering plot**

##### 2.22.4 Prophage-Host Prediction

Prediction of potential hosts for the prophage contigs.

| virus | host\_genome | host\_taxonomy | main\_method | confidence\_score | additional\_methods |
| --- | --- | --- | --- | --- | --- |
| NC\_021184.1\|provirus\_19778\_35564 | RS\_GCF\_000233715.2 | d\_\_Bacteria;p\_\_Bacillota\_B;c\_\_Desulfotomaculia;o\_\_Desulfotomaculales;f\_\_Desulfallaceae;g\_\_Sporotomaculum;s\_\_Sporotomaculum gibsoniae | blast | 95.5 | iPHoP-RF;84.20 |
| NC\_021184.1\|provirus\_19778\_35564 | RS\_GCF\_008124625.1 | d\_\_Bacteria;p\_\_Bacillota\_B;c\_\_Desulfotomaculia;o\_\_Desulfotomaculales;f\_\_Desulfallaceae;g\_\_Sporotomaculum;s\_\_Sporotomaculum thermosapovorans | iPHoP-RF | 93.4 | blast;90.30 |
| NC\_021184.1\|provirus\_19778\_35564 | GB\_GCA\_016841645.1 | d\_\_Bacteria;p\_\_Bacillota\_B;c\_\_Desulfotomaculia;o\_\_Desulfotomaculales;f\_\_Desulfallaceae;g\_\_Sporotomaculum;s\_\_Sporotomaculum geothermicum\_B | blast | 90.9 | iPHoP-RF;74.80 |
| NC\_021184.1\|provirus\_19778\_35564 | GB\_GCA\_016841205.1 | d\_\_Bacteria;p\_\_Bacillota\_B;c\_\_Desulfotomaculia;o\_\_Desulfotomaculales;f\_\_Desulfallaceae;g\_\_Sporotomaculum;s\_\_Sporotomaculum geothermicum\_A | blast | 90.6 | iPHoP-RF;70.90 |
| NC\_021184.1\|provirus\_19778\_35564 | GB\_GCA\_016841905.1 | d\_\_Bacteria;p\_\_Bacillota\_B;c\_\_Desulfotomaculia;o\_\_Desulfotomaculales;f\_\_Desulfallaceae;g\_\_Sporotomaculum;s\_\_Sporotomaculum geothermicum\_C | blast | 90.4 | iPHoP-RF;51.20 |
| NC\_021184.1\|provirus\_19778\_35564 | RS\_GCF\_009932395.1 | d\_\_Bacteria;p\_\_Bacillota\_B;c\_\_Desulfotomaculia;o\_\_Desulfotomaculales;f\_\_Desulfallaceae;g\_\_Sporotomaculum;s\_\_Sporotomaculum syntrophicum | blast | 90.3 | iPHoP-RF;83.30 |
| NC\_021184.1\|provirus\_4471243\_4518329 | RS\_GCF\_000233715.2 | d\_\_Bacteria;p\_\_Bacillota\_B;c\_\_Desulfotomaculia;o\_\_Desulfotomaculales;f\_\_Desulfallaceae;g\_\_Sporotomaculum;s\_\_Sporotomaculum gibsoniae | blast | 98.1 | iPHoP-RF;73.20 |
| NC\_021184.1\|provirus\_4565036\_4615358 | RS\_GCF\_000233715.2 | d\_\_Bacteria;p\_\_Bacillota\_B;c\_\_Desulfotomaculia;o\_\_Desulfotomaculales;f\_\_Desulfallaceae;g\_\_Sporotomaculum;s\_\_Sporotomaculum gibsoniae | blast | 98.2 | iPHoP-RF;77.30 |
